# The consequences of reciprocally exchanging the genomic sites of Integration Host Factor (IHF) subunit production for subunit stoichiometry and bacterial physiology in *Salmonella enterica* serovar Typhimurium

**DOI:** 10.1101/2021.09.13.459255

**Authors:** German Pozdeev, Michael C Beckett, Aalap Mogre, Nicholas R Thomson, Charles J Dorman

## Abstract

Integration host factor (IHF) is a heterodimeric nucleoid-associated protein that plays roles in bacterial nucleoid architecture and genome-wide gene regulation. The *ihfA* and *ihfB* genes encode the subunits and are located 350 kilobase pairs apart, in the Right replichore of the *Salmonella* chromosome. IHF is composed of one IhfA and one IhfB subunit. Despite this 1:1 stoichiometry, mass spectrometry revealed that IhfB is produced in 2-fold excess over IhfA. We re-engineered *Salmonella* to exchange reciprocally the protein-coding regions of *ihfA* and *ihfB*, such that each relocated protein-encoding region was driven by the expression signals of the other’s gene. Mass spectrometry showed that in this ‘rewired’ strain, IhfA is produced in excess over IhfB, correlating with enhanced stability of the hybrid *ihfB-ihfA* mRNA that was expressed from the *ihfB* promoter. Nevertheless, the rewired strain grew at a similar rate to the wild type, had identical cell morphology, and was similar in competitive fitness. However, compared to the wild type, it was less motile, had growth-phase-specific reductions in SPI-1 and SPI-2 gene expression and was engulfed at a higher rate by RAW macrophage. Our data show that while exchanging the physical locations of its *ihf* genes and the rewiring of their regulatory circuitry are well tolerated in *Salmonella*, genes involved in the production of type 3 secretion systems exhibit dysregulation accompanied by altered phenotypes.

**IMPACT STATEMENT:** Integration Host Factor (IHF) is an abundant nucleoid-associated protein that organises DNA architecturally, influencing gene expression globally in *Salmonella* and other bacteria. IHF is composed of two related, non-identical, subunits, produced by genes that are 350 kilobase pairs apart. Each *ihf* gene has unique expression controls and is embedded in a complex genetic network that supports mRNA translation. Given that the subunits are thought to be required in a 1:1 ratio to form functional IHF, we were surprised by this physical and regulatory separation. We rewired the *Salmonella* genome so that each subunit was produced using the other’s regulatory signals and gene location. This revealed a high degree of tolerance to the effects of this rewiring. However, we discovered that bacterial motility was disrupted, as was the expression of virulence genes that have been acquired by horizontal gene transfer. Proteomic analysis using mass spectroscopy (MS) showed the extent of the alterations to cell composition. Our MS data also demonstrated that the subunits of IHF are not produced in a 1:1 ratio in either the wild type or the rewired strain. We discuss this finding in terms of the ability of each subunit to stabilise its partner.

**DATA SUMMARY:**

1. Whole genome sequence data for strain OrfSwap*^ihfA-ihfB^* are available from the European Nucleotide Archive with accession number ERS4653309.
2. Data from mass spectrometry analyses are available via ProteomeXchange with identifier PXD027465 (login: and password: GVeNIUB2).
3. All supporting data have been provided in the article or through supplementary data files.

**Repositories:** Whole genome sequence data for strain OrfSwap*^ihfA-ihfB^* are available from the European Nucleotide Archive with accession number ERS4653309. Data from mass spectrometry analyses are available via ProteomeXchange with identifier PXD027465 (login: and password: GVeNIUB2).

## INTRODUCTION

The Integration Host Factor, IHF, is a member of the nucleoid-associated protein (NAP) family of nucleic acid binding proteins [1, 2, 3]. NAPs are important contributors to the maintenance of nucleoid architecture and the global control of gene expression, transcriptionally and posttranscriptionally [4–9]. IHF is a heterodimer comprised of the closely related proteins IhfA and IhfB [10–12], encoded by the *ihfA* and *ihfB* genes, respectively, known previously as *himA* and *himD* (*hip*) [12–14]. IHF is an abundant protein, reaching its maximum concentration at the transition from exponential growth to stationary phase in bacteria growing in batch culture [15–17]. It binds to DNA with the consensus sequence: WATCAANNNNTTR (where W is A or T, R is purine, and N is any base) [18, 19]. This protein was discovered originally as a host-encoded factor for integration of bacteriophage lambda into, and excision from, the *Escherichia coli* chromosome [10, 11, 16, 20, 21]. It binds to three sites in the lambda genome, using its DNA bending activity to organise the phage DNA appropriately for site-specific integration by Int, a tyrosine-integrase recombinase. IHF uses two of these three sites to promote the reverse reaction – the Int-catalysed excision of lambda DNA from the bacterial chromosome [16, 22].

Following the discovery that IHF is required for the lambda site-specific recombination system, this NAP was found to play roles in other recombination processes [23–27], in transposition [28, 29], chromosome replication [30, 31], integrative and conjugative element (ICE) transmission [32], plasmid and phage replication [33, 34], plasmid transfer [35], CRISPR-Cas acquisition of novel DNA elements [36, 37] and the regulation of transcription [38–43]. IHF influences the expression of large numbers of genes in several bacterial species [44–50]. Although IHF is not essential for bacterial life, knockout mutations in *ihfA* and *ihfB* result in widespread dysregulation of transcription [44, 46, 47]. IHF seems to be an architectural component of most systems that it influences, playing this role by bending DNA by up to 180° [2, 3, 51–53]. These sharp bends promote contact between DNA sequences located upstream and downstream of the IHF-bound site and between any proteins that may be attached to those flanking sequences. Thus, IHF-dependent nucleoprotein complexes lie at the heart of many of the molecular processes listed above. For example, it can promote contact between DNA-bound transcription factors and RNA polymerase, with positive or negative effects on transcription initiation, depending on the distribution of the participating proteins along the bent DNA [54].

*ihfA* and *ihfB* are located in the Right replichore of the *Salmonella* chromosome, 350 kbp apart (Fig. 1a) [55, 56]. The *ihfB* gene is in the Right macro-domain and *ihfA* is in the Ter macrodomain. Each *ihf* gene is part of a complex operon that includes genes involved in various aspects of bacterial metabolism: *ihfA* has its own promoter but is also transcribed with the *pheST* operon, encoding the phenylalanine tRNA synthetase (Fig. 1b) [15, 57] while *ihfB* has its own promoter and is co-transcribed with *rpsA*, the gene encoding the S1 ribosomal protein (Fig. 1b) [15, 58, 59]. At first glance, this genetic arrangement seems counterintuitive. Why produce the two IHF subunits from genes that are separated physically on the chromosome and are regulated independently? Is the physical location of each gene significant for the life of the bacterium, or might other arrangements work just as well? Several lines of evidence suggest that gene location on bacterial chromosomes is important [60–66]. For example, gene distance from the origin of chromosome replication (*oriC*) influences gene copy number during rapid growth. This is because several rounds of replication are initiated, producing more copies of *oriC*-proximal genes than there are copies of genes close to the terminus of replication, in a phenomenon known as replication-associated gene dosage effects [60, 67, 68]. In addition, the regulatory regimes that control the expression of genes, including genes that encode NAPs, are important for bacterial physiology [69]. Many NAPS exert a pervasive influence on the biology of the bacterium, and if their regulatory patterns are disturbed, significant impacts on cell physiology can result [60].

**Fig 1.**
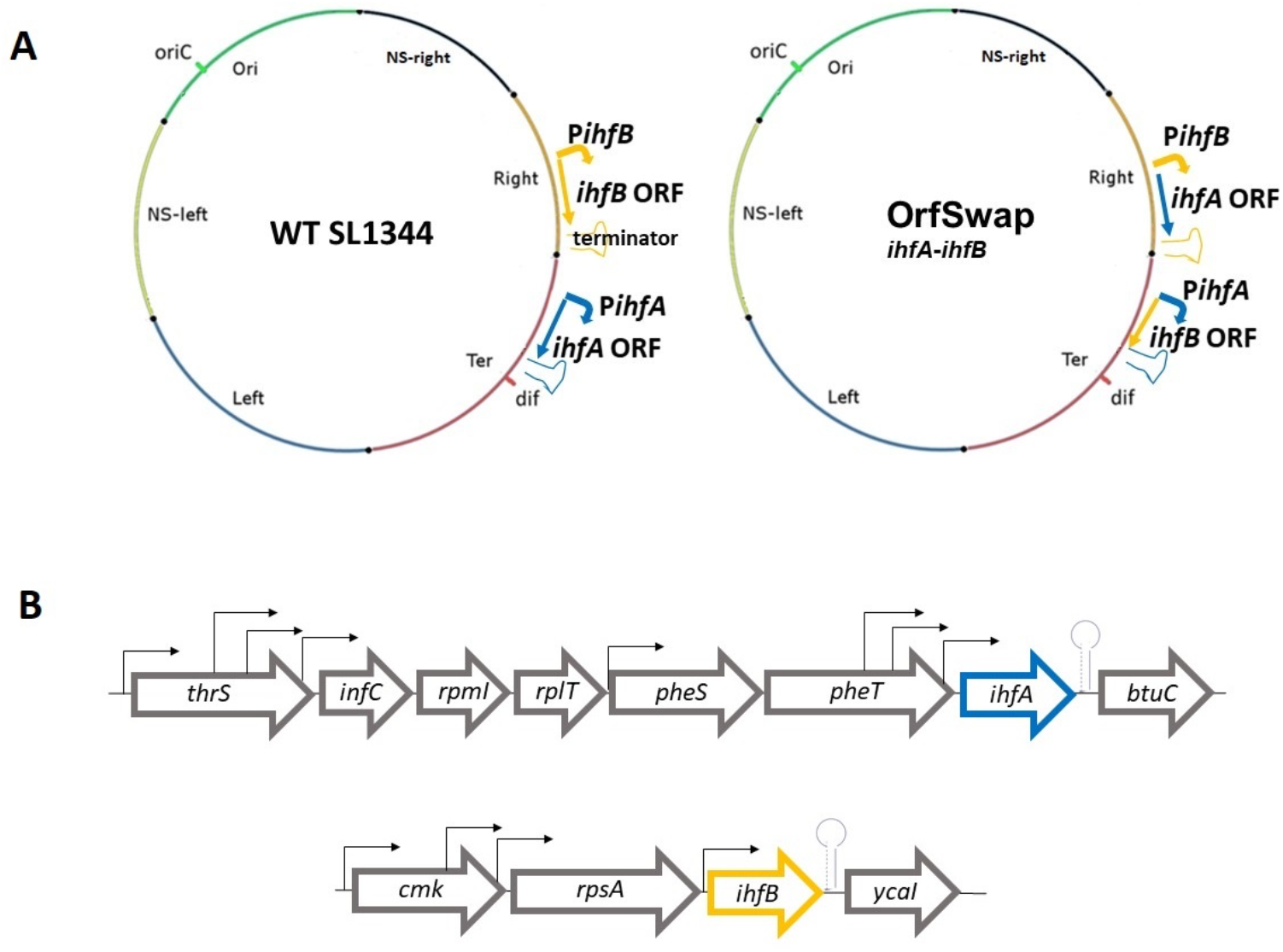
Construction of OrfSwap*^ihfA-ihfB^*, a derivative of wild type *S.* Typhimurium strain SL1344 with reciprocally exchanged *ihfA* and *ihfB* ORFs. A) Simplified maps of the chromosomes of WT SL1344 and its OrfSwap*^ihfA-ihfB^* derivative. The coloured segments represent the macrodomain structure of the chromosome and the locations of the origin of replication (*oriC*) and the *dif* site within the chromosome replication terminus are indicated. The *ihfA* gene (gold) is located in the Ter macrodomain of the WT and the *ihfB* gene (blue) is in the Right macrodomain. The angled arrows represent the transcriptional promoters and the inverted repeat symbols show the positions of the terminators. The straight arrows indicate the direction of transcription and the positions of the open reading frames of each *ihf* gene. B) A summary of the genetic neighbourhoods of the *ihfA* and *ihfB* genes in the wild type strain. The *ihfA* gene is located between the *pheT* and the *btuC* genes and is transcribed from the same DNA strand; *ihfB* is located between the *rpsA* and *ycaI* genes and is transcribed from the same DNA strand. Transcription promoters are illustrated using angled arrows and block arrows show directions of transcription, with stem-loop structures indicating the positions of transcription terminators, based on similarities to the *E. coli* orthologues listed at *Ecocyc.org* for *ihfA* (https://ecocyc.org/ECOLI/substring-search?type=NIL&object=ihfA) and *ihfB* (https://ecocyc.org/gene?orgid=ECOLI&id=EG10441#tab=TU).

Homodimers of IhfA or IhfB bind *in vitro* to the same DNA sequences as heterodimeric IHF, but the protein-DNA complexes are much less stable than those formed by the heterodimer [70]. Data from experiments where *ihfA* or *ihfB* were overexpressed separately *in vivo* revealed a mutual dependency of the subunits for stability: *ihfA* overexpressed in the absence of IhfB results in unstable peptides and *ihfB* overexpressed in the absence of IhfA results in insoluble peptides. In contrast, overexpression of *ihfA* and *ihfB* in the same cell, either from separate, compatible plasmids, or from an *ihfBA* operon on a single plasmid, results in the production of functional IHF [12]. These data suggest that coordinated expression of *ihfA* and *ihfB* is likely to be essential for the production of functional IHF protein, yet these genes are regulated independently and are separated by hundreds of kilobases on the chromosome.

We wished to investigate the significance of *ihfA* and *ihfB* gene position, and the consequences for bacterial physiology, of IHF production from relocated and rewired *ihf* genes. Therefore, we swapped the protein-encoding portion of the *ihfA* and *ihfB* genes, and therefore their physical location, but left the regulatory regions of the original genes intact. We monitored IHF protein production by mass spectroscopy. Our data revealed that the IhfB subunit of IHF is produced at a higher level than IhfA in wildtype *Salmonella* and that this pattern is reversed in the rewired strain. Differences in mRNA stability were found to underlie these differences in IHF subunit production. The reciprocal exchange of the protein coding regions of *ihfA* and *ihfB* altered bacterial motility, the expression of SPI-1 and SPI-2 encoded virulence genes and *Salmonella* pathogenicity. We discuss our findings in the context of chromosome organisation and IHF biology.

## METHODS

### Bacterial strains and culture conditions

The bacterial strains used in this study were derivatives of *S.* Typhimurium strain SL1344 and their details are listed in Table 1; plasmids are described in Table 2. Bacteriophage P22 HT 105/1 *int-*201 was used for generalized transduction during strain construction [71]. Phage lysates were filter-sterilized and stored at 4°C in the dark. Bacterial strains were stored as 35% glycerol stocks at −80°C and freshly streaked on agar plates for each biological replicate. A single colony was used to inoculate 4 ml LB broth and this was then incubated for 18 h. This overnight culture was sub-cultured into fresh 25 ml LB broth normalizing to an OD_600_ of 0.003, unless otherwise stated, and grown to the required growth phase. The standard growth conditions for all bacterial strains were 37°C, 200 rpm, unless otherwise stated.

**Table 1.**
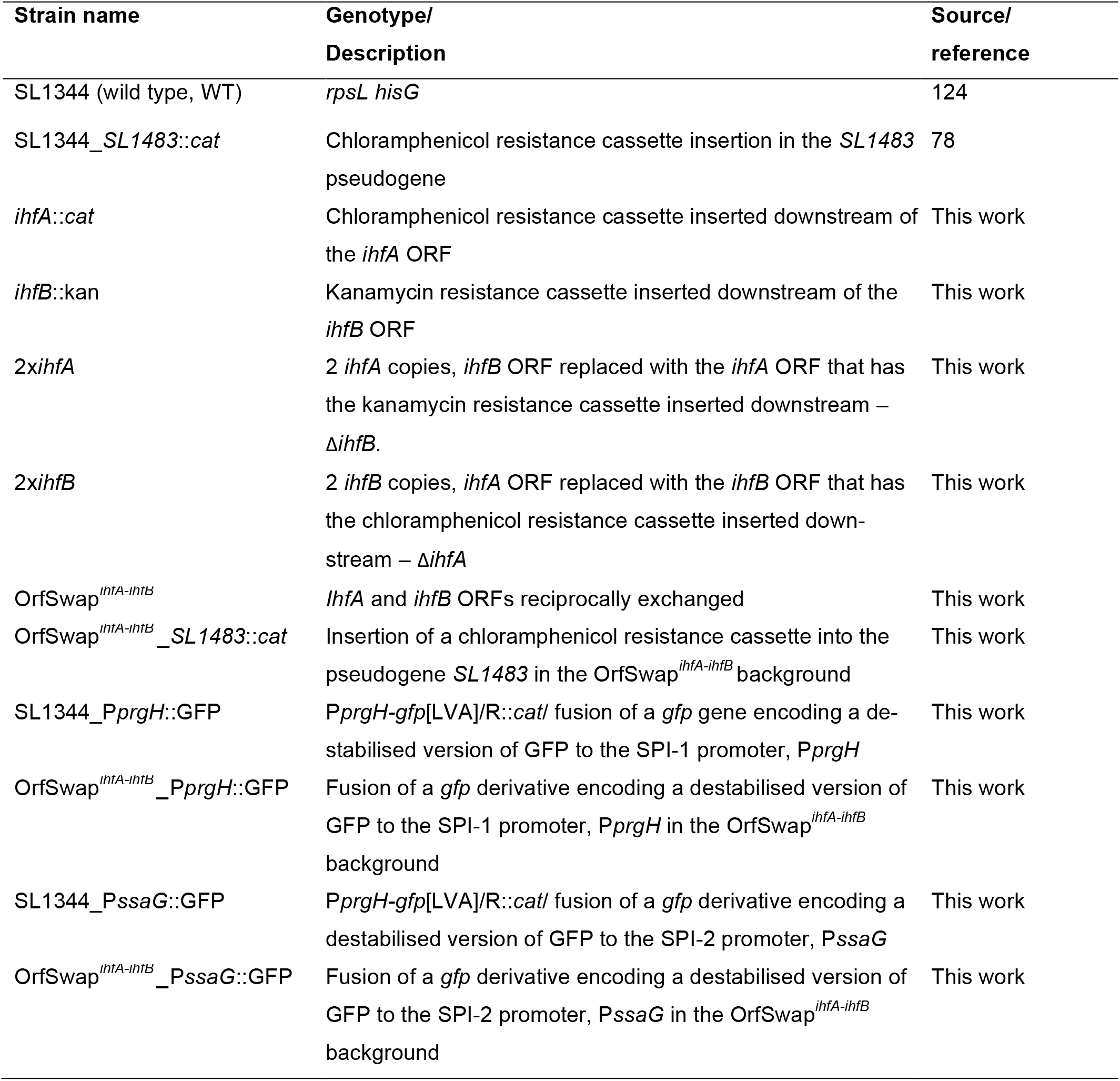
Bacterial strains

**Table 2.** Plasmids used in this study

| Plasmid name | Description | Reference |
| --- | --- | --- |
| pKD3 | Amp<sup>R</sup> (Carb<sup>R</sup>), Cm<sup>R</sup> | 72 |
| pKD4 | Amp<sup>R</sup> (Carb<sup>R</sup>), Kan<sup>R</sup> | 72 |
| pKD46 | Amp<sup>R</sup> (Carb<sup>R</sup>), $\lambda$ Red genes $\gamma$, $\beta$, *exo*<br>under the control of an arabinose induc-<br>ible promoter | 72 |
| pCP20 | Amp<sup>R</sup> (Carb<sup>R</sup>), Cm<sup>R</sup>, FLP recombinase<br>expressing, temperature sensitive repli-<br>con | 77 |

Abbreviations: Amp<sup>R</sup> (Carb<sup>R</sup>), ampicillin (carbenicillin) resistance; Cm<sup>R</sup>, chloramphenicol re- sistance; Kan<sup>R</sup>, kanamycin resistance.

To measure growth characteristics of a bacterial culture, an overnight culture was adjusted to an OD_600_ of 0.003 in 25 ml of fresh LB broth and grown at the standard conditions for 24 h in the appropriate liquid medium. The optical density of the culture at OD_600_ was measured at 1-h intervals for the first 3 hours and then every 30 min until 8 hours; the last reading was taken at 24 h. Measurements were taken using a Thermo Scientific BioMate 3S spectrophotometer with liquid cultures in plastic cuvettes.

The growth characteristics of bacterial cultures in LB broth were also measured by viable counts. The culture was grown in the same way as for spectrophotometry, and an aliquot was taken at 2 h, 4 h, 6 h, 8 h and 24 h, serially diluted and spread on LB agar plates to give between 30 and 300 colonies after overnight incubation at 37°C. The bacterial colony counts were expressed as colony forming units per millilitre (cfu ml^-1^).

### Lambda Red recombineering

The Lambda Red-mediated recombination system was extensively used for strain construction [72]. Normally, most bacteria cannot be transformed with linear exogenous DNA as it gets efficiently eliminated by bacterial restriction-modification (R-M) systems or intracellular exonucleases [73, 74]. One way to aid such transformation is to use strains deficient in some of the R-M components or exonucleases [75]; however, this is not a universal method. Bacteriophages use their own recombination systems to ensure incorporation of their genetic material into bacterial DNA. For example, lambda phage carries the Red system that is composed of three genes: *γ*, *β* and *exo*, that code for Gam, Bet and Exo protein products, respectively. Bet and Exo promote homologous recombination, while Gam inhibits the host RecBCD exonuclease V [76]. Plasmid pKD46 (Table 2) carries the Lambda Red system under the control of an arabinose-inducible promoter. The replication of plasmid pKD46 is temperature-sensitive, allowing to be eliminated by growth at the non-permissive temperature once the recombination process is completed [76].

### Generation of the OrfSwap*^ihfA-ihfB^* strain

To create the OrfSwap*^ihfA-ihfB^* strain, intermediate 2x*ihfA* and 2x*ihfB* strains were first constructed that contained, respectively, two copies of the *ihfA* gene or two copies of the *ihfB* gene (Table 1). Positioning *ihfA* in place of *ihfB* to construct 2x*ihfA* is schematically shown in Fig. S1 to represent the general strategy employed throughout this project. Briefly, a gene to be moved was tagged with an antibiotic resistance gene using plasmid pKD3 (carrying a chloramphenicol resistance cassette) or plasmid pKD4 (carrying a kanamycin resistance cassette) as a template (Table 2). For 2x*ihfA* construction, a pair of primers (infA.cmR.Pfwd and infA.cmR.Prev; Table S1) was used to amplify a chloramphenicol resistance cassette, producing an amplicon that had a first overhang homologous to a region immediately downstream of the gene of interest (*ihfA*). All insertions of the antibiotic resistance cassettes and amplicons in the subsequent steps were typically made three nucleotides downstream of the stop codon to avoid disrupting the terminator sequence and excluding other regulatory regions. The second overhang was designed so as to delete 2-4 nucleotides from the target region to make insertion more efficient. PCR was carried out using high-fidelity Phusion DNA Polymerase. The resulting amplicon was purified with the Qiagen PCR purification kit and transformed by electroporation into a strain harbouring pKD46. This strain was grown with arabinose at 30°C to drive the expression of the Lambda Red system encoded on pKD46 (Table 2) [72]. The integrity of successful transformant colonies was confirmed by PCR with “confirmation” primers and sequenced by Sanger sequencing. Taking 2x*ihfA* as an example, the *ihfA*::Cm^R^ construct was similarly amplified with primers that had overhangs homologous to regions upstream and downstream of the *ihfB* ORF (infB.int.Pfwd and infB.int.Prev; Table S1). The amplicon was purified and transformed into the WT strain harbouring pKD46, successful transformations were confirmed as described above. Alongside the 2x*ihfA*, a 2x*ihfB* strain was constructed using a similar approach but using a kanamycin resistance cassette for selection. A P22 lysate of the 2x*ihfB* was used to transduce the *ihfB*::kan^R^ into the 2x*ihfA* background to yield the OrfSwap*^ihfA-ihfB^*. Antibiotic resistance cassettes were removed by the FLP-mediated site-specific recombination with aid of pCP20 (Table 2) [77]. The resulting strain had its *ihfA* ORF under the control of the regulatory region of *ihfB* and the *ihfB* ORF under the control of the regulatory region of *ihfA* (Table 1).

IHF plays a role in tyrosine-integrase-mediated site-specific recombination [13, 20]. Therefore, attempts to remove the antibiotic resistance cassettes flanked by FRT sites by FLP-integrase-mediated site-specific recombination, in intermediate strains where the IHF heterodimer was not present (2x*ihfA* and 2x*ihfB*), failed. Hence, it should be noted that the 2x strains final versions of these strains contain antibiotic resistance cassettes, while the cassettes were removed from the final OrfSwap*^ihfA-ihfB^* strain to avoid potential fitness costs associated with carrying those cassettes.

### Bacterial motility assays

Assays were carried out precisely as described to minimize technical variation: 0.3% LB agar was melted in a 100 ml bottle in a Tyndall steamer for 50 min, allowed to cool in a 55°C water bath for 20 min, six plates were poured and left to dry near a Bunsen flame for 25 min. 1 µl of bacterial overnight culture was pipetted under the agar surface, with two inocula per plate. Plates were placed in a 37°C incubator without stacking to ensure equal oxygen access. After 5 h, the diameters of the resulting swarm zones were measured and expressed as the ratio of the WT zone to that of the mutant.

### Competitive fitness assays

Flasks of broth were inoculated with the pair of competing bacterial strains in a 1:1 ratio. Derivatives of each competitor were constructed that carried a chloramphenicol acetyl transferase (*cat*) gene cassette within the transcriptionally silent pseudogene *SL1483*. This *cat* insertion is known to be neutral in its effects on bacterial fitness [78] and allows the marked strain to be distinguished from its unmarked competitor. Competitions were run, in which wild type SL1344 was the marked strain or in which OrfSwap*^ihfA-ihfB^* was the marked strain. Strains to be competed were pre-conditioned in separate 25 ml cultures for 24 h without antibiotics. Then 10^5^ cells of each strain were mixed in 25 ml of fresh LB broth and grown as a mixed culture for another 24 h. The number of colony forming units was determined by plating the mixture on chloramphenicol-containing plates and on plates with no antibiotic at T=0 h and T=24 h. WT SL1344 was competed against its SL1344 derivative OrfSwap*^ihfA-ihfB^ SL1483::cat* and, as a control, SL1344_*SL1483::cat* was competed against the OrfSwap*^ihfA-ihfB^* strain. Competitive fitness was calculated according to the formula:

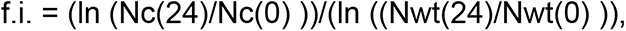

where Nc(0) and Nc(24) are the initial and final counts of a competitor and Nwt(0) and Nwt(24) are initial and final counts of the WT. Competitor is a strain other than the WT. F.i.<1 means that the competitor is less fit than the WT, f.i.>1 indicates the opposite.

### DNA isolation for whole genome sequencing

To obtain high-quality chromosomal DNA for whole genome sequencing, a basic phenol-chloroform method was used [79]. Two ml of an overnight culture were centrifuged at 16000 x g for 1 min to harvest cells and the cell pellet was resuspended in 400 µl of TE buffer pH 8 (100 mM Tris-HCl pH 8.0, 10 mM EDTA pH 8.0 (BDH, Poole, England)). 1% SDS and 2 mg/ml proteinase K were added and incubated for 2 h at 37°C to complete lysis. DNA was isolated by the addition of 1 volume phenol pH 8.0: chloroform: isoamyl alcohol (AppliChem, Darmstadt, Germany) (25:24:1) to the reaction in a phase lock tube. After thorough mixing, the sample was centrifuged at 16000 x g for 15 min at 4 °C to separate the aqueous phase. The upper aqueous layer containing DNA was collected and the phenol: chloroform extraction was repeated two more times. To remove contaminants and to precipitate the DNA, sodium acetate pH 5.2 at 0.3 M and isopropanol at 60% of the final volume were added and kept for 1 h at −20°C. DNA was pelleted by centrifugation at 16000 x g for 15 min at 4°C. The DNA pellet was washed with 70% ethanol, dried at 37°C until translucent and resuspended in 100 µl TE pH 8.0. The DNA concentration was determined spectrophotometrically following removal of RNA contamination by treatment with 100 mg/ml RNase A for 30 min at 37°C. Phenol-chloroform extraction was then performed as above. To precipitate DNA, 0.5 M of ammonium acetate (Merck, Darmstadt, Germany) and a half volume of isopropanol were added and incubated for 2 h at −20°C. DNA was pelleted by centrifugation at 16000 x g for 15 min at 4°C. The DNA pellet was washed twice with 70% ethanol, dried at 37°C until translucent and resuspended in 50 µl water. The sample was run on an agarose gel to check for degradation. The concentration of DNA extracted was determined by measuring absorbance at 260 nm on a DeNovix DS-11 spectrophotometer (Wilmington, Delaware, USA). The shape of the absorbance curve was ensured to have a clear peak at 260 nm. The purity of samples was assessed by the ratio of A_260_/A_280_ – a measure of protein and phenol contamination and A_260_/A_230_ – a measure of contaminants such as EDTA, where both should be as close as possible to 2. Only high-quality samples were chosen for further work.

### Whole genome sequencing

Whole genome sequencing, using Illumina next generation sequencing technology (Sanger Institute, Hinxton, Cambridge, UK), was performed on final versions of the constructed strains to ensure that no compensatory mutations had been introduced. SNP calling was performed by aligning reads to the reference SL1344 sequence NC_016810.1 using Breseq software [80]. Whole genome sequence data for strain OrfSwap*^ihfA-ihfB^* are available from the European Nucleotide Archive with accession number ERS4653309.

### Mammalian cell culture conditions

RAW264.7 murine macrophages were maintained in Dulbecco’s Modified Eagle’s Medium (DMEM), (Sigma, catalogue number D6429) supplemented with 10% foetal bovine serum (FBS) in a humidified 37°C, 5% CO_2_ tissue-culture incubator grown in 75 cm^3^ tissue-culture flasks. When approximately 80% confluent growth was achieved, cells were split to a fresh flask. Cells within the 9-16 passage number range were used for infections. All media and PBS used for cell culture were pre-warmed to 37°C. To split cells, old DMEM was removed and the monolayer was rinsed with 10 ml of sterile PBS. Fresh DMEM (10 ml) was pipetted into the flask and the monolayer was scraped gently with a cell scraper to dislodge the cells. Scraped cells were centrifuged at 450 x g for 5 min in an Eppendorf 5810R centrifuge and the cell pellet was resuspended in 5 ml DMEM+FBS. One ml of the cell suspension was added to 14 ml of fresh DMEM+FBS in a 75 cm^3^ flask, gently rocked to mix and incubated at 37°C, 5% CO_2_. To seed cells for infection, cells were treated as for splitting. After resuspension in 5 ml DMEM+FBS, viable cells were counted on a haemocytometer using trypan blue exclusion dye. A 24-well tissue culture plate was filled with 500 µl DMEM+FBS. 1.5×10^5^ cells were added to each well, gently rocked to mix and incubated at 37°C, 5% CO_2_ for 24 h.

### Protein sample preparation for Mass Spectrometry

An overnight culture was adjusted to an OD_600_ of 0.003 in 25 ml of LB broth and grown to the required growth stage. A volume of cells, equivalent to a cell density of 1  ml of culture at an OD_600_ of 2, was harvested by centrifugation at 3220 x g, 10 min, 4°C. The cell pellet was resuspended in 600 µl of 8 M urea and moved into a sonication tube. The sample was sonicated on ice with 10 rounds of sonication pulses at an amplitude of 10 µm for 20 s with 20 s resting periods between. Resulting lysate was transferred into a fresh Eppendorf tube and centrifuged at 1500 x g for 15 min at 4°C to remove cell debris. 400 µl of a supernatant was transferred into a fresh tube and centrifuged at 9300 x g for 10 min at 4°C. 200 µl of a supernatant was transferred into a fresh tube and kept at −20°C. The protein concentration was determined by Bradford assay [79].

To reduce disulphide bridges, 100 µg of protein in 50 µl total volume was treated with 5 mM of dithiothreitol (DTT) shaking at 350 rpm for 10 min at 60°C. To prevent cysteine bridges re-forming, exposed cysteine residues were alkylated by adding 10 mM of iodoacetamide (IAA) for 30 min in the dark. To quench the reaction, dilute 8 M urea and neutralize pH in order to enable trypsinisation, 150 µl of 200 mM of ammonium bicarbonate was added. 20 µl of the sample preparation was transferred into a fresh tube, 2 µg of trypsin (Roche, Mannheim, Germany) was added and incubated at 350 rpm for 18 h at 37°C. Trypsin digestion was stopped by adding 0.1% trifluoroacetic acid (TFA). Protein samples for mass spectrometry were purified and concentrated using ZipTip_C-18_ (Merck, Millipore, Darmstadt, Germany).

### Mass Spectrometry and data analysis

Complex trypsin-digested peptide mix was first separated by 1 h high performance liquid chromatography (HPLC) gradient using reverse phase C18 columns on an Dionex Ultimate 3000 UPLC (ThermoFisher Scientific) followed by a run on Q-Exactive (ThermoFisher Scientific) mass spectrometer located at the Mass Spectrometry Core, Conway Institute, UCD, Ireland.

Mass spectrometry output files in the RAW format were loaded into the MaxQuant version 1.6.10.43. The RAW files were searched against the *Salmonella enterica* serovar Typhimurium str. SL1344 complete proteome sequence obtained from UniProt (proteome ID UP000008962). Carbamidomethylation of cysteine was chosen as a variable modification, while oxidation of methionine and N-terminus acetylation were chosen as fixed modification. Trypsin digestion was selected with a maximum of two missed cleavages. Label Free Quantification (Fast LFQ) mode was selected in group-specific parameters and iBAQ quantification was added in global parameters. Other settings were default. The search was performed using the Andromeda peptide search engine integrated into the MaxQuant.

### Protein ratio determination

Intensity-based absolute quantification values (iBAQ) were used to calculate the ratios of IhfA to IhfB. The iBAQ intensity values are generated by the MaxQuant algorithm that calculates the sum of all peptide peak intensities divided by the number of theoretically observable peptides during tryptic digestion [81]. The protein entries corresponding to IhfA and IhfB were identified in the proteinGroups.txt file. IhfA to IhfB ratios were determined by dividing the corresponding iBAQ intensity values.

### Statistical analysis of mass spectrometry

Label-free quantification (LFQ) intensity values were used to compare the protein abundance between the samples (Table S2). These values were obtained by the delayed normalisation algorithm that eliminates differences that arise from separate sample processing and the extraction of maximum peptide ratio information by using only common peptides for the pair-wise sample comparison [82]. Statistical analysis was performed in Perseus (version 1.6.10.50). Potential contaminants, proteins only identified by site and reverse hits were filtered out. Annotations were downloaded from annotations.perseus-framework.org. The LFQ intensity values were log_2_-transformed, and the samples were grouped to account for the experimental design. A protein was considered to be present for the purpose of the analysis if it was detected in at least two samples in any of the sample groups. A two-sided Student’s T-test was performed on the data prior to imputation, using a false discovery rate (FDR) = 0.01 and S_0_ = 1. Principal component analysis (PCA) of the dataset was carried out to ensure that the MS samples cluster according to the strain. The proteins that were differentially expressed at a statistically significant level were filtered out. Hierarchical clustering was performed on these proteins (the log_2_-transformed LFQ values were normalized by Z-score) to distinguish between up and downregulated proteins. For all other proteins, missing LFQ intensity values were replaced by imputation from the normal distribution. Two-sided Student’s T-test was performed on these imputed data using the same FDR and S_0_ parameters. Manual review was performed on proteins that were found to be differentially expressed after imputation to assess the effect of imputation on statistical significance. This was necessary because among proteins that were detected in at least two samples in one strain and only in one sample in the other strain and where detected LFQ values were similar, imputation frequently resulted in the creation of false conclusions about statistical significance. As a result, statistically significant up- and downregulated proteins, as well as the unique proteins, were deemed to be differentially expressed.

## RESULTS

### Constructing OrfSwap*^ihfA-ihfB^*, a derivative of *S*. Typhimurium with the protein coding regions of its *ihf* genes exchanged reciprocally

The *ihfA* and *ihfB* genes are located on the positive strand of the *S*. Typhimurium SL1344 chromosome and are separated by approximately 350 kbp [56, 83]. The *ihfA* and *ihfB* genes in *Salmonella* and *E. coli* are at different locations on the chromosome and possess their own regulatory regions (Fig. 1b) [15, 59]. A derivative of the *S.* Typhimurium wildtype strain SL1344 was constructed in which the complete protein-coding regions (CDS) of the genes were exchanged reciprocally. Here, the protein-coding region of *ihfA* and *ihfB* were brought under the transcriptional control of each other’s 5’ and 3’ regulatory regions (Fig. 1a). These CDS exchanges resulted in the production of hybrid mRNA species consisting of the 5’ and 3’ untranslated regions (UTRs) of one gene and an mRNA corresponding to the CDS of the other. In addition, the IhfA protein was now produced from a transcript expressed at the genomic location previously occupied by the wild type *ihfB* gene, and *ihfB* was transcribed from the erstwhile genomic location of *ihfA* (Fig. 1). This strain was designated OrfSwap*^ihfA-ihfB^* (Fig. 1a; Table 1). The process of strain construction, which relied on the Lambda Red recombination system [72], is described in detail in METHODS. By whole genome sequencing the OrfSwap*^ihfA-ihfB^* strain *ihfA* and *ihfB* genes were identical to the parental strain, but also carried harmless SNPs/ rearrangements in the shufflon region. The SL1344 parental strain used in this study has two previously described SNPs, in *manX* (E95V) and *menC* (L148L), compared to the serotype reference strain genome [64].

### Expression patterns of the rearranged *ihfA* and *ihfB* genes

The effects of these gene rearrangements on *ihf* transcription were tested by RT-PCR at different stages of growth using pairs of gene-specific primers: SL_ihfA_qPCR_Pf and SL_ihfA_qPCR_Pr for *ihfA*, and SL_2ihfB_qPCR_Pf and SL_2ihfB_qPCR_Prev for *ihfB* (Table S1). Transcript levels were calculated relative to *hemX* – a gene whose expression does not change under the growth conditions used here [83].

In WT *S.* Typhimurium strain SL1344 (SL1344, Table 1), the pattern of the *ihfA* and *ihfB* mRNA expression resembled that of IHF protein production in *E. coli* [17, 84, 85], peaking at the 7-h time point, corresponding to the early stationary phase of growth (Fig. 2). In the OrfSwap*^ihfA-ihfB^* strain, *ihfA* gene expression dropped at 5 h and 7 h compared to SL1344 but increased at 2 h, while *ihfB* gene expression matched that in the SL1344, except for a decrease at 7 h. Note that in the OrfSwap*^ihfA-ihfB^* strain, *ihfA* gene expression refers to the levels of the 5’-*ihfB*[UTR]-*ihfA*[ORF]-3’ hybrid mRNA and *ihfB* gene expression refers to the levels of the 5’-*ihfA*[UTR]-*ihfB*[ORF]-3’ hybrid mRNA, since qPCR primers are CDS-specific. These results indicated that the CDS of *ihfA* and *ihfB* under the control of each other’s 5’ and 3’ non-translated regulatory regions did not result in exchange in gene expression characteristics. Instead, they had acquired unique, rescheduled gene expression patterns.

**Fig. 2.**
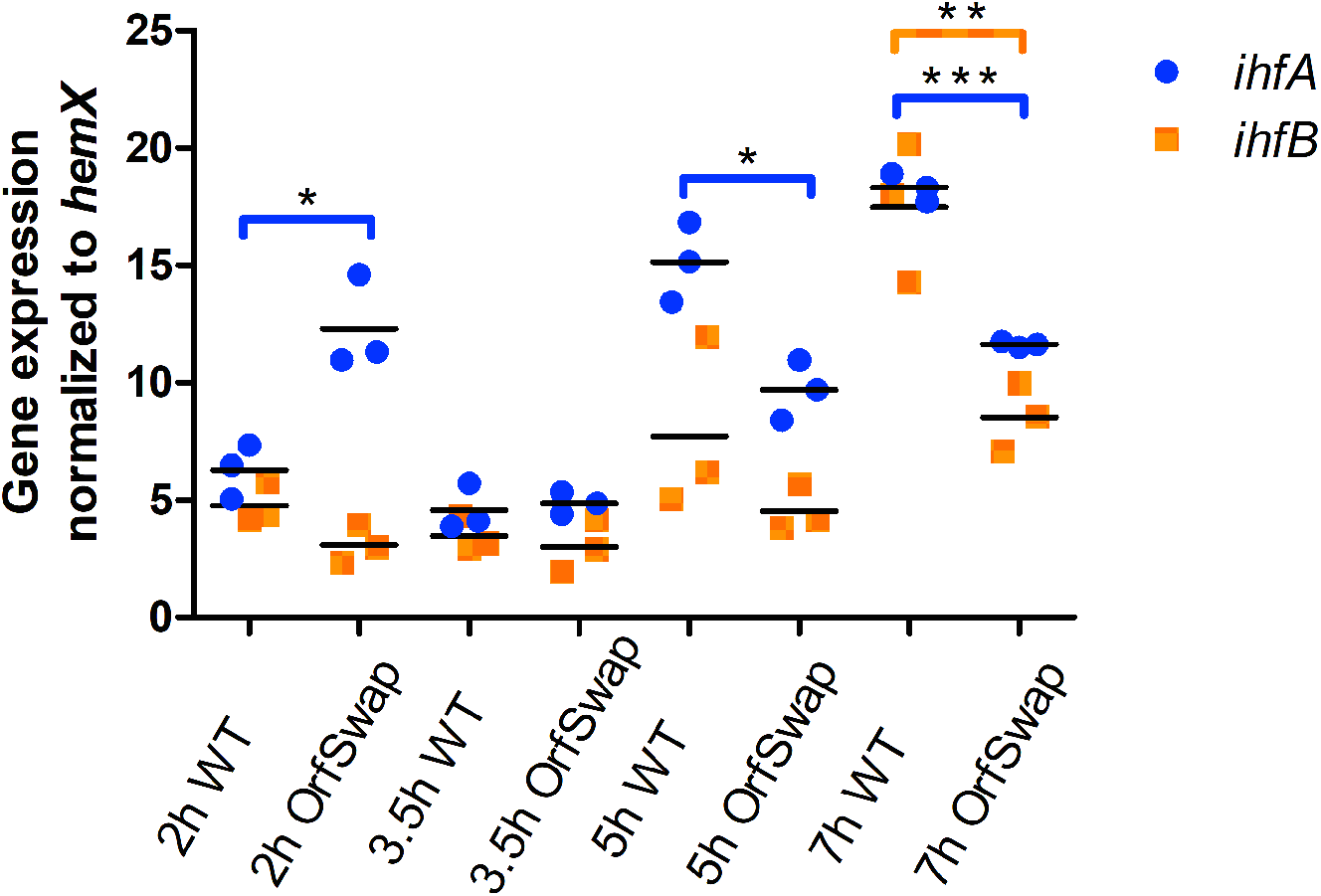
Expression patterns of the relocated *ihfA* and *ihfB* genes. Cells were grown in LB broth at 37°C with aeration and samples were taken at 2 h, 3.5 h, 5 h and 7 h representing lag, exponential, exponential-stationary transition and early stationary growth phases, respectively. Expression of the *ihfA* gene peaked at later time points (5 h and 7 h) in the WT (SL1344) and at 2 h in the OrfSwap*^ihfA-ihfB^* strain. Expression of *ihfB* was similar to that of *ihfA* and increased as bacteria grew. The *ihfB* gene was transcribed at a higher level in the WT than in the OrfSwap*^ihfA-ihfB^* strain at 5 h and 7 h. Each plot is the result of three biological replicates; statistical significance was found by Student’s unpaired T-test, where P<0.05.

### Assessing the impacts of the *ihfA* and *ihfB* rearrangements on downstream gene expression

Experiments were conducted to investigate the effect of the *ihf* gene relocation on the expression of downstream genes. RNA-seq data show no read-through transcription from *ihfA* or *ihfB* into downstream genes in SL1344 [83], indicating that the *ihfA and ihfB* gene transcription terminators are effective in SL1344.

We investigated gene expression of the genes immediately downstream from the repositioned *ihfA* and *ihfB* genes in the OrfSwap*^ihfA-ihfB^* strain by RT-qPCR. The *btuC* gene, encoding the permease of the vitamin B12 transport system, is located 3’ to *ihfA* in the WT and 3’ to the 5’-*ihfA*[UTR]-*ihfB*[ORF]-3’ hybrid locus in the OrfSwap*^ihfA-ihfB^* strain. The *ycaI* gene, encoding a putative competence-related protein, is located 3’ to *ihfB* in SL1344 and 3’ to the 5’-*ihfB*[UTR]-*ihfA*[ORF]-3’ hybrid locus in the OrfSwap*^ihfA-ihfB^* strain. Expression of *btuC* and *ycaI* was measured by RT-qPCR using the specific primer pairs (SL_btuC_qPCR_Pf and SL_btuC_qPCR_Pr; SL_ycaI_qPCR_Pf and SL_ycaI_qPCR_Pr, Table S1) for *btuC* and *ycaI*, respectively. Transcription of both downstream genes had increased in the OrfSwap*^ihfA-ihfB^*, compared to SL1344 (Fig. S3). With the exception of *btuC* at the 2-h time-point (when no difference was found) small-but-statistically-significant increases in downstream gene expression were detected. The fold changes in gene expression were well below 1, relative to *hemX* expression, suggesting that they were unlikely to be physiologically important.

### IhfA and IhfB production is unequal in both the SL1344 and the OrfSwap*^ihfA-ihfB^* strains

The accurate absolute molar quantities of IHF subunits in SL1344 and the OrfSwap*^ihfA-ihfB^* strain proteomes were compared by mass spectrometry (MS; see METHODS) [86]. Data from MS analyses are summarised in Table 2. Samples for protein preparation were taken from LB cultures at the 7-h timepoint of the growth cycle, corresponding to the early stationary phase of the growth cycle (the point where IHF production is at its maximum). It is also the time point at which *ihfA* and *ihfB* gene expression had been measured by RTqPCR (Fig. 2). IhfB to IhfA ratios were determined by dividing the corresponding iBAQ intensity values. This was done separately for each biological replicate, with the resulting ratios being averaged and plotted (Fig. 3). The results showed that in SL1344, the IhfB subunit was present at approximately twice the molar quantity of IhfA protein. However, in the OrfSwap*^ihfA-ihfB^* strain, the IhfA protein was produced at a molar quantity that exceeded that of the untagged IhfB subunit, a reversal of the pattern seen in SL1344 (Fig. 3).

**Fig. 3.**
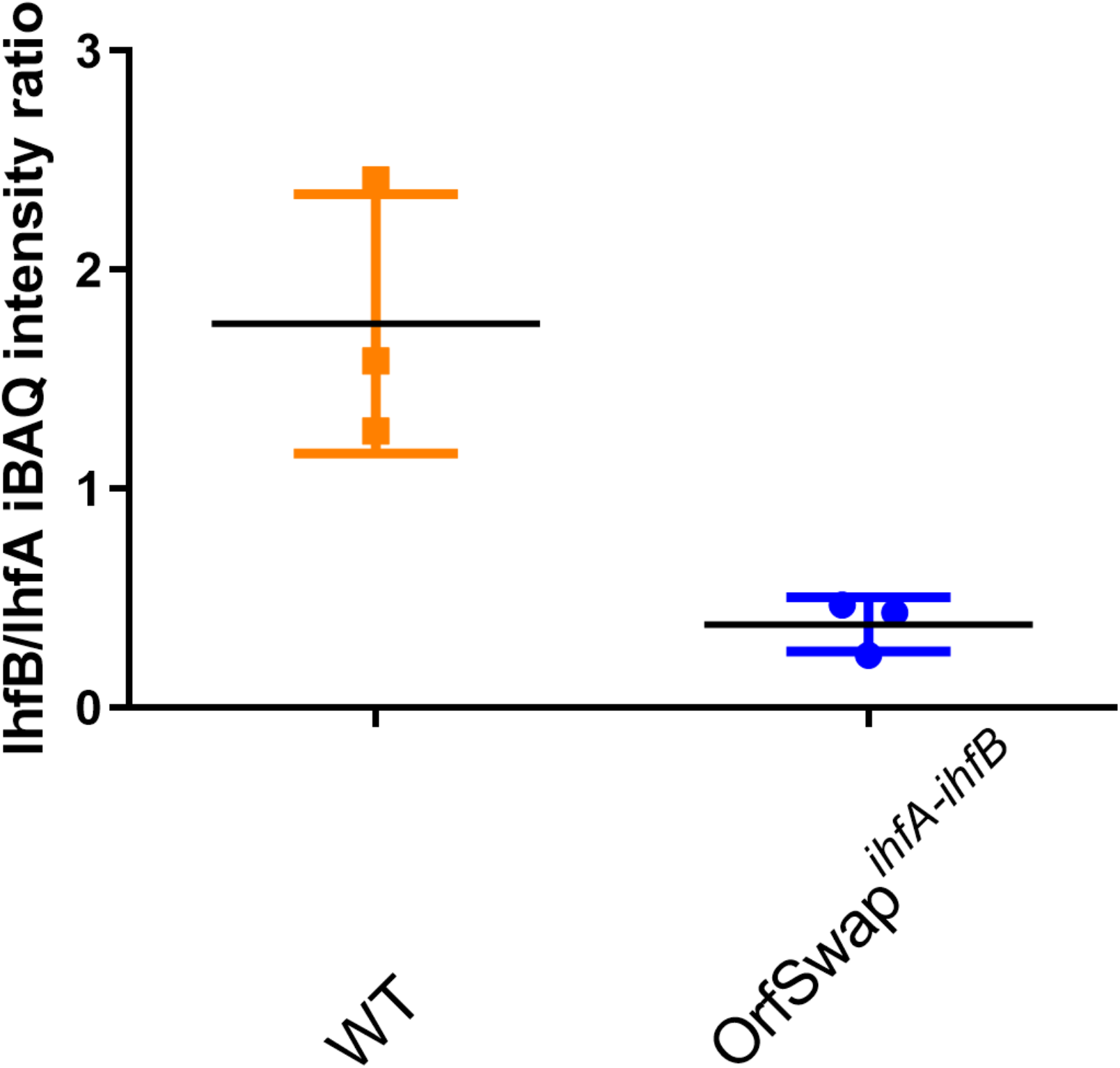
Stoichiometery of IHF subunit production in the wild type (SL1344) and the OrfSwap*^ihfA-ihfB^* strain. Mass spectrometry was performed with the WT and OrfSwap*^ihfA-ihfB^* strains at 7 h, corresponding to the early stationary phase of growth. iBAQ intensity values of the MaxQuant analysis output were used to determine the ratios of IhfB to IhfA subunits in each strain.

### Comparing the proteomes of SL1344 and OrfSwap*^ihfA-ihfB^* strains

A total of 884 proteins were detected by MS in at least two samples of either strain. The full dataset has been uploaded to ProteomeXchange (identifier PXD027465). There were 213 proteins that were downregulated significantly in the OrfSwap*^ihfA-ihfB^* strain, compared to SL1344, and 83 that were significantly upregulated (Table S2). In light of the reduced quantity of IhfB relative to IhfA in the OrfSwap*^ihfA-ihfB^* strain, it was interesting to note that the B subunit of IHF’s paralogue HU was also present in reduced molar quantities.

Of the 126 proteins significantly downregulated in the OrfSwap*^ihfA-ihfB^* strain, 22 were involved in translation, 53 were involved in metabolic pathways, 16 proteins were associated with the cell envelope and transport, 12 proteins were involved in genetic information processing, five in stress responses, seven in motility and chemotaxis, four in pathogenesis, two in the bacterial cytoskeleton, and five have no known function. In addition, 48 proteins were significantly upregulated in the OrfSwap*^ihfA-ihfB^* strain. Of these, 15 proteins were involved in metabolism, nine were associated with the outer membrane, two were involved in genetic information processing, four in translation, one in motility and chemotaxis, one in bacterial cytoskeleton, one in pathogenesis, five in stress resistance and ten proteins with no known function.

### Differential mRNA stability correlates with the relative IHF subunit levels in the wild type and the OrfSwap*^ihfA-ihfB^* strain

Since swapping the protein-coding regions of *ihfA* and *ihfB* had created hybrid mRNA in the OrfSwap*^ihfA-ihfB^* strain (Fig. 1) we investigated the possibility that differences in wild type and hybrid *ihfA* and *ihfB* mRNA stabilities might account for the differences in protein production. Rifampicin treatment [87] was used to arrest transcription and then total RNA was harvested at fixed time intervals. RT-qPCR, using primers specific for *ihfA* and *ihfB* mRNA (Table S1), was used to amplify the transcripts of these genes [88]. Messenger RNA half-life (T½) was measured as described by Chen *et al.* [89]. In SL1344 at 3.5 h, the T½ of *ihfA* mRNA was 2.78 min while that of *ihfB* mRNA was 3.54 min (Fig. 4a). Interestingly, at 7 h, T½ *ihfB* mRNA was 4.93 min, indicating significantly greater stability than *ihfA* mRNA, where T½ was 2.19 min (Fig. 4b). This difference was consistent with the almost two-fold excess of IhfB protein over IhfA at 7 h (Fig. 3). However, at this time point in the OrfSwap*^ihfA-ihfB^* strain the relative stabilities of *ihfA* and *ihfB* mRNA were found to be similar, with *ihfA* mRNA becoming slightly more stable than *ihfB* mRNA. At 3.5 h, the T½ of *ihfA* mRNA was 2.62 min, while the T½ of *ihfB* mRNA was 2.32 min (Fig. 4c). At 7 h, the T½ of *ihfA* mRNA was 3.38 min, while the T½ of *ihfB* mRNA was 2.64 min (Fig. 4d). These OrfSwap*^ihfA-ihfB^* strain data showed that the hybrid mRNAs generated when the protein-coding segments of the *ihfA* and *ihfB* genes were exchanged, had half-lives that differed from those of the native genes in SL1344. Therefore, relative mRNA stability correlated with the levels of IHF proteins in SL1344 and in the OrfSwap*^ihfA-ihfB^* strain. Overall, our data reveal that reciprocally exchanging *ihfA* and *ihfB* resulted in IhfA being produced in excess of IhfB, a reversal of the SL1344 production pattern. What were the physiological consequences of this reversal?

**Fig. 4.**
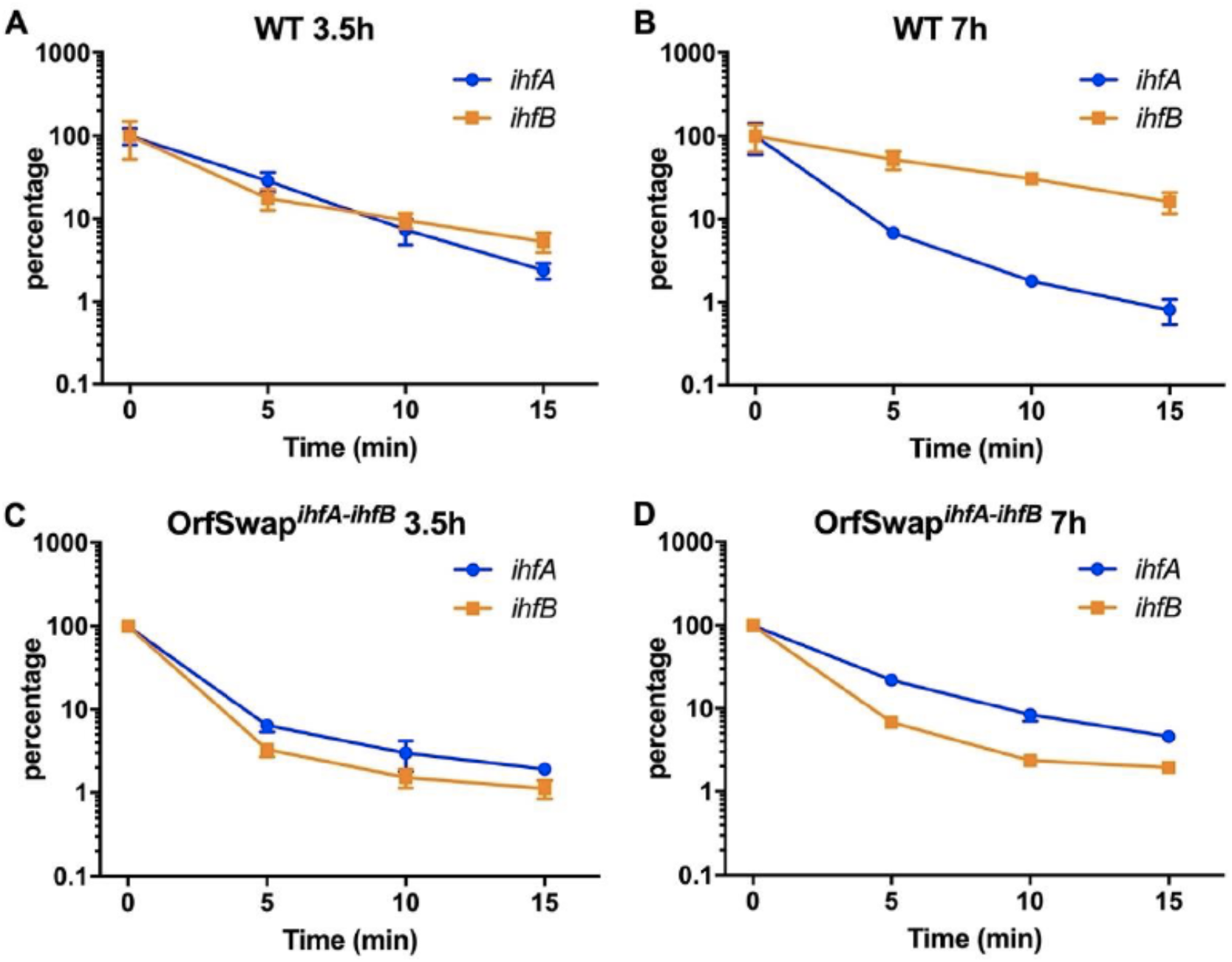
Differential mRNA stability accounts for the relative IHF subunit levels in the wild type (SL1344) and in the OrfSwap*^ihfA-ihfB^* strain. The abundance of mRNA was measured by RT-qPCR at 5, 10 and 15 min after the addition of 500 µg/ml rifampicin (0 min). A) Stabilities of *ihfA* and *ihfB* mRNA at 3.5 h in the WT, B) Stabilities of *ihfA* and *ihfB* mRNA at 7 h in the WT, C) Stabilities of *ihfA* and *ihfB* mRNA at 3.5 h in the OrfSwap*^ihfA-ihfB^* strain, D) Stabilities of *ihfA* and *ihfB* mRNA at 7 h in the OrfSwap*^ihfA-ihfB^* strain. Data are presented as percentages of the Time 0 value (100%). Mean and standard deviation of three biological replicates is shown.

### The growth, morphological, and competitive fitness characteristics of strains with repositioned and rewired *ihf* genes

To assess the impact of the reciprocal exchanges of the coding regions of the IHF-encoding genes on the growth characteristics of *Salmonella* Typhimurium, the growth profiles of the OrfSwap*^ihfA-ihfB^* strain was monitored in liquid medium by optical density at 600 nm (OD_600_) and compared to SL1344. The growth pattern of the OrfSwap*^ihfA-ihfB^* strain was identical to that of the SL1344 (Fig. S4a). Growth was also measured by calculating the number of colony-forming units in a liquid culture by spreading aliquots onto agar plates at fixed time intervals. Once again, no differences were seen between OrfSwap*^ihfA-ihfB^* strain and the WT (Fig. S4b). Similarly, SL1344 and the OrfSwap*^ihfA-ihfB^* strain had identical cell morphologies (Fig. S5) and were equal in competitive fitness (Fig. S6).

### Motility in SL1344 and OrfSwap*^ihfA-ihfB^* strain

As a master regulator, IHF influences the expression of multiple genes, including that of the entire motility regulon: IHF is a positive regulator of flagellar gene expression at all stages of flagellum production [44]. The motilities of the OrfSwap*^ihfA-ihfB^* strain and SL1344 were compared by measuring the diameters of motility zones after 5 h of incubation at 37°C on 0.3% LB agar. Motility differences were recorded as the fold change relative to the SL1344. In keeping with the known positive role of IHF in motility, the OrfSwap*^ihfA-ihfB^* displayed a statistically significant decrease in motility compared to SL1344 (Fig. 5). The motility phenotype of the OrfSwap*^ihfA-ihfB^* strain resembled that previously described for an SL1344 *ihfA ihfB* double mutant [44], indicating that the motility system is sensitive to even modest adjustments to IHF subunit production patterns.

**Fig. 5.**
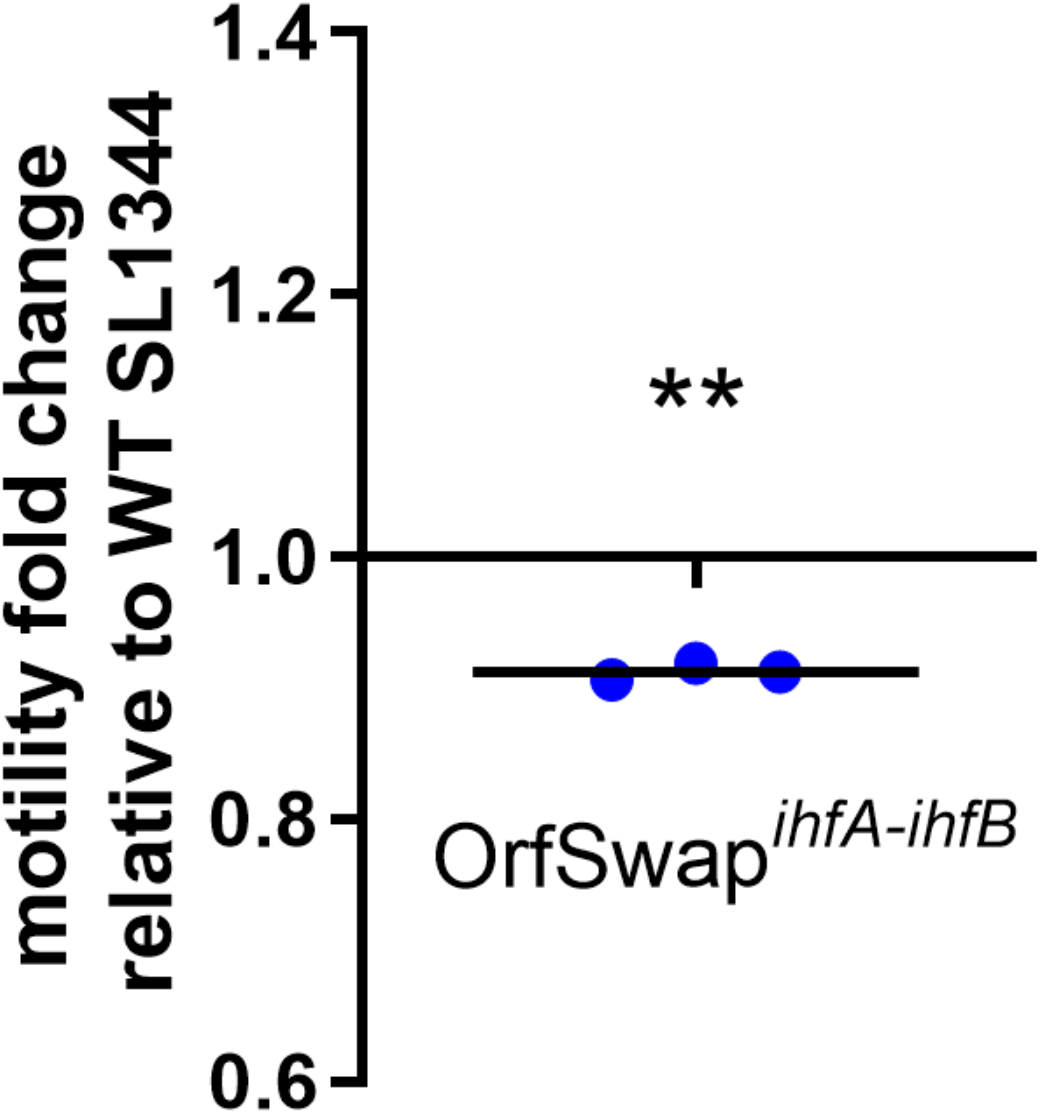
Motility in the WT (SL1344) and OrfSwap*^ihfA-ihfB^* strain. Diameters of swimming motility zones were measured after 5 h incubation at 37°C on soft 0.3% LB agar. The OrfSwap*^ihfA-ihfB^* strain was less motile than the WT as indicated by values below 1. Significance was found by one sample T-test, where P<0.05.

### SPI-1 and SPI-2 gene expression in SL1344 and OrfSwap*^ihfA-ihfB^* strains

*Salmonella* pathogenicity islands (SPIs) are horizontally acquired genetic regions on the *Salmonella* chromosome [90–93]. SPI-1 and SPI-2 each encode a type III secretion system (T3SS) and type III secreted effector proteins. These mediate *Salmonella’s* virulence via invasion of epithelial cells and survival within macrophages respectively [94, 95]. Complex, interconnected regulatory networks control the expression of the SPI genes [96–99]. IHF affects expression of both SPI-1 and SPI-2 positively and appears to play a coordinating role [44].

Fusions of the *gfp* reporter gene to the promoters of *prgH* and of *ssaG*, genes that encode needle components of SPI-1 and SPI-2, respectively, were used as proxies of SPI-1 and SPI-2 transcription [83, 100], allowing these to be compared in SL1344 and OrfSwap*^ihfA-ihfB^* strain. Details of the strain constructions and genotypes (Table 1) are given in METHODS.

Green-fluorescent-protein (GFP)-mediated fluorescence at 528 nm was measured every 20 min over 24 h and adjusted to the OD_600_ of the culture. The expression of the reporter fusions derivatives of SL1344 was in agreement with published data [101]: SPI-1 expression peaked during mid-exponential growth phase (Fig. 6a), while SPI-2 gene expression peaked in early stationary phase and plateaued thereafter (Fig. 6b). In the OrfSwap*^ihfA-ihfB^* strain, expression of both pathogenicity island genes was almost identical to that seen in SL1344 wild type control except for a small, but statistically significant, decrease during late-stationary phase in SPI-1 gene expression (Fig. 6a) and during mid-exponential phase in SPI-2 genes (Fig. 6b).

**Fig. 6.**
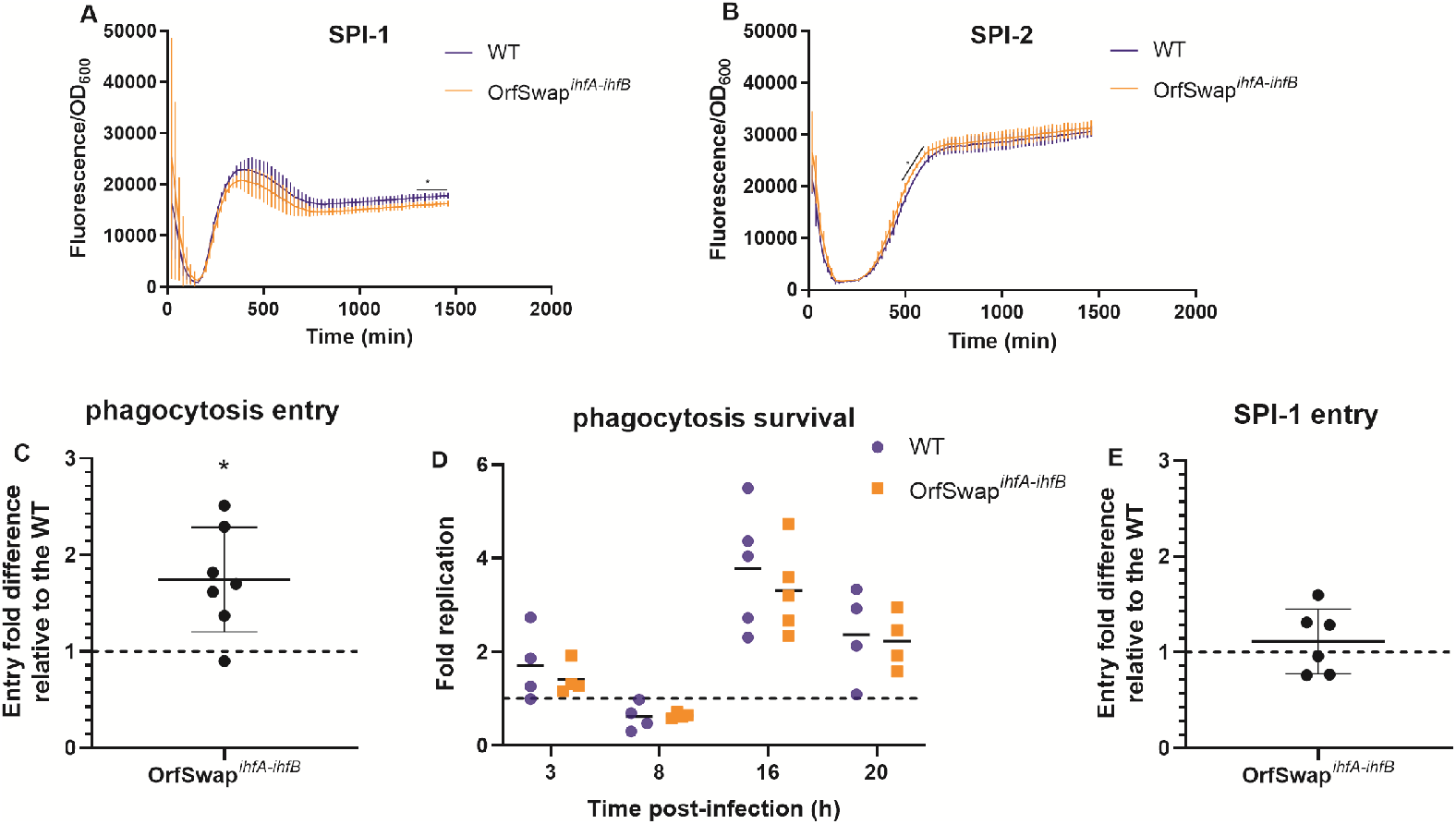
SPI-1 and SPI-2 gene expression in the WT (SL1344) and OrfSwap*^ihfA-ihfB^* strains and infection and survival in murine macrophages. *Salmonella* pathogenicity island (SPI) expression was measured with aid of *gfp* reporter fusions. Fluorescence at 528 nm, calculated relative to OD_600_, was recorded at 20-min intervals over 24 h. A) SPI-1 expression in the OrfSwap*^ihfA-ihfB^* strain was slightly lower than in the WT during the late stationary growth phase. B) SPI-2 expression in the OrfSwap*^ihfA-ihfB^* strain was slightly lower than in the WT during the mid-exponential phase. All plots are the results of at least three biological replicates and error bars represent standard deviation. Asterisks represent statistical significance for the time points highlighted with black curves. Significance was found by Student’s unpaired T-test, where P<0.05. C) RAW264.7 cells were infected with SPI-2 induced bacteria, grown to stationary phase and complement-opsonised to promote phagocytosis. Entry was measured by enumerating CFUs 1 h post-infection. In each replicate, CFUs were calculated relative to the infection mix and WT.. Mean and standard deviation are shown. Significance was found by one sample T-test, where P<0.05. D) Survival and replication were measured by enumerating CFUs at 3 h, 8 h, 16 h and 20 h post-infection. Fold replication represents the CFU recovered at the particular time point divided by the CFU at 1 h. Mean and individual replicates are shown. No significant difference was found by unpaired Student’s T-test. E) RAW264.7 cells were infected with SPI-1 induced bacteria, grown to the mid-exponential phase of growth to promote SPI-1 mediated active entry. Entry was measured by enumerating CFUs 1 h post-infection. In each replicate, CFUs were calculated relative to the infection mix and WT. Mean and standard deviation are shown. No significant differences were found by one sample T-test, where P<0.05.

### Infection and survival of SL1344 and OrfSwap*^ihfA-ihfB^* strains in murine macrophages

RAW264.7 murine macrophages are a well-studied cell line, frequently used for studying *Salmonella* infection. Gentamycin protection assays were carried out to assess the performance of SL1344 and OrfSwap*^ihfA-ihfB^* strains at different stages of the infection process: efficiency of entry to the macrophage, intracellular survival, and replication over a 20-h period post-infection. Survival and replication of *Salmonella* in macrophages is dependent upon the mechanism of entry [102]. The two main modes of entry utilised by *Salmonella* are: (1) SPI-1 mediated active invasion and (2) passive, host-mediated phagocytosis. Both modes are important as they occur in *Salmonella’*s natural host environment. Macrophages may encounter highly invasive *Salmonella*, in which SPI-1 genes are induced, as the bacteria emerge from the intestinal epithelial cell barrier [103, 104]. But *Salmonella* can also be taken up by macrophage via phagocytosis. This occurs when the microbe escapes its original macrophage replication niche while expressing the SPI-2 type III secretion system needle on its surface or when *Salmonella* is disseminated to the circulatory system during systemic infection [105].

To achieve active entry to macrophage via SPI-1, bacteria were grown to midexponential growth phase in LB broth to maximize SPI-1 expression [106]. To ensure passive entry via phagocytosis, bacteria were grown to late stationary growth phase and complement-opsonised [107]. Invasive SPI-1 induced bacteria were used at a multiplicity of infection (MOI) of 5, while opsonised SPI-2 induced bacteria were used at a MOI of 20 to achieve similar entry numbers. The 1-h post-infection time point was chosen as a reference point for invasion.

To promote phagocytosis by RAW264.7 macrophages, these cells were infected with SPI-2-induced and opsonised *Salmonella*. Both infection entry efficiency and long-term intracellular survival were tested. SL1344 was used as a reference strain in all experiments. Counts of bacterial cells of the OrfSwap*^ihfA-ihfB^* strain internalized by macrophages 1 h post infection were calculated relative to the infection mix and also relative SL1344. The data obtained showed that the OrfSwap*^ihfA-ihfB^* strain was phagocytosed more readily than SL1344 (Fig. 6c). This enhanced phagocytosis suggested that the cell membrane composition of the OrfSwap*^ihfA-ihfB^* is different from that of SL1344. This may have allowed more efficient complement deposition on the microbial cell surface and hence better phagocytosis.

*Salmonella* survival and replication inside a macrophage were tested using the same infection model. CFUs within macrophages were counted at further time points and calculated relative to CFUs at 1 h. For both strain analysed, after slight initial growth at 3 h post infection, a killing event was observed at 8 h post-infection (p.i). At 16 h p.i., *Salmonella* appeared to adapt and replicate significantly, before undergoing a further decline by 20 h p.i., possibly caused by macrophage cell death [108]. When the OrfSwap*^ihfA-ihfB^* strain was compared to the WT, no significant difference were found, at any of the time points tested (Fig. 6d).

The lack of any difference in survival rates between the strains may seem to be inconsistent with the observation that the expression levels of the SPI genes in the repositioned strains were slightly lower than in SL1344 (Fig. 6a, 6b). However, SPI expression depends on the mechanism of *Salmonella* uptake by the macrophage. Following phagocytosis of *Salmonella*, both SPI-1 and SPI-2 regulon genes are expressed to a much smaller magnitude than when *Salmonella* invades macrophages using the SPI-1-encoded T3SS [102]. Therefore, an infection model with SPI-1-induced bacteria was tested. No differences in the active SPI-1 mediated entry was found between SL1344 and the OrfSwap*^ihfA-ihfB^* strain (Fig. 6e). This suggests that the magnitude of the SPI expression changes observed in Fig. 6a and Fig. 6b was too low to cause detectable alterations in the ability of the bacterium to infect mammalian macrophages.

## DISCUSSION

A gene’s location on the bacterial chromosome is known to be an important factor in establishing the appropriate spatiotemporal expression pattern of that gene and its influence on physiology [60-62, 64, 65, 69, 109]. The most influential parameter is the distance of the gene from the origin of chromosome replication, *oriC* [60, 67, 68, 69]. In rapidly growing bacteria with multiple rounds of replication underway, genes closest to *oriC* will be present in more copies than those closer to the terminus. The resulting differences in gene dosage contribute to variations in the output of the gene product at different stages of the growth cycle. As stationary phase approaches and replication ceases, *oriC*-proximal genes lose their copy number advantage over those closer to the terminus. Even so, moving *fis*, the gene encoding the FIS NAP, from its normal *oriC*-proximal position to a location near the terminus had a relatively mild impact of the physiology of the bacterium in *S*. Typhimurium [60] and in *E. coli* [65] compared with the strong phenotypic effects seen when *fis* has both the chromosomal location and the regulatory signals of the terminus-proximal *dps* gene [60].

In this study, we have explored the significance of the chromosomal location of each gene that encodes one of the two subunits of IHF in *S.* Typhimurium, and the significance of each gene’s individual regulatory pattern, for the physiology of the bacterium. To do this, we exchanged, reciprocally, the protein-coding regions of the *ihfA* and *ihfB* genes. The exchange placed each gene under the control of the expression signals of the other gene (Fig. 1). Whole genome sequencing confirmed that the only changes made to the chromosome in the OrfSwap*^ihfA-ihfB^* strain were those introduced CDS exchange. The *ihf* genes are in a quadrant of the chromosome that is close to the replication terminus, and IHF is normally expressed maximally at the transition from exponential growth to stationary phase [15, 17]. In the OrfSwap*^ihfA-ihfB^* strain, each sequence has moved to a location that was approximately 350 kbp away from its native position within the Right replichore of the chromosome.

IHF contributes to many vital processes in the bacterial cell, yet it is not an essential protein. Bacteria with deletions of both the *ihfA* and *ihfB* genes are viable, although they exhibit dysregulation of many systems [44, 46]. This suggests that IHF may be redundant, or partially redundant in some molecular processes to which it contributes, or that the contribution made by IHF is dispensable, at least under some conditions. The related nucleoid-associated protein, HU, also a heterodimeric DNA binding protein, and with a structure that is quite similar to that of IHF, may perform some IHF roles when IHF itself is unavailable. Although it is a non-specific DNA binding protein, HU can substitute functionally for IHF in bacteriophage lambda integrase-mediated DNA excision, but not integration [110]. This is because HU provides the missing architectural function in the lambda intasome in the absence of IHF [111]. The regulons of IHF and HU in *S*. Typhimurium show overlapping, but non-identical, memberships [44, 112] suggesting partial, rather than complete functional redundancy between the two NAPs. The suggestion that redundancy between IHF and HU is only partial is supported by work showing that *E. coli* mutants devoid of HU have phenotypes that cannot be complemented by overexpression of IHF [113]. Our proteomic data revealed that, like IhfB, the B subunit of IHF’s paralogue HU, was also present at a reduced level in the OrfSwap*^ihfA-ihfB^* strain. This finding was consistent with transcriptomic data showing that the gene encoding HU’s B subunit, *hupB*, is down regulated in the absence of IhfB at early stationary phase [44]. Perhaps this adjustment to HU subunit production ameliorated the impact of inverting the relative abundances of IhfA and IhfB in the rewired strain.

The mutual dependency of IhfA and IhfB on one another for the assembly of stable IHF protein indicates that it is important to maintain the relative levels of these proteins within certain limits. IhfA is especially unstable in the absence of IhfB and can form IhfA homodimers on DNA only when present at very high concentrations [70]. Our finding that IhfB is produced in excess over IhfA may represent an attempt by the cell to address this issue. This situation is in marked contrast to that of HU, also an alpha-beta heterodimeric protein, but one that also exists both as stable alpha-homodimers and beta-homodimers, each of which has distinct cellular functions [114]. Transcriptomic data from *hupA, hupB* and *hupA hupB* knockout mutants indicate that each HU subunit has a substantial regulon, but that these overlap incompletely with each other [112].

A need to produce IhfB and IhfA at different levels may also explain why the *ihfA* and *ihfB* genes are not organised in a single operon. While it is known that operon-encoded gene products can be tuned at the level of translation initiation to achieve a required stoichiometry [115], it may also be advantageous to achieve the required balance by differential mRNA stability, and/or by differential transcription, by expressing the messages from separate transcription units under distinct control regimes. The case of IHF production seems even more sophisticated when we take into account the observation that the *ihfA* and *ihfB* genes are embedded in operons that express components of the cell’s translational machinery. It has been proposed previously that this might account for the presence of *ihfA* (*himA*) in an operon with *pheST* [116]. In this way, a rise or fall in demand for translation capacity (a proxy for metabolic flux) can feed back onto a regulator (IHF) that modulates many key cellular processes. A connection between IHF production and translation capacity is consistent with the observation that transcription of both *ihfA* and *ihfB* responds to (p)ppGpp, a signal that indicates an accumulation of uncharged tRNA molecules, and hence over-capacity in the translation apparatus [15]. The *ihfA* promoter lies within the *pheT* gene and is autoregulated by IHF, as is *ihfB* transcription [15]. The *pheST-ihfA* operon is controlled by transcription attenuation in response to phenylalanine and by the SOS response, indicating sensitivity to changes in translational capacity and to processes involved in DNA damage repair. Certainly, genes encoding components of the DNA metabolism and the translation machinery are well represented in the *Salmonella* IHF transcriptome [44] and their products in the proteome (Table S2). Most translation-associated proteins were present in reduced amounts in the OrfSwap*^ihfA-ihfB^* strain, mimicking the impact of loss of IhfB on the transcriptome at early stationary phase [44]. The majority of proteins with an altered production level in the OrfSwap*^ihfA-ihfB^* strain were present in lower amounts than in the wild type SL1344 strain. The reduction in translation-associated proteins may reflect an attempt to keep overall protein production within acceptable limits, minimising the number and severity of impacts on phenotype.

Despite the reversal of the relative levels of IhfA and IhfB in the OrfSwap*^ihfA-ihfB^* strain, our data show that giving each *ihf* gene the transcription control region and the chromosome location of the other had, at best, subtle effects on *Salmonella* physiology. For example, the reduced expression of the rod-shape-determining cytoskeletal protein MreB in stationary phase, which correlated with the previously described reduction in *mreB* transcription in an *ihfB* knock-out mutant at the same phase of growth [44], did not produce an obvious change to cell morphology (Fig. S5). On the other hand, the wild type cells had already lost their rod shape at this stage of growth, in agreement with previous work on growth phase and morphology and the down-regulation of *mreB* expression with cessation of growth [117]. Where phenotypic changes did occur, they involved processes where T3SS plays a prominent part: motility, host cell invasion and macrophage survival. All three of these are part of the H-NS regulon [118–121] while SPI-1 and SPI-2 consist of genetic elements that have been acquired by horizontal gene transfer [120, 121]. Their encoded T3SS exhibit a marked hypersensitivity to changes to the expression of NAPs and the control of DNA topology [44, 60, 66, 96, 112, 122, 123], indicating that they belong to a genomic category that has an acute dependency on wildtype patterns of global gene control. This may be because they specify gene products that are physiologically expensive to produce.

## AUTHOR STATEMENTS

### Authors and contributors

GP (performed all of the experiments, gathered, analysed, interpreted and curated the data, and co-wrote the paper), AM (curated the whole genome sequencing [WGS] data, analysed and interpreted the experimental data and co-wrote the paper) MCB (contributed to the acquisition of the proteomic data and their interpretation), NRT (generated the WGS data, contributed the funding for WGS, and co-wrote the paper), CJD (acquired funding for the work, devised and managed the project, provided team supervision, analysed and interpreted the experimental results and co-wrote the paper).

