## Supplemental Figs 1-6 and Table S1 for "The consequences of reciprocally exchanging the genomic sites of Integration Host Factor (IHF) subunit production for subunit stoichiometry and bacterial physiology in *Salmonella enterica* serovar Typhimurium"

### SUPPLEMENTARY FILES

**Table S1** Oligonucleotides used in this study

| Name | 5'-3' sequence |
|---|---|
| **Confirmation Primers**: | |
| SL_ihfA_check_Pfwd | CAACCGTACACTCGAAGAAGAG |
| SL_ihfA_check_Prev | GAACGTTTCGTCGCTGTTG |
| SL_ihfB_check_Pfrw | GCAATGGCTGAAGCATTCAAAG |
| SL_ihfB_check_Prev | GACCGTCGTTATCTTCATAGACAC |
| **qPCR primers** | |
| SL_ihfA_qPCR_Pf | GATAAGCTTGGGCTTAGCA |
| SL_ihfA_qPCR_Pr | GAGAGTTTCACCTGCTCAC |
| SL_2ihfB_qPCR_Pf | CAATCTCACATTCCCGCTAAG |
| SL_2ihfB_qPCR_Prev | GCGGATTTCAATACGCTCG |
| SL_btuC_qPCR_Pf | GGCTACACTTCTGAGCTTATG |
| SL_btuC_qPCR_Pr | GGTTCAGCAAGTGGGTTT |
| SL_ycaI_qPCR_Pf | GTCAGCGTATGCGTAGTATG |
| SL_ycaI_qPCR_Pr | CAAAGCAAACATGCCAGAC |
| RT_hemX_F | CGCCTGACGGTATGTTTCTT |
| RT_hemX_R | CCCAACCAGGACGTCTATTTAC |
| SL_ihfB_qPCR_Pf | CATATGGCCTCGACTCTTG |
| SL_ihfB_qPCR_Pr | TCCTTCCAGTTCCACTTTATC |
| ***ihfA* ORF insertion into *infB* locus** | |
| infA.cmR.Pfwd | GAGCCGGGTCGAAAACGCTTCGCCCAAAGAAGAGTAATCAGTGTAGGCTGGAGCTGCTTC |
| infA.cmR.Prev | GCGTATCTGCCGCAATACACCCTGATGGATGTTATGCCTGCATATGAATATCCTCCTTAG |
| infB.int.Pfwd | ACGGCTGCAGCCAATTTGCCTTTAAGGAACCGGAGGAATCATGGCGCTTACAAAAGCTG |
| infB.int.Prev | CGGTGCTTTTTTCGGGTTCAAGTTTTGCGTTAAAACCCTGCATATGAATATCCTCCTTAG |
| ***ihfB* ORF insertion into *ihfA* locus** | |
| ihfB.kanR.Pfwd | AGAACTGCGCGATCGCGCCAATATTTACGGTTAAGTTTTAGTGTAGGCTGGAGCTGCTTC |
| ihfB.kanR.Prev | CAAAACTTGAACCCGAAAAAAGCACCGTCAGGGTGCTTTTCATATGAATATCCTCCTTAG |
| ihfA.int.Pfwd | AAAAGAGCGATTCCAGGCATCATTGAGGGATTGAACCTATGACCAAGTCAGAATTGATTG |
| ihfA.int.Prev | GCAATACACCCTGATGGATGTTATGCCTGGATCTGACATA |

| | |
|---|---|
| | TGAATATCCTCCTTAGTTCC |
| ***ihfB*::kan insertion downstream of *ihfA*** | |
| ihfA.int.ihfB::kan_Pf | GAGCCGGGTCGAAAACGCTTCGCCCAAAGAAGAGTAATCATTGCCTTTAAGGAACCGGAG |
| ihfA.int.ihfB::kan_Prev | CGTATCTGCCGCAATACACCCTGATGGATGTTATGCCTGGCATATGAATATCCTCCTTAG |
| ***ihfA*::Cm insertion downstream of *ihfB*** | |
| ihfB.int.ihfA::Cm_Pf | TAAAGAACTGCGCGATCGCGCCAATATTTACGGTTAAGTTCAGGCATCATTGAGGGATTG |
| ihfB.int.ihfA::Cm_Prev | AGCACCCTGACGGTGCTTTTTTCGGGTTCAAGTTTTGCGTCCTGCATATGAATATCCTCC |
| **Deletion mutations – kan[r] insertions** | |
| Kan_ihfA_Pf | AAAGAGCGATTCCAGGCATCATTGAGGGATTGAACCTATGGTGTAGGCTGGAGCTGCTTC |
| Kan_ihfA_Prev | ATGTTATGCCTGGATCTGATTACTCTTCTTTGGGCGAAGCCATATGAATATCCTCCTTAG |
| Kan_ihfB_Pf | GCTGCAGCCAATTTGCCTTTAAGGAACCGGAGGAATCATGGTGTAGGCTGGAGCTGCTTC |
| Kan_ihfB_Prev | TCAAGTTTTGCGTTAAAACTTAACCGTAAATATTGGCGCGCATATGAATATCCTCCTTAG |

For strain construction primers – the black portion is an annealing end; the red portion is an overhanging end. All primers were designed in this study.

**Table S2**. A listing of the proteins that displayed altered levels of production when the proteomes of the wild type and the OrfSwap$^{ihfA\text{-}ihfB}$ strains were compared. The table has three parts: Up-regulated proteins, Down-regulated proteins and All proteins. A detailed legend is provided at the top of the table, explaining the colour code used to distinguish the different functional categories, and the rationale used for the inclusion of proteins and for their assignment to the Up-regulated or Down-regulated lists.

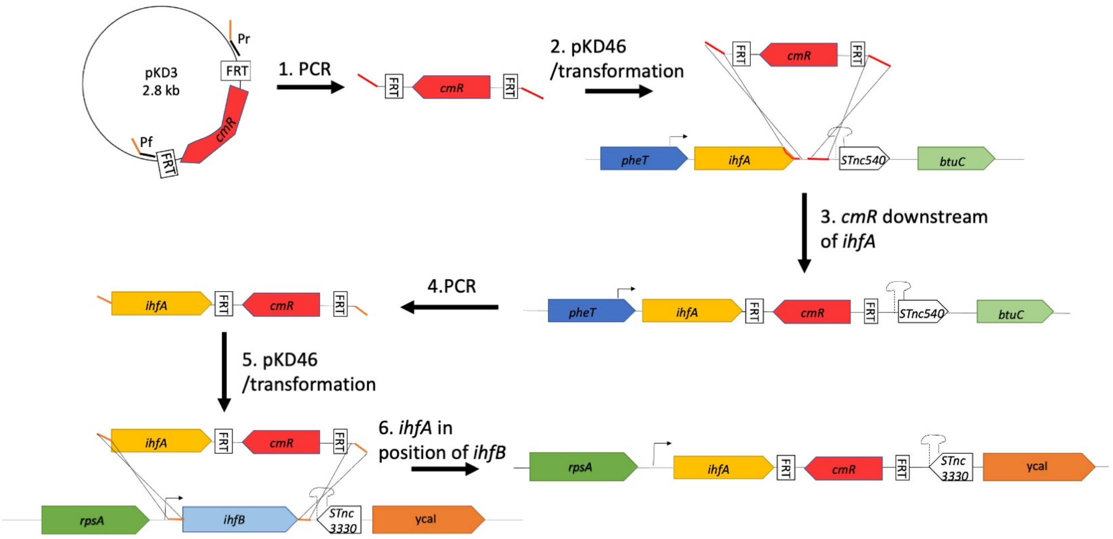


**Fig. S1. Strain construction strategy.** The strategy is illustrated using the example of placing the open reading frame of *ihfA* in the position normally occupied by the open reading frame of *ihfB*. 1. A chloramphenicol resistance cassette was amplified by PCR from the carrier plasmid pKD3 with a pair of primers that had overhangs (depicted in red) homologous to the region downstream of *ihfA*. 2. A linear PCR product was purified and transformed into wild type SL1344 harbouring pKD46. 3. In a number of cases, the linear product was inserted into the target region by Lambda-Red-mediated recombination, yielding an intermediate strain with a Cm$^{R}$-tagged *ihfA*. 4. In another round of PCR, the *ihfA*-Cm$^{R}$ construct was amplified with a pair of primers that had overhangs homologous to the regions just upstream and downstream of *ihfB*. 5. A linear PCR product was purified and transformed into the wild-type SL1344 strain harbouring pKD46. 6. In a number of cases, the linear product was inserted into the target region by Lambda-Red-mediated recombination, yielding a strain with *ihfA*-Cm$^{R}$ in place of *ihfB*. Where required, the Cm$^{R}$ resistance cassette was then removed by pCP20-mediated FLP site-specific recombination.

**A**

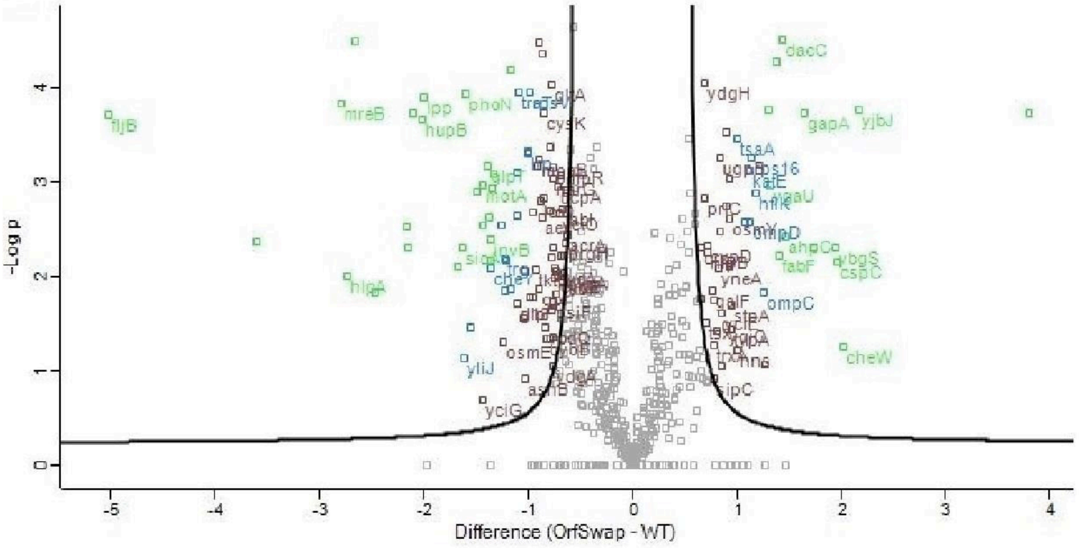


**B**

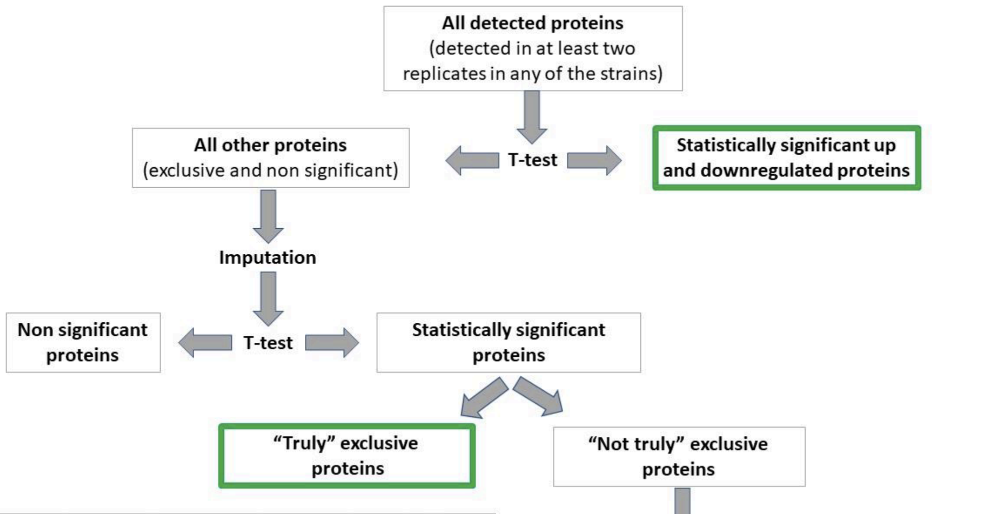


| | WT_1 | WT_2 | WT_3 | OrfSwap_1 | OrfSwap_2 | OrfSwap_3 |
|---|---|---|---|---|---|---|
| "Truly" exclusive | + | + | + | - | - | - |
| "Truly" exclusive | - | - | - | + | + | - |
| "Not truly" exclusive | + | + | + | - | + | - |
| "Not truly" exclusive | + | - | - | + | - | + |

**C**

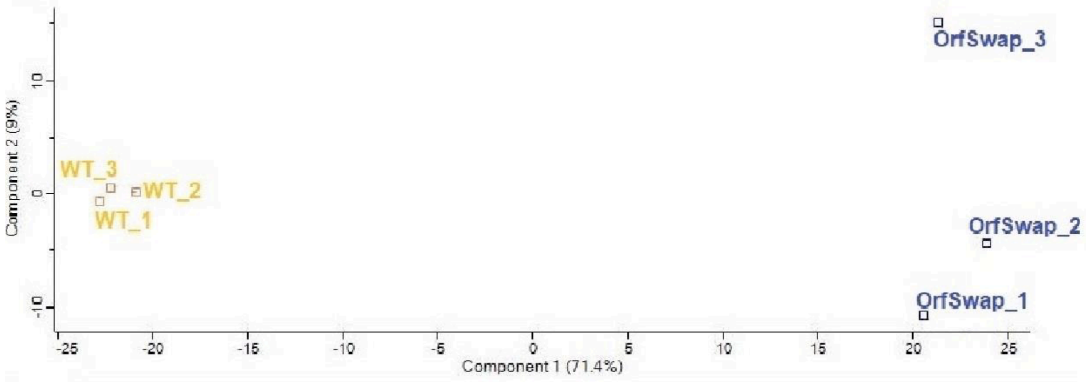

**Fig. S2.** Mass spectrometry data of the WT (SL1344) and the OrfSwap$^{ihfA\text{-}ihfB}$. Data were obtained from three biological replicates of each strain grown in LB for 7 h. A) Volcano plot of the differentially expressed genes before imputation. The differences in log2-transformed LFQ intensity were plotted against negative log10-transformed p-values of the two-sided T-test with the S0 = 1. Green represents FDR = 0.001, blue and green represent FDR = 0.01, brown, blue and green represent FDR = 0.05. The analysis was performed in Perseus 1.6.14.0. B) The flow diagram of the strategy used to determine differentially expressed proteins (read from the top). The proteins in green frames were deemed differentially expressed. The insert table defines “truly” exclusive and “not truly” exclusive proteins. “+” represents presence of a protein in a sample, “-“ indicates that a protein was not detected in a sample by MS. C) Principal component analysis of the three WT and OrfSwap$^{ihfA\text{-}ihfB}$ biological replicates shows clustering of the MS samples. The analysis was performed in Perseus 1.6.14.0.

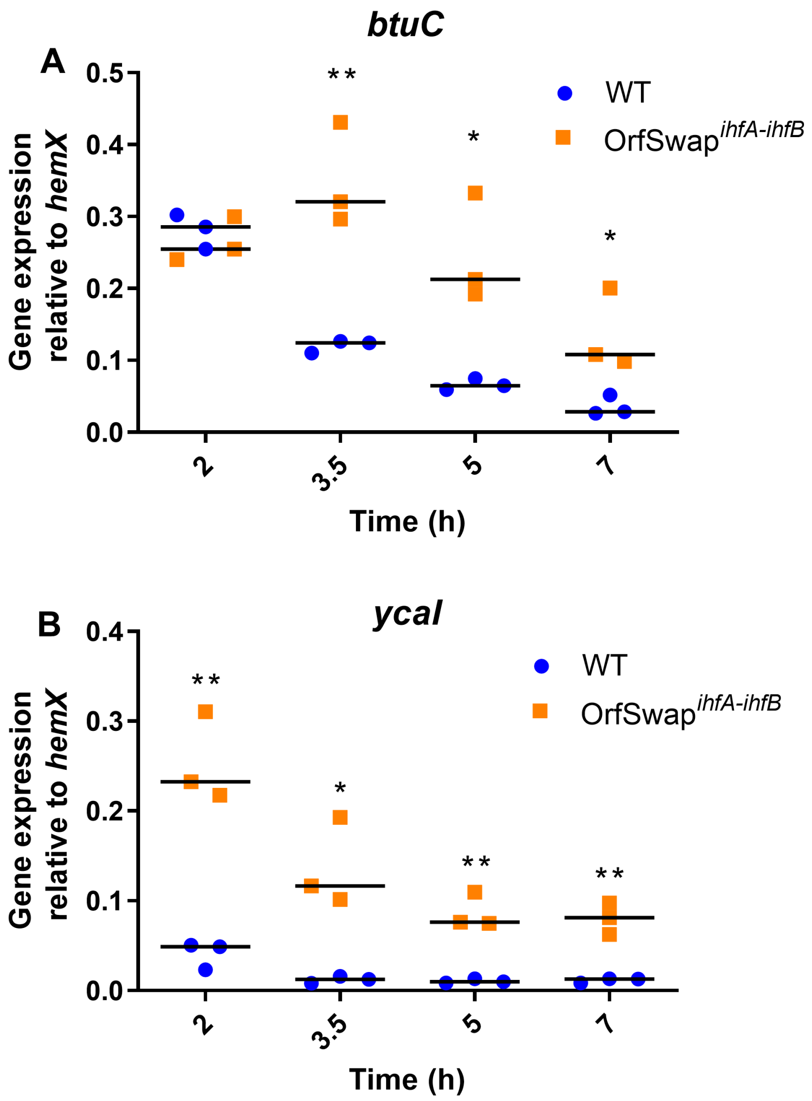


**Fig. S3.** Expression of genes downstream of *ihfA* and *ihfB*. A) Gene expression of *btuC* (a gene downstream of *ihfA*) was measured using RT-qPCR in the WT SL1344 and in the OrfSwap$^{ihfA\text{-}ihfB}$ strains. B) Gene expression of *ycaI* (a gene downstream of *ihfB*) was measured using RT-qPCR in the WT SL1344 and in the OrfSwap$^{ihfA\text{-}ihfB}$ strains. The time points represent lag (2 h), mid-exponential (3.5 h), transition from exponential to stationary (5 h) and early stationary (7 h) growth phases, respectively. All plots are results three biological replicates. Significance was found by unpaired Student's T-test, where $P<0.05$.

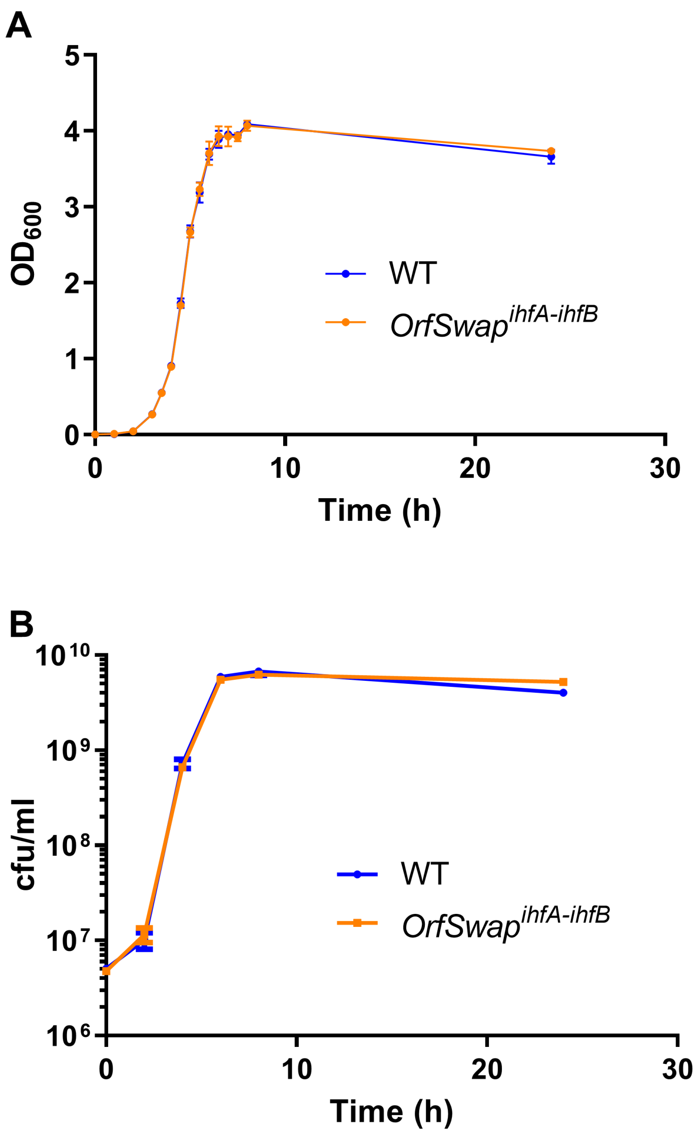


**Fig. S4.** The growth kinetics of strains with repositioned and rewired *ihf* genes Comparisons of the growth patterns of the wild type (SL1344) and OrfSwap$^{ihfA\text{-}ihfB}$, strains, measured by: A. Optical density measurement at 600 nm, with readings taken every hour until 3 h, then every 30 min until 8 h and finally at 24 h. B. Cell viability measurements made by spreading dilutions of bacterial cultures onto agar plates, incubating at 37°C and counting colonies.

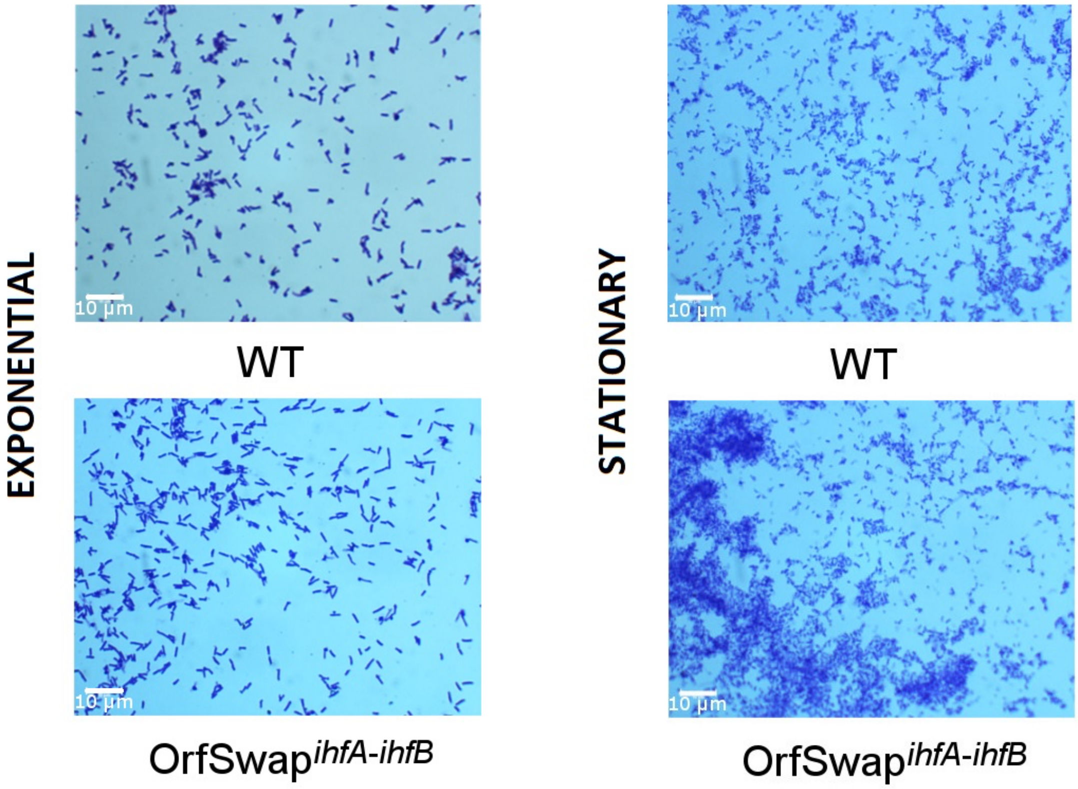


**Fig. S5. Morphologies of strains with repositioned and rewired *ihf* genes.** Bacteria were harvested, washed with PBS, heat-fixed, stained with crystal violet and viewed under 1000x magnification with oil immersion lens. Strain morphologies at mid-exponential and stationary growth phases. All images are representative of three biological replicates. The 10 µm bar is given for reference.

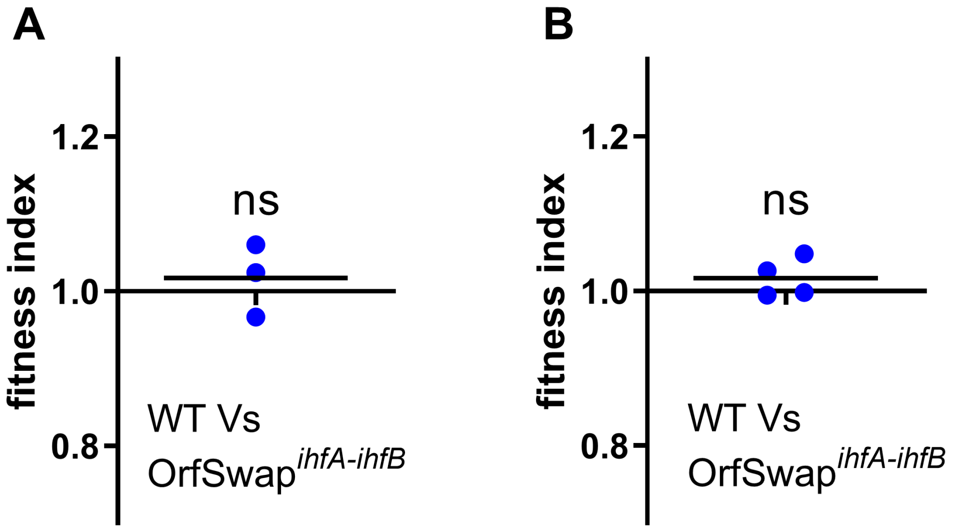


**Fig. S6.** The competitive fitness characteristics of strains with repositioned and rewired *ihf* genes. Fitness of the OrfSwap$^{ihfA\text{-}ihfB}$ strain relative to the WT SL1344 in LB broth supplemented with a) 0.171 mM NaCl or b) 0.3 mM NaCl and grown for 24 h at 200 rpm at 37°C. Fitness index = 1 means that the competed strains were equally fit, f.i. < 1 indicates that the competitor strain (OrfSwap$^{ihfA\text{-}ihfB}$) was less fit than the WT, f.i. > 1 indicates that the competitor was more fit than the WT. One sample T-test was used to determine significance, where $p<0.05$.
