## Supplemental Table S2 for "The consequences of reciprocally exchanging the genomic sites of Integration Host Factor (IHF) subunit production for subunit stoichiometry and bacterial physiology in *Salmonella enterica* serovar Typhimurium"

### LEGEND:

Colour code:

- Metabolism
- Translation
- Genetic information processing
- Division and cytoskeleton
- Stress response and environmental monitoring
- Motility and chemotaxis
- Pathogenicity
- Outer membrane and transport
- Function unknown

*Salmonella* SL1344 WT Vs OrfSwap*ihfA-ihfB* proteome comparison data.
The data was obtained using LC-MS/MS on QExactive and represents three biological replicates. The RAW-files were analysed with MaxQuant and its output was analysed statistically in Perseus.

A protein was considered as present in a strain if it was detected in at least two biological replicates in that strain.

A protein was considered as exclusive for a strain if it was detected in at least two biological replicates in that strain and in none of the samples in the other strain.

The sections named "Downregulated in OrfSwap" and "Upregulated OrfSwap" are subsets of the main dataset shown within the section named "All proteins".

In the "All proteins" the protein entries are arranged in the order of increasing t-test q-value within each cluster. In the other sheets the protein entries are arranged in categories

The differentially expressed proteins (Upregulated and Downregulated) were determined from the unimputed data

The imputation was done for all other proteins, including unique and non differencially expressed.

| | | | |
|---|---|---|---|
| **Identification table** | | **The identification table was constructed by analysing the RAW file in MaxQuant, each row contains the group of proteins that could be reconstructed from a set of peptides.** | |
| | ***Protein identifiers*** | ***Protein identifiers include UniProtKB nomenclature and sequence informations from MaxQuant analysis.*** <br> ***Annotations were obtained from the UniProtKB (downloaded and imported via Perseus). Multiple types of annotation serve the purpose of annoting as many identified. proteins as possible.*** | |
| | | UniProt accession number | |
| | | Kegg name | |
| | | Pfam name | |
| | | Uniprot full protein name | |
| | | Kegg pathway or Kegg Brite | Identifiers found manually to enable categorisation and colour-coding |
| | | Peptides | The unique plus razor peptide sequences associated with the protein group. |
| | | Unique peptides | The unique peptide sequences associated with the protein group. |
| | | Sequence coverage [%] | Percentage of the sequence that is covered by the identified unique and |
| | | Unique sequence coverage [%] | Percentage of the sequence that is covered by the identified unique |
| | | Mol. weight [kDa] | Molecular weight of the leading protein sequence contained in the protein group. |
| | ***Quantification Values (Log2 transformed)*** | ***Label-Free quantification values after normalisation by MaxLFQ algorithm of MaxQuant.*** | |
| | | LFQ intensity OrfSwap_1 | |
| | | LFQ intensity OrfSwap_2 | |
| | | LFQ intensity OrfSwap_3 | |
| | | LFQ intensity WT_1 | |
| | | LFQ intensity WT_2 | |
| | | LFQ intensity WT_3 | |

| | | |
|---|---|---|
| ***Statistical analysis*** | ***Dataset filtration and the statistical analysis was performed in Perseus (vesion 1.6.14.0).*** | |
| | Student's T-test Significant OrfSwap_WT | "+" means that the difference in LFQ intensities was found to be statistically |
| | -Log Student's T-test p-value OrfSwap_WT | -Log10 of the p-value obtained with the unpaired, two-tailed Student's T-test. The |
| | Student's T-test q-value OrfSwap_WT | T-test q-value used to sort the protein entries in the assending order |
| | Student's T-test Difference OrfSwap_WT | Log2(Mean of the OrfSwap LFQ intensities) - Log2(Mean of the WT LFQ |

### Downregulated in OrfSwap$^{ihfA\text{-}ihfB}$

| Uniprot accession number | LFQ intensity WT_1 | LFQ intensity WT_2 | LFQ intensity WT_3 | LFQ intensity OrfSwap _1 | LFQ intensity OrfSwap _2 | LFQ intensity OrfSwap _3 | Kegg name | Kegg pathway or Kegg Brite | -Log Student's T-test p-value OrfSwap _WT | Student's T-test q-value OrfSwap_WT | Student's T-test Difference OrfSwap _WT | Cluster | Peptides | Unique peptides | Sequence coverage [%] | Unique sequence coverage [%] | Mol. weight [kDa] |
|---|---|---|---|---|---|---|---|---|---|---|---|---|---|---|---|---|---|
| A0A0H3NGR4 | 32.7418 | 31.674 | 31.7475 | 28.604 | 28.3166 | 26.9522 | rpsQ; 30S ribosomal subunit protein S17 | Translation | 2.57886 | 0 | -4.0968 | Downregulated in OrfSwap | 9 | 9 | 82.1 | 82.1 | 9.722 |
| A0A0H3NST4 | 31.7894 | 31.5064 | 31.5212 | 28.8763 | 29.1345 | 28.8631 | rpmC; 50S ribosomal subunit protein L29 | Translation | 4.49763 | 0 | -2.6477 | Downregulated in OrfSwap | 5 | 5 | 66.7 | 66.7 | 7.26 |
| A0A0H3NIE3 | 30.0971 | 28.3135 | 29.8914 | 27.1019 | 32.4188 | 26.5908 | rplP; 50S ribosomal subunit protein L16 | Translation | 1.82868 | 0 | -2.4522 | Downregulated in OrfSwap | 8 | 8 | 49.3 | 49.3 | 15.19 |
| A0A0H3NIE5 | 30.8955 | 30.9135 | 31.0007 | 28.7123 | 29.1455 | 28.6563 | rplW; 50S ribosomal subunit protein L23 | Translation | 3.73176 | 0 | -2.0985 | Downregulated in OrfSwap | 9 | 9 | 53 | 53 | 11.21 |
| A0A0H3NE85 | 29.9222 | 29.4707 | 29.7858 | 28.452 | 28.5802 | 26.6467 | glnS; glutaminyl-tRNA synthetase | Aminoacyl-tRNA biosynthesis | 1.34309 | 0 | -1.8333 | Downregulated in OrfSwap | 18 | 18 | 34.2 | 34.2 | 63.54 |
| A0A0H3NJ43 | 31.6863 | 32.1787 | 31.9562 | 30.2151 | 30.5471 | 30.5796 | rpmG; 50S ribosomal subunit protein L33 | Translation | 2.90247 | 0 | -1.4931 | Downregulated in OrfSwap | 6 | 6 | 76.4 | 76.4 | 6.372 |
| A0A0H3NSR3 | 32.4492 | 32.3885 | 32.4912 | 31.3934 | 30.9909 | 30.6419 | rpsK; 30S ribosomal subunit protein S11 | Translation | 2.54994 | 0 | -1.4342 | Downregulated in OrfSwap | 12 | 12 | 62.7 | 62.7 | 11.73 |
| A0A0H3NV54 | 32.0063 | 32.0412 | 32.1672 | 30.6266 | 30.543 | 31.0402 | rplJ; 50S ribosomal subunit protein L10 | Translation | 2.93437 | 0 | -1.335 | Downregulated in OrfSwap | 21 | 21 | 94.5 | 94.5 | 17.8 |

### Downregulated in OrfSwap$^{ihfA\text{-}ihfB}$

| Uniprot accession number | LFQ intensity WT_1 | LFQ intensity WT_2 | LFQ intensity WT_3 | LFQ intensity OrfSwap _1 | LFQ intensity OrfSwap _2 | LFQ intensity OrfSwap _3 | Kegg name | Kegg pathway or Kegg Brite | -Log Student's T-test p-value OrfSwap _WT | Student's T-test q-value OrfSwap_WT | Student's T-test Difference OrfSwap _WT | Cluster | Peptides | Unique peptides | Sequence coverage [%] | Unique sequence coverage [%] | Mol. weight [kDa] |
|---|---|---|---|---|---|---|---|---|---|---|---|---|---|---|---|---|---|
| A0A0H3NSS7 | 32.0121 | 31.7291 | 31.9239 | 30.3509 | 30.5665 | 30.7686 | rpsN; 30S ribosomal subunit protein S14 | Translation | 3.08089 | 0 | -1.3264 | Downregulated in OrfSwap | 12 | 12 | 74 | 74 | 11.06 |
| A0A0H3NHU8 | 29.7108 | 29.6223 | 29.3445 | 28.4701 | 28.4556 | 27.994 | aspS; Aspartyl-tRNA synthetase | Aminoacyl-tRNA biosynthesis | 2.55053 | 0 | -1.2526 | Downregulated in OrfSwap | 23 | 23 | 40.8 | 40.8 | 65.67 |
| A0A0H3NMG1 | 31.9654 | 32.1224 | 31.9136 | 30.861 | 30.8724 | 30.7978 | rplC; 50S ribosomal subunit protein L3 | Translation | 4.18211 | 0 | -1.1567 | Downregulated in OrfSwap | 23 | 23 | 62.7 | 62.7 | 22.25 |
| A0A0H3NIV5 | 31.8072 | 31.42 | 31.5218 | 30.6804 | 30.4625 | 30.3041 | rpsR; 30s ribosomal subunit protein S18 | Translation | 2.63955 | 0.0042 | -1.1007 | Downregulated in OrfSwap | 12 | 12 | 66.7 | 66.7 | 8.986 |
| A0A0H3NGR9 | 31.7872 | 31.1082 | 31.6181 | 30.2059 | 30.7111 | 30.1044 | rpsS; 30S ribosomal subunit protein S19 | Translation | 1.86311 | 0.0057 | -1.164 | Downregulated in OrfSwap | 10 | 10 | 76.1 | 76.1 | 10.42 |
| A0A0H3NBF9 | 31.181 | 30.985 | 30.9943 | 30.0884 | 30.0778 | 30.0504 | rpsV; 30S ribosomal protein S22 | Translation | 3.95445 | 0.0058 | -0.9812 | Downregulated in OrfSwap | 6 | 6 | 68.1 | 68.1 | 5.369 |
| A0A0H3NGR0 | 31.2695 | 31.4951 | 31.3384 | 29.9184 | 30.058 | 30.8259 | rl18; L18 Ribosomal Protein | Translation | 1.71707 | 0.0069 | -1.1002 | Downregulated in OrfSwap | 17 | 17 | 66.7 | 66.7 | 12.77 |
| A0A0H3NID8 | 33.1423 | 33.1852 | 33.1794 | 32.3236 | 32.2048 | 32.3316 | rpsE; 30S ribosomal subunit protein S5 | Translation | 4.47141 | 0.0087 | -0.8824 | Downregulated in OrfSwap | 23 | 23 | 94.6 | 94.6 | 17.6 |
| A0A0H3NNN3 | 31.5293 | 31.3719 | 31.3278 | 30.6503 | 30.4286 | 30.4347 | rplY; 50s ribosomal protein L25 | Translation | 3.16623 | 0.0099 | -0.9051 | Downregulated in OrfSwap | 8 | 8 | 79.8 | 79.8 | 10.54 |

### Downregulated in OrfSwap$^{ihfA-ihfB}$

| Uniprot accession number | LFQ intensity WT_1 | LFQ intensity WT_2 | LFQ intensity WT_3 | LFQ intensity OrfSwap_1 | LFQ intensity OrfSwap_2 | LFQ intensity OrfSwap_3 | Kegg name | Kegg pathway or Kegg Brite | -Log Student's T-test p-value OrfSwap_WT | Student's T-test q-value OrfSwap_WT | Student's T-test Difference OrfSwap_WT | Cluster | Peptides | Unique peptides | Sequence coverage [%] | Unique sequence coverage [%] | Mol. weight [kDa] |
|---|---|---|---|---|---|---|---|---|---|---|---|---|---|---|---|---|---|
| A0A0H3NGQ8 | 30.3574 | 30.8635 | 30.7354 | 25.6563 | 25.6792 | 24.6995 | rpsH; 30S ribosomal subunit protein S8 | Translation | 3.92466 | 0 | -5.3071 | WT unique | 10 | 10 | 65.4 | 65.4 | 14.13 |
| A0A0H3NFA7 | 28.7392 | 28.7823 | 28.4217 | 25.2814 | 24.5944 | 25.0508 | rmf; ribosome modulation factor (protein E) | Ribosome biogenesis | 4.03355 | 0 | -3.6722 | WT unique | 5 | 5 | 72.7 | 72.7 | 6.572 |
| A0A0H3NGD6 | 29.358 | 29.2159 | 29.0969 | 25.8424 | 26.1382 | 24.759 | yfiA; hypothetical sigma(54) modulation protein | genetic information processing; Ribosome biogenesis | 2.98906 | 0 | -3.6437 | WT unique | 7 | 7 | 57.1 | 57.1 | 12.65 |
| A0A0H3NHB7 | 28.5665 | 28.3951 | 28.6813 | 26.0194 | 25.0436 | 24.6852 | glyQ; glycine-tRNA synthetase, alpha subunit | Aminoacyl-tRNA biosynthesis | 2.89842 | 0.0022 | -3.2982 | WT unique | 8 | 8 | 35.3 | 35.3 | 34.75 |
| A0A0H3NLU6 | 27.6788 | 27.998 | 27.8323 | 25.7191 | 25.0009 | 24.8476 | prfB; peptide chain release factor 2 (RF-2) | Translation factors | 3.1323 | 0.0034 | -2.6471 | WT unique | 4 | 4 | 12.8 | 12.8 | 40.13 |
| A0A0H3NA20 | 33.5083 | 33.2456 | 33.4173 | 31.482 | 31.5555 | 31.1095 | hupB; DNA-binding protein HU-beta | Genetic information processing; Chromosome and associated proteins; Regulation of transcription | 3.65513 | 0 | -2.0081 | Downregulated in OrfSwap | 13 | 13 | 61.1 | 61.1 | 9.24 |

### Downregulated in OrfSwap$^{ihfA\text{-}ihfB}$

| Uniprot accession number | LFQ intensity WT_1 | LFQ intensity WT_2 | LFQ intensity WT_3 | LFQ intensity OrfSwap _1 | LFQ intensity OrfSwap _2 | LFQ intensity OrfSwap _3 | Kegg name | Kegg pathway or Kegg Brite | -Log Student's T-test p-value OrfSwap _WT | Student's T-test q-value OrfSwap_WT | Student's T-test Difference OrfSwap _WT | Cluster | Peptides | Unique peptides | Sequence coverage [%] | Unique sequence coverage [%] | Mol. weight [kDa] |
|---|---|---|---|---|---|---|---|---|---|---|---|---|---|---|---|---|---|
| A0A0H3NJP8 | 30.1782 | 30.793 | 29.9731 | 27.9155 | 29.2027 | 29.1776 | himD; integration host factor beta-subunit | Genetic information processing; Chromosome and associated proteins; Regulation of transcription | 1.46395 | 0 | -1.5495 | Downregulated in OrfSwap | 7 | 7 | 58.5 | 58.5 | 10.62 |
| A0A0H3N9T3 | 28.8994 | 28.9935 | 28.0829 | 27.311 | 27.4755 | 27.5136 | mukB; cell division protein | Genetic information processing; Chromosome and associated proteins; Chromosome partitioning proteins | 1.84349 | 0.0041 | -1.2252 | Downregulated in OrfSwap | 32 | 32 | 20.4 | 20.4 | 170 |
| A0A0H3NEX8 | 30.5707 | 30.4008 | 30.4136 | 29.3314 | 29.4919 | 29.601 | lrp; leucine-responsive regulatory protein | Genetic information processing; Chromosome and associated proteins; Regulation of transcription | 3.3062 | 0.0055 | -0.987 | Downregulated in OrfSwap | 13 | 13 | 43.3 | 43.3 | 18.86 |
| A0A0H3NDG3 | 28.6884 | 28.7489 | 28.5633 | 24.1678 | 23.9782 | 25.7234 | topA; DNA topoisomerase I, omega protein I | Genetic information processing; Isomerases | 2.72284 | 0 | -4.0437 | WT unique | 21 | 21 | 29.9 | 29.9 | 97.3 |

### Downregulated in OrfSwap$^{ihfA-ihfB}$

| Uniprot accession number | LFQ intensity WT_1 | LFQ intensity WT_2 | LFQ intensity WT_3 | LFQ intensity OrfSwap _1 | LFQ intensity OrfSwap _2 | LFQ intensity OrfSwap _3 | Kegg name | Kegg pathway or Kegg Brite | -Log Student's T-test p-value OrfSwap _WT | Student's T-test q-value OrfSwap_WT | Student's T-test Difference OrfSwap _WT | Cluster | Peptides | Unique peptides | Sequence coverage [%] | Unique sequence coverage [%] | Mol. weight [kDa] |
|---|---|---|---|---|---|---|---|---|---|---|---|---|---|---|---|---|---|
| A0A0H3NVG2 | 28.6116 | 28.2947 | 28.6507 | 24.9511 | 24.0273 | 24.9687 | ssb; single-strand DNA-binding protein | Genetic Information Processing; DNA replication; mismatch repair | 3.51576 | 0 | -3.87 | WT unique | 7 | 7 | 28.4 | 28.4 | 19.07 |
| A0A0H3NIJ8 | 28.3055 | 27.5692 | 28.1214 | 23.8103 | 24.296 | 24.8487 | gyrI; hypothetical DNA gyrase inhibitory protein | Genetic information processing; Replication and repair | 3.22903 | 0 | -3.6804 | WT unique | 6 | 6 | 33.5 | 33.5 | 18.06 |
| A0A0H3NHP0 | 28.5757 | 28.605 | 28.4982 | 24.9141 | 24.6981 | 25.3174 | gyrB; DNA gyrase subunit B | Genetic information processing; Isomerases | 4.38502 | 0 | -3.5831 | WT unique | 19 | 19 | 18.5 | 18.5 | 89.9 |
| A0A0H3NHZ9 | 28.2363 | 28.1298 | 28.2169 | 24.7374 | 24.8031 | 25.0236 | polA; DNA polymerase I | Genetic Information Processing; DNA replication | 5.45381 | 0 | -3.3396 | WT unique | 11 | 11 | 14.7 | 14.7 | 103.1 |
| A0A0H3NV13 | 28.3698 | 28.3983 | 28.1877 | 24.6317 | 25.4968 | 25.175 | menG; menaquinone biosynthesis protein | Genetic information processing; Bacterial mRNA degradation factor | 3.60455 | 0 | -3.2174 | WT unique | 7 | 7 | 42.2 | 42.2 | 17.37 |

### Downregulated in OrfSwap$^{ihfA\text{-}ihfB}$

| Uniprot accession number | LFQ intensity WT_1 | LFQ intensity WT_2 | LFQ intensity WT_3 | LFQ intensity OrfSwap_1 | LFQ intensity OrfSwap_2 | LFQ intensity OrfSwap_3 | Kegg name | Kegg pathway or Kegg Brite | -Log Student's T-test p-value OrfSwap_WT | Student's T-test q-value OrfSwap_WT | Student's T-test Difference OrfSwap_WT | Cluster | Peptides | Unique peptides | Sequence coverage [%] | Unique sequence coverage [%] | Mol. weight [kDa] |
|---|---|---|---|---|---|---|---|---|---|---|---|---|---|---|---|---|---|
| A0A0H3NF89 | 27.8378 | 28.1916 | 28.2758 | 25.0186 | 25.2627 | 25.0742 | hilA; invasion protein regulator | Genetic information procession; regulation of transcription | 4.38728 | 0 | -2.9832 | WT unique | 15 | 15 | 27.5 | 27.5 | 63.04 |
| A0A0H3NXF0 | 27.0453 | 27.1707 | 27.0913 | 24.39 | 24.0192 | 24.4176 | parA; plasmid partition protein A | Genetic information processing; Plasmid DNA re;lication | 4.53077 | 0.0026 | -2.8269 | WT unique | 7 | 7 | 17.7 | 17.7 | 44.5 |
| A0A0H3N7T4 | 29.9047 | 31.6913 | 31.6005 | 28.2944 | 28.1613 | 28.5464 | hlpA; outer membrane protein OmpH precursor | Unclassified: signaling and cellular processes | 2.00355 | 0 | -2.7315 | Downregulated in OrfSwap | 10 | 10 | 57.8 | 57.8 | 17.91 |
| A0A0H3NCM8 | 36.1877 | 36.0946 | 36.0741 | 33.8888 | 34.3397 | 34.1492 | lpp; major outer membrane lipoprotein | Peptidoglycan biosynthesis and degradation proteins | 3.9079 | 0 | -1.9929 | Downregulated in OrfSwap | 11 | 11 | 66.7 | 66.7 | 8.391 |
| A0A0H3NJJ9 | 31.283 | 31.5632 | 31.3099 | 29.8331 | 30.2186 | 29.972 | glpT; glycerol-3-phosphate transporter | Phosphate and organophosphate transporters | 3.17835 | 0 | -1.3775 | Downregulated in OrfSwap | 16 | 16 | 22.3 | 22.3 | 50.18 |
| A0A0H3P177 | 30.7412 | 30.7921 | 30.9653 | 29.7188 | 29.7217 | 29.7917 | traT; conjugative transfer, surface exclusion protein | conjugative transfer, surface exclusion protein | 3.95515 | 0 | -1.0888 | Downregulated in OrfSwap | 12 | 12 | 46.1 | 46.1 | 26.17 |

### Downregulated in OrfSwap$^{ihfA\text{-}ihfB}$

| Uniprot accession number | LFQ intensity WT_1 | LFQ intensity WT_2 | LFQ intensity WT_3 | LFQ intensity OrfSwap_1 | LFQ intensity OrfSwap_2 | LFQ intensity OrfSwap_3 | Kegg name | Kegg pathway or Kegg Brite | -Log Student's T-test p-value OrfSwap_WT | Student's T-test q-value OrfSwap_WT | Student's T-test Difference OrfSwap_WT | Cluster | Peptides | Unique peptides | Sequence coverage [%] | Unique sequence coverage [%] | Mol. weight [kDa] |
|---|---|---|---|---|---|---|---|---|---|---|---|---|---|---|---|---|---|
| A0A0H3N9F1 | 27.69 | 27.6129 | 27.6158 | 26.3882 | 26.8304 | 25.9986 | rcsF; RcsF protein | Signal transduction; Two-component system | 2.15823 | 0.002 | -1.2339 | Downregulated in OrfSwap | 4 | 4 | 43.3 | 43.3 | 14.19 |
| A0A0H3NAN8 | 31.902 | 33.0883 | 33.0054 | 30.981 | 31.6268 | 31.6706 | osmE; osmotically inducible lipoprotein E precursor | Unclassified: signaling and cellular processes; Structural proteins | 1.31131 | 0.007 | -1.2391 | Downregulated in OrfSwap | 8 | 8 | 72.6 | 72.6 | 12.16 |
| A0A0H3NNU6 | 29.0333 | 28.3244 | 28.0774 | 27.2803 | 27.4396 | 27.4219 | glpF; glycerol uptake facilitator protein | Transporters; Aquaporins and small neutral solute transporters | 1.70913 | 0.0088 | -1.0977 | Downregulated in OrfSwap | 3 | 3 | 8.5 | 8.5 | 29.69 |
| A0A0H3NK15 | 28.1404 | 27.9784 | 28.0586 | 24.8839 | 24.6116 | 24.3896 | putP; sodium/proline symporter (proline permease) | Transporters; Electrochemical potential-driven transporters | 4.65997 | 0 | -3.4308 | WT unique | 5 | 5 | 12.4 | 12.4 | 54.32 |
| A0A0H3NIF8 | 27.8286 | 27.4577 | 28.0793 | 24.9015 | 24.4912 | 25.5211 | ybaY; conserved hypothetical lipoprotein | General function prediction only | 2.89026 | 0.0016 | -2.8172 | WT unique | 5 | 5 | 64 | 64 | 19.48 |
| A0A0H3NEM7 | 29.5823 | 27.6187 | 28.1154 | 25.0247 | 24.5587 | 25.0029 | ompX; outer membrane protein x precursor | Transporters; Pores ion channels | 2.37775 | 0.0019 | -3.5767 | WT unique | 5 | 5 | 31.6 | 31.6 | 18.49 |

### Downregulated in OrfSwap$^{ihfA\text{-}ihfB}$

| Uniprot accession number | LFQ intensity WT_1 | LFQ intensity WT_2 | LFQ intensity WT_3 | LFQ intensity OrfSwap_1 | LFQ intensity OrfSwap_2 | LFQ intensity OrfSwap_3 | Kegg name | Kegg pathway or Kegg Brite | -Log Student's T-test p-value OrfSwap_WT | Student's T-test q-value OrfSwap_WT | Student's T-test Difference OrfSwap_WT | Cluster | Peptides | Unique peptides | Sequence coverage [%] | Unique sequence coverage [%] | Mol. weight [kDa] |
|---|---|---|---|---|---|---|---|---|---|---|---|---|---|---|---|---|---|
| A0A0H3NHY1 | 28.0233 | 27.8087 | 27.981 | 24.9447 | 24.3699 | 25.3016 | tatA; sec-independent protein translocase protein | Protein export; Bacterial secretion system | 3.40779 | 0.0024 | -3.0656 | WT unique | 3 | 3 | 40.5 | 40.5 | 8.944 |
| A0A0H3N7S7 | 27.3886 | 27.2631 | 26.843 | 25.1768 | 24.2187 | 25.046 | hypothetical lipoprotein | General function prediction only | 2.62825 | 0.0043 | -2.3511 | WT unique | 2 | 2 | 47.4 | 47.4 | 8.217 |
| A0A0H3NM92 | 27.4465 | 26.681 | 27.3945 | 23.5621 | 25.1878 | 23.7653 | yhbG; probable ABC transport protein, ATP-binding component | ABC transporters | 2.21157 | 0.0046 | -3.0023 | WT unique | 6 | 6 | 32 | 32 | 26.8 |
| A0A0H3NF33 | 28.3266 | 28.2222 | 28.4165 | 25.5974 | 25.7336 | 26.3525 | proX; glycine betaine-binding periplasmic protein precursor | ABC transporters | 3.27601 | 0.0048 | -2.4273 | WT unique | 5 | 5 | 22.1 | 22.1 | 36.15 |
| A0A0H3NA56 | 27.2624 | 27.2044 | 27.1894 | 25.5529 | 24.6316 | 25.0441 | copA; copper-transporting ATPase | Translocases | 2.88093 | 0.0066 | -2.1426 | WT unique | 3 | 3 | 4.8 | 4.8 | 87.91 |
| A0A0H3NQ36 | 27.8297 | 28.0537 | 27.914 | 25.4824 | 26.366 | 24.7049 | yfiO; outer membrane protein assembly complex subunit yfiO | Transporters; Pores ion channels | 2.12122 | 0.009 | -2.4147 | WT unique | 11 | 11 | 51.4 | 51.4 | 27.83 |
| A0A0H3NMB9 | 30.3836 | 30.1613 | 30.2868 | 27.6154 | 27.7263 | 27.1208 | mreB; rod shape-determining protein | Cytoskeleton proteins | 3.84113 | 0 | -2.7897 | Downregulated in OrfSwap | 17 | 17 | 44.4 | 44.4 | 36.95 |

### Downregulated in OrfSwap$^{ihfA-ihfB}$

| Uniprot accession number | LFQ intensity WT_1 | LFQ intensity WT_2 | LFQ intensity WT_3 | LFQ intensity OrfSwap _1 | LFQ intensity OrfSwap _2 | LFQ intensity OrfSwap _3 | Kegg name | Kegg pathway or Kegg Brite | -Log Student's T-test p-value OrfSwap _WT | Student's T-test q-value OrfSwap_WT | Student's T-test Difference OrfSwap _WT | Cluster | Peptides | Unique peptides | Sequence coverage [%] | Unique sequence coverage [%] | Mol. weight [kDa] |
|---|---|---|---|---|---|---|---|---|---|---|---|---|---|---|---|---|---|
| A0A0H3NII6 | 27.3902 | 27.2823 | 27.188 | 25.7563 | 26.7898 | 26.2175 | damX; cytoplasmic membrane protein with SPOR domain | Cytoskeleton proteins | 1.55996 | 0.0087 | -1.0323 | Downregulated in OrfSwap | 9 | 9 | 26.8 | 26.8 | 45.46 |
| A0A0H3NPT7 | 31.5046 | 31.621 | 31.2891 | 29.0566 | 28.9341 | 29.9531 | glyA; serine hydroxymethyltransferase | Glycine, serine and threonine metabolism | 2.52096 | 0 | -2.157 | Downregulated in OrfSwap | 31 | 31 | 54.7 | 54.7 | 45.45 |
| A0A0H3NL37 | 30.5684 | 30.3859 | 30.3076 | 28.6955 | 28.5933 | 27.52 | pepA; cytosol aminopeptidase | Glutathione metabolism | 2.30291 | 0 | -2.151 | Downregulated in OrfSwap | 16 | 16 | 40 | 40 | 54.95 |
| A0A0H3N9G9 | 29.1035 | 29.0712 | 29.4407 | 28.3664 | 26.8333 | 26.6494 | yliJ; glutathione s-transferase family protein | Glutathione metabolism | 1.58456 | 0 | -1.9221 | Downregulated in OrfSwap | 9 | 9 | 39.4 | 39.4 | 23.69 |
| A0A0H3NIM9 | 30.1236 | 30.0961 | 30.0237 | 28.5506 | 28.2989 | 28.6372 | phoN; nonspecific acid phosphatase precursor | Riboflavin metabolism | 3.93825 | 0 | -1.5855 | Downregulated in OrfSwap | 12 | 12 | 56.4 | 56.4 | 28.38 |
| A0A0H3NIE8 | 29.9499 | 29.6014 | 30.1481 | 28.5622 | 28.3124 | 28.7206 | pfkA; 6-phosphofructokinase | Glycolysis / Gluconeogenesis | 2.62874 | 0 | -1.3681 | Downregulated in OrfSwap | 11 | 11 | 34.1 | 34.1 | 34.92 |
| A0A0H3NIA0 | 27.6627 | 27.4573 | 27.337 | 26.3432 | 26.2754 | 26.5256 | nanE; hypothetical N-acetylmannosamine-6-phosphate 2-epimerase 2 (ec 5.1.3.9) (mannac-6-p epimerase 2) | Amino sugar and nucleotide sugar metabolism | 3.09807 | 0.0019 | -1.1043 | Downregulated in OrfSwap | 4 | 4 | 16.2 | 16.2 | 24.03 |
| A0A0H3NC31 | 32.3683 | 32.2929 | 32.2204 | 31.3583 | 31.1325 | 31.4097 | hypothetical glutamate dehydrogenase | Alanine, aspartate and glutamate metabolism | 3.32191 | 0.0056 | -0.9937 | Downregulated in OrfSwap | 33 | 33 | 68 | 68 | 48.04 |

### Downregulated in OrfSwap$^{ihfA-ihfB}$

| Uniprot accession number | LFQ intensity WT_1 | LFQ intensity WT_2 | LFQ intensity WT_3 | LFQ intensity OrfSwap_1 | LFQ intensity OrfSwap_2 | LFQ intensity OrfSwap_3 | Kegg name | Kegg pathway or Kegg Brite | -Log Student's T-test p-value OrfSwap_WT | Student's T-test q-value OrfSwap_WT | Student's T-test Difference OrfSwap_WT | Cluster | Peptides | Unique peptides | Sequence coverage [%] | Unique sequence coverage [%] | Mol. weight [kDa] |
|---|---|---|---|---|---|---|---|---|---|---|---|---|---|---|---|---|---|
| A0A0H3NE44 | 30.9651 | 31.1186 | 30.9708 | 30.2448 | 30.1718 | 30.1105 | cysK; cysteine synthase A | Cysteine and methionine metabolism | 3.72936 | 0.009 | -0.8425 | Downregulated in OrfSwap | 19 | 19 | 61.3 | 61.3 | 34.54 |
| A0A0H3N8Z2 | 28.8973 | 28.1287 | 28.6084 | 27.0186 | 28.0261 | 27.3169 | asnB; asparagine synthetase B | Alanine, aspartate and glutamate metabolism | 1.36444 | 0.0091 | -1.0909 | Downregulated in OrfSwap | 14 | 14 | 37.7 | 37.7 | 62.57 |
| A0A0H3NJ96 | 28.6777 | 28.4375 | 28.6262 | 27.7626 | 27.5856 | 27.7064 | moaB; molybdenum cofactor biosynthesis protein B | Folate biosynthesis | 3.24616 | 0.0096 | -0.8956 | Downregulated in OrfSwap | 9 | 9 | 53.5 | 53.5 | 18.54 |
| A0A0H3NQU8 | 29.0522 | 29.275 | 29.3292 | 23.7036 | 25.7339 | 24.6615 | luxS; S-ribosylhomocysteinase | Cysteine and methionine metabolism; Quorum sensing | 2.79951 | 0 | -4.5191 | WT unique | 12 | 12 | 51.5 | 51.5 | 19.31 |
| A0A0H3NE99 | 28.5288 | 28.3718 | 28.2848 | 23.9503 | 24.5388 | 24.2508 | guaB; inosine-5'-monophosphate dehydrogenase | Purine metabolism | 4.63693 | 0 | -4.1485 | WT unique | 6 | 6 | 16.8 | 16.8 | 51.95 |
| A0A0H3NAA4 | 29.0752 | 29.0026 | 28.8242 | 25.1261 | 24.8356 | 24.5857 | ptsG; PTS system, glucose-specific IIBC component | Glycolysis / Gluconeogenesis | 4.73321 | 0 | -4.1182 | WT unique | 12 | 12 | 17.8 | 17.8 | 50.5 |
| A0A0H3N866 | 28.7187 | 28.8576 | 28.5595 | 25.0252 | 24.8678 | 23.9194 | proC; pyrroline-5-carboxylate reductase | Arginine and proline metabolism | 3.49222 | 0 | -4.1078 | WT unique | 9 | 9 | 28.3 | 28.3 | 28.06 |
| A0A0H3NUL5 | 28.7276 | 28.6842 | 28.7611 | 25.5831 | 24.6112 | 24.0706 | fadA; small (beta) subunit of the fatty acid-oxidizing multienzyme complex | Fatty acid degradation | 3.06609 | 0 | -3.9693 | WT unique | 10 | 10 | 24 | 24 | 41 |

#### Downregulated in OrfSwap$^{ihfA\text{-}ihfB}$

| Uniprot accession number | LFQ intensity WT_1 | LFQ intensity WT_2 | LFQ intensity WT_3 | LFQ intensity OrfSwap_1 | LFQ intensity OrfSwap_2 | LFQ intensity OrfSwap_3 | Kegg name | Kegg pathway or Kegg Brite | -Log Student's T-test p-value OrfSwap_WT | Student's T-test q-value OrfSwap_WT | Student's T-test Difference OrfSwap_WT | Cluster | Peptides | Unique peptides | Sequence coverage [%] | Unique sequence coverage [%] | Mol. weight [kDa] |
|---|---|---|---|---|---|---|---|---|---|---|---|---|---|---|---|---|---|
| A0A0H3N894 | 28.706 | 28.594 | 28.3689 | 24.9273 | 24.2324 | 24.8206 | accA; acetyl-coenzyme A carboxylase carboxyl transferase subunit alpha | Fatty acid biosynthesis | 4.0911 | 0 | -3.8962 | WT unique | 11 | 11 | 33.5 | 33.5 | 35.34 |
| A0A0H3NE87 | 29.2233 | 28.8877 | 29.059 | 25.8753 | 25.633 | 24.6033 | dapA; Dihydrodipicolinate synthase | Lysine biosynthesis | 3.10565 | 0 | -3.6861 | WT unique | 11 | 11 | 50.7 | 50.7 | 31.29 |
| A0A0H3NKD9 | 28.874 | 28.8394 | 28.8375 | 25.0415 | 25.2096 | 25.2921 | gcpE; 4-hydroxy-3-methylbut-2-en-1-yl diphosphate synthase | Terpenoid backbone biosynthesis | 5.98834 | 0 | -3.6692 | WT unique | 10 | 10 | 30.6 | 30.6 | 40.55 |
| A0A0H3NNJ9 | 28.3533 | 28.0967 | 28.0895 | 24.9876 | 24.6338 | 24.2107 | yeiG; hypothetical esterase | Energy metabolism; Methane metabolism | 3.91879 | 0 | -3.5691 | WT unique | 6 | 6 | 16.8 | 16.8 | 31.97 |
| A0A0H3NJY4 | 28.5546 | 27.9497 | 28.3016 | 24.6025 | 24.6516 | 25.1013 | conserved hypothetical protein | Function unknown | 3.9075 | 0 | -3.4835 | WT unique | 5 | 5 | 59.4 | 59.4 | 14.91 |
| A0A0H3NP42 | 27.7515 | 27.821 | 27.7342 | 24.8067 | 24.609 | 24.1086 | pdxB; Erythronate-4-phosphate dehydrogenase | Vitamin B6 metabolism | 4.00288 | 0 | -3.2608 | WT unique | 12 | 12 | 36 | 36 | 41.3 |
| A0A0H3N9A8 | 28.3787 | 28.2061 | 28.2145 | 24.8662 | 25.5054 | 24.8415 | yadR; conserved hypothetical protein | General function prediction only, iron-sulfur cluster assembly | 3.84972 | 0 | -3.1954 | WT unique | 4 | 4 | 36.8 | 36.8 | 12.1 |

### Downregulated in OrfSwap$^{ihfA\text{-}ihfB}$

| Uniprot accession number | LFQ intensity WT_1 | LFQ intensity WT_2 | LFQ intensity WT_3 | LFQ intensity OrfSwap_1 | LFQ intensity OrfSwap_2 | LFQ intensity OrfSwap_3 | Kegg name | Kegg pathway or Kegg Brite | -Log Student's T-test p-value OrfSwap_WT | Student's T-test q-value OrfSwap_WT | Student's T-test Difference OrfSwap_WT | Cluster | Peptides | Unique peptides | Sequence coverage [%] | Unique sequence coverage [%] | Mol. weight [kDa] |
|---|---|---|---|---|---|---|---|---|---|---|---|---|---|---|---|---|---|
| A0A0H3N7F3 | 28.9772 | 28.8858 | 29.0756 | 25.9895 | 26.1626 | 25.6027 | carB; carbamoyl-phosphate synthase large chain | Alanine, aspartate and glutamate metabolism | 4.20908 | 0 | -3.0612 | WT unique | 27 | 27 | 30 | 30 | 118.1 |
| A0A0H3ND10 | 28.026 | 28.0589 | 27.9616 | 25.0731 | 24.8611 | 25.3712 | speG; spermidine N1-acetyltransferase | Arginine and proline metabolism | 4.37503 | 0 | -2.9137 | WT unique | 4 | 4 | 31.7 | 31.7 | 22.05 |
| A0A0H3NEE2 | 28.0983 | 28.0177 | 28.012 | 24.4238 | 25.4347 | 25.6358 | pdxJ; Pyridoxal phosphate biosynthetic protein pdxJ | Vitamin B6 metabolism | 2.80505 | 0.0017 | -2.8779 | WT unique | 6 | 6 | 31.7 | 31.7 | 26.34 |
| A0A0H3NKF9 | 28.5283 | 28.6327 | 28.4137 | 24.8514 | 26.2431 | 25.5099 | iscS; Cysteine desulfurase | Thiamine metabolism | 2.73854 | 0.0017 | -2.9901 | WT unique | 10 | 10 | 28.7 | 28.7 | 45.08 |
| A0A0H3NJP0 | 27.8652 | 27.9186 | 27.7703 | 24.9528 | 24.6525 | 25.6034 | nuoA; NADH dehydrogenase I chain A | Energy metabolism; Oxidative phosphorylation | 3.21549 | 0.0018 | -2.7818 | WT unique | 3 | 3 | 20.4 | 20.4 | 16.49 |
| A0A0H3NMS7 | 27.9204 | 28.1168 | 28.1086 | 24.1996 | 25.4548 | 25.2623 | kdgK; 2-dehydro-3-deoxygluconokinase | Pentose phosphate pathway | 2.83176 | 0.0019 | -3.0764 | WT unique | 7 | 7 | 23.3 | 23.3 | 34.09 |
| A0A0H3NDH4 | 27.2824 | 27.7954 | 27.7062 | 25.0347 | 24.7559 | 25.0799 | dld; D-lactate dehydrogenase | Pyruvate metabolism | 3.82589 | 0.0019 | -2.6378 | WT unique | 11 | 11 | 21.9 | 21.9 | 65.05 |
| E1WA38 | 28.2192 | 28.2349 | 28.2159 | 24.8905 | 24.4284 | 25.8341 | csiD; Protein csiD | Lysine degradation | 2.80837 | 0.002 | -3.1723 | WT unique | 15 | 15 | 44.9 | 44.9 | 37.23 |

### Downregulated in OrfSwap$^{ihfA\text{-}ihfB}$

| Uniprot accession number | LFQ intensity WT_1 | LFQ intensity WT_2 | LFQ intensity WT_3 | LFQ intensity OrfSwap_1 | LFQ intensity OrfSwap_2 | LFQ intensity OrfSwap_3 | Kegg name | Kegg pathway or Kegg Brite | -Log Student's T-test p-value OrfSwap_WT | Student's T-test q-value OrfSwap_WT | Student's T-test Difference OrfSwap_WT | Cluster | Peptides | Unique peptides | Sequence coverage [%] | Unique sequence coverage [%] | Mol. weight [kDa] |
|---|---|---|---|---|---|---|---|---|---|---|---|---|---|---|---|---|---|
| A0A0H3NBV8 | 27.5574 | 27.6786 | 27.4421 | 24.4987 | 24.0018 | 25.1294 | adhC; alcohol dehydrogenase class III (ec 1.1.1.1) (glutathione-dependent formaldehyde dehydrogenase) (ec 1.2.1.1) (fdh) (faldh) | Glycolysis / Gluconeogenesis | 3.08261 | 0.002 | -3.016 | WT unique | 8 | 8 | 25.3 | 25.3 | 39.26 |
| A0A0H3NIE9 | 28.1517 | 28.4912 | 27.8466 | 25.0388 | 24.0571 | 25.4402 | bfr; bacterioferritin | Porphyrin and chlorophyll metabolism | 2.74011 | 0.002 | -3.3178 | WT unique | 5 | 5 | 38 | 38 | 18.36 |
| A0A0H3NDN6 | 28.6483 | 28.4957 | 28.3476 | 24.3859 | 24.7148 | 25.9737 | conserved hypothetical protein | General function prediction only | 2.67382 | 0.0021 | -3.4724 | WT unique | 7 | 7 | 87.7 | 87.7 | 6.482 |
| A0A0H3N8M9 | 27.8033 | 27.7418 | 27.6006 | 24.7167 | 24.7796 | 25.2736 | prpB; carboxyvinyl-carboxyphosphonate phosphorylmutase | Propanoate metabolism | 3.9401 | 0.0023 | -2.7919 | WT unique | 8 | 8 | 30.2 | 30.2 | 32 |
| A0A0H3NEV4 | 28.0449 | 28.0206 | 27.8034 | 24.3547 | 24.6691 | 25.4018 | mrp; conserved hypothetical protein | Iron-sulfur cluster binding | 3.22496 | 0.0023 | -3.1478 | WT unique | 5 | 5 | 14.9 | 14.9 | 39.93 |
| A0A0H3NH18 | 27.6486 | 27.9082 | 27.7491 | 25.184 | 25.0045 | 25.0861 | glgC; glucose-1-phosphate adenylyltransferase | Starch and sucrose metabolism | 5.08647 | 0.0025 | -2.6771 | WT unique | 10 | 10 | 26.7 | 26.7 | 48.46 |

### Downregulated in OrfSwap$^{ihfA-ihfB}$

| Uniprot accession number | LFQ intensity WT_1 | LFQ intensity WT_2 | LFQ intensity WT_3 | LFQ intensity OrfSwap_1 | LFQ intensity OrfSwap_2 | LFQ intensity OrfSwap_3 | Kegg name | Kegg pathway or Kegg Brite | -Log Student's T-test p-value OrfSwap_WT | Student's T-test q-value OrfSwap_WT | Student's T-test Difference OrfSwap_WT | Cluster | Peptides | Unique peptides | Sequence coverage [%] | Unique sequence coverage [%] | Mol. weight [kDa] |
|---|---|---|---|---|---|---|---|---|---|---|---|---|---|---|---|---|---|
| A0A0H3NJW2 | 27.7927 | 27.6008 | 27.4754 | 25.0249 | 25.4541 | 25.35 | yihX; hypothetical haloacid dehalogenase-like hydrolase | Glycolysis / Gluconeogenesis | 3.91326 | 0.0033 | -2.3466 | WT unique | 7 | 7 | 31.2 | 31.2 | 22.72 |
| A0A0H3NC52 | 27.1407 | 27.5607 | 27.3897 | 25.033 | 25.0758 | 24.6399 | yciK; hypothetical oxidoreductase | General function prediction only | 3.72735 | 0.0034 | -2.4475 | WT unique | 11 | 11 | 44.3 | 44.3 | 28.07 |
| A0A0H3NFF4 | 27.9475 | 27.8398 | 27.8899 | 25.7076 | 25.377 | 24.8891 | accD; acetyl-CoA carboxylase beta subunit | Fatty acid biosynthesis; Pyruvate biosynthesis | 3.36595 | 0.0035 | 0.62947 | WT unique | 11 | 11 | 30.3 | 30.3 | 33.22 |
| A0A0H3NJ06 | 27.1276 | 26.7796 | 26.978 | 24.9425 | 25.0646 | 24.9363 | nagA; N-acetylglucosamine-6-phosphate deacetylase | Amino sugar and nucleotide sugar metabolism | 4.26617 | 0.0044 | -1.9806 | WT unique | 9 | 9 | 24 | 24 | 41.11 |
| A0A0H3NFV9 | 28.1103 | 27.8001 | 28.067 | 24.8644 | 24.5463 | 26.245 | yggV; HAM1 protein homolog | Purine metabolism | 2.19515 | 0.0044 | -2.7739 | WT unique | 7 | 7 | 42.1 | 42.1 | 21.03 |
| A0A0H3NFW7 | 27.3607 | 27.3418 | 27.3665 | 25.1336 | 24.0358 | 25.281 | selD; Selenide,water dikinase | Selenocompound metabolism | 2.53023 | 0.0045 | -2.5396 | WT unique | 6 | 6 | 20.5 | 20.5 | 36.44 |
| A0A0H3NPK1 | 28.5238 | 28.6371 | 28.7289 | 26.7585 | 24.7101 | 24.8773 | purC; phosphoribosylaminoimidazole-succinocarboxamide synthase | Purine metabolism | 2.07091 | 0.0047 | -3.1813 | WT unique | 8 | 8 | 40.5 | 40.5 | 26.91 |
| A0A0H3NMA2 | 28.0305 | 27.3128 | 26.9765 | 24.8722 | 24.2002 | 25.0001 | nanA; N-acetylneuraminate lyase | Amino sugar and nucleotide sugar metabolism | 2.63866 | 0.0049 | -2.7491 | WT unique | 9 | 9 | 25.3 | 25.3 | 32.48 |

### Downregulated in OrfSwap$^{ihfA-ihfB}$

| Uniprot accession number | LFQ intensity WT_1 | LFQ intensity WT_2 | LFQ intensity WT_3 | LFQ intensity OrfSwap _1 | LFQ intensity OrfSwap _2 | LFQ intensity OrfSwap _3 | Kegg name | Kegg pathway or Kegg Brite | -Log Student's T-test p-value OrfSwap _WT | Student's T-test q-value OrfSwap_WT | Student's T-test Difference OrfSwap _WT | Cluster | Peptides | Unique peptides | Sequence coverage [%] | Unique sequence coverage [%] | Mol. weight [kDa] |
|---|---|---|---|---|---|---|---|---|---|---|---|---|---|---|---|---|---|
| A0A0H3NHD1 | 28.4447 | 28.3724 | 28.5579 | 24.6204 | 25.3611 | 26.4047 | galU; glucose-1-phosphate uridylyltransferase | Pentose and glucuronate interconversions | 2.34557 | 0.0049 | -2.9963 | WT unique | 8 | 8 | 28.1 | 28.1 | 32.91 |
| A0A0H3NHQ9 | 28.6889 | 28.2751 | 28.4719 | 25.4666 | 25.0056 | 26.4073 | glmS; glucosamine--fructose-6-phosphate aminotransferase | Alanine, aspartate and glutamate metabolism | 2.57392 | 0.005 | -2.8522 | WT unique | 18 | 18 | 36 | 36 | 66.88 |
| A0A0H3NXX3 | 28.4199 | 28.5144 | 28.6493 | 25.9601 | 24.7315 | 26.1741 | sulII; dihydropteroate synthase type-2 (dihydropteroate synthase type ii) (dhps) (dihydropteroate pyrophosphorylase type ii) | Folate biosynthesis | 2.51304 | 0.0051 | -2.906 | WT unique | 10 | 10 | 46.1 | 46.1 | 28.47 |
| A0A0H3NFD4 | 28.1221 | 27.8415 | 27.8771 | 24.4831 | 26.1489 | 25.4712 | conserved hypothetical protein | General function prediction only | 2.19969 | 0.0054 | -2.5792 | WT unique | 5 | 5 | 26.1 | 26.1 | 20.91 |
| A0A0H3NFY4 | 27.7238 | 27.55 | 27.8754 | 25.3731 | 25.7989 | 25.9303 | uxaC; Uronate isomerase | Pentose and glucuronate interconversions | 3.32641 | 0.0065 | -2.0156 | WT unique | 8 | 8 | 19.1 | 19.1 | 53.61 |
| A0A0H3NHK3 | 27.6659 | 27.6868 | 27.6859 | 25.1752 | 24.3323 | 26.0989 | lpxA; Acyl-[acyl-carrier-protein]--UDP-N-acetylglucos amine O-acyltransferase | Lipopolysaccharide biosynthesis | 2.08071 | 0.0077 | -2.4774 | WT unique | 7 | 7 | 42.4 | 42.4 | 28.09 |

### Downregulated in OrfSwap$^{ihfA-ihfB}$

| Uniprot accession number | LFQ intensity WT_1 | LFQ intensity WT_2 | LFQ intensity WT_3 | LFQ intensity OrfSwap _1 | LFQ intensity OrfSwap _2 | LFQ intensity OrfSwap _3 | Kegg name | Kegg pathway or Kegg Brite | -Log Student's T-test p-value OrfSwap _WT | Student's T-test q-value OrfSwap_WT | Student's T-test Difference OrfSwap _WT | Cluster | Peptides | Unique peptides | Sequence coverage [%] | Unique sequence coverage [%] | Mol. weight [kDa] |
|---|---|---|---|---|---|---|---|---|---|---|---|---|---|---|---|---|---|
| A0A0H3NBZ4 | 26.9379 | 27.0027 | 27.1399 | 24.308 | 25.3498 | 24.948 | purU; formyltetrahydrofolate deformylase | Glyoxylate and dicarboxylate metabolism | 2.65475 | 0.0078 | -2.1582 | WT unique | 6 | 6 | 31.4 | 31.4 | 31.83 |
| A0A0H3NIR2 | 27.2161 | 27.0739 | 26.9158 | 24.063 | 25.4908 | 24.8361 | glgB; 1,4-alpha-glucan branching enzyme | Starch and sucrose metabolism | 2.2413 | 0.0086 | -2.272 | WT unique | 8 | 8 | 13.2 | 13.2 | 84.27 |
| A0A0H3NFW8 | 27.2732 | 27.2466 | 26.9594 | 25.1129 | 23.7361 | 25.3218 | yggX; conserved hypothetical protein | General function prediction only | 2.06332 | 0.0089 | -2.4361 | WT unique | 8 | 8 | 60.4 | 60.4 | 10.9 |
| A0A0H3NL58 | 31.575 | 31.0535 | 31.4667 | 30.0823 | 29.8715 | 29.2774 | sicA; type III secretion-associated chaperone | Pathogenisis; Type III secretion system | 2.30646 | 0 | -1.6213 | Downregulated in OrfSwap | 9 | 9 | 57.6 | 57.6 | 19.22 |
| A0A0H3NF99 | 29.6203 | 30.149 | 29.8881 | 28.7604 | 28.6278 | 28.1962 | invB; chaperone protein for type III secretion system effectors | signaling and cellular processes; Type III secretion system | 2.39382 | 0 | -1.3577 | Downregulated in OrfSwap | 4 | 4 | 41.5 | 41.5 | 14.92 |
| A0A0H3NF82 | 29.3335 | 29.2609 | 28.8951 | 25.5643 | 24.8829 | 25.2882 | prgI; type III secretion system apparatus | Pathogenisis; Type III secretion system | 4.0845 | 0 | -3.918 | WT unique | 6 | 6 | 76.2 | 76.2 | 8.857 |
| A0A0H3NF08 | 28.7756 | 28.6228 | 28.4086 | 25.7285 | 24.0574 | 25.3258 | pipB2; Type III secretion system effector protein, Contributes to Sif formation | Pathogenisis; Type III secretion system | 2.64216 | 0.0022 | -3.5652 | WT unique | 6 | 6 | 19.1 | 19.1 | 37.27 |
| A0A0H3N7I0 | 29.6161 | 29.4072 | 28.9529 | 28.0106 | 28.2545 | 27.656 | dnaJ; DnaJ protein | Chaperones and folding catalysts; Heat shock proteins | 2.17533 | 0 | -1.3517 | Downregulated in OrfSwap | 17 | 17 | 52 | 52 | 41.31 |

### Downregulated in OrfSwap$^{ihfA\text{-}ihfB}$

| Uniprot accession number | LFQ intensity WT_1 | LFQ intensity WT_2 | LFQ intensity WT_3 | LFQ intensity OrfSwap_1 | LFQ intensity OrfSwap_2 | LFQ intensity OrfSwap_3 | Kegg name | Kegg pathway or Kegg Brite | -Log Student's T-test p-value OrfSwap_WT | Student's T-test q-value OrfSwap_WT | Student's T-test Difference OrfSwap_WT | Cluster | Peptides | Unique peptides | Sequence coverage [%] | Unique sequence coverage [%] | Mol. weight [kDa] |
|---|---|---|---|---|---|---|---|---|---|---|---|---|---|---|---|---|---|
| A0A0H3NI84 | 30.4356 | 30.2439 | 30.5114 | 29.2889 | 29.3882 | 29.6524 | hslU; heat shock protein | Chaperones and folding catalysts, Heat Shock Proteins | 2.67979 | 0.0084 | -0.9538 | Downregulated in OrfSwap | 30 | 30 | 52.6 | 52.6 | 49.67 |
| A0A0H3NIA1 | 29.6019 | 28.155 | 29.2111 | 23.7973 | 24.9327 | 25.0807 | sspA; stringent starvation protein A | Genetic information processing; Other transcription-related factors | 2.75028 | 0 | -4.3858 | WT unique | 7 | 7 | 42.5 | 42.5 | 24.25 |
| A0A0H3NK16 | 28.2763 | 28.4899 | 28.302 | 24.2363 | 24.8478 | 25.7316 | cpxR; two-component response regulatory protein | Cationic antimicrobial peptide (CAMP) resistance | 2.83224 | 0.0023 | -3.4175 | WT unique | 6 | 6 | 24.6 | 24.6 | 26.27 |
| A0A0H3NH59 | 29.95 | 29.663 | 30.6645 | 28.1231 | 24.2934 | 25.0869 | uspA; universal stress protein A | Unclassified: signaling and cellular processes | 1.61774 | 0.0047 | -4.258 | WT unique | 4 | 4 | 32.6 | 32.6 | 16.08 |
| A0A0H3NEZ8 | 32.4713 | 32.8971 | 33.0801 | 28.3487 | 27.1865 | 27.8584 | fljB; flagellin | Flagellar assembly | 3.71197 | 0 | -5.0183 | Downregulated in OrfSwap | 35 | 13 | 58.7 | 28.7 | 52.54 |
| A0A0H3NHX2 | 27.9494 | 28.2715 | 28.5072 | 26.7369 | 26.9076 | 26.8092 | motA; motility protein A | Bacterial chemotaxis | 2.96442 | 0 | -1.4248 | Downregulated in OrfSwap | 7 | 7 | 23.9 | 23.9 | 30.07 |
| A0A0H3NDY8 | 27.3737 | 27.6916 | 28.2152 | 26.5006 | 26.568 | 26.155 | cheY; chemotaxis protein CheY | Bacterial chemotaxis | 2.09063 | 0 | -1.3523 | Downregulated in OrfSwap | 3 | 3 | 25.6 | 25.6 | 14.13 |

### Downregulated in OrfSwap$^{ihfA\text{-}ihfB}$

| Uniprot accession number | LFQ intensity WT_1 | LFQ intensity WT_2 | LFQ intensity WT_3 | LFQ intensity OrfSwap_1 | LFQ intensity OrfSwap_2 | LFQ intensity OrfSwap_3 | Kegg name | Kegg pathway or Kegg Brite | -Log Student's T-test p-value OrfSwap_WT | Student's T-test q-value OrfSwap_WT | Student's T-test Difference OrfSwap_WT | Cluster | Peptides | Unique peptides | Sequence coverage [%] | Unique sequence coverage [%] | Mol. weight [kDa] |
|---|---|---|---|---|---|---|---|---|---|---|---|---|---|---|---|---|---|
| A0A0H3NGY0 | 29.5236 | 29.4539 | 30.046 | 28.3766 | 28.2561 | 28.726 | trg; methyl-accepting chemotaxis protein III (mcp-iii) (ribose and galactose chemoreceptor protein) | methyl-accepting chemotaxis protein III, ribose and galactose sensor receptor | 2.1919 | 0.0019 | -1.2216 | Downregulated in OrfSwap | 17 | 17 | 36.2 | 36.2 | 58.3 |
| A0A0H3NBY0 | 30.6015 | 30.544 | 30.4386 | 25.4402 | 24.576 | 25.617 | ymdF; conserved hypothetical protein | Function unknown | 4.08698 | 0 | -5.3169 | WT unique | 7 | 4 | 80 | 40 | 5.744 |
| A0A0H3NMF2 | 28.4841 | 28.5426 | 28.8528 | 25.4479 | 26.306 | 26.057 | cheZ; chemotaxis protein CheZ | Cell motility; bacterial chemotaxis | 3.18639 | 0.0017 | -2.6895 | WT unique | 5 | 5 | 30.8 | 30.8 | 23.92 |
| A0A0H3NCT3 | 28.7151 | 28.4911 | 28.7421 | 24.105 | 24.1709 | 26.1074 | fliB; lysine-N-methylase (ec 2.1.1.-) (lysine N-methyltransferase) | Flagellar assembly proteins | 2.36452 | 0.0021 | -3.855 | WT unique | 15 | 15 | 37.4 | 37.4 | 45.41 |
| A0A0H3NCM2 | 29.3458 | 28.7468 | 28.8794 | 25.7071 | 25.144 | 24.286 | yebF; hypothetical exported protein | Function unknown | 3.02499 | 0 | -3.945 | WT unique | 4 | 4 | 30.8 | 30.8 | 12.81 |
| A0A0H3NP68 | 27.4177 | 27.2361 | 27.986 | 24.8642 | 24.1721 | 25.8293 | yfcZ; conserved hypothetical protein | Function unknown | 2.08834 | 0.0067 | -2.5914 | WT unique | 2 | 2 | 21.3 | 21.3 | 10.29 |
| A0A0H3NIA9 | 29.3405 | 30.2187 | 29.9449 | 28.0922 | 28.5838 | 27.844 | yajQ; conserved hypothetical protein | Function unknown | 2.09715 | 0 | -1.6613 | Downregulated in OrfSwap | 11 | 11 | 64.5 | 64.5 | 19.02 |
| A0A0H3NC84 | 28.9834 | 28.8614 | 28.8789 | 24.7212 | 23.7339 | 25.5601 | conserved hypothetical protein | Function unknown | 2.87937 | 0 | -4.2361 | WT unique | 2 | 2 | 40.5 | 40.5 | 8.818 |

### Downregulated in OrfSwap$^{ihfA\text{-}ihfB}$

| Uniprot accession number | LFQ intensity WT_1 | LFQ intensity WT_2 | LFQ intensity WT_3 | LFQ intensity OrfSwap_1 | LFQ intensity OrfSwap_2 | LFQ intensity OrfSwap_3 | Kegg name | Kegg pathway or Kegg Brite | -Log Student's T-test p-value OrfSwap_WT | Student's T-test q-value OrfSwap_WT | Student's T-test Difference OrfSwap_WT | Cluster | Peptides | Unique peptides | Sequence coverage [%] | Unique sequence coverage [%] | Mol. weight [kDa] |
|---|---|---|---|---|---|---|---|---|---|---|---|---|---|---|---|---|---|
| A0A0H3NEL3 | 28.0153 | 28.1529 | 27.8157 | 25.0612 | 25.6774 | 25.4243 | yeeX; conserved hypothetical protein | Function unknown | 3.66641 | 0.0018 | -2.607 | WT unique | 5 | 5 | 45 | 45 | 13.07 |

### Upregulated in OrfSwap$^{ihfA-ihfB}$

| Uniprot accession number | LFQ intensity WT_1 | LFQ intensity WT_2 | LFQ intensity WT_3 | LFQ intensity OrfSwap_1 | LFQ intensity OrfSwap_2 | LFQ intensity OrfSwap_3 | Kegg name | Kegg pathway or Kegg brite | -Log Student's T-test p-value OrfSwap_WT | Student's T-test q-value OrfSwap_WT | Student's T-test Difference OrfSwap_WT | Cluster | Peptides | Unique peptides | Sequence coverage [%] | Unique sequence coverage [%] | Mol. weight [kDa] |
|---|---|---|---|---|---|---|---|---|---|---|---|---|---|---|---|---|---|
| A0A0H3NG20 | 25.2056 | 25.2049 | 26.1781 | 28.3448 | 28.4186 | 28.4297 | pheS; phenylalanyl-tRNA synthetase alpha chain | Aminoacyl-tRNA biosynthesis | 3.03908 | 0.0017 | 2.86815 | OrfSwap unique | 14 | 14 | 38.5 | 38.5 | 36.754 |
| A0A0H3NGG0 | 30.1529 | 30.0201 | 30.1486 | 31.4566 | 31.1316 | 31.1566 | rps16; 30S ribosomal protein S16. Chloroplast 30S ribosomal protein S16 | Translation | 3.2644 | 0 | 1.1411 | Upregulated in OrfSwap | 8 | 8 | 69.5 | 69.5 | 9.2345 |
| A0A0H3NPN3 | 31.1217 | 31.0677 | 31.2535 | 32.3426 | 32.4188 | 32.6086 | rplI; 50s ribosomal subunit protein L9 | Translation | 3.76832 | 0 | 1.30903 | Upregulated in OrfSwap | 18 | 18 | 81.9 | 81.9 | 15.784 |
| A0A0H3NGC8 | 30.0888 | 29.6599 | 29.5225 | 30.7001 | 30.5954 | 30.8335 | infB; protein chain initiation factor 2 | Genetic information processing; Translation factors | 2.17973 | 0.0093 | 0.9526 | Upregulated in OrfSwap | 39 | 39 | 40.2 | 40.2 | 97.401 |
| A0A0H3NM83 | 26.0313 | 24.7489 | 24.6708 | 28.3626 | 28.1992 | 28.2908 | greA; transcription elongation factor | Genetic information processing; Transcription machinery | 2.67419 | 0.0018 | 3.13388 | OrfSwap unique | 8 | 8 | 57.6 | 57.6 | 17.656 |
| A0A0H3NF40 | 28.4045 | 28.5753 | 28.6486 | 29.9265 | 29.9591 | 29.8785 | recA; RecA protein | Genetic Information Processing; Homologous recombination | 4.26537 | 0 | 1.37856 | Upregulated in OrfSwap | 13 | 13 | 33.4 | 33.4 | 37.944 |
| A0A0H3NU95 | 24.1466 | 24.6449 | 24.9173 | 28.9477 | 29.1212 | 28.9527 | rbsB; D-ribose- | ABC | 4.3509 | 0 | 4.43759 | OrfSwap | 8 | 8 | 38.2 | 38.2 | 30.962 |
| A0A0H3N846 | 25.308 | 25.8357 | 24.0535 | 29.0087 | 29.57 | 29.4279 | yahO; probable secreted protein | General function prediction only | 2.81447 | 0 | 4.26981 | OrfSwap unique | 6 | 6 | 70.3 | 70.3 | 9.9193 |

### Upregulated in OrfSwap$^{ihfA\text{-}ihfB}$

| Uniprot accession number | LFQ intensity WT_1 | LFQ intensity WT_2 | LFQ intensity WT_3 | LFQ intensity OrfSwap_1 | LFQ intensity OrfSwap_2 | LFQ intensity OrfSwap_3 | Kegg name | Kegg pathway or Kegg brite | -Log Student's T-test p-value OrfSwap_WT | Student's T-test q-value OrfSwap_WT | Student's T-test Difference OrfSwap_WT | Cluster | Peptides | Unique peptides | Sequence coverage [%] | Unique sequence coverage [%] | Mol. weight [kDa] |
|---|---|---|---|---|---|---|---|---|---|---|---|---|---|---|---|---|---|
| A0A0H3NWY8 | 24.8129 | 24.2013 | 24.3819 | 27.7246 | 27.4345 | 27.3382 | ydgA; putative transposase | General function prediction only | 3.83124 | 0.0026 | 3.03376 | OrfSwap unique | 3 | 3 | 10.6 | 10.6 | 39.469 |
| A0A0H3NC70 | 25.2653 | 24.6206 | 25.243 | 27.4892 | 27.5189 | 27.1944 | potD; spermidine/putrescine-binding periplasmic protein precursor | ABC transporters | 3.25371 | 0.0046 | 2.35785 | OrfSwap unique | 3 | 3 | 12.1 | 12.1 | 39.021 |
| A0A0H3NJL5 | 24.3533 | 24.4186 | 25.5102 | 27.2967 | 27.3367 | 26.9127 | lolA; outer membrane lipoprotein carrier protein precursor | Unclassified: signaling and cellular processes | 2.4297 | 0.0054 | 2.42131 | OrfSwap unique | 3 | 3 | 27 | 27 | 22.612 |
| A0A0H3NEU6 | 25.3553 | 24.7999 | 24.8673 | 26.9986 | 26.8371 | 26.8071 | artI; arginine-binding periplasmic protein 1 precursor | ABC transporters | 3.27318 | 0.0085 | 1.87347 | OrfSwap unique | 7 | 7 | 25.5 | 25.5 | 27.022 |
| A0A0H3NBQ0 | 30.5868 | 30.5254 | 30.309 | 31.4155 | 31.4943 | 31.8875 | ompD; Outer membrane protein | Transporters; Pores ion channels | 2.58229 | 0.0018 | 1.12537 | Upregulated in OrfSwap | 12 | 10 | 28.5 | 24 | 41.296 |
| A0A0H3NJI9 | 30.7017 | 30.7541 | 30.6564 | 31.4542 | 31.9453 | 32.5101 | ompC; outer membrane protein C | Transporters; Pores ion channels; Environmental Information Processing | 1.8395 | 0.0018 | 1.26579 | Upregulated in OrfSwap | 12 | 12 | 34.7 | 34.7 | 41.238 |
| A0A0H3NLC0 | 32.3121 | 32.679 | 32.2518 | 33.3078 | 33.4153 | 33.326 | osmY; Putative periplasmic protein | Transporters | 2.61189 | 0.0097 | 0.93539 | Upregulated in OrfSwap | 23 | 23 | 80 | 80 | 21.449 |

### Upregulated in OrfSwap$^{ihfA\text{-}ihfB}$

| Uniprot accession number | LFQ intensity WT_1 | LFQ intensity WT_2 | LFQ intensity WT_3 | LFQ intensity OrfSwap_1 | LFQ intensity OrfSwap_2 | LFQ intensity OrfSwap_3 | Kegg name | Kegg pathway or Kegg brite | -Log Student's T-test p-value OrfSwap_WT | Student's T-test q-value OrfSwap_WT | Student's T-test Difference OrfSwap_WT | Cluster | Peptides | Unique peptides | Sequence coverage [%] | Unique sequence coverage [%] | Mol. weight [kDa] |
|---|---|---|---|---|---|---|---|---|---|---|---|---|---|---|---|---|---|
| A0A0H3NE03 | 25.343 | 25.5248 | 25.6053 | 27.2793 | 27.0265 | 27.3546 | zipA; cell division protein | Genetic information processing; Chromosome partitioning proteins | 3.78753 | 0.0084 | 1.7291 | OrfSwap unique | 8 | 8 | 24.1 | 24.1 | 36.334 |
| A0A0H3NH66 | 24.3823 | 25.5681 | 24.8062 | 28.2356 | 28.3379 | 28.0672 | hypothetical phosphosugar-binding protein | General function prediction only | 3.12149 | 0.0025 | 3.29469 | OrfSwap unique | 9 | 9 | 30.5 | 30.5 | 36.275 |
| A0A0H3NAS5 | 25.2665 | 25.4222 | 24.2232 | 27.3577 | 27.4283 | 27.2575 | btuE; hypothetical glutathione peroxidase/vitamin B12 transport periplasmic protein BtuE | Glutathione methabolism | 2.47889 | 0.0053 | 2.37718 | OrfSwap unique | 5 | 5 | 27.3 | 27.3 | 20.423 |
| A0A0H3NAW7 | 24.6386 | 24.9372 | 24.5953 | 27.7094 | 27.4717 | 26.171 | yeaD; conserved hypothetical protein | Glycolysis / Gluconeogenesis | 2.08965 | 0.0087 | 2.39366 | OrfSwap unique | 8 | 8 | 32 | 32 | 32.56 |
| A0A0H3NKM9 | 25.0394 | 24.3908 | 25.3689 | 26.991 | 27.0506 | 26.9852 | conserved hypothetical protein | General function prediction only | 2.70649 | 0.0088 | 2.07589 | OrfSwap unique | 4 | 4 | 16.4 | 16.4 | 32.6 |
| A0A0H3NC50 | 27.8115 | 27.5382 | 28.3911 | 29.4543 | 29.1744 | 29.3421 | fabF; 3-oxoacyl-[acyl-carrier-protein] synthase II | Fatty acid biosynthesis | 2.22593 | 0 | 1.41002 | Upregulated in OrfSwap | 9 | 9 | 31.5 | 31.5 | 42.952 |
| A0A0H3NK91 | 27.7463 | 28.5556 | 28.5165 | 32.1998 | 32.1434 | | acpP; acyl carrier protein | Fatty acid biosynthesis | 3.73518 | 0 | 3.7919 | Upregulated in OrfSwap | 6 | 6 | 44.9 | 44.9 | 8.6394 |

### Upregulated in OrfSwap$^{ihfA-ihfB}$

| Uniprot accession number | LFQ intensity WT_1 | LFQ intensity WT_2 | LFQ intensity WT_3 | LFQ intensity OrfSwap _1 | LFQ intensity OrfSwap _2 | LFQ intensity OrfSwap _3 | Kegg name | Kegg pathway or Kegg brite | -Log Student's T-test p-value OrfSwap _WT | Student's T-test q-value OrfSwap_WT | Student's T-test Difference OrfSwap _WT | Cluster | Peptides | Unique peptides | Sequence coverage [%] | Unique sequence coverage [%] | Mol. weight [kDa] |
|---|---|---|---|---|---|---|---|---|---|---|---|---|---|---|---|---|---|
| A0A0H3NCD1 | 31.539 | 31.9071 | 31.7337 | 33.2541 | 33.4612 | 33.4254 | gapA; glyceraldehyde 3-phosphate dehydrogenase A | Glycolysis / Gluconeogenesis | 3.7373 | 0 | 1.65362 | Upregulated in OrfSwap | 32 | 32 | 72.2 | 72.2 | 35.586 |
| A0A0H3NE21 | 29.2556 | 29.901 | 29.2351 | 31.0944 | 30.7422 | 30.9484 | ahpC; alkyl hydroperoxide reductase c22 protein | Oxidoreductases | 2.42911 | 0 | 1.46445 | Upregulated in OrfSwap | 13 | 13 | 60.4 | 60.4 | 20.747 |
| A0A0H3NER6 | 27.2281 | 27.3842 | 27.2575 | 28.6301 | 28.743 | 28.7964 | dacC; D-alanyl-D-alanine carboxypeptidase (penicillin-binding protein 6 precursor) | Peptidoglycan biosynthesis | 4.51034 | 0 | 1.43325 | Upregulated in OrfSwap | 11 | 10 | 33.2 | 31.5 | 43.664 |
| A0A0H3NAZ4 | 31.0164 | 31.2607 | 30.9925 | 32.0745 | 32.3669 | 32.1952 | katE; catalase HPII | Tryptophan metabolism; Oxidoreductases | 3.12961 | 0.002 | 1.12233 | Upregulated in OrfSwap | 35 | 35 | 38.5 | 38.5 | 83.625 |
| A0A0H3NFA9 | 27.3998 | 27.4101 | 27.2386 | 28.5102 | 28.1303 | 28.6346 | nuoI; NADH dehydrogenase I chain I | Oxidative phosphorylation | 2.57816 | 0.0039 | 1.07551 | Upregulated in OrfSwap | 6 | 6 | 35 | 35 | 20.52 |
| A0A0H3NDI1 | 31.9457 | 32.0802 | 31.8739 | 33.0886 | 32.863 | 32.9742 | tsaA; alkyl hydroperoxide reductase | Oxidoreductases | 3.46395 | 0.004 | 1.00865 | Upregulated in OrfSwap | 18 | 18 | 85 | 85 | 22.317 |
| A0A0H3NQ37 | 29.5465 | 29.4173 | 29.2858 | 30.3373 | 30.293 | 30.3327 | deoB; phosphopentomutase | Pentose phosphate pathway | 3.53176 | 0.0083 | 0.90444 | Upregulated in OrfSwap | 14 | 14 | 34.9 | 34.9 | 44.243 |
| A0A0H3NLM4 | 31.7169 | 31.9022 | 32.0796 | 32.8699 | 32.8041 | 32.8256 | eno; Enolase | Glycolysis / Gluconeogenesis | 3.03011 | 0.0086 | 0.9336 | Upregulated in OrfSwap | 31 | 31 | 53.9 | 53.9 | 45.598 |

### Upregulated in OrfSwap$^{ihfA-ihfB}$

| Uniprot accession number | LFQ intensity WT_1 | LFQ intensity WT_2 | LFQ intensity WT_3 | LFQ intensity OrfSwap _1 | LFQ intensity OrfSwap _2 | LFQ intensity OrfSwap _3 | Kegg name | Kegg pathway or Kegg brite | -Log Student's T-test p-value OrfSwap _WT | Student's T-test q-value OrfSwap_WT | Student's T-test Difference OrfSwap _WT | Cluster | Peptides | Unique peptides | Sequence coverage [%] | Unique sequence coverage [%] | Mol. weight [kDa] |
|---|---|---|---|---|---|---|---|---|---|---|---|---|---|---|---|---|---|
| A0A0H3NEC8 | 30.8924 | 30.8775 | 30.9494 | 31.6659 | 31.7066 | 32.0441 | sucD; succinyl-CoA synthetase alpha chain | Citrate cycle (TCA cycle) | 2.74524 | 0.0092 | 0.89915 | Upregulated in OrfSwap | 15 | 15 | 55.7 | 55.7 | 29.775 |
| E1WAE4 | 25.1487 | 25.4062 | 25.2768 | 28.1918 | 28.2009 | 27.9643 | invH; outer membrane lipoprotein | Pathogenisis; Type III secretion system | 4.91785 | 0 | 2.8418 | OrfSwap unique | 8 | 8 | 61.9 | 61.9 | 16.455 |
| A0A0H3NUM9 | 24.4841 | 24.7401 | 24.4677 | 29.4549 | 29.3737 | 29.2042 | dsbA; thiol:disulfide interchange protein | Cationic antimicrobial peptide (CAMP) resistance | 5.69805 | 0 | 4.78033 | OrfSwap unique | 5 | 5 | 25.1 | 25.1 | 22.911 |
| A0A0H3NSV8 | 24.5662 | 24.8477 | 24.3022 | 29.528 | 29.355 | 29.5609 | fkpA; FKBP-type peptidyl-prolyl isomerase | Chaperones and folding catalysts | 5.06819 | 0 | 4.90925 | OrfSwap unique | 10 | 10 | 46.7 | 46.7 | 28.945 |
| A0A0H3NTP1 | 25.868 | 25.0593 | 25.0986 | 28.8673 | 29.0164 | 28.9986 | secB; protein-export protein SecB | Protein export; Quorum sensing | 3.76292 | 0 | 3.61874 | OrfSwap unique | 4 | 4 | 29 | 29 | 17.245 |
| A0A0H3NHM6 | 28.5469 | 27.5633 | 27.5161 | 30.047 | 29.9764 | 29.4713 | cspC; cold shock-like protein CspC | Genetic information processing | 2.16304 | 0 | 1.95614 | Upregulated in OrfSwap | 5 | 5 | 92.8 | 92.8 | 7.4023 |
| A0A0H3NJ18 | 28.3419 | 28.0642 | 27.8659 | 29.1779 | 29.3549 | 29.284 | hflK; HflK protein | Peptidases and inhibitors | 2.88272 | 0 | 1.1816 | Upregulated in OrfSwap | 19 | 19 | 46.5 | 46.5 | 45.629 |
| A0A0H3NMF8 | 26.7153 | 26.9684 | 29.0557 | 29.7577 | 29.4506 | 29.5774 | cheW; purine binding chemotaxis protein | Bacterial chemotaxis | 1.26613 | 0 | 2.01542 | Upregulated in OrfSwap | 4 | 4 | 19.2 | 19.2 | 18.049 |
| A0A0H3NEJ1 | 25.5714 | 24.7956 | 23.8308 | 29.5425 | 29.4367 | 29.4363 | predicted bacteriophage protein | Function unknown | 3.14465 | 0 | 4.73926 | OrfSwap unique | 14 | 14 | 38.5 | 38.5 | 39.937 |

### Upregulated in OrfSwap$^{ihfA\text{-}ihfB}$

| Uniprot accession number | LFQ intensity WT_1 | LFQ intensity WT_2 | LFQ intensity WT_3 | LFQ intensity OrfSwap _1 | LFQ intensity OrfSwap _2 | LFQ intensity OrfSwap _3 | Kegg name | Kegg pathway or Kegg brite | -Log Student's T-test p-value OrfSwap _WT | Student's T-test q-value OrfSwap_WT | Student's T-test Difference OrfSwap _WT | Cluster | Peptides | Unique peptides | Sequence coverage [%] | Unique sequence coverage [%] | Mol. weight [kDa] |
|---|---|---|---|---|---|---|---|---|---|---|---|---|---|---|---|---|---|
| A0A0H3NE67 | 24.618 | 25.2927 | 24.8986 | 27.9469 | 28.0963 | 28.3956 | ybeL; conserved hypothetical protein | Function unknown | 3.77159 | 0 | 3.20983 | OrfSwap unique | 8 | 8 | 52.2 | 52.2 | 18.41 |
| A0A0H3NLS8 | 25.108 | 24.3935 | 25.5648 | 30.6841 | 31.1864 | 31.0913 | yciF; conserved hypothetical protein | Function unknown | 4.04403 | 0 | 5.96513 | OrfSwap unique | 11 | 11 | 41.9 | 41.9 | 18.652 |
| A0A0H3NF25 | 25.2004 | 25.4046 | 25.2523 | 28.8745 | 29.2612 | 29.1161 | ygaM; conserved hypothetical protein | Function unknown | 5.10974 | 0 | 3.79816 | OrfSwap unique | 7 | 7 | 44.6 | 44.6 | 12.36 |
| A0A0H3NCY7 | 30.7232 | 30.0231 | 31.0721 | 32.3475 | 32.3975 | 32.284 | yaeH; conserved hypothetical protein | Function unknown | 2.30173 | 0 | 1.73687 | Upregulated in OrfSwap | 19 | 19 | 87.5 | 87.5 | 15.094 |
| A0A0H3NF10 | 30.8769 | 30.6399 | 30.364 | 31.8636 | 32.0274 | 31.9489 | ygaU; conserved hypothetical protein | Function unknown | 2.97523 | 0 | 1.31971 | Upregulated in OrfSwap | 17 | 17 | 94 | 94 | 16.121 |
| A0A0H3NEE4 | 26.3683 | 26.8307 | 26.4211 | 28.2491 | 28.7436 | 28.6547 | ybgS; hypothetical exported protein | Function unknown | 3.1679 | 0 | 2.00908 | Upregulated in OrfSwap | 5 | 5 | 21.1 | 21.1 | 13.33 |
| A0A0H3NIT0 | 29.7515 | 29.3757 | 29.4742 | 31.6182 | 31.925 | 31.5568 | yjbJ; conserved hypothetical protein | Function unknown | 3.76288 | 0 | 2.16622 | Upregulated in OrfSwap | 10 | 10 | 78.6 | 78.6 | 8.4594 |
| A0A0H3NAP8 | 27.4251 | 27.4918 | 27.1906 | 28.463 | 28.4791 | 27.9832 | ycfP; conserved hypothetical protein | Function unknown | 2.13582 | 0.0088 | 0.93927 | Upregulated in OrfSwap | 11 | 11 | 62.2 | 62.2 | 21.077 |
| A0A0H3NFS9 | 26.7922 | 25.9608 | 25.6528 | 26.5278 | 27.9579 | 27.7568 | conserved hypothetical protein | Function unknown | 1.07008 | 0.0095 | 1.27892 | Upregulated in OrfSwap | 5 | 5 | 53.7 | 53.7 | 12.094 |

### All proteins

| Uniprot accession number | LFQ intensity WT_1 | LFQ intensity WT_2 | LFQ intensity WT_3 | LFQ intensity OrfSwap_1 | LFQ intensity OrfSwap_2 | LFQ intensity OrfSwap_3 | Pfam name | Uniprot full protein name | Student's T-test Significant OrfSwap_WT | -Log Student's T-test p-value OrfSwap_WT | Student's T-test q-value OrfSwap_WT | Student's T-test Difference OrfSwap_WT | Cluster | Peptides | Unique peptides | Sequence coverage [%] | Unique sequence coverage [%] | Mol. weight [kDa] |
|---|---|---|---|---|---|---|---|---|---|---|---|---|---|---|---|---|---|---|
| A0A0H3NE85 | 29.9222 | 29.4707 | 29.7858 | 28.452 | 28.5802 | 26.6467 | tRNA-synt_1c;tRNA-synt_1c_C | Glutamine--tRNA ligase {ECO:0000256\|HAMAP-Rule:MF_00126} | + | 1.343093457 | 0 | -1.833314896 | Downregulated in OrfSwap | 18 | 18 | 34.2 | 34.2 | 63.54 |
| A0A0H3NJP8 | 30.1782 | 30.793 | 29.9731 | 27.9155 | 29.2027 | 29.1776 | Bac_DNA_binding | Integration host factor subunit beta {ECO:0000256\|HAMAP-Rule:MF_0038 | + | 1.463951498 | 0 | -1.549506505 | Downregulated in OrfSwap | 7 | 7 | 58.5 | 58.5 | 10.62 |
| A0A0H3N9G9 | 29.1035 | 29.0712 | 29.4407 | 28.3664 | 26.8333 | 26.6494 | GST_N_3 | | + | 1.584562813 | 0 | -1.922110875 | Downregulated in OrfSwap | 9 | 9 | 39.4 | 39.4 | 23.69 |
| A0A0H3NIE3 | 30.0971 | 28.3135 | 29.8914 | 27.1019 | 27.2526 | 26.5908 | Ribosomal_L16 | 50S ribosomal protein L16 {ECO:0000256\|HAMAP-Rule:MF_01342, ECO:0000256\|RuleBase:RU004414} | + | 1.828675667 | 0 | -2.452248255 | Downregulated in OrfSwap | 8 | 8 | 49.3 | 49.3 | 15.19 |
| A0A0H3N7T4 | 29.9047 | 31.6913 | 31.6005 | 28.2944 | 32.4188 | 28.5464 | OmpH | | + | 2.003551414 | 0 | -2.731469472 | Downregulated in OrfSwap | 10 | 10 | 57.8 | 57.8 | 17.91 |
| A0A0H3NDY8 | 27.3737 | 27.6916 | 28.2152 | 26.5006 | 26.568 | 26.155 | Response_re | | + | 2.090633807 | 0 | -1.352275848 | Downreg | 3 | 3 | 25.6 | 25.6 | 14.13 |
| A0A0H3NIA9 | 29.3405 | 30.2187 | 29.9449 | 28.0922 | 28.5838 | 27.844 | DUF520 | UPF0234 protein YajQ {ECO:0000256\|HAMAP-Rule:MF_00632} | + | 2.09714984 | 0 | -1.661343257 | Downregulated in OrfSwap | 11 | 11 | 64.5 | 64.5 | 19.02 |
| A0A0H3N7I0 | 29.6161 | 29.4072 | 28.9529 | 28.0106 | 28.2545 | 27.656 | DnaJ;DnaJ_C;DnaJ_CXXCXGXG | Chaperone protein DnaJ {ECO:0000256\|HAMAP-Rule:MF_01152} | + | 2.175334423 | 0 | -1.35169665 | Downregulated in OrfSwap | 17 | 17 | 52 | 52 | 41.31 |

### All proteins

| Uniprot accession number | LFQ intensity WT_1 | LFQ intensity WT_2 | LFQ intensity WT_3 | LFQ intensity OrfSwap_1 | LFQ intensity OrfSwap_2 | LFQ intensity OrfSwap_3 | Pfam name | Uniprot full protein name | Student's T-test Significant OrfSwap_WT | -Log Student's T-test p-value OrfSwap_WT | Student's T-test q-value OrfSwap_WT | Student's T-test Difference OrfSwap_WT | Cluster | Peptides | Unique peptides | Sequence coverage [%] | Unique sequence coverage [%] | Mol. weight [kDa] |
|---|---|---|---|---|---|---|---|---|---|---|---|---|---|---|---|---|---|---|
| A0A0H3NL37 | 30.5684 | 30.3859 | 30.3076 | 28.6955 | 28.5933 | 27.52 | Peptidase_M17;Peptidase_M17_N | Probable cytosol aminopeptidase {ECO:0000256\|HAMAP-Rule:MF_00181} | + | 2.302905915 | 0 | -2.151018143 | Downregulated in OrfSwap | 16 | 16 | 40 | 40 | 54.95 |
| A0A0H3NL58 | 31.575 | 31.0535 | 31.4667 | 30.0823 | 29.8715 | 29.2774 | TPR_3 | | + | 2.30645588 | 0 | -1.621303558 | Downregulated in OrfSwap | 9 | 9 | 57.6 | 57.6 | 19.22 |
| A0A0H3NF99 | 29.6203 | 30.149 | 29.8881 | 28.7604 | 28.6278 | 28.1962 | Invas_SpaK | | + | 2.39382399 | 0 | -1.357708613 | Downregulated in OrfSwap | 4 | 4 | 41.5 | 41.5 | 14.92 |
| A0A0H3NPT7 | 31.5046 | 31.621 | 31.2891 | 29.0566 | 28.9341 | 29.9531 | SHMT | Serine hydroxymethyltransferase {ECO:0000256\|HAMAP-Rule:MF_00051} | + | 2.520955388 | 0 | -2.156953176 | Downregulated in OrfSwap | 31 | 31 | 54.7 | 54.7 | 45.45 |
| A0A0H3NSR3 | 32.4492 | 32.3885 | 32.4912 | 31.3934 | 30.9909 | 30.6419 | Ribosomal_S11 | 30S ribosomal protein S11 {ECO:0000256\|HAMAP-Rule:MF_01310} | + | 2.54993601 | 0 | -1.434219996 | Downregulated in OrfSwap | 12 | 12 | 62.7 | 62.7 | 11.73 |
| A0A0H3NHU8 | 29.7108 | 29.6223 | 29.3445 | 28.4701 | 28.4556 | 27.994 | GAD;tRNA_anti-codon;tRNA-synt_2 | Aspartate--tRNA ligase {ECO:0000256\|HAMAP-Rule:MF_00044} | + | 2.550533202 | 0 | -1.252607346 | Downregulated in OrfSwap | 23 | 23 | 40.8 | 40.8 | 65.67 |

### All proteins

| Uniprot accession number | LFQ intensity WT_1 | LFQ intensity WT_2 | LFQ intensity WT_3 | LFQ intensity OrfSwap_1 | LFQ intensity OrfSwap_2 | LFQ intensity OrfSwap_3 | Pfam name | Uniprot full protein name | Student's T-test Significant OrfSwap_WT | -Log Student's T-test p-value OrfSwap_WT | Student's T-test q-value OrfSwap_WT | Student's T-test Difference OrfSwap_WT | Cluster | Peptides | Unique peptides | Sequence coverage [%] | Unique sequence coverage [%] | Mol. weight [kDa] |
|---|---|---|---|---|---|---|---|---|---|---|---|---|---|---|---|---|---|---|
| A0A0H3NGR4 | 32.7418 | 31.674 | 31.7475 | 28.604 | 28.3166 | 26.9522 | Ribosomal_S17 | 30S ribosomal protein S17 {ECO:0000256\|HAMAP-Rule:MF_01345, ECO:0000256\|RuleBase:RU003873} | + | 2.578855578 | 0 | -4.096844355 | Downregulated in OrfSwap | 9 | 9 | 82.1 | 82.1 | 9.722 |
| A0A0H3NIE8 | 29.9499 | 29.6014 | 30.1481 | 28.5622 | 28.3124 | 28.7206 | PFK | ATP-dependent 6-phosphofructokinase {ECO:0000256\|HAMAP-Rule:MF_00339} | + | 2.628737137 | 0 | -1.368057251 | Downregulated in OrfSwap | 11 | 11 | 34.1 | 34.1 | 34.92 |
| A0A0H3NJ43 | 31.6863 | 32.1787 | 31.9562 | 30.2151 | 30.5471 | 30.5796 | Ribosomal_L33 | 50S ribosomal protein L33 {ECO:0000256\|HAMAP-Rule:MF_00294} | + | 2.902472088 | 0 | -1.493129094 | Downregulated in OrfSwap | 6 | 6 | 76.4 | 76.4 | 6.372 |
| A0A0H3NV54 | 32.0063 | 32.0412 | 32.1672 | 30.6266 | 30.543 | 31.0402 | Ribosomal_L10 | 50S ribosomal protein L10 {ECO:0000256\|HAMAP-Rule:MF_00362, ECO:0000256\|RuleBase:RU003636, ECO:0000256\|SAAS:SAAS01073696} | + | 2.934367538 | 0 | -1.334978104 | Downregulated in OrfSwap | 21 | 21 | 94.5 | 94.5 | 17.8 |
| A0A0H3NHX2 | 27.9494 | 28.2715 | 28.5072 | 26.7369 | 26.9076 | 26.8092 | MotA_ExbB | | + | 2.964422464 | 0 | -1.424840927 | Downregulated in OrfSwap | 7 | 7 | 23.9 | 23.9 | 30.07 |

### All proteins

| Uniprot accession number | LFQ intensity WT_1 | LFQ intensity WT_2 | LFQ intensity WT_3 | LFQ intensity OrfSwap_1 | LFQ intensity OrfSwap_2 | LFQ intensity OrfSwap_3 | Pfam name | Uniprot full protein name | Student's T-test Significant OrfSwap_WT | -Log Student's T-test p-value OrfSwap_WT | Student's T-test q-value OrfSwap_WT | Student's T-test Difference OrfSwap_WT | Cluster | Peptides | Unique peptides | Sequence coverage [%] | Unique sequence coverage [%] | Mol. weight [kDa] |
|---|---|---|---|---|---|---|---|---|---|---|---|---|---|---|---|---|---|---|
| A0A0H3NSS7 | 32.0121 | 31.7291 | 31.9239 | 30.3509 | 30.5665 | 30.7686 | Ribosomal_S14 | 30S ribosomal protein S14 {ECO:0000256\|HAMAP-Rule:MF_00537} | + | 3.080886517 | 0 | -1.326375326 | Downregulated in OrfSwap | 12 | 12 | 74 | 74 | 11.06 |
| A0A0H3NJJ9 | 31.283 | 31.5632 | 31.3099 | 29.8331 | 30.2186 | 29.972 | MFS_1 | | + | 3.178347516 | 0 | -1.377480189 | Downregulated in OrfSwap | 16 | 16 | 22.3 | 22.3 | 50.18 |
| A0A0H3NA20 | 33.5083 | 33.2456 | 33.4173 | 31.482 | 31.5555 | 31.1095 | Bac_DNA_binding | | + | 3.655128384 | 0 | -2.008061091 | Downregulated in OrfSwap | 13 | 13 | 61.1 | 61.1 | 9.24 |
| A0A0H3NEZ8 | 32.4713 | 32.8971 | 33.0801 | 28.3487 | 27.1865 | 27.8584 | Flagellin_C;Flagellin_D3;Flagellin_N | Flagellin {ECO:0000256\|RuleBase:RU362073} | + | 3.711973952 | 0 | -5.018304189 | Downregulated in OrfSwap | 35 | 13 | 58.7 | 28.7 | 52.54 |
| A0A0H3NIE5 | 30.8955 | 30.9135 | 31.0007 | 28.7123 | 29.1455 | 28.6563 | Ribosomal_L23 | 50S ribosomal protein L23 {ECO:0000256\|HAMAP-Rule:MF_01369, ECO:0000256\|RuleBase:RU003935} | + | 3.731757614 | 0 | -2.098535538 | Downregulated in OrfSwap | 9 | 9 | 53 | 53 | 11.21 |
| A0A0H3NMB9 | 30.3836 | 30.1613 | 30.2868 | 27.6154 | 27.7263 | 27.1208 | MreB_Mbl | | + | 3.841132216 | 0 | -2.789749146 | Downregulated in OrfSwap | 17 | 17 | 44.4 | 44.4 | 36.95 |
| A0A0H3NCM8 | 36.1877 | 36.0946 | 36.0741 | 33.8888 | 34.3397 | 34.1492 | LPP | | + | 3.907902788 | 0 | -1.992894491 | Downregulated in OrfSwap | 11 | 11 | 66.7 | 66.7 | 8.391 |
| A0A0H3NIM9 | 30.1236 | 30.0961 | 30.0237 | 28.5506 | 28.2989 | 28.6372 | PAP2 | Acid phosphatase {ECO:0000256\|PIRNR:PIRNR000897} | + | 3.938247353 | 0 | -1.585529327 | Downregulated in OrfSwap | 12 | 12 | 56.4 | 56.4 | 28.38 |
| A0A0H3P177 | 30.7412 | 30.7921 | 30.9653 | 29.7188 | 29.7217 | 29.7917 | TraT | | + | 3.955146529 | 0 | -1.08882459 | Downregulated in OrfSwap | 12 | 12 | 46.1 | 46.1 | 26.17 |

### All proteins

| Uniprot accession number | LFQ intensity WT_1 | LFQ intensity WT_2 | LFQ intensity WT_3 | LFQ intensity OrfSwap_1 | LFQ intensity OrfSwap_2 | LFQ intensity OrfSwap_3 | Pfam name | Uniprot full protein name | Student's T-test Significant OrfSwap_WT | -Log Student's T-test p-value OrfSwap_WT | Student's T-test q-value OrfSwap_WT | Student's T-test Difference OrfSwap_WT | Cluster | Peptides | Unique peptides | Sequence coverage [%] | Unique sequence coverage [%] | Mol. weight [kDa] |
|---|---|---|---|---|---|---|---|---|---|---|---|---|---|---|---|---|---|---|
| A0A0H3NMG1 | 31.9654 | 32.1224 | 31.9136 | 30.861 | 30.8724 | 30.7978 | Ribosomal_L3 | 50S ribosomal protein L3 {ECO:0000256\|HAMAP-Rule:MF_01325, ECO:0000256\|RuleBase:RU003906} | + | 4.182113557 | 0 | -1.156721115 | Downregulated in OrfSwap | 23 | 23 | 62.7 | 62.7 | 22.25 |
| A0A0H3NST4 | 31.7894 | 31.5064 | 31.5212 | 28.8763 | 29.1345 | 28.8631 | Ribosomal_L29 | 50S ribosomal protein L29 {ECO:0000256\|HAMAP-Rule:MF_00374} | + | 4.497632166 | 0 | -2.647713343 | Downregulated in OrfSwap | 5 | 5 | 66.7 | 66.7 | 7.26 |
| A0A0H3NIA0 | 27.6627 | 27.4573 | 27.337 | 26.3432 | 26.2754 | 26.5256 | NanE | Putative N-acetylmannosamine-6-phosphate 2-epimerase {ECO:0000256\|HAMAP-Rule:MF_01235} | + | 3.098065414 | 0.00187755 | -1.104253133 | Downregulated in OrfSwap | 4 | 4 | 16.2 | 16.2 | 24.03 |
| A0A0H3NGY0 | 29.5236 | 29.4539 | 30.046 | 28.3766 | 28.2561 | 28.726 | HAMP;MCPsignal;TarH | | + | 2.191899928 | 0.00191667 | -1.221564611 | Downregulated in OrfSwap | 17 | 17 | 36.2 | 36.2 | 58.3 |
| A0A0H3N9F1 | 27.69 | 27.6129 | 27.6158 | 26.3882 | 26.8304 | 25.9986 | RcsF | Outer membrane lipoprotein RcsF {ECO:0000256\|HAMAP-Rule:MF_00976} | + | 2.158226751 | 0.00195745 | -1.233858744 | Downregulated in OrfSwap | 4 | 4 | 43.3 | 43.3 | 14.19 |

### All proteins

| Uniprot accession number | LFQ intensity WT_1 | LFQ intensity WT_2 | LFQ intensity WT_3 | LFQ intensity OrfSwap _1 | LFQ intensity OrfSwap _2 | LFQ intensity OrfSwap _3 | Pfam name | Uniprot full protein name | Student's T-test Significant OrfSwap_WT | -Log Student's T-test p-value OrfSwap_WT | Student's T-test q-value OrfSwap_WT | Student's T-test Difference OrfSwap_WT | Cluster | Peptides | Unique peptides | Sequence coverage [%] | Unique sequence coverage [%] | Mol. weight [kDa] |
|---|---|---|---|---|---|---|---|---|---|---|---|---|---|---|---|---|---|---|
| A0A0H3N9T3 | 28.8994 | 28.9935 | 28.0829 | 27.311 | 27.4755 | 27.5136 | MukB;MukB_hinge | Chromosome partition protein MukB {ECO:0000256\|HAMAP-Rule:MF_01800, ECO:0000256\|SAAS:SAAS00073551} | + | 1.843494959 | 0.00407547 | -1.225211461 | Downregulated in OrfSwap | 32 | 32 | 20.4 | 20.4 | 170 |
| A0A0H3NIV5 | 31.8072 | 31.42 | 31.5218 | 30.6804 | 30.4625 | 30.3041 | Ribosomal_S18 | 30S ribosomal protein S18 {ECO:0000256\|HAMAP-Rule:MF_00270} | + | 2.639548458 | 0.00415385 | -1.100667318 | Downregulated in OrfSwap | 12 | 12 | 66.7 | 66.7 | 8.986 |
| A0A0H3NEX8 | 30.5707 | 30.4008 | 30.4136 | 29.3314 | 29.4919 | 29.601 | AsnC_trans_reg | | + | 3.306204862 | 0.00549153 | -0.986962636 | Downregulated in OrfSwap | 13 | 13 | 43.3 | 43.3 | 18.86 |
| A0A0H3NC31 | 32.3683 | 32.2929 | 32.2204 | 31.3583 | 31.1325 | 31.4097 | ELFV_dehydrog;ELFV_dehydrog_N | Glutamate dehydrogenase {ECO:0000256\|PIRNR:PIRNR000185} | + | 3.321907404 | 0.00558621 | -0.993742625 | Downregulated in OrfSwap | 33 | 33 | 68 | 68 | 48.04 |
| A0A0H3NGR9 | 31.7872 | 31.1082 | 31.6181 | 30.2059 | 30.7111 | 30.1044 | Ribosomal_S19 | 30S ribosomal protein S19 {ECO:0000256\|HAMAP-Rule:MF_00531} | + | 1.86311149 | 0.00568421 | -1.164014181 | Downregulated in OrfSwap | 10 | 10 | 76.1 | 76.1 | 10.42 |
| A0A0H3NBF9 | 31.181 | 30.985 | 30.9943 | 30.0884 | 30.0778 | 30.0504 | Ribosomal_S22 | | + | 3.954447716 | 0.00578571 | -0.981247584 | Downregulated in OrfSwap | 6 | 6 | 68.1 | 68.1 | 5.369 |
| A0A0H3NGR0 | 31.2695 | 31.4951 | 31.3384 | 29.9184 | 30.058 | 30.8259 | Ribosomal_L18p | 50S ribosomal protein L18 {ECO:0000256\|HAMAP-Rule:MF_01337} | + | 1.717069468 | 0.00688525 | -1.100200653 | Downregulated in OrfSwap | 17 | 17 | 66.7 | 66.7 | 12.77 |

### All proteins

| Uniprot accession number | LFQ intensity WT_1 | LFQ intensity WT_2 | LFQ intensity WT_3 | LFQ intensity OrfSwap_1 | LFQ intensity OrfSwap_2 | LFQ intensity OrfSwap_3 | Pfam name | Uniprot full protein name | Student's T-test Significant OrfSwap_WT | -Log Student's T-test p-value OrfSwap_WT | Student's T-test q-value OrfSwap_WT | Student's T-test Difference OrfSwap_WT | Cluster | Peptides | Unique peptides | Sequence coverage [%] | Unique sequence coverage [%] | Mol. weight [kDa] |
|---|---|---|---|---|---|---|---|---|---|---|---|---|---|---|---|---|---|---|
| A0A0H3NAN8 | 31.902 | 33.0883 | 33.0054 | 30.981 | 31.6268 | 31.6706 | SmpA_OmlA | | + | 1.311313664 | 0.007 | -1.239124934 | Downregulated in OrfSwap | 8 | 8 | 72.6 | 72.6 | 12.16 |
| A0A0H3NI84 | 30.4356 | 30.2439 | 30.5114 | 29.2889 | 29.3882 | 29.6524 | AAA;AAA_2;ClpB_D2-small | ATP-dependent protease ATPase subunit HslU {ECO:0000256\|HAMAP-Rule:MF_00249} | + | 2.67979255 | 0.00843077 | -0.95381546 | Downregulated in OrfSwap | 30 | 30 | 52.6 | 52.6 | 49.67 |
| A0A0H3NID8 | 33.1423 | 33.1852 | 33.1794 | 32.3236 | 32.2048 | 32.3316 | Ribosomal_S5;Ribosomal_S5_C | 30S ribosomal protein S5 {ECO:0000256\|HAMAP-Rule:MF_01307, ECO:0000256\|SAAS:SAAS00382345} | + | 4.471412442 | 0.00869841 | -0.882354736 | Downregulated in OrfSwap | 23 | 23 | 94.6 | 94.6 | 17.6 |
| A0A0H3NII6 | 27.3902 | 27.2823 | 27.188 | 25.7563 | 26.7898 | 26.2175 | SPOR | Cell division protein DamX {ECO:0000256\|HAMAP-Rule:MF_02021} | + | 1.559957557 | 0.00872727 | -1.032308578 | Downregulated in OrfSwap | 9 | 9 | 26.8 | 26.8 | 45.46 |
| A0A0H3NNU6 | 29.0333 | 28.3244 | 28.0774 | 27.2803 | 27.4396 | 27.4219 | MIP | | + | 1.709125629 | 0.00883871 | -1.097738266 | Downregulated in OrfSwap | 3 | 3 | 8.5 | 8.5 | 29.69 |
| A0A0H3NE44 | 30.9651 | 31.1186 | 30.9708 | 30.2448 | 30.1718 | 30.1105 | PALP | Cysteine synthase {ECO:0000256\|RuleBase:RU003985} | + | 3.729357185 | 0.00896 | -0.842487971 | Downregulated in OrfSwap | 19 | 19 | 61.3 | 61.3 | 34.54 |
| A0A0H3N8Z2 | 28.8973 | 28.1287 | 28.6084 | 27.0186 | 28.0261 | 27.3169 | Asn_synthase;GATase_7 | | + | 1.364441509 | 0.00908108 | -1.090922038 | Downregulated in OrfSwap | 14 | 14 | 37.7 | 37.7 | 62.57 |

### All proteins

| Uniprot accession number | LFQ intensity WT_1 | LFQ intensity WT_2 | LFQ intensity WT_3 | LFQ intensity OrfSwap _1 | LFQ intensity OrfSwap _2 | LFQ intensity OrfSwap _3 | Pfam name | Uniprot full protein name | Student's T-test Significant OrfSwap_WT | -Log Student's T-test p-value OrfSwap_WT | Student's T-test q-value OrfSwap_WT | Student's T-test Difference OrfSwap_WT | Cluster | Peptides | Unique peptides | Sequence coverage [%] | Unique sequence coverage [%] | Mol. weight [kDa] |
|---|---|---|---|---|---|---|---|---|---|---|---|---|---|---|---|---|---|---|
| A0A0H3NJ96 | 28.6777 | 28.4375 | 28.6262 | 27.7626 | 27.5856 | 27.7064 | MoCF_biosynth | Molybdenum cofactor biosynthesis protein B {ECO:0000256\|PIRNR:PIRNR006443} | + | 3.246156856 | 0.0096 | -0.895645777 | Downregulated in OrfSwap | 9 | 9 | 53.5 | 53.5 | 18.54 |
| A0A0H3NNN3 | 31.5293 | 31.3719 | 31.3278 | 30.6503 | 30.4286 | 30.4347 | Ribosomal_L25p | 50S ribosomal protein L25 {ECO:0000256\|HAMAP-Rule:MF_01336, ECO:0000256\|SAAS:SAAS01080246} | + | 3.16622692 | 0.00988235 | -0.905138652 | Downregulated in OrfSwap | 8 | 8 | 79.8 | 79.8 | 10.54 |
| A0A0H3N846 | 25.308 | 25.8357 | 24.0535 | 29.0087 | 29.57 | 29.4279 | DUF1471 | | + | 2.814469841 | 0 | 4.269806544 | OrfSwap unique | 6 | 6 | 70.3 | 70.3 | 9.919 |
| A0A0H3NEJ1 | 25.5714 | 24.7956 | 23.8308 | 29.5425 | 29.4367 | 29.4363 | AAA_31 | | + | 3.144651399 | 0 | 4.739256541 | OrfSwap unique | 14 | 14 | 38.5 | 38.5 | 39.94 |
| A0A0H3NTP1 | 25.868 | 25.0593 | 25.0986 | 28.8673 | 29.0164 | 28.9986 | SecB | Protein-export protein SecB {ECO:0000256\|HAMAP-Rule:MF_00821} | + | 3.762922289 | 0 | 3.618738174 | OrfSwap unique | 4 | 4 | 29 | 29 | 17.25 |
| A0A0H3NE67 | 24.618 | 25.2927 | 24.8986 | 27.9469 | 28.0963 | 28.3956 | DUF1451 | | + | 3.771589213 | 0 | 3.209830602 | OrfSwap unique | 8 | 8 | 52.2 | 52.2 | 18.41 |
| A0A0H3NLS8 | 25.108 | 24.3935 | 25.5648 | 30.6841 | 31.1864 | 31.0913 | DUF892 | | + | 4.044030582 | 0 | 5.965133031 | OrfSwap unique | 11 | 11 | 41.9 | 41.9 | 18.65 |
| A0A0H3NU95 | 24.1466 | 24.6449 | 24.9173 | 28.9477 | 29.1212 | 28.9527 | Peripla_BP_4 | | + | 4.35090215 | 0 | 4.437585831 | OrfSwap unique | 8 | 8 | 38.2 | 38.2 | 30.96 |
| E1WAE4 | 25.1487 | 25.4062 | 25.2768 | 28.1918 | 28.2009 | 27.9643 | InvH | Invasion lipoprotein invH | + | 4.917853333 | 0 | 2.841798147 | OrfSwap unique | 8 | 8 | 61.9 | 61.9 | 16.46 |

### All proteins

| Uniprot accession number | LFQ intensity WT_1 | LFQ intensity WT_2 | LFQ intensity WT_3 | LFQ intensity OrfSwap _1 | LFQ intensity OrfSwap _2 | LFQ intensity OrfSwap _3 | Pfam name | Uniprot full protein name | Student's T-test Significant OrfSwap_WT | -Log Student's T-test p-value OrfSwap_WT | Student's T-test q-value OrfSwap_WT | Student's T-test Difference OrfSwap_WT | Cluster | Peptides | Unique peptides | Sequence coverage [%] | Unique sequence coverage [%] | Mol. weight [kDa] |
|---|---|---|---|---|---|---|---|---|---|---|---|---|---|---|---|---|---|---|
| A0A0H3NSV8 | 24.5662 | 24.8477 | 24.3022 | 29.528 | 29.355 | 29.5609 | FKBP_C;FKBP_N | Peptidyl-prolyl cis-trans isomerase {ECO:0000256 \|RuleBase:RU003915} | + | 5.068186758 | 0 | 4.90924708 | OrfSwap unique | 10 | 10 | 46.7 | 46.7 | 28.95 |
| A0A0H3NF25 | 25.2004 | 25.4046 | 25.2523 | 28.8745 | 29.2612 | 29.1161 | DUF883 | | + | 5.109736498 | 0 | 3.798163732 | OrfSwap unique | 7 | 7 | 44.6 | 44.6 | 12.36 |
| A0A0H3NUM9 | 24.4841 | 24.7401 | 24.4677 | 29.4549 | 29.3737 | 29.2042 | DSBA | Thiol:disulfide interchange protein {ECO:0000256 \|PIRNR:PIRNR001488} | + | 5.698047309 | 0 | 4.780326843 | OrfSwap unique | 5 | 5 | 25.1 | 25.1 | 22.91 |
| A0A0H3NG20 | 25.2056 | 25.2049 | 26.1781 | 28.3448 | 28.4186 | 28.4297 | Phe_tRNA-synt_N;tRNA-synt_2d | Phenylalanine--tRNA ligase alpha subunit {ECO:0000256 \|HAMAP-Rule:MF_00281} | + | 3.03908037 | 0.00174194 | 2.868146261 | OrfSwap unique | 14 | 14 | 38.5 | 38.5 | 36.75 |
| A0A0H3NM83 | 26.0313 | 24.7489 | 24.6708 | 28.3626 | 28.1992 | 28.2908 | GreA_GreB;GreA_GreB_N | Transcription elongation factor GreA {ECO:0000256 \|HAMAP-Rule:MF_00105, ECO:0000256\|RuleBase:RU000556, ECO:0000256\|SAAS:SAAS01033802} | + | 2.674189994 | 0.00183051 | 3.133878072 | OrfSwap unique | 8 | 8 | 57.6 | 57.6 | 17.66 |
| A0A0H3NH66 | 24.3823 | 25.5681 | 24.8062 | 28.2356 | 28.3379 | 28.0672 | SIS | | + | 3.121494818 | 0.00245455 | 3.294691086 | OrfSwap unique | 9 | 9 | 30.5 | 30.5 | 36.28 |
| A0A0H3NWY8 | 24.8129 | 24.2013 | 24.3819 | 27.7246 | 27.4345 | 27.3382 | Transposase_31 | | + | 3.831240474 | 0.00263415 | 3.033763885 | OrfSwap unique | 3 | 3 | 10.6 | 10.6 | 39.47 |

### All proteins

| Uniprot accession number | LFQ intensity WT_1 | LFQ intensity WT_2 | LFQ intensity WT_3 | LFQ intensity OrfSwap_1 | LFQ intensity OrfSwap_2 | LFQ intensity OrfSwap_3 | Pfam name | Uniprot full protein name | Student's T-test Significant OrfSwap_WT | -Log Student's T-test p-value OrfSwap_WT | Student's T-test q-value OrfSwap_WT | Student's T-test Difference OrfSwap_WT | Cluster | Peptides | Unique peptides | Sequence coverage [%] | Unique sequence coverage [%] | Mol. weight [kDa] |
|---|---|---|---|---|---|---|---|---|---|---|---|---|---|---|---|---|---|---|
| A0A0H3NC70 | 25.2653 | 24.6206 | 25.243 | 27.4892 | 27.5189 | 27.1944 | SBP_bac_8 | Putrescine-binding periplasmic protein {ECO:0000256\|PIRNR:PIRNR019574} | + | 3.253710513 | 0.00455696 | 2.357851028 | OrfSwap unique | 3 | 3 | 12.1 | 12.1 | 39.02 |
| A0A0H3NAS5 | 25.2665 | 25.4222 | 24.2232 | 27.3577 | 27.4283 | 27.2575 | GSHPx | Thioredoxin/glutathione peroxidase BtuE {ECO:0000256\|HAMAP-Rule:MF_02061} | + | 2.478893215 | 0.00530233 | 2.377180099 | OrfSwap unique | 5 | 5 | 27.3 | 27.3 | 20.42 |
| A0A0H3NJL5 | 24.3533 | 24.4186 | 25.5102 | 27.2967 | 27.3367 | 26.9127 | LolA | Outer-membrane lipoprotein carrier protein {ECO:0000256\|HAMAP-Rule:MF_00240, ECO:0000256\|SAAS:SAAS00846918} | + | 2.429697029 | 0.00542857 | 2.42130661 | OrfSwap unique | 3 | 3 | 27 | 27 | 22.61 |
| A0A0H3NE03 | 25.343 | 25.5248 | 25.6053 | 27.2793 | 27.0265 | 27.3546 | ZipA_C | Cell division protein ZipA {ECO:0000256\|HAMAP-Rule:MF_00509, ECO:0000256\|RuleBase:RU003612, ECO:0000256\|SAAS:SAAS00309506} | + | 3.787532877 | 0.00836364 | 1.729096095 | OrfSwap unique | 8 | 8 | 24.1 | 24.1 | 36.33 |

### All proteins

| Uniprot accession number | LFQ intensity WT_1 | LFQ intensity WT_2 | LFQ intensity WT_3 | LFQ intensity OrfSwap_1 | LFQ intensity OrfSwap_2 | LFQ intensity OrfSwap_3 | Pfam name | Uniprot full protein name | Student's T-test Significant OrfSwap_WT | -Log Student's T-test p-value OrfSwap_WT | Student's T-test q-value OrfSwap_WT | Student's T-test Difference OrfSwap_WT | Cluster | Peptides | Unique peptides | Sequence coverage [%] | Unique sequence coverage [%] | Mol. weight [kDa] |
|---|---|---|---|---|---|---|---|---|---|---|---|---|---|---|---|---|---|---|
| A0A0H3NEU6 | 25.3553 | 24.7999 | 24.8673 | 26.9986 | 26.8371 | 26.8071 | SBP_bac_3 | | + | 3.273182969 | 0.00853608 | 1.87346522 | OrfSwap unique | 7 | 7 | 25.5 | 25.5 | 27.02 |
| A0A0H3NAW7 | 24.6386 | 24.9372 | 24.5953 | 27.7094 | 27.4717 | 26.171 | Aldose_epim | Putative glucose-6-phosphate 1-epimerase {ECO:0000256\|PIRNR:PIRNR016020} | + | 2.089646539 | 0.00871579 | 2.393657049 | OrfSwap unique | 8 | 8 | 32 | 32 | 32.56 |
| A0A0H3NKM9 | 25.0394 | 24.3908 | 25.3689 | 26.991 | 27.0506 | 26.9852 | Fructosamin_kin | | + | 2.706491711 | 0.00880851 | 2.075890223 | OrfSwap unique | 4 | 4 | 16.4 | 16.4 | 32.6 |
| A0A0H3NIV6 | 25.663 | 26.3983 | 24.5004 | 27.7438 | 28.0012 | 27.8798 | AsmA | | + | 1.87158564 | 0.00952 | 2.354379018 | OrfSwap unique | 7 | 7 | 11.4 | 11.4 | 74.72 |
| A0A0H3NMF8 | 26.7153 | 26.9684 | 29.0557 | 29.7577 | 29.4506 | 29.5774 | CheW | | + | 1.266134466 | 0 | 2.015424093 | Upregulated in OrfSwap | 4 | 4 | 19.2 | 19.2 | 18.05 |
| A0A0H3NHM6 | 28.5469 | 27.5633 | 27.5161 | 30.047 | 29.9764 | 29.4713 | CSD | | + | 2.163040941 | 0 | 1.956142426 | Upregulated in OrfSwap | 5 | 5 | 92.8 | 92.8 | 7.402 |
| A0A0H3NC50 | 27.8115 | 27.5382 | 28.3911 | 29.4543 | 29.1744 | 29.3421 | ketoacyl-synt;Ketoacyl-synt_C | 3-oxoacyl-[acyl-carrier-protein] synthase 2 {ECO:0000256\|PIRNR:PIRNR000447} | + | 2.225925156 | 0 | 1.410021464 | Upregulated in OrfSwap | 9 | 9 | 31.5 | 31.5 | 42.95 |
| A0A0H3NCY7 | 30.7232 | 30.0231 | 31.0721 | 32.3475 | 32.3975 | 32.284 | DUF3461 | UPF0325 protein YaeH {ECO:0000256\|HAMAP-Rule:MF_01519} | + | 2.30172554 | 0 | 1.73686854 | Upregulated in OrfSwap | 19 | 19 | 87.5 | 87.5 | 15.09 |
| A0A0H3NE21 | 29.2556 | 29.901 | 29.2351 | 31.0944 | 30.7422 | 30.9484 | 1-cysPrx_C;AhpC-TSA | | + | 2.429109252 | 0 | 1.464451472 | Upregulated in OrfSwap | 13 | 13 | 60.4 | 60.4 | 20.75 |
| A0A0H3NJ18 | 28.3419 | 28.0642 | 27.8659 | 29.1779 | 29.3549 | 29.284 | Band_7;HflK_N | Protein HflK {ECO:0000256\|RuleBase:RU364113} | + | 2.882721164 | 0 | 1.181601842 | Upregulated in OrfSwap | 19 | 19 | 46.5 | 46.5 | 45.63 |

#### All proteins

| Uniprot accession number | LFQ intensity WT_1 | LFQ intensity WT_2 | LFQ intensity WT_3 | LFQ intensity OrfSwap_1 | LFQ intensity OrfSwap_2 | LFQ intensity OrfSwap_3 | Pfam name | Uniprot full protein name | Student's T-test Significant OrfSwap_WT | -Log Student's T-test p-value OrfSwap_WT | Student's T-test q-value OrfSwap_WT | Student's T-test Difference OrfSwap_WT | Cluster | Peptides | Unique peptides | Sequence coverage [%] | Unique sequence coverage [%] | Mol. weight [kDa] |
|---|---|---|---|---|---|---|---|---|---|---|---|---|---|---|---|---|---|---|
| A0A0H3NF10 | 30.8769 | 30.6399 | 30.364 | 31.8636 | 32.0274 | 31.9489 | BON;LysM | | + | 2.975232309 | 0 | 1.319713593 | Upregulated in OrfSwap | 17 | 17 | 94 | 94 | 16.12 |
| A0A0H3NEE4 | 26.3683 | 26.8307 | 26.4211 | 28.2491 | 28.7436 | 28.6547 | YbgS | | + | 3.167895985 | 0 | 2.009081523 | Upregulated in OrfSwap | 5 | 5 | 21.1 | 21.1 | 13.33 |
| A0A0H3NGG0 | 30.1529 | 30.0201 | 30.1486 | 31.4566 | 31.1316 | 31.1566 | Ribosomal_S16 | 30S ribosomal protein S16 {ECO:0000256\|HAMAP-Rule:MF_00385} | + | 3.264399837 | 0 | 1.141096115 | Upregulated in OrfSwap | 8 | 8 | 69.5 | 69.5 | 9.235 |
| A0A0H3NK91 | 27.7463 | 28.5556 | 28.5165 | 32.1998 | 32.1434 | 31.8508 | PP-binding | Acyl carrier protein {ECO:0000256\|HAMAP-Rule:MF_01217, ECO:0000256\|RuleBase:RU003545} | + | 3.735178125 | 0 | 3.79189682 | Upregulated in OrfSwap | 6 | 6 | 44.9 | 44.9 | 8.639 |
| A0A0H3NCD1 | 31.539 | 31.9071 | 31.7337 | 33.2541 | 33.4612 | 33.4254 | Gp_dh_C;Gp_dh_N | Glyceraldehyde-3-phosphate dehydrogenase {ECO:0000256\|RuleBase:RU361160} | + | 3.737301543 | 0 | 1.653623581 | Upregulated in OrfSwap | 32 | 32 | 72.2 | 72.2 | 35.59 |
| A0A0H3NIT0 | 29.7515 | 29.3757 | 29.4742 | 31.6182 | 31.925 | 31.5568 | CsbD | | + | 3.762880496 | 0 | 2.16621844 | Upregulated in OrfSwap | 10 | 10 | 78.6 | 78.6 | 8.459 |
| A0A0H3NPN3 | 31.1217 | 31.0677 | 31.2535 | 32.3426 | 32.4188 | 32.6086 | Ribosomal_L9_C;Ribosomal_L9_N | 50S ribosomal protein L9 {ECO:0000256\|HAMAP-Rule:MF_00503} | + | 3.768319647 | 0 | 1.309034348 | Upregulated in OrfSwap | 18 | 18 | 81.9 | 81.9 | 15.78 |

### All proteins

| Uniprot accession number | LFQ intensity WT_1 | LFQ intensity WT_2 | LFQ intensity WT_3 | LFQ intensity OrfSwap _1 | LFQ intensity OrfSwap _2 | LFQ intensity OrfSwap _3 | Pfam name | Uniprot full protein name | Student's T-test Significant OrfSwap_WT | -Log Student's T-test p-value OrfSwap_WT | Student's T-test q-value OrfSwap_WT | Student's T-test Difference OrfSwap_WT | Cluster | Peptides | Unique peptides | Sequence coverage [%] | Unique sequence coverage [%] | Mol. weight [kDa] |
|---|---|---|---|---|---|---|---|---|---|---|---|---|---|---|---|---|---|---|
| A0A0H3NF40 | 28.4045 | 28.5753 | 28.6486 | 29.9265 | 29.9591 | 29.8785 | RecA | Protein RecA {ECO:0000256\|HAMAP-Rule:MF_00268, ECO:0000256\|RuleBase:RU000526, ECO:0000256\|SAAS:SAAS00013832} | + | 4.265369542 | 0 | 1.378559113 | Upregulated in OrfSwap | 13 | 13 | 33.4 | 33.4 | 37.94 |
| A0A0H3NER6 | 27.2281 | 27.3842 | 27.2575 | 28.6301 | 28.743 | 28.7964 | PBP5_C;Peptidase_S11 | | + | 4.510338045 | 0 | 1.43325297 | Upregulated in OrfSwap | 11 | 10 | 33.2 | 31.5 | 43.66 |
| A0A0H3NBQ0 | 30.5868 | 30.5254 | 30.309 | 31.4155 | 31.4943 | 31.8875 | Porin_1 | | + | 2.582288253 | 0.00180392 | 1.125371297 | Upregulated in OrfSwap | 12 | 10 | 28.5 | 24 | 41.3 |
| A0A0H3NJI9 | 30.7017 | 30.7541 | 30.6564 | 31.4542 | 31.9453 | 32.5101 | Porin_1 | | + | 1.839498289 | 0.00184 | 1.265792211 | Upregulated in OrfSwap | 12 | 12 | 34.7 | 34.7 | 41.24 |
| A0A0H3NAZ4 | 31.0164 | 31.2607 | 30.9925 | 32.0745 | 32.3669 | 32.1952 | Catalase;Catalase-rel | Catalase {ECO:0000256\|PIRNR:PIRNR038927, ECO:0000256\|RuleBase:RU000498} | + | 3.129606045 | 0.002 | 1.122330348 | Upregulated in OrfSwap | 35 | 35 | 38.5 | 38.5 | 83.63 |
| A0A0H3NFA9 | 27.3998 | 27.4101 | 27.2386 | 28.5102 | 28.1303 | 28.6346 | Fer4_7 | NADH-quinone oxidoreductase subunit I {ECO:0000256\|HAMAP-Rule:MF_01351} | + | 2.578158117 | 0.00392727 | 1.075511932 | Upregulated in OrfSwap | 6 | 6 | 35 | 35 | 20.52 |
| A0A0H3NDI1 | 31.9457 | 32.0802 | 31.8739 | 33.0886 | 32.863 | 32.9742 | 1-cysPrx_C;AhpC-TSA | | + | 3.463949199 | 0.004 | 1.008647919 | Upregulated in OrfSwap | 18 | 18 | 85 | 85 | 22.32 |

### All proteins

| Uniprot accession number | LFQ intensity WT_1 | LFQ intensity WT_2 | LFQ intensity WT_3 | LFQ intensity OrfSwap_1 | LFQ intensity OrfSwap_2 | LFQ intensity OrfSwap_3 | Pfam name | Uniprot full protein name | Student's T-test Significant OrfSwap_WT | -Log Student's T-test p-value OrfSwap_WT | Student's T-test q-value OrfSwap_WT | Student's T-test Difference OrfSwap_WT | Cluster | Peptides | Unique peptides | Sequence coverage [%] | Unique sequence coverage [%] | Mol. weight [kDa] |
|---|---|---|---|---|---|---|---|---|---|---|---|---|---|---|---|---|---|---|
| A0A0H3NQ37 | 29.5465 | 29.4173 | 29.2858 | 30.3373 | 30.293 | 30.3327 | Metalloenzyme | Phosphopentomutase {ECO:0000256\|HAMAP-Rule:MF_00740, ECO:0000256\|SAAS:SAAS00847420} | + | 3.531763124 | 0.00830303 | 0.904441198 | Upregulated in OrfSwap | 14 | 14 | 34.9 | 34.9 | 44.24 |
| A0A0H3NLM4 | 31.7169 | 31.9022 | 32.0796 | 32.8699 | 32.8041 | 32.8256 | Enolase_C;Enolase_N | Enolase {ECO:0000256\|HAMAP-Rule:MF_00318} | + | 3.030113139 | 0.0085625 | 0.9335982 | Upregulated in OrfSwap | 31 | 31 | 53.9 | 53.9 | 45.6 |
| A0A0H3NAP8 | 27.4251 | 27.4918 | 27.1906 | 28.463 | 28.4791 | 27.9832 | UPF0227 | UPF0227 protein YcfP {ECO:0000256\|HAMAP-Rule:MF_01047} | + | 2.135815079 | 0.00884211 | 0.939271927 | Upregulated in OrfSwap | 11 | 11 | 62.2 | 62.2 | 21.08 |
| A0A0H3NEC8 | 30.8924 | 30.8775 | 30.9494 | 31.6659 | 31.7066 | 32.0441 | CoA_binding;Ligase_CoA | Succinate--CoA ligase [ADP-forming] subunit alpha {ECO:0000256\|HAMAP-Rule:MF_01988} | + | 2.745241936 | 0.00920548 | 0.899145126 | Upregulated in OrfSwap | 15 | 15 | 55.7 | 55.7 | 29.78 |

### All proteins

| Uniprot accession number | LFQ intensity WT_1 | LFQ intensity WT_2 | LFQ intensity WT_3 | LFQ intensity OrfSwap_1 | LFQ intensity OrfSwap_2 | LFQ intensity OrfSwap_3 | Pfam name | Uniprot full protein name | Student's T-test Significant OrfSwap_WT | -Log Student's T-test p-value OrfSwap_WT | Student's T-test q-value OrfSwap_WT | Student's T-test Difference OrfSwap_WT | Cluster | Peptides | Unique peptides | Sequence coverage [%] | Unique sequence coverage [%] | Mol. weight [kDa] |
|---|---|---|---|---|---|---|---|---|---|---|---|---|---|---|---|---|---|---|
| A0A0H3NGC8 | 30.0888 | 29.6599 | 29.5225 | 30.7001 | 30.5954 | 30.8335 | GTP_EFTU;IF-2;IF2_assoc;IF2_N | Translation initiation factor IF-2 {ECO:0000256\|HAMAP-Rule:MF_00100, ECO:0000256\|RuleBase:RU000644, ECO:0000256\|SAAS:SAAS00048520} | + | 2.179727241 | 0.00933333 | 0.952603658 | Upregulated in OrfSwap | 39 | 39 | 40.2 | 40.2 | 97.4 |
| A0A0H3NFS9 | 26.7922 | 25.9608 | 25.6528 | 26.5278 | 27.9579 | 27.7568 | DUF326 | | + | 1.070079388 | 0.00946479 | 1.278924306 | Upregulated in OrfSwap | 5 | 5 | 53.7 | 53.7 | 12.09 |
| A0A0H3NLC0 | 32.3121 | 32.679 | 32.2518 | 33.3078 | 33.4153 | 33.326 | BON | | + | 2.611889253 | 0.00973913 | 0.935385386 | Upregulated in OrfSwap | 23 | 23 | 80 | 80 | 21.45 |
| A0A0H3NDG3 | 28.6884 | 28.7489 | 28.5633 | 24.1678 | 23.9782 | 25.7234 | Topo_Zn_Ribbon;Topoisom_bac;Toprim;zf-C4_Topoisom | DNA topoisomerase 1 {ECO:0000256\|HAMAP-Rule:MF_00952} | + | 2.722839775 | 0 | -4.04370753 | WT unique | 21 | 21 | 29.9 | 29.9 | 97.3 |
| A0A0H3NIA1 | 29.6019 | 28.155 | 29.2111 | 23.7973 | 24.9327 | 25.0807 | GST_C;GST_N | | + | 2.750280955 | 0 | -4.38580958 | WT unique | 7 | 7 | 42.5 | 42.5 | 24.25 |
| A0A0H3NQU8 | 29.0522 | 29.275 | 29.3292 | 23.7036 | 25.7339 | 24.6615 | LuxS | S-ribosylhomocysteine lyase {ECO:0000256\|HAMAP-Rule:MF_00091, ECO:0000256\|SAAS:SAAS00962992} | + | 2.79951216 | 0 | -4.519123077 | WT unique | 12 | 12 | 51.5 | 51.5 | 19.31 |
| A0A0H3NC84 | 28.9834 | 28.8614 | 28.8789 | 24.7212 | 23.7339 | 25.5601 | DUF1480 | | + | 2.879372119 | 0 | -4.23614947 | WT unique | 2 | 2 | 40.5 | 40.5 | 8.818 |

### All proteins

| Uniprot accession number | LFQ intensity WT_1 | LFQ intensity WT_2 | LFQ intensity WT_3 | LFQ intensity OrfSwap_1 | LFQ intensity OrfSwap_2 | LFQ intensity OrfSwap_3 | Pfam name | Uniprot full protein name | Student's T-test Significant OrfSwap_WT | -Log Student's T-test p-value OrfSwap_WT | Student's T-test q-value OrfSwap_WT | Student's T-test Difference OrfSwap_WT | Cluster | Peptides | Unique peptides | Sequence coverage [%] | Unique sequence coverage [%] | Mol. weight [kDa] |
|---|---|---|---|---|---|---|---|---|---|---|---|---|---|---|---|---|---|---|
| A0A0H3NGD6 | 29.358 | 29.2159 | 29.0969 | 25.8424 | 26.1382 | 24.759 | Ribosomal_S30AE | | + | 2.989062073 | 0 | -3.643734614 | WT unique | 7 | 7 | 57.1 | 57.1 | 12.65 |
| A0A0H3NCM2 | 29.3458 | 28.7468 | 28.8794 | 25.7071 | 25.144 | 24.286 | YebF | Protein YebF {ECO:0000256\|HAMAP-Rule:MF_01435} | + | 3.024988175 | 0 | -3.944953283 | WT unique | 4 | 4 | 30.8 | 30.8 | 12.81 |
| A0A0H3NUL5 | 28.7276 | 28.6842 | 28.7611 | 25.5831 | 24.6112 | 24.0706 | Thiolase_C;Thiolase_N | 3-ketoacyl-CoA thiolase {ECO:0000256\|HAMAP-Rule:MF_01620} | + | 3.066086255 | 0 | -3.969276428 | WT unique | 10 | 10 | 24 | 24 | 41 |
| A0A0H3NE87 | 29.2233 | 28.8877 | 29.059 | 25.8753 | 25.633 | 24.6033 | DHDPS | 4-hydroxy-tetrahydrodipicolinate synthase {ECO:0000256\|HAMAP-Rule:MF_00418} | + | 3.105653931 | 0 | -3.686118444 | WT unique | 11 | 11 | 50.7 | 50.7 | 31.29 |
| A0A0H3NIJ8 | 28.3055 | 27.5692 | 28.1214 | 23.8103 | 24.296 | 24.8487 | GyrI-like | DNA gyrase inhibitor {ECO:0000256\|HAMAP-Rule:MF_01896} | + | 3.229027489 | 0 | -3.680379232 | WT unique | 6 | 6 | 33.5 | 33.5 | 18.06 |
| A0A0H3N866 | 28.7187 | 28.8576 | 28.5595 | 25.0252 | 24.8678 | 23.9194 | F420_oxidored;P5CR_dimer | Pyrroline-5-carboxylate reductase {ECO:0000256\|HAMAP-Rule:MF_01925, ECO:0000256\|RuleBase:RU003903} | + | 3.492224845 | 0 | -4.107822418 | WT unique | 9 | 9 | 28.3 | 28.3 | 28.06 |

### All proteins

| Uniprot accession number | LFQ intensity WT_1 | LFQ intensity WT_2 | LFQ intensity WT_3 | LFQ intensity OrfSwap_1 | LFQ intensity OrfSwap_2 | LFQ intensity OrfSwap_3 | Pfam name | Uniprot full protein name | Student's T-test Significant OrfSwap_WT | -Log Student's T-test p-value OrfSwap_WT | Student's T-test q-value OrfSwap_WT | Student's T-test Difference OrfSwap_WT | Cluster | Peptides | Unique peptides | Sequence coverage [%] | Unique sequence coverage [%] | Mol. weight [kDa] |
|---|---|---|---|---|---|---|---|---|---|---|---|---|---|---|---|---|---|---|
| A0A0H3NVG2 | 28.6116 | 28.2947 | 28.6507 | 24.9511 | 24.0273 | 24.9687 | SSB | Single-stranded DNA-binding protein {ECO:0000256\|HAMAP-Rule:MF_00984, ECO:0000256\|RuleBase:RU000524} | + | 3.515759464 | 0 | -3.869991302 | WT unique | 7 | 7 | 28.4 | 28.4 | 19.07 |
| A0A0H3NV13 | 28.3698 | 28.3983 | 28.1877 | 24.6317 | 25.4968 | 25.175 | RraA-like | Regulator of ribonuclease activity A {ECO:0000256\|HAMAP-Rule:MF_00471, ECO:0000256\|SAAS:SAAS00850171} | + | 3.604549246 | 0 | -3.217415492 | WT unique | 7 | 7 | 42.2 | 42.2 | 17.37 |
| A0A0H3N9A8 | 28.3787 | 28.2061 | 28.2145 | 24.8662 | 25.5054 | 24.8415 | Fe-S_biosyn | Iron-sulfur cluster insertion protein ErpA {ECO:0000256\|HAMAP-Rule:MF_01380} | + | 3.849718581 | 0 | -3.195384979 | WT unique | 4 | 4 | 36.8 | 36.8 | 12.1 |
| A0A0H3NJY4 | 28.5546 | 27.9497 | 28.3016 | 24.6025 | 24.6516 | 25.1013 | CoA_binding_2 | | + | 3.907500457 | 0 | -3.483498255 | WT unique | 5 | 5 | 59.4 | 59.4 | 14.91 |
| A0A0H3NNJ9 | 28.3533 | 28.0967 | 28.0895 | 24.9876 | 24.6338 | 24.2107 | Esterase | S-formylglutathione hydrolase {ECO:0000256\|RuleBase:RU363068} | + | 3.91878525 | 0 | -3.56914266 | WT unique | 6 | 6 | 16.8 | 16.8 | 31.97 |

### All proteins

| Uniprot accession number | LFQ intensity WT_1 | LFQ intensity WT_2 | LFQ intensity WT_3 | LFQ intensity OrfSwap_1 | LFQ intensity OrfSwap_2 | LFQ intensity OrfSwap_3 | Pfam name | Uniprot full protein name | Student's T-test Significant OrfSwap_WT | -Log Student's T-test p-value OrfSwap_WT | Student's T-test q-value OrfSwap_WT | Student's T-test Difference OrfSwap_WT | Cluster | Peptides | Unique peptides | Sequence coverage [%] | Unique sequence coverage [%] | Mol. weight [kDa] |
|---|---|---|---|---|---|---|---|---|---|---|---|---|---|---|---|---|---|---|
| A0A0H3NGQ8 | 30.3574 | 30.8635 | 30.7354 | 25.6563 | 25.6792 | 24.6995 | Ribosomal_S8 | 30S ribosomal protein S8 {ECO:0000256\|HAMAP-Rule:MF_01302} | + | 3.92465837 | 0 | -5.307102203 | WT unique | 10 | 10 | 65.4 | 65.4 | 14.13 |
| A0A0H3NP42 | 27.7515 | 27.821 | 27.7342 | 24.8067 | 24.609 | 24.1086 | 2-Hacid_dh;2-Hacid_dh_C;DUF3410 | Erythronate-4-phosphate dehydrogenase {ECO:0000256\|HAMAP-Rule:MF_01825, ECO:0000256\|SAAS:SAAS01082383} | + | 4.002882954 | 0 | -3.260825475 | WT unique | 12 | 12 | 36 | 36 | 41.3 |
| A0A0H3NFA7 | 28.7392 | 28.7823 | 28.4217 | 25.2814 | 24.5944 | 25.0508 | RMF | Ribosome modulation factor {ECO:0000256\|HAMAP-Rule:MF_00919} | + | 4.033545435 | 0 | -3.672176997 | WT unique | 5 | 5 | 72.7 | 72.7 | 6.572 |
| A0A0H3NF82 | 29.3335 | 29.2609 | 28.8951 | 25.5643 | 24.8829 | 25.2882 | T3SS_needle_F | | + | 4.084497533 | 0 | -3.918017705 | WT unique | 6 | 6 | 76.2 | 76.2 | 8.857 |
| A0A0H3NBY0 | 30.6015 | 30.544 | 30.4386 | 25.4402 | 24.576 | 25.617 | KGG | | + | 4.086975817 | 0 | -5.316932042 | WT unique | 7 | 4 | 80 | 40 | 5.744 |

### All proteins

| Uniprot accession number | LFQ intensity WT_1 | LFQ intensity WT_2 | LFQ intensity WT_3 | LFQ intensity OrfSwap _1 | LFQ intensity OrfSwap _2 | LFQ intensity OrfSwap _3 | Pfam name | Uniprot full protein name | Student's T-test Significant OrfSwap_WT | -Log Student's T-test p-value OrfSwap_WT | Student's T-test q-value OrfSwap_WT | Student's T-test Difference OrfSwap_WT | Cluster | Peptides | Unique peptides | Sequence coverage [%] | Unique sequence coverage [%] | Mol. weight [kDa] |
|---|---|---|---|---|---|---|---|---|---|---|---|---|---|---|---|---|---|---|
| A0A0H3N894 | 28.706 | 28.594 | 28.3689 | 24.9273 | 24.2324 | 24.8206 | ACCA | Acetyl-coenzyme A carboxylase carboxyl transferase subunit alpha {ECO:0000256\|HAMAP-Rule:MF_00823, ECO:0000256\|SAAS:SAAS00403330} | + | 4.09109926 | 0 | -3.896204631 | WT unique | 11 | 11 | 33.5 | 33.5 | 35.34 |
| A0A0H3N7F3 | 28.9772 | 28.8858 | 29.0756 | 25.9895 | 26.1626 | 25.6027 | CPSase_L_D2;CPSase_L_D3;MGS | Carbamoyl-phosphate synthase large chain {ECO:0000256\|HAMAP-Rule:MF_01210} | + | 4.209077058 | 0 | -3.061215719 | WT unique | 27 | 27 | 30 | 30 | 118.1 |
| A0A0H3ND10 | 28.026 | 28.0589 | 27.9616 | 25.0731 | 24.8611 | 25.3712 | Acetyltransf_3 | | + | 4.375028891 | 0 | -2.913706462 | WT unique | 4 | 4 | 31.7 | 31.7 | 22.05 |
| A0A0H3NHP0 | 28.5757 | 28.605 | 28.4982 | 24.9141 | 24.6981 | 25.3174 | DNA_gyraseB;DNA_gyraseB_C;HATPase_c;Toprim | DNA gyrase subunit B {ECO:0000256\|HAMAP-Rule:MF_01898} | + | 4.385015726 | 0 | -3.58312734 | WT unique | 19 | 19 | 18.5 | 18.5 | 89.9 |
| A0A0H3NF89 | 27.8378 | 28.1916 | 28.2758 | 25.0186 | 25.2627 | 25.0742 | Trans_reg_C | | + | 4.387283653 | 0 | -2.983248393 | WT unique | 15 | 15 | 27.5 | 27.5 | 63.04 |

### All proteins

| Uniprot accession number | LFQ intensity WT_1 | LFQ intensity WT_2 | LFQ intensity WT_3 | LFQ intensity OrfSwap_1 | LFQ intensity OrfSwap_2 | LFQ intensity OrfSwap_3 | Pfam name | Uniprot full protein name | Student's T-test Significant OrfSwap_WT | -Log Student's T-test p-value OrfSwap_WT | Student's T-test q-value OrfSwap_WT | Student's T-test Difference OrfSwap_WT | Cluster | Peptides | Unique peptides | Sequence coverage [%] | Unique sequence coverage [%] | Mol. weight [kDa] |
|---|---|---|---|---|---|---|---|---|---|---|---|---|---|---|---|---|---|---|
| A0A0H3NE99 | 28.5288 | 28.3718 | 28.2848 | 23.9503 | 24.5388 | 24.2508 | CBS;IMPDH | Inosine-5'-monophosphate dehydrogenase {ECO:0000256\|HAMAP-Rule:MF_01964, ECO:0000256\|RuleBase:RU003928} | + | 4.636932613 | 0 | -4.148470561 | WT unique | 6 | 6 | 16.8 | 16.8 | 51.95 |
| A0A0H3NK15 | 28.1404 | 27.9784 | 28.0586 | 24.8839 | 24.6116 | 24.3896 | SSF | | + | 4.659973067 | 0 | -3.430778503 | WT unique | 5 | 5 | 12.4 | 12.4 | 54.32 |
| A0A0H3NAA4 | 29.0752 | 29.0026 | 28.8242 | 25.1261 | 24.8356 | 24.5857 | PTS_EIIB;PTS_EIIC | | + | 4.733206178 | 0 | -4.118205388 | WT unique | 12 | 12 | 17.8 | 17.8 | 50.5 |
| A0A0H3NHZ9 | 28.2363 | 28.1298 | 28.2169 | 24.7374 | 24.8031 | 25.0236 | 5_3_exonuc;5_3_exonuc_N;DNA_pol_A;DNA_pol_A_exo1 | DNA polymerase I {ECO:0000256\|RuleBase:RU004460} | + | 5.453809893 | 0 | -3.339597702 | WT unique | 11 | 11 | 14.7 | 14.7 | 103.1 |
| A0A0H3NKD9 | 28.874 | 28.8394 | 28.8375 | 25.0415 | 25.2096 | 25.2921 | GcpE | 4-hydroxy-3-methylbut-2-en-1-yl diphosphate synthase (flavodoxin) {ECO:0000256\|HAMAP-Rule:MF_00159} | + | 5.988338982 | 0 | -3.669188182 | WT unique | 10 | 10 | 30.6 | 30.6 | 40.55 |
| A0A0H3NIF8 | 27.8286 | 27.4577 | 28.0793 | 24.9015 | 24.4912 | 25.5211 | YscW | | + | 2.890261087 | 0.00163636 | -2.817237854 | WT unique | 5 | 5 | 64 | 64 | 19.48 |

### All proteins

| Uniprot accession number | LFQ intensity WT_1 | LFQ intensity WT_2 | LFQ intensity WT_3 | LFQ intensity OrfSwap_1 | LFQ intensity OrfSwap_2 | LFQ intensity OrfSwap_3 | Pfam name | Uniprot full protein name | Student's T-test Significant OrfSwap_WT | -Log Student's T-test p-value OrfSwap_WT | Student's T-test q-value OrfSwap_WT | Student's T-test Difference OrfSwap_WT | Cluster | Peptides | Unique peptides | Sequence coverage [%] | Unique sequence coverage [%] | Mol. weight [kDa] |
|---|---|---|---|---|---|---|---|---|---|---|---|---|---|---|---|---|---|---|
| A0A0H3NEE2 | 28.0983 | 28.0177 | 28.012 | 24.4238 | 25.4347 | 25.6358 | PdxJ | Pyridoxine 5'-phosphate synthase {ECO:0000256\|HAMAP-Rule:MF_00279, ECO:0000256\|SAAS:SAAS00958133} | + | 2.805050289 | 0.00166154 | -2.877882004 | WT unique | 6 | 6 | 31.7 | 31.7 | 26.34 |
| A0A0H3NMF2 | 28.4841 | 28.5426 | 28.8528 | 25.4479 | 26.306 | 26.057 | CheZ | Protein phosphatase CheZ {ECO:0000256\|PIRNR:PIRNR002884} | + | 3.186390898 | 0.0016875 | -2.689495722 | WT unique | 5 | 5 | 30.8 | 30.8 | 23.92 |
| A0A0H3NKF9 | 28.5283 | 28.6327 | 28.4137 | 24.8514 | 26.2431 | 25.5099 | Aminotran_5 | Cysteine desulfurase IscS {ECO:0000256\|HAMAP-Rule:MF_00331, ECO:0000256\|SAAS:SAAS01084503} | + | 2.738538415 | 0.00171429 | -2.990080516 | WT unique | 10 | 10 | 28.7 | 28.7 | 45.08 |
| A0A0H3NEL3 | 28.0153 | 28.1529 | 27.8157 | 25.0612 | 25.6774 | 25.4243 | DUF496 | UPF0265 protein YeeX {ECO:0000256\|HAMAP-Rule:MF_00683} | + | 3.666412189 | 0.00177049 | -2.60696729 | WT unique | 5 | 5 | 45 | 45 | 13.07 |
| A0A0H3NJP0 | 27.8652 | 27.9186 | 27.7703 | 24.9528 | 24.6525 | 25.6034 | Oxidored_q4 | NADH-quinone oxidoreductase subunit A {ECO:0000256\|HAMAP-Rule:MF_01394} | + | 3.215486492 | 0.0018 | -2.781788508 | WT unique | 3 | 3 | 20.4 | 20.4 | 16.49 |

### All proteins

| Uniprot accession number | LFQ intensity WT_1 | LFQ intensity WT_2 | LFQ intensity WT_3 | LFQ intensity OrfSwap_1 | LFQ intensity OrfSwap_2 | LFQ intensity OrfSwap_3 | Pfam name | Uniprot full protein name | Student's T-test Significant OrfSwap_WT | -Log Student's T-test p-value OrfSwap_WT | Student's T-test q-value OrfSwap_WT | Student's T-test Difference OrfSwap_WT | Cluster | Peptides | Unique peptides | Sequence coverage [%] | Unique sequence coverage [%] | Mol. weight [kDa] |
|---|---|---|---|---|---|---|---|---|---|---|---|---|---|---|---|---|---|---|
| A0A0H3NMS7 | 27.9204 | 28.1168 | 28.1086 | 24.1996 | 25.4548 | 25.2623 | PfkB | | + | 2.831756761 | 0.00186207 | -3.07637914 | WT unique | 7 | 7 | 23.3 | 23.3 | 34.09 |
| A0A0H3NDH4 | 27.2824 | 27.7954 | 27.7062 | 25.0347 | 24.7559 | 25.0799 | FAD_binding_4;Lact-deh-memb | Quinone-dependent D-lactate dehydrogenase {ECO:0000256\|HAMAP-Rule:MF_02092} | + | 3.825885449 | 0.00189474 | -2.637835185 | WT unique | 11 | 11 | 21.9 | 21.9 | 65.05 |
| A0A0H3NEM7 | 29.5823 | 27.6187 | 28.1154 | 25.0247 | 24.5587 | 25.0029 | OMP_b-brl | | + | 2.377749563 | 0.00192857 | -3.576712926 | WT unique | 5 | 5 | 31.6 | 31.6 | 18.49 |
| E1WA38 | 28.2192 | 28.2349 | 28.2159 | 24.8905 | 24.4284 | 25.8341 | CsiD | Protein CsiD | + | 2.808369885 | 0.00196364 | -3.172341029 | WT unique | 15 | 15 | 44.9 | 44.9 | 37.23 |
| A0A0H3NBV8 | 27.5574 | 27.6786 | 27.4421 | 24.4987 | 24.0018 | 25.1294 | ADH_N;ADH_zinc_N | S-(hydroxymethyl)glutathione dehydrogenase {ECO:0000256\|RuleBase:RU362016} | + | 3.082606597 | 0.002 | -3.016041438 | WT unique | 8 | 8 | 25.3 | 25.3 | 39.26 |
| A0A0H3NIE9 | 28.1517 | 28.4912 | 27.8466 | 25.0388 | 24.0571 | 25.4402 | Ferritin | Bacterioferritin {ECO:0000256\|PIRNR:PIRNR002560, ECO:0000256\|RuleBase:RU000623} | + | 2.740111586 | 0.00203774 | -3.31780688 | WT unique | 5 | 5 | 38 | 38 | 18.36 |
| A0A0H3NCT3 | 28.7151 | 28.4911 | 28.7421 | 24.105 | 24.1709 | 26.1074 | | | + | 2.364522234 | 0.00207692 | -3.854970296 | WT unique | 15 | 15 | 37.4 | 37.4 | 45.41 |
| A0A0H3NDN6 | 28.6483 | 28.4957 | 28.3476 | 24.3859 | 24.7148 | 25.9737 | GnsAB_toxin | | + | 2.673817233 | 0.00211765 | -3.472373327 | WT unique | 7 | 7 | 87.7 | 87.7 | 6.482 |

### All proteins

| Uniprot accession number | LFQ intensity WT_1 | LFQ intensity WT_2 | LFQ intensity WT_3 | LFQ intensity OrfSwap_1 | LFQ intensity OrfSwap_2 | LFQ intensity OrfSwap_3 | Pfam name | Uniprot full protein name | Student's T-test Significant OrfSwap_WT | -Log Student's T-test p-value OrfSwap_WT | Student's T-test q-value OrfSwap_WT | Student's T-test Difference OrfSwap_WT | Cluster | Peptides | Unique peptides | Sequence coverage [%] | Unique sequence coverage [%] | Mol. weight [kDa] |
|---|---|---|---|---|---|---|---|---|---|---|---|---|---|---|---|---|---|---|
| A0A0H3NHB7 | 28.5665 | 28.3951 | 28.6813 | 26.0194 | 25.0436 | 24.6852 | tRNA-synt_2e | Glycine--tRNA ligase alpha subunit {ECO:0000256\|HAMAP-Rule:MF_00254} | + | 2.898418954 | 0.00216 | -3.298238118 | WT unique | 8 | 8 | 35.3 | 35.3 | 34.75 |
| A0A0H3NF08 | 28.7756 | 28.6228 | 28.4086 | 25.7285 | 24.0574 | 25.3258 | Pentapeptide | | + | 2.642161931 | 0.00220408 | -3.565162659 | WT unique | 6 | 6 | 19.1 | 19.1 | 37.27 |
| A0A0H3N8M9 | 27.8033 | 27.7418 | 27.6006 | 24.7167 | 24.7796 | 25.2736 | | 2-methylisocitrate lyase {ECO:0000256\|HAMAP-Rule:MF_01939} | + | 3.940097405 | 0.00225 | -2.791903814 | WT unique | 8 | 8 | 30.2 | 30.2 | 32 |
| A0A0H3NK16 | 28.2763 | 28.4899 | 28.302 | 24.2363 | 24.8478 | 25.7316 | Response_reg;Trans_reg_C | | + | 2.83223627 | 0.00229787 | -3.417505264 | WT unique | 6 | 6 | 24.6 | 24.6 | 26.27 |
| A0A0H3NEV4 | 28.0449 | 28.0206 | 27.8034 | 24.3547 | 24.6691 | 25.4018 | ParA | Iron-sulfur cluster carrier protein {ECO:0000256\|HAMAP-Rule:MF_02040} | + | 3.224963767 | 0.00234783 | -3.147785823 | WT unique | 5 | 5 | 14.9 | 14.9 | 39.93 |
| A0A0H3NHY1 | 28.0233 | 27.8087 | 27.981 | 24.9447 | 24.3699 | 25.3016 | MttA_Hcf106 | Sec-independent protein translocase protein TatA {ECO:0000256\|HAMAP-Rule:MF_00236} | + | 3.407789974 | 0.0024 | -3.065576553 | WT unique | 3 | 3 | 40.5 | 40.5 | 8.944 |

### All proteins

| Uniprot accession number | LFQ intensity WT_1 | LFQ intensity WT_2 | LFQ intensity WT_3 | LFQ intensity OrfSwap_1 | LFQ intensity OrfSwap_2 | LFQ intensity OrfSwap_3 | Pfam name | Uniprot full protein name | Student's T-test Significant OrfSwap_WT | -Log Student's T-test p-value OrfSwap_WT | Student's T-test q-value OrfSwap_WT | Student's T-test Difference OrfSwap_WT | Cluster | Peptides | Unique peptides | Sequence coverage [%] | Unique sequence coverage [%] | Mol. weight [kDa] |
|---|---|---|---|---|---|---|---|---|---|---|---|---|---|---|---|---|---|---|
| A0A0H3NH18 | 27.6486 | 27.9082 | 27.7491 | 25.184 | 25.0045 | 25.0861 | NTP_transferase | Glucose-1-phosphate adenylyltransferase {ECO:0000256\|HAMAP-Rule:MF_00624} | + | 5.086470567 | 0.00251163 | -2.677073161 | WT unique | 10 | 10 | 26.7 | 26.7 | 48.46 |
| A0A0H3NXF0 | 27.0453 | 27.1707 | 27.0913 | 24.39 | 24.0192 | 24.4176 | AAA_31 | | + | 4.530770445 | 0.00257143 | -2.826892853 | WT unique | 7 | 7 | 17.7 | 17.7 | 44.5 |
| A0A0H3NJW2 | 27.7927 | 27.6008 | 27.4754 | 25.0249 | 25.4541 | 25.35 | | | + | 3.913261434 | 0.00331429 | -2.346627553 | WT unique | 7 | 7 | 31.2 | 31.2 | 22.72 |
| A0A0H3NLU6 | 27.6788 | 27.998 | 27.8323 | 25.7191 | 25.0009 | 24.8476 | PCRF;RF-1 | Peptide chain release factor 2 {ECO:0000256\|HAMAP-Rule:MF_00094, ECO:0000256\|SAAS:SAAS00385830} | + | 3.132296213 | 0.00336232 | -2.647134781 | WT unique | 4 | 4 | 12.8 | 12.8 | 40.13 |
| A0A0H3NC52 | 27.1407 | 27.5607 | 27.3897 | 25.033 | 25.0758 | 24.6399 | adh_short | | + | 3.72734944 | 0.00341176 | -2.447487513 | WT unique | 11 | 11 | 44.3 | 44.3 | 28.07 |
| A0A0H3NFF4 | 27.9475 | 27.8398 | 27.8899 | 25.7076 | 25.377 | 24.8891 | Carboxyl_trans | Acetyl-coenzyme A carboxylase carboxyl transferase subunit beta {ECO:0000256\|HAMAP-Rule:MF_01395} | + | 3.365945137 | 0.00346269 | -2.567860285 | WT unique | 11 | 11 | 30.3 | 30.3 | 33.22 |
| A0A0H3N7S7 | 27.3886 | 27.2631 | 26.843 | 25.1768 | 24.2187 | 25.046 | DUF903 | | + | 2.62825327 | 0.00433735 | -2.351103465 | WT unique | 2 | 2 | 47.4 | 47.4 | 8.217 |
| A0A0H3NJ06 | 27.1276 | 26.7796 | 26.978 | 24.9425 | 25.0646 | 24.9363 | Amidohydro_1 | | + | 4.266165726 | 0.00439024 | -1.980592092 | WT unique | 9 | 9 | 24 | 24 | 41.11 |

### All proteins

| Uniprot accession number | LFQ intensity WT_1 | LFQ intensity WT_2 | LFQ intensity WT_3 | LFQ intensity OrfSwap_1 | LFQ intensity OrfSwap_2 | LFQ intensity OrfSwap_3 | Pfam name | Uniprot full protein name | Student's T-test Significant OrfSwap_WT | -Log Student's T-test p-value OrfSwap_WT | Student's T-test q-value OrfSwap_WT | Student's T-test Difference OrfSwap_WT | Cluster | Peptides | Unique peptides | Sequence coverage [%] | Unique sequence coverage [%] | Mol. weight [kDa] |
|---|---|---|---|---|---|---|---|---|---|---|---|---|---|---|---|---|---|---|
| A0A0H3NFV9 | 28.1103 | 27.8001 | 28.067 | 24.8644 | 24.5463 | 26.245 | Ham1p_like | dITP/XTP pyrophosphatase {ECO:0000256\|HAMAP-Rule:MF_01405} | + | 2.195146588 | 0.00444444 | -2.773890177 | WT unique | 7 | 7 | 42.1 | 42.1 | 21.03 |
| A0A0H3NFW7 | 27.3607 | 27.3418 | 27.3665 | 25.1336 | 24.0358 | 25.281 | AIRS;AIRS_C | Selenide, water dikinase {ECO:0000256\|HAMAP-Rule:MF_00625} | + | 2.530225191 | 0.0045 | -2.539550781 | WT unique | 6 | 6 | 20.5 | 20.5 | 36.44 |
| A0A0H3NM92 | 27.4465 | 26.681 | 27.3945 | 23.5621 | 25.1878 | 23.7653 | ABC_tran;BCA_ABC_TP_C | | + | 2.21157288 | 0.00461538 | -3.002256393 | WT unique | 6 | 6 | 32 | 32 | 26.8 |
| A0A0H3NPK1 | 28.5238 | 28.6371 | 28.7289 | 26.7585 | 24.7101 | 24.8773 | SAICAR_synt | Phosphoribosylaminoimidazole-succinocarboxamide synthase {ECO:0000256\|HAMAP-Rule:MF_00137, ECO:0000256\|SAAS:SAAS00709724} | + | 2.07091113 | 0.00467532 | -3.181280772 | WT unique | 8 | 8 | 40.5 | 40.5 | 26.91 |
| A0A0H3NH59 | 29.95 | 29.663 | 30.6645 | 28.1231 | 24.2934 | 25.0869 | Usp | Universal stress protein {ECO:0000256\|PIRNR:PIRNR006276} | + | 1.617735849 | 0.00473684 | -4.258038203 | WT unique | 4 | 4 | 32.6 | 32.6 | 16.08 |
| A0A0H3NF33 | 28.3266 | 28.2222 | 28.4165 | 25.5974 | 25.7336 | 26.3525 | OpuAC | | + | 3.276010385 | 0.0048 | -2.42727534 | WT unique | 5 | 5 | 22.1 | 22.1 | 36.15 |

### All proteins

| Uniprot accession number | LFQ intensity WT_1 | LFQ intensity WT_2 | LFQ intensity WT_3 | LFQ intensity OrfSwap_1 | LFQ intensity OrfSwap_2 | LFQ intensity OrfSwap_3 | Pfam name | Uniprot full protein name | Student's T-test Significant OrfSwap_WT | -Log Student's T-test p-value OrfSwap_WT | Student's T-test q-value OrfSwap_WT | Student's T-test Difference OrfSwap_WT | Cluster | Peptides | Unique peptides | Sequence coverage [%] | Unique sequence coverage [%] | Mol. weight [kDa] |
|---|---|---|---|---|---|---|---|---|---|---|---|---|---|---|---|---|---|---|
| A0A0H3NMA2 | 28.0305 | 27.3128 | 26.9765 | 24.8722 | 24.2002 | 25.0001 | DHDPS | N-acetylneuraminate lyase {ECO:0000256\|HAMAP-Rule:MF_01237} | + | 2.638656357 | 0.00486486 | -2.749066035 | WT unique | 9 | 9 | 25.3 | 25.3 | 32.48 |
| A0A0H3NHD1 | 28.4447 | 28.3724 | 28.5579 | 24.6204 | 25.3611 | 26.4047 | NTP_transferase | UTP--glucose-1-phosphate uridylyltransferase {ECO:0000256\|RuleBase:RU361259} | + | 2.34557034 | 0.00493151 | -2.996271133 | WT unique | 8 | 8 | 28.1 | 28.1 | 32.91 |
| A0A0H3NHQ9 | 28.6889 | 28.2751 | 28.4719 | 25.4666 | 25.0056 | 26.4073 | SIS | Glutamine--fructose-6-phosphate aminotransferase [isomerizing] {ECO:0000256\|HAMAP-Rule:MF_00164, ECO:0000256\|SAAS:SAAS00887593} | + | 2.573924888 | 0.005 | -2.852156957 | WT unique | 18 | 18 | 36 | 36 | 66.88 |
| A0A0H3NXX3 | 28.4199 | 28.5144 | 28.6493 | 25.9601 | 24.7315 | 26.1741 | Pterin_bind | Dihydropteroate synthase {ECO:0000256\|RuleBase:RU361205} | + | 2.513036698 | 0.00507042 | -2.905951182 | WT unique | 10 | 10 | 46.1 | 46.1 | 28.47 |
| A0A0H3NFD4 | 28.1221 | 27.8415 | 27.8771 | 24.4831 | 26.1489 | 25.4712 | NUDIX | | + | 2.199686839 | 0.00536471 | -2.579163233 | WT unique | 5 | 5 | 26.1 | 26.1 | 20.91 |

### All proteins

| Uniprot accession number | LFQ intensity WT_1 | LFQ intensity WT_2 | LFQ intensity WT_3 | LFQ intensity OrfSwap_1 | LFQ intensity OrfSwap_2 | LFQ intensity OrfSwap_3 | Pfam name | Uniprot full protein name | Student's T-test Significant OrfSwap_WT | -Log Student's T-test p-value OrfSwap_WT | Student's T-test q-value OrfSwap_WT | Student's T-test Difference OrfSwap_WT | Cluster | Peptides | Unique peptides | Sequence coverage [%] | Unique sequence coverage [%] | Mol. weight [kDa] |
|---|---|---|---|---|---|---|---|---|---|---|---|---|---|---|---|---|---|---|
| A0A0H3NFY4 | 27.7238 | 27.55 | 27.8754 | 25.3731 | 25.7989 | 25.9303 | UxaC | Uronate isomerase {ECO:0000256\|HAMAP-Rule:MF_00675, ECO:0000256\|SAAS:SAAS00387330} | + | 3.326408363 | 0.00651685 | -2.015642802 | WT unique | 8 | 8 | 19.1 | 19.1 | 53.61 |
| A0A0H3NA56 | 27.2624 | 27.2044 | 27.1894 | 25.5529 | 24.6316 | 25.0441 | HMA | | + | 2.880933449 | 0.00659091 | -2.142559687 | WT unique | 3 | 3 | 4.8 | 4.8 | 87.91 |
| A0A0H3NP68 | 27.4177 | 27.2361 | 27.986 | 24.8642 | 24.1721 | 25.8293 | DUF406 | | + | 2.088339157 | 0.00666667 | -2.591416677 | WT unique | 2 | 2 | 21.3 | 21.3 | 10.29 |
| A0A0H3NHK3 | 27.6659 | 27.6868 | 27.6859 | 25.1752 | 24.3323 | 26.0989 | Acetyltransf_11;Hexapep | Acyl-[acyl-carrier-protein]--UDP-N-acetylglucosamine O-acyltransferase {ECO:0000256\|HAMAP-Rule:MF_00387, ECO:0000256\|SAAS:SAAS00720086} | + | 2.080713991 | 0.00773626 | -2.477420171 | WT unique | 7 | 7 | 42.4 | 42.4 | 28.09 |
| A0A0H3NBZ4 | 26.9379 | 27.0027 | 27.1399 | 24.308 | 25.3498 | 24.948 | ACT;Formyl_trans_N | Formyltetrahydrofolate deformylase {ECO:0000256\|HAMAP-Rule:MF_01927} | + | 2.654751214 | 0.00782222 | -2.158186595 | WT unique | 6 | 6 | 31.4 | 31.4 | 31.83 |
| A0A0H3NW85 | 28.0508 | 28.1018 | 27.9655 | 26.683 | 24.4023 | 25.2755 | ADH_N;ADH_zinc_N | | + | 1.750443961 | 0.00844898 | -2.585781097 | WT unique | 6 | 6 | 15.3 | 15.3 | 36.37 |

### All proteins

| Uniprot accession number | LFQ intensity WT_1 | LFQ intensity WT_2 | LFQ intensity WT_3 | LFQ intensity OrfSwap_1 | LFQ intensity OrfSwap_2 | LFQ intensity OrfSwap_3 | Pfam name | Uniprot full protein name | Student's T-test Significant OrfSwap_WT | -Log Student's T-test p-value OrfSwap_WT | Student's T-test q-value OrfSwap_WT | Student's T-test Difference OrfSwap_WT | Cluster | Peptides | Unique peptides | Sequence coverage [%] | Unique sequence coverage [%] | Mol. weight [kDa] |
|---|---|---|---|---|---|---|---|---|---|---|---|---|---|---|---|---|---|---|
| A0A0H3NIR2 | 27.2161 | 27.0739 | 26.9158 | 24.063 | 25.4908 | 24.8361 | Alpha-amylase;Alpha-amylase_C;CBM_48 | 1,4-alpha-glucan branching enzyme GlgB {ECO:0000256\|HAMAP-Rule:MF_00685} | + | 2.241296067 | 0.008625 | -2.271966298 | WT unique | 8 | 8 | 13.2 | 13.2 | 84.27 |
| A0A0H3NFW8 | 27.2732 | 27.2466 | 26.9594 | 25.1129 | 23.7361 | 25.3218 | Iron_traffic | Probable Fe(2+)-trafficking protein {ECO:0000256\|HAMAP-Rule:MF_00686, ECO:0000256\|SAAS:SAAS00944244} | + | 2.063320717 | 0.00890323 | -2.436124166 | WT unique | 8 | 8 | 60.4 | 60.4 | 10.9 |
| A0A0H3NQ36 | 27.8297 | 28.0537 | 27.914 | 25.4824 | 26.366 | 24.7049 | YfiO | Outer membrane protein assembly factor BamD {ECO:0000256\|HAMAP-Rule:MF_00922} | + | 2.121224925 | 0.009 | -2.414691289 | WT unique | 11 | 11 | 51.4 | 51.4 | 27.83 |
| A0A0H3NF42 | 30.0099 | 30.1892 | 29.9482 | 31.1324 | 30.6373 | 31.1868 | CsrA | Translational regulator CsrA {ECO:0000256\|HAMAP-Rule:MF_00167} | | 2.110175741 | 0.01020513 | 0.936394374 | | 5 | 5 | 60.7 | 60.7 | 6.856 |
| A0A0H3NH34 | 29.4942 | 29.2628 | 29.4929 | 30.2239 | 30.3227 | 30.2263 | SBP_bac_8 | | | 3.261561987 | 0.01238554 | 0.840986252 | | 21 | 21 | 54.2 | 54.2 | 47.48 |

#### All proteins

| Uniprot accession number | LFQ intensity WT_1 | LFQ intensity WT_2 | LFQ intensity WT_3 | LFQ intensity OrfSwap_1 | LFQ intensity OrfSwap_2 | LFQ intensity OrfSwap_3 | Pfam name | Uniprot full protein name | Student's T-test Significant OrfSwap_WT | -Log Student's T-test p-value OrfSwap_WT | Student's T-test q-value OrfSwap_WT | Student's T-test Difference OrfSwap_WT | Cluster | Peptides | Unique peptides | Sequence coverage [%] | Unique sequence coverage [%] | Mol. weight [kDa] |
|---|---|---|---|---|---|---|---|---|---|---|---|---|---|---|---|---|---|---|
| A0A0H3NGS2 | 29.4645 | 30.1701 | 30.0329 | 30.5837 | 30.8961 | 31.0258 | Histone_HNS | DNA-binding protein {ECO:0000256\|PIRNR:PIRNR002096} | | 1.697695842 | 0.01287356 | 0.946023305 | | 11 | 10 | 64.7 | 64.7 | 15.49 |
| A0A0H3NBY9 | 31.9832 | 31.5969 | 31.9028 | 32.1941 | 32.8316 | 33.473 | Histone_HNS | DNA-binding protein H-NS | | 1.218906194 | 0.01436559 | 1.005273183 | | 15 | 14 | 78.1 | 78.1 | 15.54 |
| A0A0H3NI16 | 27.6488 | 28.3233 | 27.9427 | 29.4012 | 28.6973 | 28.678 | EFG_C;GTP_EFTU;GTP_EFTU_D2 | | | 1.440816254 | 0.01468132 | 0.953917185 | | 17 | 17 | 25.4 | 25.4 | 67.38 |
| A0A0H3NM23 | 27.9091 | 28.2594 | 27.9195 | 28.7835 | 28.8104 | 29.0007 | DAO | D-amino acid dehydrogenase {ECO:0000256\|HAMAP-Rule:MF_01202} | | 2.47396129 | 0.01501124 | 0.835480372 | | 12 | 12 | 31.7 | 31.7 | 47.88 |
| A0A0H3NS22 | 26.7961 | 26.9383 | 27.664 | 27.9264 | 27.9802 | 28.2151 | BMFP | | | 1.485916209 | 0.01671579 | 0.907743454 | | 6 | 6 | 34.3 | 34.3 | 11.55 |
| A0A0H3NNT6 | 29.8448 | 29.6204 | 29.9592 | 30.8839 | 30.6119 | 30.4071 | Peripla_BP_4 | | | 2.08002455 | 0.01783333 | 0.826202393 | | 16 | 16 | 67.4 | 67.4 | 36.76 |
| A0A0H3NDI2 | 30.6845 | 30.0856 | 30.1439 | 30.8764 | 31.2152 | 31.3879 | DUF892 | | | 1.612781361 | 0.02028283 | 0.8551356 | | 13 | 13 | 60.7 | 60.7 | 18.97 |
| A0A0H3NH98 | 27.917 | 28.2032 | 27.9584 | 28.9134 | 29.0741 | 28.4617 | FtsZ_C;Tubulin | Cell division protein FtsZ {ECO:0000256\|HAMAP-Rule:MF_00909, ECO:0000256\|RuleBase:RU000631} | | 1.746819822 | 0.02095238 | 0.790217082 | | 11 | 11 | 33.9 | 33.9 | 40.32 |
| A0A0H3NB60 | 28.4676 | 28.4943 | 28.4257 | 29.2163 | 29.0839 | 29.1553 | DUF1471 | | | 4.046458115 | 0.02135922 | 0.689332962 | | 15 | 15 | 45.2 | 45.2 | 33.92 |
| A0A0H3NFV7 | 28.7218 | 28.442 | 28.4799 | 29.2774 | 29.4329 | 29.0932 | Lipoprotein_18 | Outer membrane protein assembly factor BamC {ECO:0000256\|HAMAP-Rule:MF_00924} | | 2.264977838 | 0.02210526 | 0.719983419 | | 8 | 8 | 31.4 | 31.4 | 36.94 |

### All proteins

| Uniprot accession number | LFQ intensity WT_1 | LFQ intensity WT_2 | LFQ intensity WT_3 | LFQ intensity OrfSwap_1 | LFQ intensity OrfSwap_2 | LFQ intensity OrfSwap_3 | Pfam name | Uniprot full protein name | Student's T-test Significant OrfSwap_WT | -Log Student's T-test p-value OrfSwap_WT | Student's T-test q-value OrfSwap_WT | Student's T-test Difference OrfSwap_WT | Cluster | Peptides | Unique peptides | Sequence coverage [%] | Unique sequence coverage [%] | Mol. weight [kDa] |
|---|---|---|---|---|---|---|---|---|---|---|---|---|---|---|---|---|---|---|
| A0A0H3NN91 | 27.0707 | 27.6736 | 27.3509 | 28.0936 | 28.2559 | 28.0508 | NTP_transferase | | | 1.84655182 | 0.02235514 | 0.768350601 | | 11 | 11 | 38.2 | 38.2 | 37.91 |
| A0A0H3NGN2 | 27.9625 | 26.6687 | 27.5368 | 28.2007 | 28.3494 | 28.2112 | Pep_deformylase | Peptide deformylase {ECO:0000256\|HAMAP-Rule:MF_00163, ECO:0000256\|SAAS:SAAS01077920} | | 1.058701655 | 0.02244068 | 0.864454905 | | 9 | 9 | 49.1 | 49.1 | 19.28 |
| A0A0H3NTD0 | 28.6127 | 28.57 | 28.485 | 29.3688 | 29.292 | 29.0954 | Peptidase_M3 | | | 2.829441039 | 0.0225 | 0.696178436 | | 22 | 22 | 38.4 | 38.4 | 76.94 |
| A0A0H3N8E3 | 28.3902 | 27.9809 | 28.3204 | 28.9499 | 28.9351 | 28.9732 | Rotamase_2 | Peptidylprolyl isomerase {ECO:0000256\|SAAS:SAAS00523066} | | 2.327322851 | 0.02290909 | 0.722227732 | | 20 | 20 | 36.8 | 36.8 | 68.13 |
| A0A0H3NUC4 | 28.8547 | 28.9954 | 29.5849 | 29.5726 | 30.0769 | 30.1247 | Thioredoxin | Thioredoxin {ECO:0000256\|PIRNR:PIRNR000077} | | 1.283053847 | 0.02520325 | 0.77971522 | | 9 | 9 | 63.3 | 63.3 | 11.81 |
| A0A0H3NDK9 | 28.0596 | 28.4086 | 28.3848 | 28.6439 | 29.0731 | 29.2789 | Channel_Tsx | | | 1.512400843 | 0.0286875 | 0.714342117 | | 4 | 4 | 19.5 | 19.5 | 32.78 |
| A0A0H3NMI2 | 30.2628 | 30.3485 | 30.4073 | 31.1242 | 31.088 | 30.7783 | cNMP_binding;HTH_Crp_2 | | | 2.300085724 | 0.02891339 | 0.657284419 | | 22 | 22 | 79 | 79 | 23.66 |
| A0A0H3NFB3 | 29.9354 | 29.5813 | 29.6782 | 30.5846 | 30.468 | 30.1431 | Complex1_30kDa;Complex1_49kDa | NADH-quinone oxidoreductase subunit C/D {ECO:0000256\|HAMAP-Rule:MF_01359, ECO:0000256\|SAAS:SAAS00014838} | | 1.771673565 | 0.03045926 | 0.666950862 | | 30 | 30 | 51.3 | 51.3 | 68.86 |

#### All proteins

| Uniprot accession number | LFQ intensity WT_1 | LFQ intensity WT_2 | LFQ intensity WT_3 | LFQ intensity OrfSwap_1 | LFQ intensity OrfSwap_2 | LFQ intensity OrfSwap_3 | Pfam name | Uniprot full protein name | Student's T-test Significant OrfSwap_WT | -Log Student's T-test p-value OrfSwap_WT | Student's T-test q-value OrfSwap_WT | Student's T-test Difference OrfSwap_WT | Cluster | Peptides | Unique peptides | Sequence coverage [%] | Unique sequence coverage [%] | Mol. weight [kDa] |
|---|---|---|---|---|---|---|---|---|---|---|---|---|---|---|---|---|---|---|
| A0A0H3NA94 | 26.5176 | 26.7955 | 26.1965 | 27.0459 | 27.2109 | 27.1412 | ACP_syn_III;ACP_syn_III_C | 3-oxoacyl-[acyl-carrier-protein] synthase 3 {ECO:0000256\|HAMAP-Rule:MF_01815} | | 1.606312358 | 0.0413913 | 0.629472733 | | 3 | 3 | 13.9 | 13.9 | 33.52 |
| A0A0H3NI68 | 28.6664 | 24.6578 | 25.5952 | 29.473 | 29.7836 | 29.3405 | Sod_Fe_C;Sod_Fe_N | Superoxide dismutase {ECO:0000256\|RuleBase:RU000414} | | 1.243886799 | 0.01407843 | 3.225894928 | | 8 | 8 | 33 | 33 | 23.08 |
| A0A0H3NE23 | 25.0808 | 24.3939 | 24.9659 | 26.398 | 26.5024 | 26.5457 | TPP_enzyme_C;TPP_enzyme_M;TPP_enzyme_N | | | 2.813567947 | 0.01497521 | 1.668464025 | | 8 | 8 | 15.8 | 15.8 | 59.89 |
| A0A0H3NH15 | 24.4488 | 25.0024 | 25.71 | 26.774 | 26.9574 | 27.1092 | Phosphorylase | Alpha-1,4 glucan phosphorylase {ECO:0000256\|RuleBase:RU000587} | | 2.12957285 | 0.0151 | 1.893123627 | | 10 | 10 | 16.1 | 16.1 | 93.34 |
| E1WAC6 | 26.0947 | 24.7987 | 24.608 | 27.8065 | 27.3001 | 26.8465 | SipA | Cell invasion protein sipA | | 1.777680873 | 0.01617857 | 2.150550207 | | 13 | 13 | 25.5 | 25.5 | 73.94 |
| A0A0H3NC58 | 24.9387 | 25.3528 | 24.8056 | 26.5451 | 26.8231 | 26.9133 | LpoB | Penicillin-binding protein activator LpoB {ECO:0000256\|HAMAP-Rule:MF_01889} | | 3.017930578 | 0.01623077 | 1.728110631 | | 2 | 2 | 13.7 | 13.7 | 22.52 |
| A0A0H3N9R4 | 25.3701 | 24.8993 | 24.8896 | 26.8754 | 26.6554 | 26.6078 | | | | 3.126506054 | 0.01632432 | 1.659876506 | | 4 | 4 | 23.3 | 23.3 | 27.55 |

### All proteins

| Uniprot accession number | LFQ intensity WT_1 | LFQ intensity WT_2 | LFQ intensity WT_3 | LFQ intensity OrfSwap_1 | LFQ intensity OrfSwap_2 | LFQ intensity OrfSwap_3 | Pfam name | Uniprot full protein name | Student's T-test Significant OrfSwap_WT | -Log Student's T-test p-value OrfSwap_WT | Student's T-test q-value OrfSwap_WT | Student's T-test Difference OrfSwap_WT | Cluster | Peptides | Unique peptides | Sequence coverage [%] | Unique sequence coverage [%] | Mol. weight [kDa] |
|---|---|---|---|---|---|---|---|---|---|---|---|---|---|---|---|---|---|---|
| A0A0H3NAD6 | 27.2166 | 24.0845 | 25.5199 | 27.8976 | 27.9859 | 28.7402 | FabA | 3-hydroxydecanoyl-[acyl-carrier-protein] dehydratase {ECO:0000256\|HAMAP-Rule:MF_00405} | | 1.291719926 | 0.01728 | 2.60088412 | | 4 | 4 | 22.1 | 22.1 | 19.05 |
| A0A0H3NHS0 | 23.3792 | 23.8045 | 25.5359 | 26.4629 | 26.4102 | 26.4053 | CopC | | | 1.529303172 | 0.01731298 | 2.1862456 | | 3 | 3 | 20.6 | 20.6 | 13.57 |
| A0A0H3NE42 | 25.2446 | 25.0955 | 24.6685 | 26.764 | 26.476 | 26.5891 | FliL | Flagellar protein FliL {ECO:0000256\|RuleBase:RU364125} | | 2.953778868 | 0.01741935 | 1.606777827 | | 5 | 5 | 27.7 | 27.7 | 17.1 |
| A0A0H3N9E1 | 25.1575 | 26.1632 | 24.6592 | 27.2943 | 27.22 | 27.3304 | YkuD | | | 1.934809318 | 0.01756098 | 1.954936345 | | 8 | 7 | 38.9 | 33.7 | 33.29 |
| A0A0H3NC08 | 25.1815 | 24.6531 | 24.0908 | 25.8722 | 26.8584 | 26.8918 | SBP_bac_5 | | | 1.840174699 | 0.01798496 | 1.898970922 | | 6 | 6 | 13.4 | 13.4 | 59.88 |
| A0A0H3NN98 | 26.3555 | 26.1995 | 24.5085 | 27.6535 | 27.6763 | 27.8611 | Hexapep;NTP_transf_3 | Bifunctional protein GlmU {ECO:0000256\|HAMAP-Rule:MF_01631} | | 1.577214607 | 0.02075912 | 2.042483648 | | 10 | 10 | 25.4 | 25.4 | 49.2 |
| A0A0H3NVY8 | 24.1393 | 23.4942 | 25.1563 | 25.8923 | 25.9946 | 25.9386 | OapA;OapA_N | | | 1.589357984 | 0.03272258 | 1.678588231 | | 3 | 3 | 13.3 | 13.3 | 23.16 |
| A0A0H3NN75 | 25.3105 | 25.4693 | 24.8091 | 26.715 | 26.5074 | 26.3481 | PGM_PMM_I;PGM_PMM_II;PGM_PMM_III;PGM_PMM_IV | | | 2.380081359 | 0.03759494 | 1.327221553 | | 6 | 6 | 17.6 | 17.6 | 52.09 |
| A0A0H3NS54 | 25.1382 | 25.6078 | 27.025 | 27.4402 | 27.693 | 27.7132 | GST_N_2 | | | 1.376266491 | 0.037725 | 1.69181633 | | 8 | 8 | 32.9 | 32.9 | 37.38 |
| A0A0H3NN94 | 24.9726 | 25.1901 | 25.7983 | 26.5901 | 26.6338 | 26.7939 | PBP_like_2 | Phosphate-binding protein PstS {ECO:0000256\|PIRNR:PIRNR002756} | | 2.218008458 | 0.03796226 | 1.352270762 | | 5 | 5 | 19.4 | 19.4 | 36.82 |

### All proteins

| Uniprot accession number | LFQ intensity WT_1 | LFQ intensity WT_2 | LFQ intensity WT_3 | LFQ intensity OrfSwap_1 | LFQ intensity OrfSwap_2 | LFQ intensity OrfSwap_3 | Pfam name | Uniprot full protein name | Student's T-test Significant OrfSwap_WT | -Log Student's T-test p-value OrfSwap_WT | Student's T-test q-value OrfSwap_WT | Student's T-test Difference OrfSwap_WT | Cluster | Peptides | Unique peptides | Sequence coverage [%] | Unique sequence coverage [%] | Mol. weight [kDa] |
|---|---|---|---|---|---|---|---|---|---|---|---|---|---|---|---|---|---|---|
| A0A0H3NIZ4 | 25.4523 | 25.8803 | 25.4605 | 26.6878 | 26.8281 | 26.8246 | DUF615 | UPF0307 protein YjgA {ECO:0000256\|HAMAP-Rule:MF_00765} | | 2.868417243 | 0.04830233 | 1.182442983 | | 4 | 4 | 24 | 24 | 21.39 |
| A0A0H3N9R9 | 24.7859 | 25.9196 | 24.6157 | 26.2652 | 26.7372 | 26.7787 | Pectinesterase | | | 1.552180834 | 0.04852071 | 1.486635208 | | 5 | 5 | 16.4 | 16.4 | 45.91 |
| A0A0H3NCJ4 | 29.9639 | 30.0466 | 30.0721 | 29.1143 | 29.1491 | 29.2203 | Anticodon_1;tRNA-synt_1;zf-FPG_IleRS | Isoleucine--tRNA ligase {ECO:0000256\|HAMAP-Rule:MF_02002} | | 4.361200998 | 0.01002985 | -0.866315206 | | 29 | 29 | 31.2 | 31.2 | 105.7 |
| A0A0H3NHC6 | 28.8402 | 28.9023 | 29.1544 | 28.0509 | 28.2174 | 28.0219 | FMN_dh | L-lactate dehydrogenase {ECO:0000256\|HAMAP-Rule:MF_01559, ECO:0000256\|SAAS:SAAS00848080} | | 2.801834128 | 0.01135802 | -0.868907293 | | 16 | 16 | 41.2 | 41.2 | 42.71 |
| A0A0H3NAM6 | 30.4473 | 30.0813 | 29.976 | 29.2138 | 29.527 | 28.8374 | PNPase_C;RNase_E_G;S1 | Ribonuclease E {ECO:0000256\|HAMAP-Rule:MF_00970} | | 1.784356446 | 0.0115 | -0.975446065 | | 25 | 25 | 33.9 | 33.9 | 119.4 |
| A0A0H3NBV6 | 28.4468 | 28.4108 | 28.3044 | 26.4044 | 27.7461 | 27.6306 | Sua5_yciO_yrdC | | | 1.227211749 | 0.01164557 | -1.127044042 | | 7 | 7 | 29.6 | 29.6 | 23.11 |
| A0A0H3NE47 | 30.1221 | 30.2894 | 30.2741 | 29.4222 | 29.5458 | 28.9395 | Transket_pyr;Transketolase_C;Transketolase_N | Transketolase {ECO:0000256\|RuleBase:RU004996} | | 2.066128731 | 0.01253659 | -0.926063538 | | 24 | 24 | 41.3 | 41.3 | 73.03 |
| A0A0H3NS38 | 29.0402 | 28.8453 | 28.9376 | 28.2048 | 28.1796 | 27.8602 | MCPsignal;PAS_3 | | | 2.638236128 | 0.01302326 | -0.85948499 | | 17 | 16 | 31.2 | 31.2 | 55.06 |

### All proteins

| Uniprot accession number | LFQ intensity WT_1 | LFQ intensity WT_2 | LFQ intensity WT_3 | LFQ intensity OrfSwap_1 | LFQ intensity OrfSwap_2 | LFQ intensity OrfSwap_3 | Pfam name | Uniprot full protein name | Student's T-test Significant OrfSwap_WT | -Log Student's T-test p-value OrfSwap_WT | Student's T-test q-value OrfSwap_WT | Student's T-test Difference OrfSwap_WT | Cluster | Peptides | Unique peptides | Sequence coverage [%] | Unique sequence coverage [%] | Mol. weight [kDa] |
|---|---|---|---|---|---|---|---|---|---|---|---|---|---|---|---|---|---|---|
| A0A0H3NSV2 | 31.9471 | 32.2101 | 32.1927 | 31.1305 | 31.3374 | 31.335 | Ribosom_S12_S23 | 30S ribosomal protein S12 {ECO:0000256\|HAMAP-Rule:MF_00403, ECO:0000256\|RuleBase:RU003623} | | 2.831728684 | 0.01317647 | -0.848960241 | | 8 | 8 | 37.9 | 37.9 | 13.74 |
| A0A0H3NGE3 | 31.4149 | 31.3505 | 31.3448 | 30.1223 | 30.2433 | 30.8881 | Ribosomal_L27 | 50S ribosomal protein L27 {ECO:0000256\|HAMAP-Rule:MF_00539} | | 1.788133456 | 0.01333333 | -0.952189128 | | 13 | 13 | 67.1 | 67.1 | 9.124 |
| A0A0H3NSU0 | 33.7902 | 33.7869 | 33.6767 | 33.0943 | 32.9148 | 32.9005 | Ribosomal_L2;Ribosomal_L2_C | 50S ribosomal protein L2 {ECO:0000256\|HAMAP-Rule:MF_01320} | | 3.372484689 | 0.01452174 | -0.781445821 | | 33 | 33 | 79.5 | 79.5 | 29.82 |
| A0A0H3N9M7 | 33.3327 | 33.4343 | 33.4866 | 32.6222 | 32.6402 | 32.6812 | Citrate_synt | Citrate synthase {ECO:0000256\|PIRNR:PIRNR001369, ECO:0000256\|RuleBase:RU003370} | | 4.038407976 | 0.01484444 | -0.770036062 | | 27 | 27 | 49.4 | 49.4 | 48.08 |
| A0A0H3NC63 | 30.1069 | 31.0358 | 30.7483 | 27.4951 | 29.5641 | 30.5403 | KGG | | | 0.694134186 | 0.01518182 | -1.430496216 | | 8 | 5 | 73.3 | 36.7 | 6.225 |
| A0A0H3NIF1 | 30.8474 | 31.0665 | 30.8207 | 29.9902 | 30.1101 | 30.2623 | AAA;Lon_C;LON_substr_bdg | Lon protease {ECO:0000256\|HAMAP-Rule:MF_01973, ECO:0000256\|PIRNR:PIRNR001174} | | 2.691743438 | 0.01689362 | -0.790679932 | | 36 | 36 | 48.3 | 48.3 | 87.41 |

### All proteins

| Uniprot accession number | LFQ intensity WT_1 | LFQ intensity WT_2 | LFQ intensity WT_3 | LFQ intensity OrfSwap_1 | LFQ intensity OrfSwap_2 | LFQ intensity OrfSwap_3 | Pfam name | Uniprot full protein name | Student's T-test Significant OrfSwap_WT | -Log Student's T-test p-value OrfSwap_WT | Student's T-test q-value OrfSwap_WT | Student's T-test Difference OrfSwap_WT | Cluster | Peptides | Unique peptides | Sequence coverage [%] | Unique sequence coverage [%] | Mol. weight [kDa] |
|---|---|---|---|---|---|---|---|---|---|---|---|---|---|---|---|---|---|---|
| A0A0H3NC01 | 27.2799 | 27.4477 | 27.3335 | 26.4947 | 26.5537 | 26.73 | Molybdopterin;Molydop_binding;Nitr_red_alph_N | | | 3.039740901 | 0.01746939 | -0.760848999 | | 10 | 7 | 10.9 | 7.6 | 140.7 |
| A0A0H3NGY1 | 30.4591 | 30.3306 | 30.1941 | 29.5685 | 29.5291 | 29.6142 | Response_reg;Trans_reg_C | | | 3.150643695 | 0.01764948 | -0.757359187 | | 15 | 15 | 60.3 | 60.3 | 27.35 |
| A0A0H3NIT2 | 27.1741 | 27.125 | 26.8546 | 26.3436 | 26.3613 | 26.1563 | ADH_N;ADH_zinc_N | | | 2.519167085 | 0.02104 | -0.764172236 | | 8 | 8 | 36.1 | 36.1 | 35.15 |
| A0A0H3N9C9 | 31.8062 | 31.1753 | 31.0093 | 30.6788 | 30.3395 | 30.4846 | Usp | | | 1.468806204 | 0.02115385 | -0.829299291 | | 11 | 11 | 80.3 | 80.3 | 15.9 |
| A0A0H3NHI7 | 30.1975 | 29.94 | 30.2231 | 29.5654 | 29.2809 | 29.2602 | Hexapep;Hexapep_2;THDPS_N_2 | 2,3,4,5-tetrahydropyridine-2,6-dicarboxylate N-succinyltransferase {ECO:0000256\|HAMAP-Rule:MF_00811} | | 2.307664926 | 0.02156863 | -0.751409531 | | 21 | 21 | 51.8 | 51.8 | 29.85 |
| A0A0H3NNZ6 | 29.769 | 29.6837 | 29.9417 | 29.1901 | 29.1186 | 28.787 | YfbU | UPF0304 protein YfbU {ECO:0000256\|HAMAP-Rule:MF_00762} | | 2.205348055 | 0.02178218 | -0.766254425 | | 11 | 11 | 60.4 | 60.4 | 19.52 |
| A0A0H3NB09 | 30.1385 | 30.1204 | 30.0623 | 29.3994 | 29.3089 | 29.5587 | HGTP_anticodon;TGS;tRNA_SAD;tRNA-synt_2b | Threonine--tRNA ligase {ECO:0000256\|HAMAP-Rule:MF_00184} | | 3.06312636 | 0.02191304 | -0.684696833 | | 20 | 20 | 37.9 | 37.9 | 73.98 |
| A0A0H3N9N5 | 31.5173 | 31.5677 | 31.6565 | 31.0705 | 30.5638 | 30.8692 | Cyt_bd_oxida_I | | | 2.088942852 | 0.02214815 | -0.745992661 | | 18 | 18 | 20.7 | 20.7 | 58.32 |

### All proteins

| Uniprot accession number | LFQ intensity WT_1 | LFQ intensity WT_2 | LFQ intensity WT_3 | LFQ intensity OrfSwap_1 | LFQ intensity OrfSwap_2 | LFQ intensity OrfSwap_3 | Pfam name | Uniprot full protein name | Student's T-test Significant OrfSwap_WT | -Log Student's T-test p-value OrfSwap_WT | Student's T-test q-value OrfSwap_WT | Student's T-test Difference OrfSwap_WT | Cluster | Peptides | Unique peptides | Sequence coverage [%] | Unique sequence coverage [%] | Mol. weight [kDa] |
|---|---|---|---|---|---|---|---|---|---|---|---|---|---|---|---|---|---|---|
| A0A0H3NLY3 | 29.9818 | 30.1203 | 29.8991 | 29.0762 | 29.2274 | 29.5412 | DAHP_synth_1 | 2-dehydro-3-deoxyphosphooctonate aldolase {ECO:0000256\|HAMAP-Rule:MF_00056} | | 2.046344687 | 0.0222521 | -0.718795141 | | 18 | 18 | 59.5 | 59.5 | 30.8 |
| A0A0H3N8D3 | 30.08 | 29.9091 | 29.8884 | 29.487 | 28.5838 | 29.342 | COX1 | | | 1.341027248 | 0.02230088 | -0.821580251 | | 7 | 7 | 10.7 | 10.7 | 74.28 |
| A0A0H3NE57 | 30.0237 | 29.838 | 29.9858 | 29.1352 | 29.3517 | 29.2492 | | | | 2.94710838 | 0.02256604 | -0.703760147 | | 14 | 14 | 46.4 | 46.4 | 27.87 |
| A0A0H3NI20 | 30.4082 | 30.0826 | 29.8654 | 29.5836 | 29.328 | 29.1807 | PsiF_repeat | | | 1.731712153 | 0.02263248 | -0.754643122 | | 13 | 13 | 67.9 | 67.9 | 11.61 |
| A0A0H3N884 | 33.4144 | 32.9879 | 33.4628 | 32.5492 | 32.5778 | 32.5255 | EF_TS | Elongation factor Ts {ECO:0000256\|HAMAP-Rule:MF_00050, ECO:0000256\|RuleBase:RU000642} | | 2.083213239 | 0.0227027 | -0.737547557 | | 33 | 33 | 58.3 | 58.3 | 30.36 |
| A0A0H3NB04 | 28.7415 | 29.368 | 28.7714 | 28.0632 | 28.2321 | 28.278 | DUF2219 | | | 1.639231949 | 0.02282759 | -0.769187927 | | 11 | 11 | 32.9 | 32.9 | 34.84 |
| A0A0H3NMU6 | 28.887 | 29.3018 | 28.9653 | 28.3582 | 28.5246 | 28.0891 | DALR_1;tRNA_synt_2f | Glycine--tRNA ligase beta subunit {ECO:0000256\|HAMAP-Rule:MF_00255} | | 1.809645925 | 0.02296667 | -0.727416356 | | 18 | 18 | 28.2 | 28.2 | 76.45 |
| A0A0H3NDV2 | 30.2311 | 29.7841 | 30.2275 | 29.4241 | 29.3817 | 29.1962 | Complex1_51K;NADH_4Fe-4S;SLBB | NADH-quinone oxidoreductase subunit F {ECO:0000256\|RuleBase:RU364066} | | 1.982877317 | 0.02311927 | -0.746898015 | | 22 | 22 | 51.7 | 51.7 | 49.25 |

### All proteins

| Uniprot accession number | LFQ intensity WT_1 | LFQ intensity WT_2 | LFQ intensity WT_3 | LFQ intensity OrfSwap_1 | LFQ intensity OrfSwap_2 | LFQ intensity OrfSwap_3 | Pfam name | Uniprot full protein name | Student's T-test Significant OrfSwap_WT | -Log Student's T-test p-value OrfSwap_WT | Student's T-test q-value OrfSwap_WT | Student's T-test Difference OrfSwap_WT | Cluster | Peptides | Unique peptides | Sequence coverage [%] | Unique sequence coverage [%] | Mol. weight [kDa] |
|---|---|---|---|---|---|---|---|---|---|---|---|---|---|---|---|---|---|---|
| A0A0H3NHQ3 | 28.9566 | 28.9824 | 28.7734 | 28.2481 | 28.0282 | 28.4049 | S-AdoMet_synt_C;S-AdoMet_synt_M;S-AdoMet_synt_N | S-adenosylmethionine synthase {ECO:0000256\|HAMAP-Rule:MF_00086} | | 2.218045841 | 0.0248 | -0.677054087 | | 9 | 9 | 23.2 | 23.2 | 41.95 |
| A0A0H3NS35 | 29.3871 | 29.4562 | 29.7045 | 28.3447 | 28.7609 | 29.1741 | Sigma70_ner;Sigma70_r1_1;Sigma70_r1_2;Sigma70_r2;Sigma70_r3;Sigma70_r4 | RNA polymerase sigma factor RpoD {ECO:0000256\|HAMAP-Rule:MF_00963} | | 1.368073385 | 0.025 | -0.756001155 | | 24 | 23 | 33.8 | 32.2 | 70.53 |
| A0A0H3NDE2 | 31.0816 | 31.3005 | 31.3142 | 30.6436 | 30.5893 | 30.4682 | | Enoyl-[acyl-carrier-protein] reductase [NADH] {ECO:0000256\|PIRNR:PIRNR000094} | | 2.72100151 | 0.02540984 | -0.665051778 | | 16 | 16 | 59.5 | 59.5 | 27.76 |
| A0A0H3NJ49 | 30.1365 | 30.4846 | 30.0738 | 29.4621 | 29.4212 | 29.6932 | Cyt_bd_oxida_II | | | 2.000896698 | 0.02561983 | -0.706164678 | | 5 | 5 | 10 | 10 | 42.36 |
| A0A0H3NGS4 | 32.4553 | 32.4053 | 32.4192 | 31.908 | 31.8163 | 31.6482 | Ribosomal_S10 | 30S ribosomal protein S10 {ECO:0000256\|HAMAP-Rule:MF_00508} | | 2.919415874 | 0.02730159 | -0.635765711 | | 17 | 17 | 75.7 | 75.7 | 11.77 |

### All proteins

| Uniprot accession number | LFQ intensity WT_1 | LFQ intensity WT_2 | LFQ intensity WT_3 | LFQ intensity OrfSwap_1 | LFQ intensity OrfSwap_2 | LFQ intensity OrfSwap_3 | Pfam name | Uniprot full protein name | Student's T-test Significant OrfSwap_WT | -Log Student's T-test p-value OrfSwap_WT | Student's T-test q-value OrfSwap_WT | Student's T-test Difference OrfSwap_WT | Cluster | Peptides | Unique peptides | Sequence coverage [%] | Unique sequence coverage [%] | Mol. weight [kDa] |
|---|---|---|---|---|---|---|---|---|---|---|---|---|---|---|---|---|---|---|
| A0A0H3NRH8 | 31.8375 | 31.5963 | 31.9141 | 31.2841 | 31.2191 | 30.7738 | GDC-P | Glycine dehydrogenase (decarboxylating) {ECO:0000256\|HAMAP-Rule:MF_00711} | | 1.679776633 | 0.02824615 | -0.690301895 | | 27 | 27 | 32.5 | 32.5 | 104.3 |
| A0A0H3NHF6 | 30.493 | 30.3734 | 30.2891 | 29.7155 | 29.8586 | 29.6959 | Pribosyl_synth;Pribosyltran_N | Ribose-phosphate pyrophosphokinase {ECO:0000256\|HAMAP-Rule:MF_00583} | | 2.884171683 | 0.02846512 | -0.628475825 | | 16 | 16 | 47.6 | 47.6 | 34.22 |
| A0A0H3NJP4 | 27.483 | 27.0882 | 27.0916 | 26.2183 | 26.6981 | 26.6607 | Aminotran_5 | Phosphoserine aminotransferase {ECO:0000256\|HAMAP-Rule:MF_00160} | | 1.579034634 | 0.02992366 | -0.695254008 | | 7 | 7 | 23.8 | 23.8 | 39.83 |
| A0A0H3NRZ3 | 29.1921 | 29.0834 | 29.0002 | 28.6126 | 28.1943 | 28.5189 | Aldo_ket_red | | | 2.030134293 | 0.02997015 | -0.64993159 | | 14 | 14 | 62.2 | 62.2 | 31 |
| A0A0H3NL53 | 29.2726 | 29.5027 | 29.5313 | 28.9162 | 28.8197 | 28.6668 | PrgH | | | 2.356635755 | 0.03019549 | -0.6346639 | | 17 | 17 | 37.5 | 37.5 | 44.46 |
| A0A0H3NDZ2 | 29.5066 | 29.2139 | 29.1225 | 28.5911 | 28.61 | 28.7069 | tRNA-synt_1c | Glutamate--tRNA ligase {ECO:0000256\|HAMAP-Rule:MF_00022} | | 2.220771972 | 0.03042424 | -0.644983927 | | 14 | 14 | 26.8 | 26.8 | 53.63 |
| A0A0H3NIG4 | 30.8775 | 30.8864 | 30.7684 | 30.0318 | 30.2758 | 30.3467 | HlyD;HlyD_D23 | | | 2.437328598 | 0.03114706 | -0.626003901 | | 18 | 18 | 53.9 | 53.9 | 42.23 |
| A0A0H3NCW1 | 29.3247 | 29.0583 | 29.2019 | 27.9322 | 28.3233 | 29.0558 | DUF945 | | | 1.052901714 | 0.03182482 | -0.757888158 | | 9 | 9 | 27.5 | 27.5 | 54.28 |

### All proteins

| Uniprot accession number | LFQ intensity WT_1 | LFQ intensity WT_2 | LFQ intensity WT_3 | LFQ intensity OrfSwap_1 | LFQ intensity OrfSwap_2 | LFQ intensity OrfSwap_3 | Pfam name | Uniprot full protein name | Student's T-test Significant OrfSwap_WT | -Log Student's T-test p-value OrfSwap_WT | Student's T-test q-value OrfSwap_WT | Student's T-test Difference OrfSwap_WT | Cluster | Peptides | Unique peptides | Sequence coverage [%] | Unique sequence coverage [%] | Mol. weight [kDa] |
|---|---|---|---|---|---|---|---|---|---|---|---|---|---|---|---|---|---|---|
| A0A0H3NDX2 | 28.6932 | 28.6669 | 28.358 | 27.2257 | 25.0228 | 26.2857 | Transcrip_reg | Probable transcriptional regulatory protein YebC {ECO:0000256\|HAMAP-Rule:MF_00693} | | 1.681168677 | 0.01394175 | -2.394609451 | | 6 | 6 | 23.2 | 23.2 | 26.42 |
| A0A0H3NBC9 | 27.388 | 26.9729 | 27.0007 | 25.3936 | 24.2108 | 25.444 | GFO_IDH_MocA;GFO_IDH_MocA_C | | | 2.111935521 | 0.01421782 | -2.104417801 | | 10 | 10 | 32.1 | 32.1 | 38.41 |
| A0A0H3NJB0 | 27.7199 | 27.3307 | 27.5285 | 24.5847 | 25.3819 | 26.3525 | | | | 1.787792723 | 0.01485246 | -2.086694082 | | 4 | 4 | 23.3 | 23.3 | 26.83 |
| A0A0H3NH16 | 26.5808 | 26.7705 | 26.8379 | 25.4858 | 24.0896 | 24.7835 | Usp | | | 2.042747907 | 0.01522689 | -1.943431218 | | 9 | 9 | 28.9 | 28.9 | 35.64 |
| A0A0H3NKJ6 | 27.1814 | 27.1489 | 27.2247 | 24.2959 | 25.5129 | 25.745 | ELFV_dehydrog;ELFV_dehydrog_N | Glutamate dehydrogenase {ECO:0000256\|PIRNR:PIRNR000185} | | 1.947601089 | 0.01535593 | -2.000388463 | | 7 | 7 | 20.8 | 20.8 | 48.57 |
| A0A0H3NHH1 | 27.4781 | 27.248 | 27.5139 | 25.3282 | 25.2525 | 26.2066 | DUF1732;YicC_N | | | 2.337213436 | 0.01548718 | -1.817578634 | | 5 | 5 | 22 | 22 | 33.13 |
| A0A0H3P1B8 | 28.3393 | 27.9341 | 28.0893 | 24.842 | 27.2703 | 24.2925 | | | | 1.344767741 | 0.01562069 | -2.652627945 | | 4 | 4 | 17.9 | 17.9 | 29.91 |
| A0A0H3N891 | 26.5909 | 26.7227 | 27.01 | 24.9929 | 25.0366 | 22.7233 | FabA | 3-hydroxyacyl-[acyl-carrier-protein] dehydratase FabZ {ECO:0000256\|HAMAP-Rule:MF_00406} | | 1.507904465 | 0.01562963 | -2.523615519 | | 4 | 4 | 34.4 | 34.4 | 17 |
| A0A0H3P1B9 | 26.2688 | 26.5714 | 26.3422 | 24.1943 | 23.9753 | 25.2289 | | | | 2.080274897 | 0.01575652 | -1.928028742 | | 4 | 4 | 33 | 33 | 10.88 |

### All proteins

| Uniprot accession number | LFQ intensity WT_1 | LFQ intensity WT_2 | LFQ intensity WT_3 | LFQ intensity OrfSwap_1 | LFQ intensity OrfSwap_2 | LFQ intensity OrfSwap_3 | Pfam name | Uniprot full protein name | Student's T-test Significant OrfSwap_WT | -Log Student's T-test p-value OrfSwap_WT | Student's T-test q-value OrfSwap_WT | Student's T-test Difference OrfSwap_WT | Cluster | Peptides | Unique peptides | Sequence coverage [%] | Unique sequence coverage [%] | Mol. weight [kDa] |
|---|---|---|---|---|---|---|---|---|---|---|---|---|---|---|---|---|---|---|
| A0A0H3NHG4 | 26.173 | 26.0895 | 26.2945 | 24.3938 | 24.3391 | 24.813 | Thymidylat_synt | Thymidylate synthase {ECO:0000256\|HAMAP-Rule:MF_00008, ECO:0000256\|SAAS:SAAS01077528} | | 3.311594927 | 0.0157757 | -1.670333862 | | 2 | 2 | 10.6 | 10.6 | 30.37 |
| A0A0H3NCS2 | 28.6065 | 29.233 | 28.5427 | 27.8326 | 25.1119 | 25.7899 | IMPDH | GMP reductase {ECO:0000256\|HAMAP-Rule:MF_00596} | | 1.40307544 | 0.01589474 | -2.549289068 | | 11 | 11 | 25.6 | 25.6 | 37.14 |
| A0A0H3NRU4 | 27.6621 | 27.5341 | 27.6077 | 24.2463 | 26.5302 | 24.6821 | UxuA | Mannonate dehydratase {ECO:0000256\|HAMAP-Rule:MF_00106, ECO:0000256\|SAAS:SAAS01090786} | | 1.600924686 | 0.01592453 | -2.448438009 | | 9 | 9 | 29.4 | 29.4 | 44.94 |
| A0A0H3N869 | 26.8627 | 26.5038 | 26.7674 | 24.4652 | 24.0253 | 25.5502 | PNP_UDP_1 | 5'-methylthioadenosine/S-adenosylhomocysteine nucleosidase {ECO:0000256\|HAMAP-Rule:MF_01684} | | 1.919046108 | 0.0160354 | -2.03101031 | | 2 | 2 | 10.3 | 10.3 | 24.45 |
| A0A0H3NDB2 | 27.5928 | 27.6972 | 27.173 | 25.9945 | 23.4339 | 25.3262 | NTP_transferase | | | 1.515715876 | 0.01607619 | -2.569473267 | | 3 | 3 | 11.3 | 11.3 | 29.04 |

#### All proteins

| Uniprot accession number | LFQ intensity WT_1 | LFQ intensity WT_2 | LFQ intensity WT_3 | LFQ intensity OrfSwap_1 | LFQ intensity OrfSwap_2 | LFQ intensity OrfSwap_3 | Pfam name | Uniprot full protein name | Student's T-test Significant OrfSwap_WT | -Log Student's T-test p-value OrfSwap_WT | Student's T-test q-value OrfSwap_WT | Student's T-test Difference OrfSwap_WT | Cluster | Peptides | Unique peptides | Sequence coverage [%] | Unique sequence coverage [%] | Mol. weight [kDa] |
|---|---|---|---|---|---|---|---|---|---|---|---|---|---|---|---|---|---|---|
| A0A0H3NNV1 | 27.1854 | 27.4283 | 27.263 | 24.3949 | 25.3322 | 25.9614 | Proteasome | ATP-dependent protease subunit HslV {ECO:0000256\|HAMAP-Rule:MF_00248, ECO:0000256\|SAAS:SAAS00347004} | | 1.958084317 | 0.01647273 | -2.062766393 | | 3 | 3 | 17 | 17 | 18.99 |
| A0A0H3NPN9 | 27.2951 | 27.1716 | 27.2677 | 24.8389 | 24.7297 | 26.2449 | Inositol_P | 3'(2'),5'-bisphosphate nucleotidase CysQ {ECO:0000256\|HAMAP-Rule:MF_02095} | | 1.804445689 | 0.01661538 | -1.973617554 | | 10 | 10 | 55.3 | 55.3 | 27.49 |
| A0A0H3NW54 | 27.107 | 27.5383 | 27.3527 | 24.8974 | 25.817 | 25.7003 | OTCace;OTCace_N | Aspartate carbamoyltransferase {ECO:0000256\|HAMAP-Rule:MF_00001} | | 2.386978285 | 0.01662385 | -1.861101786 | | 7 | 7 | 28.9 | 28.9 | 34.38 |
| A0A0H3NBE5 | 26.553 | 26.9426 | 26.9616 | 25.237 | 24.442 | 25.4605 | PNTB | NAD(P) transhydrogenase subunit beta {ECO:0000256\|PIRNR:PIRNR000204} | | 2.20597105 | 0.01674419 | -1.772576014 | | 6 | 6 | 18 | 18 | 48.87 |

### All proteins

| Uniprot accession number | LFQ intensity WT_1 | LFQ intensity WT_2 | LFQ intensity WT_3 | LFQ intensity OrfSwap_1 | LFQ intensity OrfSwap_2 | LFQ intensity OrfSwap_3 | Pfam name | Uniprot full protein name | Student's T-test Significant OrfSwap_WT | -Log Student's T-test p-value OrfSwap_WT | Student's T-test q-value OrfSwap_WT | Student's T-test Difference OrfSwap_WT | Cluster | Peptides | Unique peptides | Sequence coverage [%] | Unique sequence coverage [%] | Mol. weight [kDa] |
|---|---|---|---|---|---|---|---|---|---|---|---|---|---|---|---|---|---|---|
| A0A0H3NDN3 | 29.8384 | 29.7879 | 29.6497 | 24.9609 | 29.4651 | 24.6295 | CLP_protease | ATP-dependent Clp protease proteolytic subunit {ECO:0000256\|HAMAP-Rule:MF_00444, ECO:0000256\|RuleBase:RU003567} | | 1.024933375 | 0.016875 | -3.406876882 | | 5 | 5 | 29.5 | 29.5 | 23.18 |
| A0A0H3NGW8 | 27.5119 | 27.3137 | 27.0426 | 24.6884 | 26.1908 | 25.1047 | PALP;Thr_synth_N | | | 1.860135889 | 0.01700787 | -1.961413066 | | 6 | 6 | 16.6 | 16.6 | 46.99 |
| A0A0H3NEY3 | 27.1921 | 26.9688 | 27.0538 | 25.3797 | 24.2029 | 25.7401 | Molybdop_Fe4S4;Molybdopterin;Molydop_binding | | | 1.860137714 | 0.01714286 | -1.964018504 | | 7 | 7 | 11.9 | 11.9 | 90.35 |
| A0A0H3NFS3 | 25.5544 | 25.7417 | 25.9775 | 24.4235 | 24.0592 | 24.2499 | Sdh5 | | | 3.14243726 | 0.01812121 | -1.513659159 | | 2 | 2 | 18.2 | 18.2 | 10.55 |
| A0A0H3ND72 | 26.9636 | 27.3248 | 27.1112 | 25.3835 | 24.9663 | 25.96 | Peptidase_M15 | D-alanyl-D-alanine dipeptidase {ECO:0000256\|HAMAP-Rule:MF_01924, ECO:0000256\|PIRNR:PIRNR026671} | | 2.283114293 | 0.01877612 | -1.696570079 | | 8 | 8 | 33.2 | 33.2 | 29 |
| A0A0H3NIV9 | 26.9193 | 26.9552 | 26.7392 | 25.8617 | 24.6887 | 24.5697 | AA_permease | | | 1.926926844 | 0.02035556 | -1.831206004 | | 3 | 3 | 11.3 | 11.3 | 51.34 |

### All proteins

| Uniprot accession number | LFQ intensity WT_1 | LFQ intensity WT_2 | LFQ intensity WT_3 | LFQ intensity OrfSwap_1 | LFQ intensity OrfSwap_2 | LFQ intensity OrfSwap_3 | Pfam name | Uniprot full protein name | Student's T-test Significant OrfSwap_WT | -Log Student's T-test p-value OrfSwap_WT | Student's T-test q-value OrfSwap_WT | Student's T-test Difference OrfSwap_WT | Cluster | Peptides | Unique peptides | Sequence coverage [%] | Unique sequence coverage [%] | Mol. weight [kDa] |
|---|---|---|---|---|---|---|---|---|---|---|---|---|---|---|---|---|---|---|
| A0A0H3N7Z5 | 27.3193 | 27.2774 | 27.0832 | 25.5472 | 25.8986 | 25.8687 | SHS2_FTSA | Cell division protein FtsA {ECO:0000256\|HAMAP-Rule:MF_02033, ECO:0000256\|PIRNR:PIRNR003101} | | 3.390001525 | 0.02091176 | -1.455176036 | | 6 | 6 | 24 | 24 | 45.31 |
| A0A0H3NBQ1 | 28.589 | 28.7803 | 28.6177 | 28.1225 | 24.3044 | 25.4425 | | | | 1.122828009 | 0.02123741 | -2.705876668 | | 11 | 11 | 33.2 | 33.2 | 41.57 |
| A0A0H3NMS0 | 28.6006 | 28.5461 | 28.6815 | 27.9712 | 25.4689 | 25.1048 | Pyr_redox_2;Pyr_redox_dim | | | 1.263332267 | 0.0213913 | -2.427788417 | | 10 | 10 | 33.6 | 33.6 | 48.69 |
| A0A0H3NDK1 | 26.8806 | 27.0653 | 26.9039 | 25.347 | 24.4355 | 25.8276 | Fer4_11;Nitr_red_bet_C | | | 1.875882696 | 0.02173793 | -1.746571859 | | 6 | 2 | 13.5 | 6.5 | 57.8 |
| A0A0H3NGW5 | 26.8528 | 26.875 | 27.1921 | 25.066 | 25.0444 | 25.9139 | tRNA-synt_1b | Tryptophan--tRNA ligase {ECO:0000256\|HAMAP-Rule:MF_00140} | | 2.222311892 | 0.02185714 | -1.631866455 | | 6 | 6 | 22.8 | 22.8 | 37.4 |
| A0A0H3NG22 | 25.9414 | 26.4309 | 26.2776 | 25.015 | 24.6645 | 24.3808 | Inositol_P | Inositol-1-monophosphatase {ECO:0000256\|RuleBase:RU364068} | | 2.551669516 | 0.02188889 | -1.529853821 | | 4 | 4 | 19.5 | 19.5 | 29.16 |
| A0A0H3NHU2 | 27.6132 | 27.7766 | 28.0045 | 27.3463 | 24.1956 | 24.2031 | Acyl-CoA_dh_1;Acyl-CoA_dh_M;Acyl-CoA_dh_N;DUF1974 | | | 1.136485512 | 0.02204196 | -2.549728394 | | 14 | 14 | 19.5 | 19.5 | 89.22 |

### All proteins

| Uniprot accession number | LFQ intensity WT_1 | LFQ intensity WT_2 | LFQ intensity WT_3 | LFQ intensity OrfSwap_1 | LFQ intensity OrfSwap_2 | LFQ intensity OrfSwap_3 | Pfam name | Uniprot full protein name | Student's T-test Significant OrfSwap_WT | -Log Student's T-test p-value OrfSwap_WT | Student's T-test q-value OrfSwap_WT | Student's T-test Difference OrfSwap_WT | Cluster | Peptides | Unique peptides | Sequence coverage [%] | Unique sequence coverage [%] | Mol. weight [kDa] |
|---|---|---|---|---|---|---|---|---|---|---|---|---|---|---|---|---|---|---|
| A0A0H3NE53 | 27.7038 | 25.6345 | 27.4268 | 25.1184 | 24.8942 | 24.5064 | DPBB_1;SPOR | Endolytic peptidoglycan transglycosylase RlpA {ECO:0000256\|HAMAP-Rule:MF_02071} | | 1.438896055 | 0.02219718 | -2.082028071 | | 9 | 9 | 33.2 | 33.2 | 36.88 |
| A0A0H3N9K7 | 27.0426 | 27.4305 | 27.0958 | 26.2217 | 25.1779 | 24.448 | SIS_2 | Phosphoheptose isomerase {ECO:0000256\|HAMAP-Rule:MF_00067, ECO:0000256\|SAAS:SAAS00846111} | | 1.645332972 | 0.02235461 | -1.907088598 | | 5 | 5 | 22.9 | 22.9 | 20.9 |
| A0A0H3NMK3 | 26.3644 | 25.9698 | 25.9825 | 24.5523 | 24.3316 | 24.9678 | YedD | | | 2.552796142 | 0.02380822 | -1.488344828 | | 4 | 4 | 50.4 | 50.4 | 15.36 |
| A0A0H3NGE4 | 24.8943 | 29.6303 | 29.632 | 24.7441 | 24.5168 | 25.5541 | polyprenyl_synt | | | 0.902286527 | 0.02663946 | -3.113884608 | | 2 | 2 | 10.5 | 10.5 | 35.27 |
| A0A0H3NRG8 | 26.6278 | 26.9292 | 26.5892 | 25.1009 | 25.4587 | 25.4955 | DsbC_N;Thioredoxin_2 | Thiol:disulfide interchange protein {ECO:0000256\|RuleBase:RU364038} | | 2.927183468 | 0.02944966 | -1.363728841 | | 3 | 3 | 18.8 | 18.8 | 24.3 |
| A0A0H3NL63 | 27.5254 | 27.5092 | 27.7644 | 25.3321 | 26.917 | 24.7195 | FliMN_C | | | 1.37495992 | 0.02964865 | -1.94346873 | | 8 | 8 | 31.4 | 31.4 | 33.79 |
| A0A0H3NK36 | 27.4525 | 26.9054 | 26.6334 | 26.0987 | 25.1185 | 24.1519 | FBPase_glpX | Fructose-1,6-bisphosphatase {ECO:0000256\|PIRNR:PIRNR004532} | | 1.426244684 | 0.03072 | -1.87406222 | | 5 | 5 | 16.1 | 16.1 | 35.66 |

### All proteins

| Uniprot accession number | LFQ intensity WT_1 | LFQ intensity WT_2 | LFQ intensity WT_3 | LFQ intensity OrfSwap_1 | LFQ intensity OrfSwap_2 | LFQ intensity OrfSwap_3 | Pfam name | Uniprot full protein name | Student's T-test Significant OrfSwap_WT | -Log Student's T-test p-value OrfSwap_WT | Student's T-test q-value OrfSwap_WT | Student's T-test Difference OrfSwap_WT | Cluster | Peptides | Unique peptides | Sequence coverage [%] | Unique sequence coverage [%] | Mol. weight [kDa] |
|---|---|---|---|---|---|---|---|---|---|---|---|---|---|---|---|---|---|---|
| A0A0H3NJ94 | 27.1268 | 27.1537 | 27.393 | 25.0585 | 24.9925 | 26.4894 | PyrI;PyrI_C | Aspartate carbamoyltransferase regulatory chain {ECO:0000256\|HAMAP-Rule:MF_00002} | | 1.585032098 | 0.03194737 | -1.711028417 | | 8 | 8 | 63.4 | 63.4 | 17.09 |
| A0A0H3NB32 | 25.892 | 25.7839 | 25.9964 | 24.8775 | 24.4441 | 24.0889 | FAD_binding_4;FAD-oxidase_C;Fer4_8 | | | 2.415259352 | 0.03215894 | -1.420576731 | | 4 | 4 | 4 | 4 | 113.1 |
| A0A0H3N968 | 28.5237 | 28.6921 | 28.7754 | 26.0807 | 24.7088 | 28.262 | DALR_2;tRNA-synt_1e | Cysteine--tRNA ligase {ECO:0000256\|HAMAP-Rule:MF_00041} | | 1.047760612 | 0.03233766 | -2.313212077 | | 13 | 13 | 26.7 | 26.7 | 52.26 |
| A0A0H3NIE2 | 26.6603 | 26.6227 | 26.9109 | 25.4335 | 24.9705 | 25.6388 | Lipoprotein_16 | | | 2.505662582 | 0.03254902 | -1.38368988 | | 4 | 4 | 18.6 | 18.6 | 22.81 |
| A0A0H3NMJ3 | 26.2889 | 26.3109 | 26.0235 | 25.3636 | 23.9945 | 24.5265 | | Phosphoglycolate phosphatase {ECO:0000256\|HAMAP-Rule:MF_00495} | | 1.741783732 | 0.03451282 | -1.579556783 | | 6 | 6 | 25.4 | 25.4 | 27.3 |
| A0A0H3NG91 | 25.8056 | 25.7487 | 25.9851 | 24.6048 | 24.7681 | 24.0916 | NAD_binding_11;NAD_binding_2 | 2-hydroxy-3-oxopropionate reductase {ECO:0000256\|HAMAP-Rule:MF_02032} | | 2.486755054 | 0.0350828 | -1.358270009 | | 4 | 4 | 20.6 | 20.6 | 30.73 |
| A0A0H3P187 | 25.0022 | 27.2752 | 27.298 | 23.8069 | 24.8642 | 24.9566 | KfrA_N | | | 1.101811983 | 0.03749068 | -1.98255221 | | 8 | 8 | 24.8 | 24.8 | 41.48 |

### All proteins

| Uniprot accession number | LFQ intensity WT_1 | LFQ intensity WT_2 | LFQ intensity WT_3 | LFQ intensity OrfSwap _1 | LFQ intensity OrfSwap _2 | LFQ intensity OrfSwap _3 | Pfam name | Uniprot full protein name | Student's T-test Significant OrfSwap_WT | -Log Student's T-test p-value OrfSwap_WT | Student's T-test q-value OrfSwap_WT | Student's T-test Difference OrfSwap_WT | Cluster | Peptides | Unique peptides | Sequence coverage [%] | Unique sequence coverage [%] | Mol. weight [kDa] |
|---|---|---|---|---|---|---|---|---|---|---|---|---|---|---|---|---|---|---|
| A0A0H3N847 | 29.4341 | 29.3314 | 29.5067 | 29.9166 | 25.3859 | 24.2991 | zf-dskA_traR | RNA polymerase-binding transcription factor DksA {ECO:0000256 \|HAMAP-Rule:MF_00926} | | 0.773746808 | 0.04079012 | -2.890249888 | | 7 | 7 | 65.6 | 65.6 | 17.52 |
| A0A0H3NFT8 | 28.0038 | 27.8389 | 27.8943 | 25.5121 | 27.5251 | 25.2172 | SIMPL | | | 1.182213724 | 0.04130061 | -1.827534993 | | 8 | 8 | 35.5 | 35.5 | 26.5 |
| A0A0H3NDR2 | 26.8972 | 27.3824 | 27.0359 | 26.1817 | 25.7731 | 25.1275 | YbaB_DNA_bd | Nucleoid-associated protein YbaB {ECO:0000256 \|HAMAP-Rule:MF_00274} | | 1.849841649 | 0.04145455 | -1.41103363 | | 3 | 3 | 20.2 | 20.2 | 12.02 |
| A0A0H3NKK1 | 26.1472 | 26.3573 | 26.3063 | 24.9607 | 24.8317 | 25.3325 | AstA | Arginine N-succinyltransferase {ECO:0000256 \|HAMAP-Rule:MF_01171, ECO:0000256\|SAAS:SAAS00376548} | | 2.781107562 | 0.04170732 | -1.228658676 | | 6 | 6 | 21.8 | 21.8 | 38.28 |
| A0A0H3NWV8 | 27.5237 | 27.1606 | 26.0319 | 24.7419 | 24.0727 | 26.2676 | StbA | | | 1.11840912 | 0.04253012 | -1.878040314 | | 6 | 6 | 18.2 | 18.2 | 35.71 |
| A0A0H3N9L5 | 27.8519 | 25.3238 | 27.8543 | 24.9774 | 25.3523 | 24.9823 | eIF-1a | Translation initiation factor IF-1 {ECO:0000256 \|HAMAP-Rule:MF_00075} | | 1.050833713 | 0.04802312 | -1.905989965 | | 3 | 3 | 50 | 50 | 8.25 |
| A0A0H3NG56 | 26.5806 | 26.8642 | 26.9045 | 24.3083 | 25.1286 | 26.143 | Peptidase_S24;Peptidase_S26 | Signal peptidase I {ECO:0000256 \|RuleBase:RU003993} | | 1.373711337 | 0.04807143 | -1.589796066 | | 6 | 6 | 21.9 | 21.9 | 35.78 |

### All proteins

| Uniprot accession number | LFQ intensity WT_1 | LFQ intensity WT_2 | LFQ intensity WT_3 | LFQ intensity OrfSwap_1 | LFQ intensity OrfSwap_2 | LFQ intensity OrfSwap_3 | Pfam name | Uniprot full protein name | Student's T-test Significant OrfSwap_WT | -Log Student's T-test p-value OrfSwap_WT | Student's T-test q-value OrfSwap_WT | Student's T-test Difference OrfSwap_WT | Cluster | Peptides | Unique peptides | Sequence coverage [%] | Unique sequence coverage [%] | Mol. weight [kDa] |
|---|---|---|---|---|---|---|---|---|---|---|---|---|---|---|---|---|---|---|
| A0A0H3NCB2 | 26.7566 | 27.1383 | 26.8464 | 26.2646 | 25.0613 | 24.9746 | Aldolase | | | 1.574986702 | 0.04835928 | -1.480258306 | | 4 | 4 | 18.3 | 18.3 | 22.19 |
| A0A0H3NPP5 | 26.7093 | 26.3499 | 26.5908 | 25.2878 | 25.1968 | 25.5966 | Mur_ligase; Mur_ligase_C;Mur_ligase_M | UDP-N-acetylmuramate--L-alanyl-gamma-D-glutamyl-meso-2,6-diaminoheptandioate ligase {ECO:0000256\|HAMAP-Rule:MF_02020} | | 2.751002554 | 0.04845977 | -1.189585368 | | 6 | 6 | 14.4 | 14.4 | 50.05 |
| A0A0H3N8V0 | 26.6722 | 26.6773 | 26.7655 | 26.3652 | 24.4104 | 23.9763 | Sec_GG;SecD_SecF | Protein-export membrane protein SecF {ECO:0000256\|HAMAP-Rule:MF_01464} | | 1.143330763 | 0.0485848 | -1.787740707 | | 3 | 3 | 15.2 | 15.2 | 35.38 |
| A0A0H3NPU7 | 27.2258 | 27.9406 | 27.8118 | 26.9054 | 26.0254 | 25.5585 | AIRS_C | Phosphoribosylformylglycinamidine synthase {ECO:0000256\|HAMAP-Rule:MF_00419} | | 1.528067484 | 0.04887059 | -1.496312459 | | 9 | 9 | 9.5 | 9.5 | 141.5 |
| A0A0H3N983 | 26.6154 | 26.2633 | 26.6974 | 25.0245 | 24.4863 | 25.7784 | GalKase_gal_bdg;GHMP_kinases_C; GHMP_kinases_N | Galactokinase {ECO:0000256\|HAMAP-Rule:MF_00246, ECO:0000256\|SAAS:SAAS00350925} | | 1.640204268 | 0.04891429 | -1.428923925 | | 6 | 6 | 14.1 | 14.1 | 41.3 |
| A0A0H3NHB8 | 26.2572 | 26.6078 | 26.5175 | 24.5257 | 25.2373 | 25.5771 | ABC_tran;oligo_HPY | | | 1.835039945 | 0.04960452 | -1.347492854 | | 5 | 5 | 21.3 | 21.3 | 37.25 |

### All proteins

| Uniprot accession number | LFQ intensity WT_1 | LFQ intensity WT_2 | LFQ intensity WT_3 | LFQ intensity OrfSwap_1 | LFQ intensity OrfSwap_2 | LFQ intensity OrfSwap_3 | Pfam name | Uniprot full protein name | Student's T-test Significant OrfSwap_WT | -Log Student's T-test p-value OrfSwap_WT | Student's T-test q-value OrfSwap_WT | Student's T-test Difference OrfSwap_WT | Cluster | Peptides | Unique peptides | Sequence coverage [%] | Unique sequence coverage [%] | Mol. weight [kDa] |
|---|---|---|---|---|---|---|---|---|---|---|---|---|---|---|---|---|---|---|
| A0A0H3NHV3 | 26.8173 | 26.425 | 26.4605 | 24.8019 | 25.8062 | 25.0528 | Mannitol_dh;Mannitol_dh_C | | | 1.836551775 | 0.04988636 | -1.347300212 | | 6 | 6 | 13.9 | 13.9 | 54.01 |
| A0A0H3NEI9 | 24.6788 | 25.5763 | 25.7075 | 26.9196 | 26.6619 | 26.4682 | TP_methylase | | | 1.758545121 | 0.05017778 | 1.362363815 | | 5 | 5 | 19.5 | 19.5 | 28.38 |
| A0A0H3NHZ2 | 23.5882 | 26.1053 | 25.3094 | 27.8079 | 27.4578 | 25.7501 | BOF | | | 0.959874181 | 0.0504581 | 2.004301707 | | 2 | 2 | 15.4 | 15.4 | 13.95 |
| A0A0H3NND7 | 27.2194 | 26.6497 | 26.8687 | 26.4906 | 24.2025 | 24.8773 | Wzz | ECA polysaccharide chain length modulation protein {ECO:0000256\|HAMAP-Rule:MF_02025} | | 1.15914086 | 0.05074157 | -1.722420375 | | 9 | 9 | 31.3 | 31.3 | 39.46 |
| A0A0H3NH51 | 25.2804 | 27.5406 | 25.9359 | 27.7959 | 28.2385 | 27.7793 | Rotamase;SurA_N | Chaperone SurA {ECO:0000256\|HAMAP-Rule:MF_01183} | | 1.152139815 | 0.05083516 | 1.685596466 | | 13 | 13 | 34.3 | 34.3 | 47.25 |
| A0A0H3NGD3 | 26.3244 | 26.432 | 26.5835 | 24.9731 | 25.5749 | 25.2074 | SecG | | | 2.481254184 | 0.05111602 | -1.19482104 | | 3 | 3 | 27.3 | 27.3 | 11.39 |
| A0A0H3N916 | 25.4089 | 27.7487 | 25.9245 | 27.6281 | 28.4857 | 28.1952 | ACR_tran | Efflux pump membrane transporter {ECO:0000256\|RuleBase:RU364070} | | 1.087662039 | 0.05197826 | 1.742331823 | | 15 | 15 | 18.9 | 18.9 | 113.6 |
| A0A0H3NFX6 | 26.6009 | 26.5169 | 26.5496 | 25.373 | 25.7267 | 24.227 | Exo_endo_phos | | | 1.479546636 | 0.0522623 | -1.446868896 | | 11 | 11 | 40.7 | 40.7 | 30.79 |
| A0A0H3NIS3 | 28.4762 | 28.8353 | 28.72 | 25.849 | 24.7427 | 28.8269 | CstA;CstA_5TM | | | 0.835263918 | 0.05236757 | -2.20429039 | | 5 | 5 | 8.1 | 8.1 | 75.04 |
| A0A0H3NI37 | 26.8197 | 26.5098 | 25.1005 | 24.5093 | 24.7057 | 24.6723 | Fer4_11;Form-deh_trans | | | 1.33073023 | 0.05333333 | -1.514240901 | | 4 | 4 | 13.3 | 13.3 | 33.06 |

### All proteins

| Uniprot accession number | LFQ intensity WT_1 | LFQ intensity WT_2 | LFQ intensity WT_3 | LFQ intensity OrfSwap_1 | LFQ intensity OrfSwap_2 | LFQ intensity OrfSwap_3 | Pfam name | Uniprot full protein name | Student's T-test Significant OrfSwap_WT | -Log Student's T-test p-value OrfSwap_WT | Student's T-test q-value OrfSwap_WT | Student's T-test Difference OrfSwap_WT | Cluster | Peptides | Unique peptides | Sequence coverage [%] | Unique sequence coverage [%] | Mol. weight [kDa] |
|---|---|---|---|---|---|---|---|---|---|---|---|---|---|---|---|---|---|---|
| A0A0H3NAH4 | 27.6135 | 27.2482 | 27.3935 | 25.39 | 24.2815 | 27.1411 | 2-Hacid_dh_C | Glyoxylate/hydroxypyruvate reductase A {ECO:0000256\|HAMAP-Rule:MF_01666} | | 1.014653931 | 0.05437433 | -1.814182281 | | 7 | 7 | 28.5 | 28.5 | 35.03 |
| A0A0H3NMJ5 | 25.7902 | 24.3714 | 24.7417 | 26.3681 | 26.3862 | 26.3453 | SKI | Shikimate kinase 1 {ECO:0000256\|HAMAP-Rule:MF_00109} | | 1.520175992 | 0.05481481 | 1.39881134 | | 4 | 4 | 31.2 | 31.2 | 19.47 |
| A0A0H3NH79 | 27.3432 | 27.3772 | 27.3122 | 25.768 | 24.533 | 26.8286 | BcsB | Cyclic di-GMP-binding protein {ECO:0000256\|RuleBase:RU365021} | | 1.158167535 | 0.05510638 | -1.634312312 | | 8 | 8 | 13.3 | 13.3 | 84.3 |
| A0A0H3N9L9 | 25.9502 | 26.21 | 26.5332 | 24.6084 | 25.5554 | 24.535 | AA_kinase;PUA | Glutamate 5-kinase {ECO:0000256\|HAMAP-Rule:MF_00456} | | 1.644869319 | 0.056 | -1.33151563 | | 4 | 4 | 14.2 | 14.2 | 39.14 |
| E1WFA0 | 26.7876 | 26.6503 | 26.6451 | 25.1754 | 25.1282 | 26.0093 | HATPase_c;PhoQ_Sensor | Virulence sensor histidine kinase PhoQ | | 1.909920908 | 0.05629319 | -1.256701152 | | 6 | 6 | 12.7 | 12.7 | 55.47 |
| A0A0H3NHD2 | 25.6791 | 27.5633 | 24.8751 | 27.8395 | 27.8671 | 27.6693 | Rhodanese | | | 1.030067945 | 0.05658947 | 1.752812068 | | 5 | 5 | 34.3 | 34.3 | 15.66 |
| A0A0H3NGF5 | 26.6926 | 25.315 | 26.9092 | 25.1083 | 23.4476 | 25.2609 | | | | 1.042786334 | 0.05830052 | -1.699963888 | | 5 | 5 | 22.7 | 22.7 | 26.27 |

### All proteins

| Uniprot accession number | LFQ intensity WT_1 | LFQ intensity WT_2 | LFQ intensity WT_3 | LFQ intensity OrfSwap_1 | LFQ intensity OrfSwap_2 | LFQ intensity OrfSwap_3 | Pfam name | Uniprot full protein name | Student's T-test Significant OrfSwap_WT | -Log Student's T-test p-value OrfSwap_WT | Student's T-test q-value OrfSwap_WT | Student's T-test Difference OrfSwap_WT | Cluster | Peptides | Unique peptides | Sequence coverage [%] | Unique sequence coverage [%] | Mol. weight [kDa] |
|---|---|---|---|---|---|---|---|---|---|---|---|---|---|---|---|---|---|---|
| A0A0H3NG39 | 27.5228 | 27.7728 | 25.8774 | 25.2644 | 25.365 | 25.9215 | DHBP_synthase | 3,4-dihydroxy-2-butanone 4-phosphate synthase {ECO:0000256\|HAMAP-Rule:MF_00180, ECO:0000256\|RuleBase:RU003843} | | 1.152438787 | 0.06338144 | -1.540630341 | | 2 | 2 | 11.5 | 11.5 | 23.31 |
| A0A0H3NA44 | 23.929 | 24.4313 | 25.4923 | 25.2591 | 26.6806 | 26.5191 | Pribosyltran | Adenine phosphoribosyltransferase {ECO:0000256\|HAMAP-Rule:MF_00004, ECO:0000256\|SAAS:SAAS01090472} | | 1.121877177 | 0.06448731 | 1.535381953 | | 5 | 5 | 35.5 | 35.5 | 19.99 |
| A0A0H3NQZ5 | 24.9021 | 25.3475 | 25.9691 | 26.0378 | 27.1628 | 27.1966 | YscJ_FliF | Lipoprotein {ECO:0000256\|RuleBase:RU364102} | | 1.328612924 | 0.06481633 | 1.39279302 | | 3 | 3 | 18.3 | 18.3 | 28.21 |
| A0A0H3NLC5 | 27.5089 | 27.7272 | 27.8351 | 24.6218 | 27.7837 | 25.0786 | DeoC | Deoxyribose-phosphate aldolase {ECO:0000256\|HAMAP-Rule:MF_00592} | | 0.874546303 | 0.06514872 | -1.862347921 | | 8 | 8 | 40 | 40 | 28.37 |
| A0A0H3NGI6 | 26.5618 | 26.9846 | 26.6625 | 26.2922 | 25.1482 | 24.6088 | PDZ;PDZ_2 | Periplasmic serine endoprotease DegP-like {ECO:0000256\|RuleBase:RU364067} | | 1.26957085 | 0.06976884 | -1.386597315 | | 7 | 7 | 19.8 | 19.8 | 47.33 |

### All proteins

| Uniprot accession number | LFQ intensity WT_1 | LFQ intensity WT_2 | LFQ intensity WT_3 | LFQ intensity OrfSwap_1 | LFQ intensity OrfSwap_2 | LFQ intensity OrfSwap_3 | Pfam name | Uniprot full protein name | Student's T-test Significant OrfSwap_WT | -Log Student's T-test p-value OrfSwap_WT | Student's T-test q-value OrfSwap_WT | Student's T-test Difference OrfSwap_WT | Cluster | Peptides | Unique peptides | Sequence coverage [%] | Unique sequence coverage [%] | Mol. weight [kDa] |
|---|---|---|---|---|---|---|---|---|---|---|---|---|---|---|---|---|---|---|
| A0A0H3NBS8 | 25.6261 | 24.9622 | 24.0856 | 25.315 | 27.0697 | 27.0594 | Aldedh | Gamma-aminobutyraldehyde dehydrogenase {ECO:0000256\|HAMAP-Rule:MF_01275} | | 1.016403918 | 0.07012121 | 1.590040843 | | 11 | 11 | 30.6 | 30.6 | 52.04 |
| A0A0H3NFF6 | 24.0395 | 25.7653 | 24.6047 | 24.9966 | 27.5487 | 27.1973 | LprI | | | 0.87433564 | 0.07058 | 1.777692159 | | 4 | 4 | 24.4 | 24.4 | 14.13 |
| A0A0H3NI34 | 24.7702 | 27.1225 | 26.9035 | 25.258 | 24.092 | 24.4808 | Aminotran_3 | Putrescine aminotransferase {ECO:0000256\|HAMAP-Rule:MF_01276} | | 0.938377052 | 0.07241791 | -1.655134837 | | 7 | 7 | 19.3 | 19.3 | 46.47 |
| A0A0H3NDP0 | 24.9765 | 26.1346 | 26.2062 | 24.8744 | 24.237 | 24.2766 | Acetyltransf_10 | | | 1.363712217 | 0.07259406 | -1.30973498 | | 2 | 2 | 15.7 | 15.7 | 17.45 |
| A0A0H3N9I0 | 24.6774 | 24.8651 | 25.6271 | 27.7416 | 24.937 | 27.7808 | Glutaredoxin | | | 0.829749407 | 0.07509804 | 1.763289769 | | 3 | 3 | 47.1 | 47.1 | 9.924 |
| A0A0H3N8Q7 | 26.4622 | 26.3468 | 26.4897 | 25.7533 | 25.3262 | 24.8466 | H2O2_YaaD | UPF0246 protein YaaA {ECO:0000256\|HAMAP-Rule:MF_00652} | | 1.875276959 | 0.07546798 | -1.124176661 | | 7 | 7 | 31.1 | 31.1 | 29.76 |
| A0A0H3NEM8 | 27.072 | 26.8661 | 26.9237 | 26.4037 | 25.725 | 25.0528 | SpoU_methylase;SpoU_sub_bind | | | 1.444219014 | 0.07939806 | -1.226771673 | | 7 | 7 | 23.8 | 23.8 | 38.08 |
| A0A0H3NKF4 | 25.2937 | 25.2508 | 24.9276 | 25.9302 | 26.1781 | 26.6182 | Fer2 | | | 2.023324884 | 0.07978537 | 1.084794998 | | 3 | 3 | 29.7 | 29.7 | 12.39 |
| A0A0H3NCC6 | 29.0604 | 28.105 | 24.992 | 24.4738 | 24.9679 | 26.5112 | MipA | | | 0.685173477 | 0.08144231 | -2.068143845 | | 3 | 3 | 13.3 | 13.3 | 27.99 |
| A0A0H3NAF5 | 26.0678 | 25.972 | 25.9988 | 25.2977 | 25.0006 | 24.5595 | Ricin_B_lectin | | | 2.094949477 | 0.08183575 | -1.060272217 | | 3 | 3 | 20.8 | 20.8 | 19.06 |
| A0A0H3NAX0 | 23.9114 | 24.6001 | 25.2838 | 25.2356 | 26.271 | 26.1589 | Aldose_epim | Aldose 1-epimerase {ECO:0000256\|PIRNR:PIRNR005096} | | 1.179130743 | 0.08888038 | 1.290044785 | | 7 | 7 | 22 | 22 | 38.55 |
| A0A0H3NG37 | 27.719 | 25.0242 | 27.6878 | 25.213 | 25.177 | 25.2385 | P-II | | | 0.83091311 | 0.09219048 | -1.600828807 | | 5 | 5 | 39.3 | 39.3 | 12.43 |

### All proteins

| Uniprot accession number | LFQ intensity WT_1 | LFQ intensity WT_2 | LFQ intensity WT_3 | LFQ intensity OrfSwap_1 | LFQ intensity OrfSwap_2 | LFQ intensity OrfSwap_3 | Pfam name | Uniprot full protein name | Student's T-test Significant OrfSwap_WT | -Log Student's T-test p-value OrfSwap_WT | Student's T-test q-value OrfSwap_WT | Student's T-test Difference OrfSwap_WT | Cluster | Peptides | Unique peptides | Sequence coverage [%] | Unique sequence coverage [%] | Mol. weight [kDa] |
|---|---|---|---|---|---|---|---|---|---|---|---|---|---|---|---|---|---|---|
| A0A0H3NPX0 | 23.5252 | 24.0426 | 24.9996 | 25.5294 | 25.5082 | 25.1831 | | | | 1.279967052 | 0.09277725 | 1.217734655 | | 7 | 7 | 6 | 6 | 139.2 |
| A0A0H3NAG4 | 25.3048 | 27.538 | 24.2249 | 25.5929 | 28.6173 | 29.003 | YccJ | | | 0.635155329 | 0.09293023 | 2.048512141 | | 7 | 7 | 80 | 80 | 8.69 |
| A0A0H3NCZ8 | 25.3782 | 25.0074 | 25.609 | 27.6286 | 25.8932 | 26.3666 | BMC | | | 1.116330371 | 0.09293458 | 1.297911962 | | 4 | 4 | 62.6 | 62.6 | 9.067 |
| A0A0H3NNJ2 | 25.2405 | 26.2008 | 26.1406 | 25.2175 | 24.6162 | 24.1032 | FAD_binding_6;NAD_binding_1 | | | 1.274011335 | 0.09337089 | -1.21498998 | | 8 | 8 | 42.9 | 42.9 | 26.35 |
| A0A0H3NEC5 | 25.0444 | 24.9369 | 24.8065 | 25.9787 | 26.1516 | 25.6015 | KH_dom-like;MMR_HSR1 | GTPase Der {ECO:0000256\|HAMAP-Rule:MF_00195, ECO:0000256\|RuleBase:RU004481, ECO:0000256\|SAAS:SAAS00336749} | | 2.291418001 | 0.09359259 | 0.981352488 | | 6 | 6 | 13.7 | 13.7 | 54.97 |
| A0A0H3NBL2 | 25.1873 | 24.5874 | 24.0586 | 25.8654 | 25.6318 | 25.668 | MukE | Chromosome partition protein MukE {ECO:0000256\|HAMAP-Rule:MF_01802} | | 1.533935294 | 0.09365899 | 1.110596975 | | 5 | 5 | 21.8 | 21.8 | 26.97 |
| A0A0H3NDN0 | 29.9278 | 29.7352 | 29.3405 | 25.0557 | 28.9076 | 29.1525 | COX_ARM;COX2 | Ubiquinol oxidase subunit 2 {ECO:0000256\|PIRNR:PIRNR000292} | | 0.664955105 | 0.09381132 | -1.962561925 | | 11 | 11 | 45.6 | 45.6 | 35.29 |
| A0A0H3NFV8 | 25.2448 | 25.4323 | 25.2348 | 24.8338 | 24.099 | 23.6225 | Aldo_ket_red | | | 1.45129366 | 0.09666055 | -1.118873596 | | 3 | 3 | 11.6 | 11.6 | 31.33 |
| A0A0H3NGG5 | 28.6111 | 28.8405 | 28.7186 | 25.6391 | 27.5374 | 28.4677 | GrpE | Protein GrpE {ECO:0000256\|HAMAP-Rule:MF_01151, ECO:0000256\|RuleBase:RU000639} | | 0.838386761 | 0.09722374 | -1.508622487 | | 5 | 5 | 19.9 | 19.9 | 21.84 |

### All proteins

| Uniprot accession number | LFQ intensity WT_1 | LFQ intensity WT_2 | LFQ intensity WT_3 | LFQ intensity OrfSwap _1 | LFQ intensity OrfSwap _2 | LFQ intensity OrfSwap _3 | Pfam name | Uniprot full protein name | Student's T-test Significant OrfSwap_WT | -Log Student's T-test p-value OrfSwap_WT | Student's T-test q-value OrfSwap_WT | Student's T-test Difference OrfSwap_WT | Cluster | Peptides | Unique peptides | Sequence coverage [%] | Unique sequence coverage [%] | Mol. weight [kDa] |
|---|---|---|---|---|---|---|---|---|---|---|---|---|---|---|---|---|---|---|
| A0A0H3NL38 | 24.3525 | 25.2378 | 24.9796 | 26.0662 | 25.6579 | 25.9567 | A_deaminase | Adenosine deaminase {ECO:0000256\|HAMAP-Rule:MF_00540, ECO:0000256\|SAAS:SAAS00331568} | | 1.634343821 | 0.10649091 | 1.036942164 | | 8 | 8 | 22.8 | 22.8 | 36.24 |
| A0A0H3NKP1 | 25.8969 | 25.9684 | 26.2434 | 24.5742 | 24.5729 | 25.658 | TetR_N | | | 1.365799178 | 0.1070852 | -1.101199468 | | 6 | 6 | 29.3 | 29.3 | 21.83 |
| A0A0H3N9V6 | 25.2582 | 25.9312 | 26.3435 | 24.8907 | 24.5466 | 24.882 | Glycos_trans_3N | | | 1.478006091 | 0.10749321 | -1.071224213 | | 5 | 5 | 25 | 25 | 35.65 |
| A0A0H3NF23 | 25.812 | 26.311 | 25.9524 | 25.6992 | 24.4143 | 24.5015 | ABC_tran;CBS | | | 1.230388482 | 0.10756757 | -1.153465907 | | 4 | 4 | 11 | 11 | 44.12 |
| A0A0H3NP03 | 23.9442 | 24.8931 | 25.2161 | 24.3016 | 27.4059 | 27.287 | Transket_pyr;Transketolase_C | | | 0.691238875 | 0.10918584 | 1.647018433 | | 5 | 5 | 16.7 | 16.7 | 34.56 |
| A0A0H3NGC2 | 25.3624 | 27.1144 | 27.1787 | 25.2026 | 24.9158 | 25.6629 | Oxidored_FMN | | | 0.954111745 | 0.10967111 | -1.291426977 | | 8 | 8 | 37 | 37 | 39.48 |
| A0A0H3NHJ2 | 25.4319 | 27.4076 | 27.498 | 25.5586 | 25.5687 | 25.2066 | AA_kinase | Uridylate kinase {ECO:0000256\|HAMAP-Rule:MF_01220} | | 0.909990268 | 0.11016071 | -1.33453687 | | 7 | 7 | 32 | 32 | 25.96 |
| A0A0H3NB20 | 26.4279 | 26.4715 | 26.3045 | 25.354 | 24.1843 | 26.0392 | Kinase-PPPase | Phosphoenolpyruvate synthase regulatory protein {ECO:0000256\|HAMAP-Rule:MF_01062} | | 1.04416986 | 0.11221145 | -1.208778381 | | 5 | 5 | 21.3 | 21.3 | 31.15 |
| A0A0H3NM81 | 27.6353 | 27.3551 | 25.0688 | 24.0035 | 25.2921 | 26.055 | Peptidase_M48 | Protease HtpX {ECO:0000256\|HAMAP-Rule:MF_00188} | | 0.709998832 | 0.11275439 | -1.569513321 | | 6 | 6 | 16.7 | 16.7 | 31.88 |

### All proteins

| Uniprot accession number | LFQ intensity WT_1 | LFQ intensity WT_2 | LFQ intensity WT_3 | LFQ intensity OrfSwap_1 | LFQ intensity OrfSwap_2 | LFQ intensity OrfSwap_3 | Pfam name | Uniprot full protein name | Student's T-test Significant OrfSwap_WT | -Log Student's T-test p-value OrfSwap_WT | Student's T-test q-value OrfSwap_WT | Student's T-test Difference OrfSwap_WT | Cluster | Peptides | Unique peptides | Sequence coverage [%] | Unique sequence coverage [%] | Mol. weight [kDa] |
|---|---|---|---|---|---|---|---|---|---|---|---|---|---|---|---|---|---|---|
| A0A0H3NML0 | 27.3238 | 25.4307 | 27.3237 | 26.0313 | 24.9017 | 25.1695 | YscJ_FliF;YscJ_FliF_C | Flagellar M-ring protein {ECO:0000256\|PIRNR:PIRNR004862} | | 0.859090417 | 0.11476522 | -1.325234095 | | 9 | 9 | 15.7 | 15.7 | 61.23 |
| A0A0H3NAP4 | 27.8525 | 28.2403 | 28.0427 | 26.1486 | 28.2544 | 25.5622 | HIT | | | 0.77658117 | 0.11991342 | -1.39013354 | | 5 | 5 | 29.4 | 29.4 | 13.27 |
| A0A0H3NIY9 | 26.0898 | 26.8666 | 24.7182 | 27.1071 | 27.0597 | 27.2228 | DcuA_DcuB | | | 0.918157384 | 0.12081034 | 1.238332748 | | 5 | 5 | 5.6 | 5.6 | 42.03 |
| A0A0H3NWK5 | 26.8614 | 25.3274 | 26.8547 | 25.1291 | 25.3041 | 25.174 | GyrI-like;HTH_18 | | | 1.049224949 | 0.12135622 | -1.14544932 | | 5 | 5 | 19.7 | 19.7 | 33.24 |
| A0A0H3NHB2 | 26.1666 | 26.0617 | 26.2191 | 25.0345 | 25.6305 | 25.1005 | Gly-zipper_Omp;OmpA | | | 1.999578692 | 0.12596581 | -0.893984477 | | 3 | 3 | 23.9 | 23.9 | 21.08 |
| A0A0H3NTI1 | 25.4874 | 26.3865 | 25.6931 | 25.3866 | 23.7789 | 24.9295 | 2-Hacid_dh;2-Hacid_dh_C | Glyoxylate/hydroxypyruvate reductase B {ECO:0000256\|HAMAP-Rule:MF_01667, ECO:0000256\|SAAS:SAAS01081393} | | 0.986121013 | 0.12634894 | -1.157340368 | | 6 | 6 | 14.8 | 14.8 | 35.25 |
| A0A0H3NUA0 | 30.192 | 29.519 | 29.3181 | 29.5925 | 28.8917 | 25.23 | DUF413 | | | 0.571865649 | 0.12686441 | -1.771619161 | | 8 | 8 | 49.1 | 49.1 | 13.08 |
| A0A0H3NI74 | 24.8199 | 24.8297 | 25.6486 | 26.8688 | 25.3003 | 26.5902 | ATP-cone | Transcriptional repressor NrdR {ECO:0000256\|HAMAP-Rule:MF_00440, ECO:0000256\|SAAS:SAAS01000175} | | 0.972618268 | 0.12827004 | 1.153696696 | | 5 | 5 | 32.9 | 32.9 | 17.2 |
| A0A0H3NB92 | 24.5152 | 25.4415 | 24.8207 | 26.4362 | 26.4865 | 25.1833 | | | | 1.029798292 | 0.12994958 | 1.109532674 | | 4 | 4 | 13.1 | 13.1 | 43.94 |

### All proteins

| Uniprot accession number | LFQ intensity WT_1 | LFQ intensity WT_2 | LFQ intensity WT_3 | LFQ intensity OrfSwap_1 | LFQ intensity OrfSwap_2 | LFQ intensity OrfSwap_3 | Pfam name | Uniprot full protein name | Student's T-test Significant OrfSwap_WT | -Log Student's T-test p-value OrfSwap_WT | Student's T-test q-value OrfSwap_WT | Student's T-test Difference OrfSwap_WT | Cluster | Peptides | Unique peptides | Sequence coverage [%] | Unique sequence coverage [%] | Mol. weight [kDa] |
|---|---|---|---|---|---|---|---|---|---|---|---|---|---|---|---|---|---|---|
| A0A0H3NDL2 | 24.9544 | 24.6555 | 24.481 | 26.8611 | 27.0398 | 24.3253 | NusB | Transcription antitermination protein NusB {ECO:0000256\|HAMAP-Rule:MF_00073, ECO:0000256\|SAAS:SAAS01079076} | | 0.709284942 | 0.13077178 | 1.378436406 | | 3 | 3 | 21.6 | 21.6 | 15.69 |
| A0A0H3NCH8 | 27.8583 | 25.1836 | 27.165 | 24.9595 | 25.4713 | 25.7506 | Phage_int_SAM_1 | | | 0.737546473 | 0.13131667 | -1.341859182 | | 4 | 4 | 30.6 | 30.6 | 21.69 |
| A0A0H3NGH3 | 27.0099 | 26.2922 | 26.7269 | 25.5173 | 25.5025 | 26.1549 | DUF386 | | | 1.472379375 | 0.13134728 | -0.951416016 | | 2 | 2 | 21.3 | 21.3 | 17.06 |
| A0A0H3NGU5 | 26.2738 | 26.2885 | 26.2957 | 23.4781 | 26.5363 | 24.7926 | Glu_cys_ligase | Glutamate--cysteine ligase {ECO:0000256\|HAMAP-Rule:MF_00578} | | 0.694501774 | 0.13466667 | -1.350312551 | | 6 | 6 | 16.6 | 16.6 | 58.38 |
| A0A0H3NDD3 | 27.3901 | 24.816 | 27.1473 | 24.266 | 26.6504 | 23.6384 | EFP;EFP_N;Elong-fact-P_C | Elongation factor P-like protein {ECO:0000256\|HAMAP-Rule:MF_00646, ECO:0000256\|SAAS:SAAS01087250} | | 0.579058447 | 0.13522314 | -1.599573135 | | 4 | 4 | 22.6 | 22.6 | 21.39 |
| A0A0H3NS81 | 24.4772 | 25.0188 | 25.517 | 26.044 | 25.7148 | 26.0803 | LppC | Penicillin-binding protein activator LpoA {ECO:0000256\|HAMAP-Rule:MF_01890} | | 1.366583025 | 0.13580408 | 0.942056656 | | 5 | 5 | 9.7 | 9.7 | 72.62 |

### All proteins

| Uniprot accession number | LFQ intensity WT_1 | LFQ intensity WT_2 | LFQ intensity WT_3 | LFQ intensity OrfSwap_1 | LFQ intensity OrfSwap_2 | LFQ intensity OrfSwap_3 | Pfam name | Uniprot full protein name | Student's T-test Significant OrfSwap_WT | -Log Student's T-test p-value OrfSwap_WT | Student's T-test q-value OrfSwap_WT | Student's T-test Difference OrfSwap_WT | Cluster | Peptides | Unique peptides | Sequence coverage [%] | Unique sequence coverage [%] | Mol. weight [kDa] |
|---|---|---|---|---|---|---|---|---|---|---|---|---|---|---|---|---|---|---|
| A0A0H3NDQ1 | 26.26 | 24.63 | 26.8721 | 25.0983 | 23.5647 | 25.146 | NADHdh | NADH-quinone oxidoreductase subunit H {ECO:0000256\|HAMAP-Rule:MF_01350} | | 0.710513958 | 0.13585246 | -1.31771342 | | 2 | 2 | 2.8 | 2.8 | 36.29 |
| A0A0H3NCV5 | 27.2081 | 25.157 | 26.2146 | 25.6843 | 24.0202 | 25.1166 | Asp_decarbox | Aspartate 1-decarboxylase {ECO:0000256\|HAMAP-Rule:MF_00446} | | 0.749611234 | 0.13721951 | -1.252854665 | | 3 | 3 | 19.8 | 19.8 | 13.89 |
| A0A0H3ND61 | 26.7003 | 25.6722 | 26.687 | 24.0431 | 24.8191 | 26.4373 | TP_methylase | | | 0.733395063 | 0.14106883 | -1.253341039 | | 2 | 2 | 7.9 | 7.9 | 25.87 |
| A0A0H3N990 | 25.7206 | 25.2827 | 25.2016 | 24.2554 | 24.9324 | 24.349 | MS_channel | | | 1.540662066 | 0.14151613 | -0.889360428 | | 2 | 2 | 6.6 | 6.6 | 42.31 |
| A0A0H3NAQ7 | 27.1126 | 25.0749 | 24.1672 | 27.195 | 27.0118 | 26.0507 | Flavodoxin_1 | Flavodoxin {ECO:0000256\|PIRNR:PIRNR038996} | | 0.622082989 | 0.16963855 | 1.300951004 | | 2 | 2 | 18.8 | 18.8 | 19.64 |
| A0A0H3NAK0 | 25.8801 | 25.6132 | 26.0892 | 25.1462 | 23.7391 | 25.5558 | DAO | N-methyl-L-tryptophan oxidase {ECO:0000256\|HAMAP-Rule:MF_00515} | | 0.858328951 | 0.17096 | -1.047143936 | | 4 | 4 | 18.5 | 18.5 | 40.66 |
| A0A0H3NE79 | 26.5038 | 26.4062 | 26.5533 | 26.0763 | 25.3978 | 25.5755 | PhoH | | | 1.746167461 | 0.17185714 | -0.804548264 | | 6 | 6 | 14.7 | 14.7 | 40.96 |
| A0A0H3NRA3 | 24.7409 | 27.6063 | 28.2429 | 27.676 | 28.404 | 28.7439 | DUF903 | | | 0.557624721 | 0.17209486 | 1.411263784 | | 6 | 6 | 56 | 56 | 8.126 |
| A0A0H3NKM0 | 24.6917 | 24.078 | 24.9324 | 25.6189 | 25.0303 | 25.7068 | | | | 1.253144021 | 0.17240945 | 0.884614944 | | 2 | 2 | 27 | 27 | 7.61 |

### All proteins

| Uniprot accession number | LFQ intensity WT_1 | LFQ intensity WT_2 | LFQ intensity WT_3 | LFQ intensity OrfSwap_1 | LFQ intensity OrfSwap_2 | LFQ intensity OrfSwap_3 | Pfam name | Uniprot full protein name | Student's T-test Significant OrfSwap_WT | -Log Student's T-test p-value OrfSwap_WT | Student's T-test q-value OrfSwap_WT | Student's T-test Difference OrfSwap_WT | Cluster | Peptides | Unique peptides | Sequence coverage [%] | Unique sequence coverage [%] | Mol. weight [kDa] |
|---|---|---|---|---|---|---|---|---|---|---|---|---|---|---|---|---|---|---|
| A0A0H3N8B1 | 28.5596 | 28.7411 | 28.974 | 25.6764 | 28.4123 | 28.3644 | DMRL_synthase | 6,7-dimethyl-8-ribityllumazine synthase {ECO:0000256\|HAMAP-Rule:MF_00178, ECO:0000256\|SAAS:SAAS01078290} | | 0.628920999 | 0.17254183 | -1.273829142 | | 5 | 5 | 38.5 | 38.5 | 16.01 |
| A0A0H3NNS0 | 26.6988 | 27.4408 | 23.9673 | 25.2401 | 24.096 | 24.6562 | GerE;Response_reg | Transcriptional regulatory protein RcsB {ECO:0000256\|HAMAP-Rule:MF_00981} | | 0.548301016 | 0.1841098 | -1.371500015 | | 8 | 8 | 33.8 | 33.8 | 23.71 |
| A0A0H3NDS8 | 28.0357 | 28.364 | 28.1628 | 27.6603 | 27.9601 | 25.5465 | Epimerase;Formyl_trans_C;Formyl_trans_N | Bifunctional polymyxin resistance protein ArnA {ECO:0000256\|HAMAP-Rule:MF_01166} | | 0.670960696 | 0.19351563 | -1.131861369 | | 13 | 13 | 25.9 | 25.9 | 73.59 |
| A0A0H3ND22 | 26.4761 | 26.0732 | 25.5251 | 24.2578 | 24.1748 | 26.2924 | Glycos_transf_2 | | | 0.680954482 | 0.19410117 | -1.116469701 | | 2 | 2 | 9.9 | 9.9 | 35.51 |
| A0A0H3NP94 | 28.1435 | 28.1915 | 28.0136 | 27.8861 | 27.6335 | 25.3902 | Acid_phosphat_B | Class B acid phosphatase {ECO:0000256\|PIRNR:PIRNR017818} | | 0.652092393 | 0.19468217 | -1.146260579 | | 5 | 5 | 27.8 | 27.8 | 26.31 |
| E1WAB4 | 27.2753 | 26.7108 | 26.9507 | 27.9331 | 24.6549 | 24.238 | | Oxygen-regulated invasion protein OrgB | | 0.508580244 | 0.20447876 | -1.370319366 | | 6 | 6 | 30.5 | 30.5 | 26.46 |

### All proteins

| Uniprot accession number | LFQ intensity WT_1 | LFQ intensity WT_2 | LFQ intensity WT_3 | LFQ intensity OrfSwap_1 | LFQ intensity OrfSwap_2 | LFQ intensity OrfSwap_3 | Pfam name | Uniprot full protein name | Student's T-test Significant OrfSwap_WT | -Log Student's T-test p-value OrfSwap_WT | Student's T-test q-value OrfSwap_WT | Student's T-test Difference OrfSwap_WT | Cluster | Peptides | Unique peptides | Sequence coverage [%] | Unique sequence coverage [%] | Mol. weight [kDa] |
|---|---|---|---|---|---|---|---|---|---|---|---|---|---|---|---|---|---|---|
| A0A0H3N804 | 26.798 | 26.3069 | 26.5577 | 26.6355 | 24.3896 | 25.5156 | Aldedh | Gamma-glutamyl phosphate reductase {ECO:0000256\|HAMAP-Rule:MF_00412} | | 0.716823703 | 0.20611494 | -1.040612539 | | 6 | 6 | 15.6 | 15.6 | 44.67 |
| A0A0H3NFW4 | 25.3982 | 25.1823 | 24.6932 | 25.311 | 26.1104 | 26.4916 | DUF469 | | | 1.018194332 | 0.20643077 | 0.879769007 | | 2 | 2 | 10.2 | 10.2 | 12.79 |
| A0A0H3NVS5 | 24.6256 | 31.7777 | 31.726 | 31.1942 | 31.672 | 31.5164 | Entericidin | | | 0.366017145 | 0.2138626 | 2.084419886 | | 1 | 1 | 39.6 | 39.6 | 4.811 |
| A0A0H3NQ63 | 25.578 | 25.996 | 25.6215 | 25.6254 | 24.1891 | 24.7301 | PRC;RimM | Ribosome maturation factor RimM {ECO:0000256\|HAMAP-Rule:MF_00014, ECO:0000256\|SAAS:SAAS00059784} | | 0.940688282 | 0.21450951 | -0.883623123 | | 2 | 2 | 10.4 | 10.4 | 20.47 |
| A0A0H3NIH1 | 24.1757 | 25.8295 | 24.8528 | 27.2459 | 26.5196 | 24.61 | MotA_ExbB | | | 0.565506485 | 0.21731818 | 1.172527313 | | 7 | 7 | 21 | 21 | 51.46 |
| A0A0H3NFC9 | 26.5628 | 24.635 | 27.2068 | 24.3334 | 25.8418 | 24.8182 | Transketolase_N | | | 0.566795133 | 0.22676692 | -1.137069066 | | 5 | 5 | 35.1 | 35.1 | 30.4 |
| A0A0H3NGW1 | 25.4597 | 24.7244 | 24.8495 | 26.5551 | 26.4944 | 24.8712 | Ribul_P_3_epim | Ribulose-phosphate 3-epimerase {ECO:0000256\|PIRNR:PIRNR001461} | | 0.740115895 | 0.22715472 | 0.96235466 | | 6 | 6 | 21.3 | 21.3 | 24.49 |
| A0A0H3NAQ1 | 28.3407 | 28.3928 | 28.3998 | 28.4462 | 23.8661 | 28.3394 | Glucosamine_iso | Glucosamine-6-phosphate deaminase {ECO:0000256\|HAMAP-Rule:MF_01241} | | 0.422136795 | 0.22963296 | -1.493870417 | | 12 | 12 | 54.1 | 54.1 | 29.63 |
| A0A0H3NTQ2 | 26.8501 | 25.1944 | 26.5695 | 24.8445 | 25.8205 | 25.1294 | Glyco_transf_9 | | | 0.73247117 | 0.23508955 | -0.939876556 | | 3 | 3 | 8.3 | 8.3 | 39.02 |

#### All proteins

| Uniprot accession number | LFQ intensity WT_1 | LFQ intensity WT_2 | LFQ intensity WT_3 | LFQ intensity OrfSwap_1 | LFQ intensity OrfSwap_2 | LFQ intensity OrfSwap_3 | Pfam name | Uniprot full protein name | Student's T-test Significant OrfSwap_WT | -Log Student's T-test p-value OrfSwap_WT | Student's T-test q-value OrfSwap_WT | Student's T-test Difference OrfSwap_WT | Cluster | Peptides | Unique peptides | Sequence coverage [%] | Unique sequence coverage [%] | Mol. weight [kDa] |
|---|---|---|---|---|---|---|---|---|---|---|---|---|---|---|---|---|---|---|
| A0A0H3NCF0 | 27.5304 | 27.9265 | 27.8633 | 24.9893 | 27.2458 | 27.8022 | Aldedh | N-succinylglutamate 5-semialdehyde dehydrogenase {ECO:0000256\|HAMAP-Rule:MF_01174} | | 0.558625973 | 0.24312268 | -1.094291687 | | 9 | 9 | 20.9 | 20.9 | 53.02 |
| A0A0H3NBQ5 | 25.2713 | 26.5377 | 26.1795 | 25.8532 | 24.6958 | 24.8239 | Molybdopterin;Molydop_binding;Nitr_red_alph_N | | | 0.763237308 | 0.2625037 | -0.871847788 | | 10 | 7 | 10.5 | 7.2 | 140.1 |
| A0A0H3NDL3 | 26.1206 | 26.5154 | 26.8377 | 24.8649 | 25.5635 | 26.4674 | LolB | Outer-membrane lipoprotein LolB {ECO:0000256\|HAMAP-Rule:MF_00233, ECO:0000256\|SAAS:SAAS00846922} | | 0.779762633 | 0.26515129 | -0.85933876 | | 6 | 6 | 26.6 | 26.6 | 23.69 |
| A0A0H3NLT0 | 24.5264 | 26.29 | 26.0654 | 24.1188 | 24.6848 | 25.2754 | Aldo_ket_red | | | 0.653099301 | 0.26892647 | -0.934234619 | | 3 | 3 | 11.6 | 11.6 | 38.56 |
| A0A0H3NPC7 | 26.8417 | 24.4158 | 26.8474 | 24.7782 | 23.771 | 26.0782 | Dyp_perox | | | 0.479728625 | 0.27107692 | -1.159184138 | | 9 | 9 | 49.5 | 49.5 | 33.05 |
| E1WAC8 | 31.0765 | 31.5064 | 32.1395 | 32.6575 | 32.5755 | 31.8486 | IpaC_SipC | Cell invasion protein SipC | | 0.913960106 | 0.27680292 | 0.786367416 | | 39 | 39 | 73.8 | 73.8 | 42.98 |
| A0A0H3NEQ4 | 27.6463 | 27.6256 | 24.3341 | 25.2324 | 26.0564 | 24.679 | SRP_SPB;SRP54;SRP54_N | Signal recognition particle protein {ECO:0000256\|HAMAP-Rule:MF_00306} | | 0.444999124 | 0.27775362 | -1.21272405 | | 7 | 7 | 19.9 | 19.9 | 49.8 |
| A0A0H3NLI8 | 25.9622 | 25.9979 | 26.1512 | 25.3924 | 25.7746 | 24.7626 | | | | 1.137347439 | 0.27797818 | -0.727197011 | | 5 | 5 | 27.7 | 27.7 | 24.37 |

### All proteins

| Uniprot accession number | LFQ intensity WT_1 | LFQ intensity WT_2 | LFQ intensity WT_3 | LFQ intensity OrfSwap_1 | LFQ intensity OrfSwap_2 | LFQ intensity OrfSwap_3 | Pfam name | Uniprot full protein name | Student's T-test Significant OrfSwap_WT | -Log Student's T-test p-value OrfSwap_WT | Student's T-test q-value OrfSwap_WT | Student's T-test Difference OrfSwap_WT | Cluster | Peptides | Unique peptides | Sequence coverage [%] | Unique sequence coverage [%] | Mol. weight [kDa] |
|---|---|---|---|---|---|---|---|---|---|---|---|---|---|---|---|---|---|---|
| A0A0H3NYF3 | 29.0904 | 29.1882 | 29.0813 | 28.62 | 28.5022 | 28.4775 | ParB;ParBc | | | 3.337119336 | 0.28036101 | -0.586761475 | | 14 | 14 | 45.2 | 45.2 | 36.85 |
| A0A0H3NMG4 | 33.0265 | 32.7985 | 32.8373 | 33.5773 | 33.4708 | 33.4116 | EFG_C;EFG_IV;GTP_EFTU;GTP_EFTU_D2 | Elongation factor G {ECO:0000256\|HAMAP-Rule:MF_00054} | | 2.661029644 | 0.28078014 | 0.599168142 | | 54 | 54 | 74.6 | 74.6 | 77.6 |
| A0A0H3ND42 | 26.3014 | 26.2693 | 26.2054 | 25.3254 | 25.515 | 25.9566 | Peptidase_M90 | Protein MtfA {ECO:0000256\|HAMAP-Rule:MF_01593} | | 1.599471196 | 0.28097842 | -0.659721375 | | 3 | 3 | 15.5 | 15.5 | 30.19 |
| A0A0H3NIB3 | 32.3545 | 32.4306 | 32.3804 | 31.8094 | 31.8131 | 31.8445 | RNA_pol_Rpb2_1;RNA_pol_Rpb2_2;RNA_pol_Rpb2_3;RNA_pol_Rpb2_45;RNA_pol_Rpb2_6;RNA_pol_Rpb2_7 | DNA-directed RNA polymerase subunit beta {ECO:0000256\|HAMAP-Rule:MF_01321, ECO:0000256\|RuleBase:RU363031} | | 4.650347385 | 0.28143772 | -0.566162109 | | 75 | 75 | 47.5 | 47.5 | 150.6 |
| A0A0H3NJI3 | 30.187 | 30.0026 | 30.0262 | 29.047 | 29.6658 | 29.5306 | DUF3828 | | | 1.541902641 | 0.28193684 | -0.657442729 | | 10 | 10 | 61.4 | 61.4 | 18.81 |
| A0A0H3NM35 | 31.4863 | 31.4237 | 31.3807 | 30.9586 | 30.8019 | 30.7604 | Ribosomal_S21 | 30S ribosomal protein S21 {ECO:0000256\|HAMAP-Rule:MF_00358, ECO:0000256\|RuleBase:RU000667} | | 3.020423703 | 0.28244286 | -0.589903514 | | 11 | 11 | 59.2 | 59.2 | 8.5 |
| A0A0H3NRJ9 | 27.5995 | 27.8001 | 24.9765 | 27.8061 | 27.8597 | 27.8589 | Rib_5-P_isom_A | Ribose-5-phosphate isomerase A {ECO:0000256\|HAMAP-Rule:MF_00170} | | 0.504579253 | 0.28292958 | 1.049494425 | | 6 | 6 | 24.2 | 24.2 | 22.9 |

### All proteins

| Uniprot accession number | LFQ intensity WT_1 | LFQ intensity WT_2 | LFQ intensity WT_3 | LFQ intensity OrfSwap_1 | LFQ intensity OrfSwap_2 | LFQ intensity OrfSwap_3 | Pfam name | Uniprot full protein name | Student's T-test Significant OrfSwap_WT | -Log Student's T-test p-value OrfSwap_WT | Student's T-test q-value OrfSwap_WT | Student's T-test Difference OrfSwap_WT | Cluster | Peptides | Unique peptides | Sequence coverage [%] | Unique sequence coverage [%] | Mol. weight [kDa] |
|---|---|---|---|---|---|---|---|---|---|---|---|---|---|---|---|---|---|---|
| A0A0H3NME1 | 32.5294 | 32.8416 | 32.7327 | 32.2579 | 32.035 | 31.9434 | RNA_pol_A_bac;RNA_pol_A_CTD;RNA_pol_L | DNA-directed RNA polymerase subunit alpha {ECO:0000256\|HAMAP-Rule:MF_00059} | | 2.050779798 | 0.28310954 | -0.622455597 | | 23 | 23 | 66.9 | 66.9 | 36.51 |
| A0A0H3NNT3 | 29.8086 | 29.6056 | 29.9577 | 29.4015 | 29.1621 | 28.8234 | FAD_binding_2 | Anaerobic glycerol-3-phosphate dehydrogenase subunit B {ECO:0000256\|HAMAP-Rule:MF_00753} | | 1.552325445 | 0.28311111 | -0.661644618 | | 9 | 9 | 22 | 22 | 45.67 |
| A0A0H3N9Z7 | 25.6854 | 27.3205 | 27.0576 | 27.2205 | 27.7599 | 27.5963 | Aldo_ket_red | | | 0.72098797 | 0.28497561 | 0.8377285 | | 7 | 7 | 24.1 | 24.1 | 36.16 |
| E1WGF0 | 27.6665 | 27.7212 | 27.4428 | 28.2598 | 28.2034 | 28.1569 | Arg_tRNA_synt_N;DALR_1;tRNA-synt_1d | Arginine--tRNA ligase | | 2.566925075 | 0.28553846 | 0.596537272 | | 14 | 14 | 22.2 | 22.2 | 64.28 |
| A0A0H3N9U1 | 25.8458 | 26.0013 | 25.0543 | 25.7115 | 23.7959 | 24.7448 | Bax1-I | | | 0.636154868 | 0.28953633 | -0.883050283 | | 2 | 2 | 7.3 | 7.3 | 25.94 |
| A0A0H3NEI6 | 26.6611 | 24.3276 | 26.6225 | 24.5827 | 24.9061 | 25.2018 | TP_methylase | | | 0.543166523 | 0.29054167 | -0.973559062 | | 5 | 5 | 31.6 | 31.6 | 25.81 |
| A0A0H3NE63 | 28.9959 | 29.0618 | 28.8589 | 28.5998 | 28.2199 | 28.2521 | Anticodon_1;tRNA-synt_1;tRNA-synt_1_2 | Leucine--tRNA ligase {ECO:0000256\|HAMAP-Rule:MF_00049} | | 1.978269681 | 0.29155862 | -0.614944458 | | 24 | 24 | 32.2 | 32.2 | 96.98 |

### All proteins

| Uniprot accession number | LFQ intensity WT_1 | LFQ intensity WT_2 | LFQ intensity WT_3 | LFQ intensity OrfSwap_1 | LFQ intensity OrfSwap_2 | LFQ intensity OrfSwap_3 | Pfam name | Uniprot full protein name | Student's T-test Significant OrfSwap_WT | -Log Student's T-test p-value OrfSwap_WT | Student's T-test q-value OrfSwap_WT | Student's T-test Difference OrfSwap_WT | Cluster | Peptides | Unique peptides | Sequence coverage [%] | Unique sequence coverage [%] | Mol. weight [kDa] |
|---|---|---|---|---|---|---|---|---|---|---|---|---|---|---|---|---|---|---|
| A0A0H3NSS0 | 32.1154 | 31.7687 | 31.7231 | 31.253 | 31.5234 | 30.8541 | Ribosomal_L30 | 50S ribosomal protein L30 {ECO:0000256\|HAMAP-Rule:MF_01371, ECO:0000256\|SAAS:SAAS00765691} | | 1.337278914 | 0.29983505 | -0.658910751 | | 9 | 9 | 71.2 | 71.2 | 6.514 |
| A0A0H3NFL8 | 28.4286 | 24.7652 | 28.0085 | 28.3327 | 28.302 | 28.0252 | Acyl_transf_1 | Malonyl CoA-acyl carrier protein transacylase {ECO:0000256\|PIRNR:PIRNR000446} | | 0.423342419 | 0.29989116 | 1.152544022 | | 8 | 8 | 42.1 | 42.1 | 32.41 |
| A0A0H3N9W0 | 26.6432 | 26.5906 | 26.2209 | 26.1248 | 24.7629 | 26.19 | ABC_tran | | | 0.752454037 | 0.30012287 | -0.792366028 | | 5 | 5 | 22.5 | 22.5 | 26.7 |
| A0A0H3NIK2 | 26.1334 | 26.2855 | 26.1473 | 25.3023 | 25.9582 | 25.3584 | EptA_B_N;Sulfatase | | | 1.404119931 | 0.30115068 | -0.64909935 | | 2 | 2 | 3.5 | 3.5 | 61.62 |
| A0A0H3NKV9 | 27.7021 | 28.575 | 28.4643 | 28.9615 | 29.0158 | 28.8089 | FKBP_C;FKBP_N | Peptidyl-prolyl cis-trans isomerase {ECO:0000256\|RuleBase:RU003915} | | 1.139618753 | 0.30158644 | 0.681596756 | | 6 | 6 | 26.8 | 26.8 | 23.74 |
| A0A0H3NJ77 | 24.8282 | 24.7419 | 25.6917 | 26.5604 | 25.9616 | 25.1376 | Cytochrom_B562 | | | 0.713393155 | 0.30237333 | 0.799273173 | | 5 | 5 | 35.9 | 35.9 | 13.91 |
| A0A0H3NR16 | 31.1396 | 30.4938 | 30.3608 | 30.2639 | 30.0953 | 29.5337 | Secretin;Secretin_N | | | 1.006616032 | 0.30240532 | -0.700386047 | | 22 | 22 | 44.7 | 44.7 | 61.77 |

### All proteins

| Uniprot accession number | LFQ intensity WT_1 | LFQ intensity WT_2 | LFQ intensity WT_3 | LFQ intensity OrfSwap_1 | LFQ intensity OrfSwap_2 | LFQ intensity OrfSwap_3 | Pfam name | Uniprot full protein name | Student's T-test Significant OrfSwap_WT | -Log Student's T-test p-value OrfSwap_WT | Student's T-test q-value OrfSwap_WT | Student's T-test Difference OrfSwap_WT | Cluster | Peptides | Unique peptides | Sequence coverage [%] | Unique sequence coverage [%] | Mol. weight [kDa] |
|---|---|---|---|---|---|---|---|---|---|---|---|---|---|---|---|---|---|---|
| A0A0H3NIB2 | 27.1682 | 27.216 | 27.1181 | 25.2549 | 24.6873 | 28.2298 | KOW;NusG | Transcription termination/antitermination protein NusG {ECO:0000256\|HAMAP-Rule:MF_00948, ECO:0000256\|RuleBase:RU000538, ECO:0000256\|SAAS:SAAS00126873} | | 0.432337135 | 0.30338462 | -1.110092163 | | 5 | 5 | 39.8 | 39.8 | 20.55 |
| A0A0H3NV02 | 25.2564 | 24.7025 | 25.5231 | 25.8333 | 25.7467 | 25.8782 | ABC_tran | | | 1.263043752 | 0.30398658 | 0.65870285 | | 5 | 5 | 12.1 | 12.1 | 55.52 |
| A0A0H3NAQ9 | 28.4056 | 28.7848 | 28.312 | 29.0704 | 29.2476 | 29.0269 | PfkB | Phosphofructokinase {ECO:0000256\|PIRNR:PIRNR000535} | | 1.737683224 | 0.30417508 | 0.614192963 | | 11 | 11 | 42.6 | 42.6 | 32.57 |
| A0A0H3NGL0 | 30.4784 | 30.3342 | 30.102 | 29.4819 | 29.8618 | 29.7324 | Biotin_carb_C;Biotin_carb_N;CPSase_L_D2 | Biotin carboxylase {ECO:0000256\|RuleBase:RU365063} | | 1.762888375 | 0.30441892 | -0.612796783 | | 26 | 26 | 58.7 | 58.7 | 46 |
| A0A0H3NEY8 | 27.0341 | 26.1375 | 27.1433 | 26.3629 | 25.9966 | 25.8627 | Radical_SAM | Pyruvate formate-lyase-activating enzyme {ECO:0000256\|RuleBase:RU362053} | | 0.926143162 | 0.30985489 | -0.697563807 | | 6 | 6 | 28.8 | 28.8 | 31.31 |
| A0A0H3NG48 | 24.83 | 26.3205 | 24.1647 | 25.6748 | 26.1081 | 26.0859 | SH3_3 | | | 0.581399966 | 0.31030189 | 0.851196925 | | 4 | 4 | 29.4 | 29.4 | 22.86 |

### All proteins

| Uniprot accession number | LFQ intensity WT_1 | LFQ intensity WT_2 | LFQ intensity WT_3 | LFQ intensity OrfSwap_1 | LFQ intensity OrfSwap_2 | LFQ intensity OrfSwap_3 | Pfam name | Uniprot full protein name | Student's T-test Significant OrfSwap_WT | -Log Student's T-test p-value OrfSwap_WT | Student's T-test q-value OrfSwap_WT | Student's T-test Difference OrfSwap_WT | Cluster | Peptides | Unique peptides | Sequence coverage [%] | Unique sequence coverage [%] | Mol. weight [kDa] |
|---|---|---|---|---|---|---|---|---|---|---|---|---|---|---|---|---|---|---|
| A0A0H3NDN5 | 27.7315 | 27.8402 | 27.7033 | 27.199 | 27.3012 | 27.1233 | ECH_1 | 1,4-dihydroxy-2-naphthoyl-CoA synthase {ECO:0000256\|HAMAP-Rule:MF_01934} | | 2.938721497 | 0.31049367 | -0.550487518 | | 6 | 6 | 22.5 | 22.5 | 31.69 |
| A0A0H3NKQ9 | 30.8766 | 31.1031 | 30.9808 | 31.4965 | 31.5644 | 31.5551 | PEP-utilizers;PEP-utilizers_C;PPDK_N | Phosphoenolpyruvate synthase {ECO:0000256\|PIRNR:PIRNR000854} | | 2.881999592 | 0.31108571 | 0.551807404 | | 34 | 34 | 43.8 | 43.8 | 88.84 |
| A0A0H3NDQ2 | 30.9814 | 31.0312 | 30.7205 | 30.3999 | 30.2942 | 30.3036 | EIIA-man;PTSIIB_sorb | | | 2.320409482 | 0.31154967 | -0.578502655 | | 19 | 19 | 50 | 50 | 34.99 |
| A0A0H3P0A4 | 27.3409 | 28.0332 | 28.0207 | 28.5268 | 28.5383 | 28.2613 | APH | | | 1.229549824 | 0.31166879 | 0.643851598 | | 6 | 6 | 25.8 | 25.8 | 29.59 |
| A0A0H3NHF2 | 29.5301 | 29.6784 | 29.5793 | 30.1368 | 30.174 | 30.1047 | Epimerase | ADP-L-glycero-D-manno-heptose-6-epimerase {ECO:0000256\|HAMAP-Rule:MF_01601} | | 3.457831618 | 0.31188424 | 0.542579651 | | 17 | 17 | 59 | 59 | 34.85 |
| A0A0H3NRN3 | 27.1606 | 24.9065 | 26.2312 | 25.5935 | 24.6902 | 25.3639 | Arginase | Agmatinase {ECO:0000256\|HAMAP-Rule:MF_01418} | | 0.552692969 | 0.31194872 | -0.8835392 | | 3 | 3 | 14.1 | 14.1 | 33.6 |
| A0A0H3NDK7 | 24.616 | 25.3165 | 25.5374 | 25.6256 | 25.6391 | 26.3262 | PAP2 | Lipid A 1-diphosphate synthase {ECO:0000256\|HAMAP-Rule:MF_01945} | | 0.913299483 | 0.3123754 | 0.707008998 | | 3 | 3 | 15.1 | 15.1 | 26.92 |

### All proteins

| Uniprot accession number | LFQ intensity WT_1 | LFQ intensity WT_2 | LFQ intensity WT_3 | LFQ intensity OrfSwap _1 | LFQ intensity OrfSwap _2 | LFQ intensity OrfSwap _3 | Pfam name | Uniprot full protein name | Student's T-test Significant OrfSwap_WT | -Log Student's T-test p-value OrfSwap_WT | Student's T-test q-value OrfSwap_WT | Student's T-test Difference OrfSwap_WT | Cluster | Peptides | Unique peptides | Sequence coverage [%] | Unique sequence coverage [%] | Mol. weight [kDa] |
|---|---|---|---|---|---|---|---|---|---|---|---|---|---|---|---|---|---|---|
| A0A0H3NNT2 | 25.7873 | 25.6011 | 26.0421 | 25.1672 | 25.0081 | 25.43 | FGGY_C;FGGY_N | Autoinducer-2 kinase {ECO:0000256\|HAMAP-Rule:MF_02053} | | 1.576105605 | 0.31266454 | -0.608431498 | | 5 | 5 | 16.2 | 16.2 | 57.44 |
| A0A0H3NMW4 | 32.0707 | 31.6516 | 31.6742 | 31.3047 | 31.0229 | 31.2639 | Aldedh | | | 1.686241053 | 0.31289032 | -0.60167249 | | 22 | 22 | 50 | 50 | 56.33 |
| A0A0H3NFI2 | 30.0068 | 29.8297 | 29.9056 | 28.7601 | 29.5165 | 29.508 | Toluene_X | | | 1.198408966 | 0.31338961 | -0.652482986 | | 6 | 6 | 21.1 | 21.1 | 47.7 |
| A0A0H3N9C5 | 30.7116 | 30.126 | 30.281 | 29.5846 | 29.7123 | 29.9631 | RRF | Ribosome-recycling factor {ECO:0000256\|HAMAP-Rule:MF_00040} | | 1.392522542 | 0.31360502 | -0.619540532 | | 16 | 16 | 70.8 | 70.8 | 20.56 |
| A0A0H3NC04 | 24.6784 | 25.6818 | 25.6799 | 25.681 | 26.0325 | 26.4916 | Macro | O-acetyl-ADP-ribose deacetylase {ECO:0000256\|HAMAP-Rule:MF_01205} | | 0.818419985 | 0.3137 | 0.721646627 | | 6 | 6 | 38.5 | 38.5 | 19.2 |
| A0A0H3NNR2 | 29.1399 | 28.9185 | 28.9672 | 28.5678 | 28.4792 | 28.2825 | Molybdop_Fe4S4;Molybdopterin;Molydop_binding | Periplasmic nitrate reductase {ECO:0000256\|HAMAP-Rule:MF_01630} | | 2.199307521 | 0.31427329 | -0.565396627 | | 21 | 21 | 27.4 | 27.4 | 92.87 |
| A0A0H3NEC0 | 26.0284 | 24.8055 | 24.443 | 27.6715 | 27.4225 | 23.7836 | NifU_N | Iron-sulfur cluster assembly scaffold protein IscU {ECO:0000256\|RuleBase:RU362089} | | 0.374064794 | 0.31428037 | 1.200246175 | | 2 | 2 | 19.5 | 19.5 | 13.82 |
| A0A0H3NMQ9 | 27.5097 | 28.1018 | 28.4852 | 28.4694 | 28.558 | 29.1997 | HAMP;MCPsignal;TarH | | | 0.907182304 | 0.31441042 | 0.710127513 | | 14 | 11 | 25.8 | 21.2 | 58.95 |
| A0A0H3NP23 | 25.4113 | 25.5028 | 25.2851 | 27.3299 | 24.3431 | 27.6633 | SBP_bac_3 | | | 0.421715867 | 0.31511111 | 1.045701981 | | 6 | 6 | 29.2 | 29.2 | 28.2 |

### All proteins

| Uniprot accession number | LFQ intensity WT_1 | LFQ intensity WT_2 | LFQ intensity WT_3 | LFQ intensity OrfSwap_1 | LFQ intensity OrfSwap_2 | LFQ intensity OrfSwap_3 | Pfam name | Uniprot full protein name | Student's T-test Significant OrfSwap_WT | -Log Student's T-test p-value OrfSwap_WT | Student's T-test q-value OrfSwap_WT | Student's T-test Difference OrfSwap_WT | Cluster | Peptides | Unique peptides | Sequence coverage [%] | Unique sequence coverage [%] | Mol. weight [kDa] |
|---|---|---|---|---|---|---|---|---|---|---|---|---|---|---|---|---|---|---|
| A0A0H3NM26 | 27.2232 | 27.5397 | 24.2186 | 27.3874 | 27.383 | 27.4248 | GSP_synth | | | 0.433341385 | 0.31538361 | 1.071232478 | | 9 | 9 | 23.3 | 23.3 | 44.98 |
| A0A0H3NYL3 | 26.9793 | 27.2143 | 27.1667 | 27.1801 | 24.7036 | 26.7236 | | | | 0.528553593 | 0.31543791 | -0.91768837 | | 4 | 4 | 12.9 | 12.9 | 51.23 |
| A0A0H3NED8 | 29.2418 | 28.8158 | 29.3514 | 29.3638 | 29.6893 | 30.5086 | OmpA | Peptidoglycan-associated protein {ECO:0000256\|HAMAP-Rule:MF_02204} | | 0.885032921 | 0.31569737 | 0.71753629 | | 11 | 11 | 55.7 | 55.7 | 18.87 |
| A0A0H3NC36 | 25.8929 | 25.8831 | 25.7791 | 24.4075 | 25.8022 | 25.2004 | Peptidase_S66 | | | 0.816322668 | 0.31575232 | -0.715019862 | | 3 | 3 | 14.5 | 14.5 | 33.35 |
| A0A0H3N7M4 | 30.8409 | 30.7064 | 30.9178 | 30.4657 | 30.0818 | 30.145 | 2-oxoacid_dh;Biotin_lipoyl;E3_binding | Acetyltransferase component of pyruvate dehydrogenase complex {ECO:0000256\|RuleBase:RU361137} | | 1.936237128 | 0.31673927 | -0.59089152 | | 27 | 27 | 38.5 | 38.5 | 66.05 |
| A0A0H3NIE0 | 32.5099 | 32.5107 | 32.535 | 32.1856 | 32.0361 | 31.5034 | Ribosomal_L5;Ribosomal_L5_C | 50S ribosomal protein L5 {ECO:0000256\|HAMAP-Rule:MF_01333} | | 1.374845087 | 0.32134154 | -0.610148748 | | 20 | 20 | 85.5 | 85.5 | 20.32 |
| A0A0H3NGB7 | 30.1684 | 30.1453 | 30.6355 | 30.7297 | 30.6896 | 31.5054 | Sod_Fe_C;Sod_Fe_N | Superoxide dismutase {ECO:0000256\|RuleBase:RU000414} | | 0.996934376 | 0.32359509 | 0.658508937 | | 7 | 7 | 30.6 | 30.6 | 21.34 |
| A0A0H3NJ22 | 26.9546 | 25.4753 | 26.9292 | 24.9533 | 25.5098 | 26.4191 | SpoU_methylase;SpoU_sub_bind | 23S rRNA (guanosine-2'-O-)-methyltransferase RlmB {ECO:0000256\|HAMAP-Rule:MF_01887} | | 0.564757971 | 0.32757187 | -0.825627009 | | 3 | 3 | 19.8 | 19.8 | 26.66 |

### All proteins

| Uniprot accession number | LFQ intensity WT_1 | LFQ intensity WT_2 | LFQ intensity WT_3 | LFQ intensity OrfSwap_1 | LFQ intensity OrfSwap_2 | LFQ intensity OrfSwap_3 | Pfam name | Uniprot full protein name | Student's T-test Significant OrfSwap_WT | -Log Student's T-test p-value OrfSwap_WT | Student's T-test q-value OrfSwap_WT | Student's T-test Difference OrfSwap_WT | Cluster | Peptides | Unique peptides | Sequence coverage [%] | Unique sequence coverage [%] | Mol. weight [kDa] |
|---|---|---|---|---|---|---|---|---|---|---|---|---|---|---|---|---|---|---|
| A0A0H3NG60 | 31.1246 | 30.9833 | 30.9039 | 31.7069 | 31.4823 | 31.4672 | 4HB_MCP_1;HAMP;MCPsignal | | | 2.255078817 | 0.3304878 | 0.548198064 | | 20 | 19 | 37.8 | 37.8 | 57.28 |
| A0A0H3NGG9 | 27.2805 | 27.4368 | 27.4415 | 27.2599 | 24.6848 | 27.4185 | AlaDh_PNT_C;AlaDh_PNT_N;PNTB_4TM | NAD(P) transhydrogenase subunit alpha {ECO:0000256\|PIRNR:PIRNR000203} | | 0.452221177 | 0.3330997 | -0.931850433 | | 7 | 7 | 15.7 | 15.7 | 54.27 |
| A0A0H3NBG3 | 31.829 | 31.7994 | 31.7629 | 31.37 | 31.2692 | 31.1868 | Gly_radical;PFL-like | | | 3.122094198 | 0.33373333 | -0.521743774 | | 51 | 50 | 55.9 | 55 | 85 |
| A0A0H3NHN7 | 26.9627 | 24.7805 | 26.8411 | 25.7637 | 24.6215 | 25.5446 | DNA_pol3_beta;DNA_pol3_beta_2;DNA_pol3_beta_3 | Beta sliding clamp {ECO:0000256\|PIRNR:PIRNR000804} | | 0.487749652 | 0.33446809 | -0.884833018 | | 7 | 7 | 21.6 | 21.6 | 40.55 |
| A0A0H3NHX6 | 28.2616 | 28.7453 | 28.768 | 28.8587 | 29.2586 | 29.4971 | PNP_UDP_1 | Uridine phosphorylase {ECO:0000256\|RuleBase:RU361131} | | 1.158433521 | 0.3359759 | 0.613174438 | | 13 | 13 | 60.1 | 60.1 | 27.14 |
| A0A0H3NEN0 | 25.6306 | 25.0737 | 24.9039 | 24.9108 | 24.5212 | 24.2924 | CM_2;PDT | | | 1.037814249 | 0.33698498 | -0.627929688 | | 5 | 5 | 15.3 | 15.3 | 42.82 |
| A0A0H3NJL8 | 29.2849 | 28.8586 | 29.0744 | 28.6002 | 28.6333 | 28.2833 | HemX | | | 1.569825888 | 0.33998802 | -0.567024867 | | 14 | 14 | 35.2 | 35.2 | 42.29 |
| A0A0H3NGG1 | 25.0403 | 26.2281 | 26.1855 | 25.7271 | 25.1543 | 24.3022 | ATP_bind_2 | RNase adapter protein RapZ {ECO:0000256\|HAMAP-Rule:MF_00636} | | 0.595806512 | 0.34570746 | -0.756775538 | | 5 | 5 | 20.8 | 20.8 | 32.46 |
| A0A0H3NB12 | 29.3751 | 29.2933 | 29.75 | 28.9677 | 29.0733 | 28.678 | B3_4;B5;FDX-ACB;tRNA_bind | Phenylalanine--tRNA ligase beta subunit {ECO:0000256\|HAMAP-Rule:MF_00283} | | 1.434327881 | 0.34578299 | -0.566399256 | | 22 | 22 | 26.9 | 26.9 | 87.32 |

### All proteins

| Uniprot accession number | LFQ intensity WT_1 | LFQ intensity WT_2 | LFQ intensity WT_3 | LFQ intensity OrfSwap_1 | LFQ intensity OrfSwap_2 | LFQ intensity OrfSwap_3 | Pfam name | Uniprot full protein name | Student's T-test Significant OrfSwap_WT | -Log Student's T-test p-value OrfSwap_WT | Student's T-test q-value OrfSwap_WT | Student's T-test Difference OrfSwap_WT | Cluster | Peptides | Unique peptides | Sequence coverage [%] | Unique sequence coverage [%] | Mol. weight [kDa] |
|---|---|---|---|---|---|---|---|---|---|---|---|---|---|---|---|---|---|---|
| A0A0H3NIX5 | 30.7994 | 31.0637 | 30.8237 | 30.3736 | 30.4551 | 30.2745 | FBPase | Fructose-1,6-bisphosphatase class 1 {ECO:0000256\|HAMAP-Rule:MF_01855} | | 2.221697296 | 0.34589971 | -0.527875264 | | 20 | 20 | 47.9 | 47.9 | 36.8 |
| A0A0H3NMF7 | 32.6046 | 32.5289 | 32.4989 | 32.0715 | 32.1487 | 31.8187 | Ribosomal_L22 | 50S ribosomal protein L22 {ECO:0000256\|HAMAP-Rule:MF_01331, ECO:0000256\|RuleBase:RU004008} | | 2.150582453 | 0.34627219 | -0.531136195 | | 14 | 14 | 65.5 | 65.5 | 12.23 |
| A0A0H3NPH0 | 33.079 | 33.3735 | 33.1417 | 32.5639 | 32.6062 | 32.819 | Cpn10 | 10 kDa chaperonin {ECO:0000256\|HAMAP-Rule:MF_00580, ECO:0000256\|RuleBase:RU000535} | | 1.959737622 | 0.34653801 | -0.535065969 | | 14 | 14 | 77.3 | 77.3 | 10.32 |
| A0A0H3NHC1 | 26.0081 | 26.1433 | 25.7886 | 25.9333 | 24.931 | 25.171 | DUF2810 | | | 0.929162051 | 0.34669436 | -0.634869258 | | 5 | 5 | 42.5 | 42.5 | 13.79 |
| A0A0H3N8D8 | 31.9814 | 31.7805 | 32.0535 | 32.4965 | 32.4402 | 32.4438 | FKBP_C;Trigger_C;Trigger_N | Trigger factor {ECO:0000256\|HAMAP-Rule:MF_00303, ECO:0000256\|RuleBase:RU003914} | | 2.472021252 | 0.3467619 | 0.521713257 | | 49 | 49 | 63.9 | 63.9 | 48.07 |
| A0A0H3NP22 | 29.2699 | 29.7339 | 29.2786 | 29.0508 | 29.0257 | 28.388 | Malate_synthase | Malate synthase {ECO:0000256\|RuleBase:RU000555} | | 1.072663022 | 0.3468 | -0.605995178 | | 22 | 22 | 41.7 | 41.7 | 60.38 |

### All proteins

| Uniprot accession number | LFQ intensity WT_1 | LFQ intensity WT_2 | LFQ intensity WT_3 | LFQ intensity OrfSwap_1 | LFQ intensity OrfSwap_2 | LFQ intensity OrfSwap_3 | Pfam name | Uniprot full protein name | Student's T-test Significant OrfSwap_WT | -Log Student's T-test p-value OrfSwap_WT | Student's T-test q-value OrfSwap_WT | Student's T-test Difference OrfSwap_WT | Cluster | Peptides | Unique peptides | Sequence coverage [%] | Unique sequence coverage [%] | Mol. weight [kDa] |
|---|---|---|---|---|---|---|---|---|---|---|---|---|---|---|---|---|---|---|
| A0A0H3NFL2 | 25.5358 | 25.9569 | 26.0008 | 24.9646 | 24.7456 | 25.833 | BRCT;DNA_ligase_aden;DNA_ligase_OB;DNA_ligase_ZBD | DNA ligase {ECO:0000256\|HAMAP-Rule:MF_01588, ECO:0000256\|RuleBase:RU000618, ECO:0000256\|SAAS:SAAS00919859} | | 0.828862218 | 0.34810496 | -0.650125504 | | 6 | 6 | 10 | 10 | 73.45 |
| A0A0H3NFT5 | 26.4884 | 24.7565 | 25.1044 | 26.1443 | 26.143 | 26.2432 | 2-Hacid_dh;2-Hacid_dh_C | | | 0.616117812 | 0.34913953 | 0.727050781 | | 11 | 11 | 30.7 | 30.7 | 43.99 |
| A0A0H3NC90 | 29.2279 | 28.66 | 29.0674 | 28.6544 | 28.2867 | 28.3176 | MliC | | | 1.287169885 | 0.35184 | -0.565535863 | | 4 | 4 | 37.7 | 37.7 | 12.58 |
| A0A0H3NJR4 | 26.2836 | 26.6316 | 25.4215 | 25.1572 | 26.2624 | 24.6731 | GST_C_4;GST_N_3 | | | 0.560531478 | 0.35278161 | -0.748007456 | | 4 | 4 | 13.1 | 13.1 | 23.95 |
| A0A0H3NJ10 | 26.7178 | 25.08 | 26.8604 | 24.7685 | 26.4335 | 24.9555 | RNase_T | Oligoribonuclease {ECO:0000256\|HAMAP-Rule:MF_00045} | | 0.463804367 | 0.35284814 | -0.833565394 | | 4 | 4 | 19.9 | 19.9 | 20.63 |
| A0A0H3NHW2 | 25.2517 | 25.3626 | 25.9418 | 25.2637 | 24.6972 | 24.7934 | DegT_DnrJ_EryC1 | dTDP-4-amino-4,6-dideoxygalactose transaminase {ECO:0000256\|HAMAP-Rule:MF_02026} | | 1.019939104 | 0.35308357 | -0.600615819 | | 5 | 5 | 10.1 | 10.1 | 42.01 |
| A0A0H3N7N9 | 23.591 | 26.3885 | 24.8966 | 24.3908 | 26.8243 | 26.6657 | PQQ | | | 0.372123264 | 0.35382659 | 1.001637777 | | 5 | 5 | 7.5 | 7.5 | 86.49 |

### All proteins

| Uniprot accession number | LFQ intensity WT_1 | LFQ intensity WT_2 | LFQ intensity WT_3 | LFQ intensity OrfSwap_1 | LFQ intensity OrfSwap_2 | LFQ intensity OrfSwap_3 | Pfam name | Uniprot full protein name | Student's T-test Significant OrfSwap_WT | -Log Student's T-test p-value OrfSwap_WT | Student's T-test q-value OrfSwap_WT | Student's T-test Difference OrfSwap_WT | Cluster | Peptides | Unique peptides | Sequence coverage [%] | Unique sequence coverage [%] | Mol. weight [kDa] |
|---|---|---|---|---|---|---|---|---|---|---|---|---|---|---|---|---|---|---|
| A0A0H3NHL6 | 29.871 | 30.0642 | 30.2287 | 30.7134 | 30.6894 | 24.3408 | GCV_H | Glycine cleavage system H protein {ECO:0000256\|HAMAP-Rule:MF_00272} | | 0.279170011 | 0.35417971 | -1.473438899 | | 6 | 6 | 14.7 | 14.7 | 13.84 |
| A0A0H3NEP5 | 24.4555 | 26.7291 | 26.6392 | 26.4679 | 27.0553 | 26.6925 | dTDP_sugar_isom | dTDP-4-dehydrorhamnose 3,5-epimerase {ECO:0000256\|RuleBase:RU364069} | | 0.449876165 | 0.36878453 | 0.797292709 | | 3 | 3 | 15.8 | 15.8 | 20.66 |
| A0A0H3NHN9 | 24.9388 | 24.6309 | 25.4308 | 25.8987 | 25.3944 | 25.4438 | GAF_2 | | | 0.957884378 | 0.36919008 | 0.578788122 | | 3 | 3 | 19.1 | 19.1 | 21.03 |
| A0A0H3NCQ6 | 31.8579 | 31.876 | 32.2091 | 31.4938 | 31.3934 | 31.5317 | HAMP;MCPsignal;TarH | | | 1.858388175 | 0.36919668 | -0.508031209 | | 24 | 20 | 45.8 | 38.2 | 59.61 |
| A0A0H3NJA1 | 24.9372 | 24.2957 | 24.9682 | 24.6222 | 25.8424 | 25.7125 | Fn3-like;Glyco_hydro_3;Glyco_hydro_3_C | | | 0.672339931 | 0.36932584 | 0.658707937 | | 6 | 6 | 7.3 | 7.3 | 83.39 |
| A0A0H3NDJ2 | 29.5019 | 29.4577 | 29.2604 | 28.8637 | 28.9534 | 28.9263 | DNA_gyraseA_C;DNA_topoisoIV | DNA gyrase subunit A {ECO:0000256\|HAMAP-Rule:MF_01897} | | 2.474236738 | 0.36965634 | -0.49221166 | | 24 | 24 | 24.1 | 24.1 | 97.06 |
| A0A0H3NF48 | 30.8338 | 30.9837 | 30.8661 | 30.5478 | 30.4151 | 30.2077 | tRNA_anti-codon;tRNA-synt_2 | Asparagine--tRNA ligase {ECO:0000256\|HAMAP-Rule:MF_00534} | | 2.008029667 | 0.36979272 | -0.504302343 | | 23 | 23 | 41.2 | 41.2 | 52.52 |

#### All proteins

| Uniprot accession number | LFQ intensity WT_1 | LFQ intensity WT_2 | LFQ intensity WT_3 | LFQ intensity OrfSwap_1 | LFQ intensity OrfSwap_2 | LFQ intensity OrfSwap_3 | Pfam name | Uniprot full protein name | Student's T-test Significant OrfSwap_WT | -Log Student's T-test p-value OrfSwap_WT | Student's T-test q-value OrfSwap_WT | Student's T-test Difference OrfSwap_WT | Cluster | Peptides | Unique peptides | Sequence coverage [%] | Unique sequence coverage [%] | Mol. weight [kDa] |
|---|---|---|---|---|---|---|---|---|---|---|---|---|---|---|---|---|---|---|
| A0A0H3NDK6 | 29.8215 | 29.8694 | 29.8068 | 29.4411 | 29.3556 | 29.2681 | Sec_GG;SecD_SecF;SecD-TM1 | Protein translocase subunit SecD {ECO:0000256\|HAMAP-Rule:MF_01463} | | 3.062788323 | 0.37020613 | -0.47763888 | | 13 | 13 | 24.6 | 24.6 | 66.6 |
| A0A0H3NDR5 | 29.1358 | 29.5594 | 29.6579 | 28.8308 | 28.8296 | 29.0868 | Acetate_kinase | Acetate kinase {ECO:0000256\|HAMAP-Rule:MF_00020} | | 1.376359432 | 0.37022222 | -0.535321554 | | 11 | 11 | 28.5 | 28.5 | 43.26 |
| A0A0H3NKV1 | 24.8146 | 26.0894 | 25.7966 | 25.0163 | 27.3644 | 26.7767 | GerE;Response_reg | | | 0.436566852 | 0.3706257 | 0.81892395 | | 5 | 5 | 25.5 | 25.5 | 24.35 |
| A0A0H3NCG3 | 32.4023 | 32.5891 | 32.7419 | 33.2272 | 33.0322 | 33.0076 | HSP70 | Chaperone protein DnaK {ECO:0000256\|HAMAP-Rule:MF_00332} | | 1.881041659 | 0.37070056 | 0.511249542 | | 43 | 43 | 51.1 | 51.1 | 69.26 |
| A0A0H3NVY3 | 31.5029 | 31.448 | 31.423 | 31.8377 | 31.8679 | 32.2001 | Ribosomal_S6 | 30S ribosomal protein S6 {ECO:0000256\|HAMAP-Rule:MF_00360, ECO:0000256\|SAAS:SAAS00020539} | | 1.901828157 | 0.37147875 | 0.510594686 | | 15 | 15 | 74 | 74 | 15.17 |
| A0A0H3NGF6 | 29.8741 | 30.0975 | 29.9962 | 29.4219 | 29.1892 | 29.7466 | Fumerase;Fumerase_C | Fumarate hydratase class I {ECO:0000256\|PIRNR:PIRNR001394} | | 1.433990882 | 0.37163636 | -0.53666687 | | 17 | 17 | 38.1 | 38.1 | 60.26 |

### All proteins

| Uniprot accession number | LFQ intensity WT_1 | LFQ intensity WT_2 | LFQ intensity WT_3 | LFQ intensity OrfSwap_1 | LFQ intensity OrfSwap_2 | LFQ intensity OrfSwap_3 | Pfam name | Uniprot full protein name | Student's T-test Significant OrfSwap_WT | -Log Student's T-test p-value OrfSwap_WT | Student's T-test q-value OrfSwap_WT | Student's T-test Difference OrfSwap_WT | Cluster | Peptides | Unique peptides | Sequence coverage [%] | Unique sequence coverage [%] | Mol. weight [kDa] |
|---|---|---|---|---|---|---|---|---|---|---|---|---|---|---|---|---|---|---|
| A0A0H3NDY0 | 27.8118 | 27.9413 | 27.7979 | 28.3695 | 28.3971 | 28.2243 | Glucokinase | Glucokinase {ECO:0000256\|HAMAP-Rule:MF_00524} | | 2.616228388 | 0.37163736 | 0.479971568 | | 9 | 9 | 35.5 | 35.5 | 34.59 |
| A0A0H3NFT9 | 27.3315 | 25.8223 | 26.957 | 27.3668 | 27.6602 | 27.1162 | MS_channel;TM_helix | | | 0.636172545 | 0.37206838 | 0.677470525 | | 7 | 7 | 18.5 | 18.5 | 31.06 |
| A0A0H3NB33 | 29.8156 | 29.0582 | 29.3894 | 30.0178 | 30.2822 | 29.6669 | SBP_bac_3 | | | 0.940058572 | 0.37450811 | 0.567873637 | | 16 | 16 | 53.6 | 53.6 | 27.26 |
| A0A0H3NIX0 | 28.4581 | 28.9836 | 28.7634 | 28.9665 | 29.3945 | 29.4674 | PMSR | Peptide methionine sulfoxide reductase MsrA {ECO:0000256\|HAMAP-Rule:MF_01401} | | 1.165982952 | 0.37451087 | 0.541081746 | | 4 | 4 | 21.7 | 21.7 | 23.4 |
| A0A0H3NF11 | 28.288 | 27.7101 | 27.705 | 28.619 | 28.5111 | 28.2079 | TctC | | | 1.117641649 | 0.37459079 | 0.544959386 | | 13 | 13 | 42.5 | 42.5 | 35.48 |
| A0A0H3NJB1 | 24.2961 | 25.3148 | 25.7578 | 26.2674 | 25.0418 | 26.1539 | CCP_MauG;Cytochrom_C;Haem_bd | | | 0.526677211 | 0.37462366 | 0.698111216 | | 4 | 4 | 10.5 | 10.5 | 51.91 |
| A0A0H3N7S5 | 24.2248 | 26.7822 | 27 | 25.18 | 25.1455 | 25.1764 | Peptidase_M24 | Methionine aminopeptidase {ECO:0000256\|HAMAP-Rule:MF_01974, ECO:0000256\|RuleBase:RU003653} | | 0.39604541 | 0.37471698 | -0.835041682 | | 7 | 7 | 26.1 | 26.1 | 29.29 |
| A0A0H3NCU4 | 28.8028 | 28.9876 | 28.6647 | 28.274 | 28.3235 | 28.3861 | S4;tRNA-synt_1b | Tyrosine--tRNA ligase {ECO:0000256\|HAMAP-Rule:MF_02006} | | 2.111575468 | 0.3750137 | -0.490507762 | | 13 | 13 | 30.7 | 30.7 | 47.28 |

### All proteins

| Uniprot accession number | LFQ intensity WT_1 | LFQ intensity WT_2 | LFQ intensity WT_3 | LFQ intensity OrfSwap_1 | LFQ intensity OrfSwap_2 | LFQ intensity OrfSwap_3 | Pfam name | Uniprot full protein name | Student's T-test Significant OrfSwap_WT | -Log Student's T-test p-value OrfSwap_WT | Student's T-test q-value OrfSwap_WT | Student's T-test Difference OrfSwap_WT | Cluster | Peptides | Unique peptides | Sequence coverage [%] | Unique sequence coverage [%] | Mol. weight [kDa] |
|---|---|---|---|---|---|---|---|---|---|---|---|---|---|---|---|---|---|---|
| A0A0H3NGP7 | 32.27 | 32.1025 | 32.5793 | 31.6611 | 31.7957 | 31.9434 | Ribosomal_L17 | 50S ribosomal protein L17 {ECO:0000256\|HAMAP-Rule:MF_01368, ECO:0000256\|RuleBase:RU000661} | | 1.482139502 | 0.37510383 | -0.517230352 | | 14 | 14 | 61.4 | 61.4 | 14.39 |
| A0A0H3NMN6 | 26.9728 | 26.9476 | 27.1091 | 27.5916 | 27.7533 | 27.2333 | Glyco_transf_5;Glycos_transf_1 | Glycogen synthase {ECO:0000256\|HAMAP-Rule:MF_00484} | | 1.480468382 | 0.37553134 | 0.516235987 | | 9 | 9 | 18.4 | 18.4 | 52.95 |
| A0A0H3NPV4 | 25.5061 | 25.8996 | 25.6377 | 25.0939 | 24.7554 | 25.5431 | LptF_LptG | | | 1.009575635 | 0.37635294 | -0.550321579 | | 3 | 3 | 10.3 | 10.3 | 39.55 |
| A0A0H3N9R0 | 26.6221 | 26.9032 | 26.5823 | 26.7371 | 24.3283 | 26.6815 | GDP_Man_Dehyd | | | 0.418958822 | 0.37719149 | -0.786911011 | | 3 | 3 | 9.5 | 9.5 | 37.13 |
| A0A0H3NPT4 | 30.8034 | 31.0893 | 30.9648 | 31.4314 | 31.4138 | 31.4364 | Ribonuc_L-PSP | | | 2.33402943 | 0.37736193 | 0.47468249 | | 13 | 13 | 74.2 | 74.2 | 13.58 |
| A0A0H3NWA8 | 28.6225 | 28.2552 | 28.6228 | 28.1144 | 27.7614 | 28.0928 | Amidohydro_1;Amidohydro_3 | Isoaspartyl dipeptidase {ECO:0000256\|PIRNR:PIRNR001238} | | 1.419481093 | 0.37794133 | -0.51064237 | | 11 | 11 | 38.5 | 38.5 | 40.32 |
| A0A0H3NMD7 | 28.0016 | 28.7409 | 28.5539 | 28.6961 | 29.0229 | 29.224 | MscL | Large-conductance mechanosensitive channel {ECO:0000256\|HAMAP-Rule:MF_00115, ECO:0000256\|SAAS:SAAS00980538} | | 0.951277797 | 0.3843183 | 0.548879623 | | 6 | 6 | 33.6 | 33.6 | 15.07 |

### All proteins

| Uniprot accession number | LFQ intensity WT_1 | LFQ intensity WT_2 | LFQ intensity WT_3 | LFQ intensity OrfSwap_1 | LFQ intensity OrfSwap_2 | LFQ intensity OrfSwap_3 | Pfam name | Uniprot full protein name | Student's T-test Significant OrfSwap_WT | -Log Student's T-test p-value OrfSwap_WT | Student's T-test q-value OrfSwap_WT | Student's T-test Difference OrfSwap_WT | Cluster | Peptides | Unique peptides | Sequence coverage [%] | Unique sequence coverage [%] | Mol. weight [kDa] |
|---|---|---|---|---|---|---|---|---|---|---|---|---|---|---|---|---|---|---|
| A0A0H3NHS8 | 27.9782 | 28.1891 | 28.0553 | 27.7289 | 27.5935 | 27.495 | Aminotran_4 | Branched-chain-amino-acid aminotransferase {ECO:0000256 \|RuleBase:RU364094} | | 2.160087535 | 0.38585752 | -0.468384425 | | 5 | 5 | 18.8 | 18.8 | 34.05 |
| A0A0H3NE22 | 28.5932 | 28.59 | 28.6226 | 29.1835 | 29.2087 | 28.8484 | SBP_bac_3 | | | 1.828981582 | 0.38667368 | 0.478285472 | | 20 | 20 | 65.8 | 65.8 | 28.81 |
| A0A0H3NI03 | 25.7341 | 27.8924 | 27.0988 | 27.3844 | 27.6637 | 27.7919 | Citrate_synt | Citrate synthase {ECO:0000256 \|PIRNR:PIRNR001369} | | 0.476747626 | 0.38687831 | 0.704919815 | | 7 | 7 | 22.6 | 22.6 | 43.17 |
| A0A0H3NE27 | 25.0927 | 23.671 | 24.7383 | 26.0999 | 25.6063 | 24.0375 | Flagellin_IN;FliD_C;FliD_N | Flagellar hook-associated protein 2 {ECO:0000256 \|RuleBase:RU362066} | | 0.422491338 | 0.38726702 | 0.747212092 | | 5 | 5 | 16.3 | 16.3 | 49.83 |
| A0A0H3NEC2 | 32.2273 | 32.2654 | 32.1921 | 31.8001 | 31.6444 | 31.8697 | FAD_binding_2;Succ_DH_flav_C | Succinate dehydrogenase flavoprotein subunit {ECO:0000256 \|RuleBase:RU362051} | | 2.54831598 | 0.38747507 | -0.456916173 | | 46 | 46 | 61.6 | 61.6 | 64.46 |
| A0A0H3NJA9 | 26.6288 | 26.4238 | 25.164 | 25.5684 | 25.0375 | 25.7051 | HlyD_D23 | UPF0194 membrane protein YbhG {ECO:0000256 \|HAMAP-Rule:MF_01304} | | 0.562651588 | 0.3896 | -0.635213852 | | 3 | 3 | 9.7 | 9.7 | 36.31 |

### All proteins

| Uniprot accession number | LFQ intensity WT_1 | LFQ intensity WT_2 | LFQ intensity WT_3 | LFQ intensity OrfSwap _1 | LFQ intensity OrfSwap _2 | LFQ intensity OrfSwap _3 | Pfam name | Uniprot full protein name | Student's T-test Significant OrfSwap_WT | -Log Student's T-test p-value OrfSwap_WT | Student's T-test q-value OrfSwap_WT | Student's T-test Difference OrfSwap_WT | Cluster | Peptides | Unique peptides | Sequence coverage [%] | Unique sequence coverage [%] | Mol. weight [kDa] |
|---|---|---|---|---|---|---|---|---|---|---|---|---|---|---|---|---|---|---|
| A0A0H3NJQ5 | 25.2138 | 26.3831 | 24.9375 | 24.7696 | 24.8684 | 25.0537 | ABC1 | Probable protein kinase UbiB {ECO:0000256\|HAMAP-Rule:MF_00414, ECO:0000256\|SAAS:SAAS00756156} | | 0.61130846 | 0.39005208 | -0.614228566 | | 3 | 3 | 5.5 | 5.5 | 63.24 |
| A0A0H3NIX1 | 30.8451 | 30.4691 | 30.6552 | 29.8376 | 30.2723 | 30.345 | SBP_bac_5 | | | 1.234688651 | 0.39081984 | -0.504822413 | | 25 | 25 | 54.6 | 54.6 | 58.46 |
| A0A0H3N9S2 | 27.8413 | 28.1169 | 27.9909 | 27.4827 | 27.6444 | 27.4284 | MmgE_PrpD | | | 1.972881379 | 0.39168992 | -0.464560191 | | 11 | 11 | 25.3 | 25.3 | 53.79 |
| A0A0H3NIA5 | 29.6583 | 30.0657 | 29.8388 | 29.3539 | 29.5253 | 29.2617 | DUF1043 | | | 1.549362244 | 0.39195939 | -0.473929723 | | 12 | 12 | 72.4 | 72.4 | 15.27 |
| A0A0H3N8J9 | 24.4042 | 24.6824 | 25.1301 | 26.3619 | 25.8523 | 24.1524 | Thioredoxin | | | 0.438674049 | 0.39205181 | 0.716641108 | | 4 | 4 | 24.6 | 24.6 | 31.81 |
| A0A0H3NGR2 | 25.0277 | 23.6795 | 25.0329 | 24.7452 | 25.8378 | 25.0943 | DUF1264 | | | 0.511112719 | 0.39240712 | 0.645799637 | | 3 | 3 | 18.8 | 18.8 | 28.86 |
| A0A0H3NJG2 | 26.9982 | 24.7662 | 26.9009 | 24.4716 | 25.5257 | 26.305 | DUF1414 | UPF0352 protein YejL {ECO:0000256\|HAMAP-Rule:MF_00816} | | 0.36494352 | 0.39263797 | -0.787671407 | | 2 | 2 | 28 | 28 | 8.231 |
| A0A0H3NA16 | 30.3915 | 30.3509 | 30.2098 | 31.0329 | 30.9962 | 30.4318 | IpgD | | | 1.169606424 | 0.3926581 | 0.502894084 | | 33 | 33 | 64.9 | 64.9 | 61.93 |
| A0A0H3NSL9 | 25.9089 | 25.2239 | 26.5892 | 25.2605 | 25.4312 | 25.2798 | ADH_N;ADH_zinc_N | | | 0.664873127 | 0.39281026 | -0.583510717 | | 4 | 4 | 18.2 | 18.2 | 34.39 |
| A0A0H3NMY0 | 29.2131 | 29.3885 | 29.6666 | 29.7992 | 30.0028 | 29.9041 | Aminotran_1_2 | 2-amino-3-ketobutyrate coenzyme A ligase {ECO:0000256\|HAMAP-Rule:MF_00985} | | 1.530636911 | 0.39301031 | 0.479313533 | | 20 | 20 | 43.2 | 43.2 | 43.03 |
| A0A0H3NCU0 | 29.0561 | 28.6392 | 29.0987 | 28.6484 | 28.4759 | 28.1973 | MarR | Transcriptional regulator SlyA {ECO:0000256\|HAMAP-Rule:MF_01819} | | 1.172405517 | 0.39325313 | -0.490815481 | | 3 | 3 | 27.4 | 27.4 | 16.7 |

#### All proteins

| Uniprot accession number | LFQ intensity WT_1 | LFQ intensity WT_2 | LFQ intensity WT_3 | LFQ intensity OrfSwap_1 | LFQ intensity OrfSwap_2 | LFQ intensity OrfSwap_3 | Pfam name | Uniprot full protein name | Student's T-test Significant OrfSwap_WT | -Log Student's T-test p-value OrfSwap_WT | Student's T-test q-value OrfSwap_WT | Student's T-test Difference OrfSwap_WT | Cluster | Peptides | Unique peptides | Sequence coverage [%] | Unique sequence coverage [%] | Mol. weight [kDa] |
|---|---|---|---|---|---|---|---|---|---|---|---|---|---|---|---|---|---|---|
| A0A0H3NAS0 | 29.4714 | 30.1419 | 29.9126 | 29.6024 | 30.9708 | 30.7887 | Ribosomal_L20 | 50S ribosomal protein L20 {ECO:0000256\|HAMAP-Rule:MF_00382, ECO:0000256\|RuleBase:RU000560} | | 0.577624941 | 0.39340816 | 0.61201032 | | 8 | 8 | 40.7 | 40.7 | 13.5 |
| A0A0H3NK87 | 32.1353 | 31.7962 | 31.8365 | 31.4439 | 31.5611 | 31.3898 | Ribosomal_L32p | 50S ribosomal protein L32 {ECO:0000256\|HAMAP-Rule:MF_00340} | | 1.744731881 | 0.39368 | -0.457740784 | | 6 | 6 | 78.9 | 78.9 | 6.446 |
| A0A0H3NAY6 | 31.3111 | 31.1354 | 31.0465 | 30.1594 | 30.9092 | 30.8725 | LTXXQ | | | 0.944783688 | 0.39423232 | -0.517256419 | | 20 | 20 | 57.8 | 57.8 | 18.23 |
| A0A0H3NFQ5 | 30.0656 | 30.3156 | 29.7532 | 30.5478 | 30.4672 | 30.553 | TAL_FSA | Transaldolase {ECO:0000256\|HAMAP-Rule:MF_00492, ECO:0000256\|SAAS:SAAS00118670} | | 1.353504815 | 0.39424121 | 0.477877299 | | 22 | 21 | 64.6 | 59.5 | 35.59 |
| A0A0H3NR39 | 26.7789 | 27.1644 | 27.2016 | 25.0572 | 27.848 | 25.9512 | Sigma70_r1_2;Sigma70_r2;Sigma70_r3;Sigma70_r4 | RNA polymerase sigma factor RpoS {ECO:0000256\|HAMAP-Rule:MF_00959} | | 0.385083767 | 0.39441432 | -0.762854894 | | 7 | 6 | 25.2 | 22.1 | 37.93 |
| A0A0H3NEN1 | 28.6549 | 28.7437 | 28.5391 | 27.8467 | 28.451 | 28.1812 | Wzz | | | 1.237468358 | 0.39469018 | -0.486274083 | | 13 | 13 | 36.7 | 36.7 | 36.26 |
| A0A0H3ND15 | 30.365 | 30.4589 | 30.0534 | 29.6643 | 30.198 | 29.4924 | KGG | | | 0.969394442 | 0.3959202 | -0.507509867 | | 6 | 3 | 73.3 | 36.7 | 6.139 |
| A0A0H3NJ23 | 28.9602 | 29.6231 | 29.1427 | 28.2891 | 28.7124 | 29.1243 | Glutaredoxin | Glutaredoxin {ECO:0000256\|RuleBase:RU364065} | | 0.789697877 | 0.39680597 | -0.533426921 | | 5 | 5 | 60.2 | 60.2 | 9.135 |
| A0A0H3NMH1 | 24.8714 | 25.5116 | 24.6611 | 25.1351 | 25.6569 | 25.8806 | YecR | | | 0.736462004 | 0.39743921 | 0.542847951 | | 4 | 4 | 42.3 | 42.3 | 12.2 |

### All proteins

| Uniprot accession number | LFQ intensity WT_1 | LFQ intensity WT_2 | LFQ intensity WT_3 | LFQ intensity OrfSwap_1 | LFQ intensity OrfSwap_2 | LFQ intensity OrfSwap_3 | Pfam name | Uniprot full protein name | Student's T-test Significant OrfSwap_WT | -Log Student's T-test p-value OrfSwap_WT | Student's T-test q-value OrfSwap_WT | Student's T-test Difference OrfSwap_WT | Cluster | Peptides | Unique peptides | Sequence coverage [%] | Unique sequence coverage [%] | Mol. weight [kDa] |
|---|---|---|---|---|---|---|---|---|---|---|---|---|---|---|---|---|---|---|
| A0A0H3NFJ4 | 25.3835 | 25.7195 | 25.4307 | 25.6598 | 24.4238 | 24.7716 | FtsJ | Ribosomal RNA large subunit methyltransferase M {ECO:0000256\|HAMAP-Rule:MF_01551, ECO:0000256\|SAAS:SAAS00058276} | | 0.662541308 | 0.39854455 | -0.559532801 | | 2 | 2 | 5.7 | 5.7 | 42.02 |
| A0A0H3N972 | 25.2918 | 24.7593 | 25.2611 | 25.9457 | 25.7165 | 25.193 | THF_DHG_CYH;THF_DHG_CYH_C | Bifunctional protein FolD {ECO:0000256\|HAMAP-Rule:MF_01576} | | 0.847573654 | 0.40159214 | 0.514312108 | | 4 | 4 | 17 | 17 | 30.87 |
| A0A0H3NHD5 | 25.1766 | 25.7142 | 26.4629 | 25.4283 | 26.9902 | 26.8931 | Mannitol_dh;Mannitol_dh_C | Mannitol-1-phosphate 5-dehydrogenase {ECO:0000256\|HAMAP-Rule:MF_00196, ECO:0000256\|SAAS:SAAS00320441} | | 0.446863092 | 0.40161765 | 0.652629852 | | 5 | 5 | 17.5 | 17.5 | 40.9 |
| A0A0H3N961 | 28.6252 | 29.0888 | 28.6125 | 28.3099 | 28.3647 | 28.2555 | PD40;TolB_N | Tol-Pal system protein TolB {ECO:0000256\|HAMAP-Rule:MF_00671, ECO:0000256\|SAAS:SAAS01034303} | | 1.360515642 | 0.40258128 | -0.465431849 | | 13 | 13 | 36.7 | 36.7 | 46.15 |

### All proteins

| Uniprot accession number | LFQ intensity WT_1 | LFQ intensity WT_2 | LFQ intensity WT_3 | LFQ intensity OrfSwap_1 | LFQ intensity OrfSwap_2 | LFQ intensity OrfSwap_3 | Pfam name | Uniprot full protein name | Student's T-test Significant OrfSwap_WT | -Log Student's T-test p-value OrfSwap_WT | Student's T-test q-value OrfSwap_WT | Student's T-test Difference OrfSwap_WT | Cluster | Peptides | Unique peptides | Sequence coverage [%] | Unique sequence coverage [%] | Mol. weight [kDa] |
|---|---|---|---|---|---|---|---|---|---|---|---|---|---|---|---|---|---|---|
| A0A0H3NNL2 | 28.5385 | 29.2797 | 29.1206 | 28.113 | 28.7161 | 28.5577 | Gln-synt_C;Gln-synt_N | Glutamine synthetase {ECO:0000256\|RuleBase:RU004356} | | 0.830903666 | 0.40357531 | -0.517328898 | | 14 | 14 | 27.5 | 27.5 | 51.79 |
| A0A0H3NM85 | 31.3551 | 31.1254 | 31.0117 | 30.6846 | 30.838 | 30.6392 | Ribosomal_L21p | 50S ribosomal protein L21 {ECO:0000256\|HAMAP-Rule:MF_01363, ECO:0000256\|RuleBase:RU000562, ECO:0000256\|SAAS:SAAS00352960} | | 1.708747288 | 0.40657213 | -0.443454107 | | 10 | 10 | 64.1 | 64.1 | 11.58 |
| A0A0H3NK11 | 30.3652 | 30.4483 | 30.1751 | 29.6985 | 30.0136 | 29.9438 | FMN_red | NAD(P)H dehydrogenase (quinone) {ECO:0000256\|HAMAP-Rule:MF_01017} | | 1.623149494 | 0.40844878 | -0.44422404 | | 13 | 13 | 60.1 | 60.1 | 20.87 |
| A0A0H3NCF8 | 31.8707 | 32.0937 | 32.0503 | 32.3298 | 32.4994 | 32.4596 | TAL_FSA | Transaldolase {ECO:0000256\|HAMAP-Rule:MF_00492, ECO:0000256\|RuleBase:RU004155, ECO:0000256\|SAAS:SAAS00118670} | | 2.118508238 | 0.4140438 | 0.424692154 | | 31 | 30 | 69.4 | 64.4 | 35.17 |

### All proteins

| Uniprot accession number | LFQ intensity WT_1 | LFQ intensity WT_2 | LFQ intensity WT_3 | LFQ intensity OrfSwap_1 | LFQ intensity OrfSwap_2 | LFQ intensity OrfSwap_3 | Pfam name | Uniprot full protein name | Student's T-test Significant OrfSwap_WT | -Log Student's T-test p-value OrfSwap_WT | Student's T-test q-value OrfSwap_WT | Student's T-test Difference OrfSwap_WT | Cluster | Peptides | Unique peptides | Sequence coverage [%] | Unique sequence coverage [%] | Mol. weight [kDa] |
|---|---|---|---|---|---|---|---|---|---|---|---|---|---|---|---|---|---|---|
| A0A0H3N9M5 | 29.0529 | 28.4709 | 28.962 | 29.3539 | 29.2217 | 29.2953 | Seryl_tRNA_N;tRNA-synt_2b | Serine--tRNA ligase {ECO:0000256\|HAMAP-Rule:MF_00176} | | 1.174834121 | 0.41476029 | 0.461701711 | | 16 | 16 | 36.7 | 36.7 | 48.58 |
| A0A0H3NCQ0 | 28.2086 | 28.3901 | 27.9394 | 27.7352 | 27.9084 | 27.5219 | | | | 1.247583593 | 0.41504854 | -0.45753034 | | 8 | 8 | 75.3 | 75.3 | 10.13 |
| A0A0H3N800 | 28.2466 | 28.2549 | 28.44 | 28.0091 | 27.8611 | 27.8087 | Pribosyltran | Xanthine phosphoribosyltransferase {ECO:0000256\|HAMAP-Rule:MF_01903} | | 2.073433401 | 0.41544231 | -0.42089208 | | 4 | 4 | 30.3 | 30.3 | 16.97 |
| A0A0H3NJ87 | 25.8867 | 26.0481 | 26.129 | 26.2228 | 26.7878 | 26.424 | PBP | | | 1.193204469 | 0.41621205 | 0.456935883 | | 5 | 5 | 36.1 | 36.1 | 17.07 |
| A0A0H3NDX8 | 29.2984 | 29.2519 | 29.2749 | 28.7748 | 28.7207 | 29.048 | SBP_bac_3 | | | 1.856945933 | 0.41698551 | -0.427244822 | | 12 | 12 | 51.2 | 51.2 | 28.38 |
| A0A0H3NJQ1 | 30.151 | 30.0739 | 29.8999 | 29.5112 | 29.6985 | 29.6581 | DRTGG;PTA_PTB | Phosphate acetyltransferase {ECO:0000256\|PIRNR:PIRNR006107} | | 1.95848958 | 0.42296403 | -0.419006348 | | 23 | 23 | 29.1 | 29.1 | 77.28 |
| A0A0H3N962 | 32.3942 | 32.4392 | 32.3383 | 32.0372 | 31.9974 | 31.9445 | Aconitase;Aconitase_2_N;Aconitase_B_N | Aconitate hydratase B {ECO:0000256\|PIRNR:PIRNR036687} | | 3.254476076 | 0.42352153 | -0.397559484 | | 52 | 52 | 51.1 | 51.1 | 93.53 |
| A0A0H3N7K6 | 24.7901 | 26.4784 | 24.808 | 25.8299 | 25.9362 | 26.0659 | Dala_Dala_lig_C;Dala_Dala_lig_N | D-alanine--D-alanine ligase {ECO:0000256\|HAMAP-Rule:MF_00047, ECO:0000256\|SAAS:SAAS01090413} | | 0.446094031 | 0.43646778 | 0.585188548 | | 4 | 4 | 16.7 | 16.7 | 32.64 |

### All proteins

| Uniprot accession number | LFQ intensity WT_1 | LFQ intensity WT_2 | LFQ intensity WT_3 | LFQ intensity OrfSwap_1 | LFQ intensity OrfSwap_2 | LFQ intensity OrfSwap_3 | Pfam name | Uniprot full protein name | Student's T-test Significant OrfSwap_WT | -Log Student's T-test p-value OrfSwap_WT | Student's T-test q-value OrfSwap_WT | Student's T-test Difference OrfSwap_WT | Cluster | Peptides | Unique peptides | Sequence coverage [%] | Unique sequence coverage [%] | Mol. weight [kDa] |
|---|---|---|---|---|---|---|---|---|---|---|---|---|---|---|---|---|---|---|
| A0A0H3N9M1 | 29.0663 | 28.6926 | 28.7837 | 28.5828 | 28.6823 | 27.8313 | Pyr_redox_2 | Thioredoxin reductase {ECO:0000256\|RuleBase:RU003881} | | 0.761366743 | 0.43729524 | -0.482051849 | | 8 | 8 | 43.2 | 43.2 | 34.68 |
| A0A0H3NJC8 | 30.7392 | 30.6228 | 30.6993 | 30.5535 | 30.372 | 29.751 | Ribosomal_L34 | 50S ribosomal protein L34 {ECO:0000256\|HAMAP-Rule:MF_00391, ECO:0000256\|SAAS:SAAS00020245} | | 0.875828673 | 0.44362945 | -0.461580912 | | 5 | 5 | 37 | 37 | 5.38 |
| A0A0H3NBB0 | 28.8936 | 29.1054 | 28.9635 | 29.399 | 29.2647 | 29.5141 | TPP_enzyme_C;TPP_enzyme_M;TPP_enzyme_N | | | 1.88123254 | 0.44364929 | 0.405054092 | | 15 | 15 | 30.8 | 30.8 | 61.68 |
| A0A0H3NIX6 | 31.3024 | 31.1102 | 31.2632 | 31.6036 | 31.7835 | 31.5034 | DeoC | | | 1.799056096 | 0.44520095 | 0.404911041 | | 23 | 23 | 51.7 | 51.7 | 38.09 |
| A0A0H3NHD4 | 29.3121 | 28.4905 | 28.5253 | 28.1258 | 28.0351 | 28.6969 | ADH_N;ADH_zinc_N | L-threonine 3-dehydrogenase {ECO:0000256\|HAMAP-Rule:MF_00627, ECO:0000256\|SAAS:SAAS00321739} | | 0.654091924 | 0.44639059 | -0.489997228 | | 8 | 8 | 23.8 | 23.8 | 37.21 |
| A0A0H3NPK6 | 29.9736 | 30.0987 | 29.933 | 29.4308 | 29.6871 | 29.6809 | HTH_12;OB_RNB;RNB;S1 | Ribonuclease R {ECO:0000256\|HAMAP-Rule:MF_01895} | | 1.828767865 | 0.44715094 | -0.402170181 | | 23 | 23 | 30.3 | 30.3 | 92.05 |

#### All proteins

| Uniprot accession number | LFQ intensity WT_1 | LFQ intensity WT_2 | LFQ intensity WT_3 | LFQ intensity OrfSwap _1 | LFQ intensity OrfSwap _2 | LFQ intensity OrfSwap _3 | Pfam name | Uniprot full protein name | Student's T-test Significant OrfSwap_WT | -Log Student's T-test p-value OrfSwap_WT | Student's T-test q-value OrfSwap_WT | Student's T-test Difference OrfSwap_WT | Cluster | Peptides | Unique peptides | Sequence coverage [%] | Unique sequence coverage [%] | Mol. weight [kDa] |
|---|---|---|---|---|---|---|---|---|---|---|---|---|---|---|---|---|---|---|
| A0A0H3NGI8 | 28.6344 | 28.734 | 28.9788 | 29.524 | 29.4103 | 28.7806 | SecIII_SopE_N;SopE_GEF | Guanine nucleotide exchange factor {ECO:0000256\|PIRNR:PIRNR034781} | | 0.836645579 | 0.448723 | 0.45590655 | | 14 | 14 | 54.6 | 54.6 | 26.64 |
| A0A0H3NLZ5 | 25.2454 | 25.8467 | 25.3979 | 25.8607 | 25.8638 | 26.0591 | CN_hydrolase | | | 1.055027231 | 0.4491897 | 0.431168874 | | 7 | 7 | 27.1 | 27.1 | 33.12 |
| A0A0H3NH30 | 27.3723 | 26.7651 | 26.8231 | 27.3003 | 27.5634 | 27.3916 | DUF2756 | | | 0.970919573 | 0.45675701 | 0.431612651 | | 3 | 3 | 29.7 | 29.7 | 15.3 |
| A0A0H3N8Y8 | 29.198 | 28.3232 | 28.4738 | 27.9828 | 27.9978 | 28.593 | AAA_2;ClpB_D2-small;zf-C4_ClpX | ATP-dependent Clp protease ATP-binding subunit ClpX {ECO:0000256\|HAMAP-Rule:MF_00175, ECO:0000256\|SAAS:SAAS01076755} | | 0.634504666 | 0.45998135 | -0.473791758 | | 14 | 14 | 36.2 | 36.2 | 46.18 |
| A0A0H3NCU6 | 24.9171 | 24.839 | 25.2067 | 24.4122 | 26.1258 | 26.098 | Pribosyltran | Hypoxanthine phosphoribosyltransferase {ECO:0000256\|RuleBase:RU364099} | | 0.410199909 | 0.46069767 | 0.55774498 | | 5 | 5 | 32 | 32 | 20.07 |
| A0A0H3NDL1 | 32.1291 | 32.5141 | 32.5472 | 32.8137 | 32.7959 | 32.7802 | GDPD | | | 1.38616466 | 0.46176334 | 0.399780273 | | 31 | 31 | 60.4 | 60.4 | 40.42 |
| A0A0H3NJJ6 | 28.4825 | 28.0058 | 28.5515 | 29.158 | 28.6594 | 24.6831 | Ribonuc_red_sm | | | 0.232998014 | 0.46659584 | -0.846429189 | | 7 | 7 | 25.8 | 25.8 | 43.55 |

### All proteins

| Uniprot accession number | LFQ intensity WT_1 | LFQ intensity WT_2 | LFQ intensity WT_3 | LFQ intensity OrfSwap_1 | LFQ intensity OrfSwap_2 | LFQ intensity OrfSwap_3 | Pfam name | Uniprot full protein name | Student's T-test Significant OrfSwap_WT | -Log Student's T-test p-value OrfSwap_WT | Student's T-test q-value OrfSwap_WT | Student's T-test Difference OrfSwap_WT | Cluster | Peptides | Unique peptides | Sequence coverage [%] | Unique sequence coverage [%] | Mol. weight [kDa] |
|---|---|---|---|---|---|---|---|---|---|---|---|---|---|---|---|---|---|---|
| A0A0H3NFX0 | 29.3556 | 29.3008 | 29.2748 | 28.9774 | 28.8877 | 28.9796 | | Uracil phosphoribosyltransferase {ECO:0000256\|HAMAP-Rule:MF_01218, ECO:0000256\|SAAS:SAAS00709136} | | 3.146644215 | 0.46662673 | -0.362157822 | | 11 | 11 | 60.1 | 60.1 | 22.53 |
| A0A0H3NJ45 | 32.506 | 32.2134 | 32.3541 | 32.1057 | 32.0156 | 31.7645 | 2-oxogl_dehyd_N;E1_dh;OxoGdeHyase_C;Transket_pyr | | | 1.393165085 | 0.46696296 | -0.395890554 | | 55 | 55 | 55.9 | 55.9 | 104.8 |
| A0A0H3ND07 | 25.4739 | 25.5155 | 25.4881 | 26.398 | 25.8727 | 25.5128 | Fer4_11;Fer4_4 | | | 0.780093639 | 0.46844037 | 0.435361226 | | 2 | 2 | 11.2 | 11.2 | 22.77 |
| A0A0H3NJ17 | 29.0512 | 29.5709 | 29.0133 | 29.6809 | 29.3306 | 29.9138 | PGM_PMM_I;PGM_PMM_II;PGM_PMM_III;PGM_PMM_IV | | | 0.804311686 | 0.46852968 | 0.429961522 | | 20 | 20 | 43.6 | 43.6 | 58.09 |
| A0A0H3NTL9 | 24.5633 | 25.8712 | 25.9415 | 24.2355 | 25.3579 | 25.1548 | GTP_EFTU;GTP_EFTU_D2;SelB-wing_2;SelB-wing_3 | | | 0.406894206 | 0.46869885 | -0.542592367 | | 2 | 2 | 4.2 | 4.2 | 68.65 |
| A0A0H3NBL7 | 30.6175 | 30.8343 | 30.6335 | 30.2831 | 30.2885 | 30.3962 | Aminotran_1_2 | Aminotransferase {ECO:0000256\|RuleBase:RU000481} | | 2.037862629 | 0.46935469 | -0.372507095 | | 21 | 21 | 54 | 54 | 43.52 |
| A0A0H3NF80 | 29.1732 | 29.2544 | 29.6584 | 29.5328 | 29.1076 | 27.899 | CCG;Fer4_8 | | | 0.430795378 | 0.47275455 | -0.515509923 | | 21 | 21 | 56.1 | 56.1 | 44.13 |
| A0A0H3N9C6 | 25.6937 | 23.7844 | 26.2041 | 25.7122 | 25.9604 | 25.7916 | Pyr_redox_2;Thioredoxin_3 | | | 0.33056297 | 0.47282916 | 0.593996048 | | 6 | 6 | 15.2 | 15.2 | 55.95 |

#### All proteins

| Uniprot accession number | LFQ intensity WT_1 | LFQ intensity WT_2 | LFQ intensity WT_3 | LFQ intensity OrfSwap_1 | LFQ intensity OrfSwap_2 | LFQ intensity OrfSwap_3 | Pfam name | Uniprot full protein name | Student's T-test Significant OrfSwap_WT | -Log Student's T-test p-value OrfSwap_WT | Student's T-test q-value OrfSwap_WT | Student's T-test Difference OrfSwap_WT | Cluster | Peptides | Unique peptides | Sequence coverage [%] | Unique sequence coverage [%] | Mol. weight [kDa] |
|---|---|---|---|---|---|---|---|---|---|---|---|---|---|---|---|---|---|---|
| A0A0H3NHZ8 | 26.6144 | 26.7297 | 26.9176 | 26.3744 | 26.2627 | 26.4917 | Ubie_methyltran | Ubiquinone/menaquinone biosynthesis C-methyltransferase UbiE {ECO:0000256\|HAMAP-Rule:MF_01813} | | 1.573079346 | 0.47402268 | -0.377675374 | | 5 | 5 | 26.3 | 26.3 | 28.14 |
| A0A0H3NJG5 | 30.088 | 30.1188 | 30.083 | 30.5688 | 30.4246 | 30.3714 | PNP_UDP_1 | Purine nucleoside phosphorylase DeoD-type {ECO:0000256\|HAMAP-Rule:MF_01627} | | 2.402830181 | 0.4760724 | 0.35835139 | | 18 | 18 | 73.2 | 73.2 | 25.98 |
| A0A0H3NEM4 | 30.8334 | 30.9856 | 30.9334 | 31.4848 | 31.1598 | 31.2281 | AAA;AAA_2;Clp_N;ClpB_D2-small | Chaperone protein ClpB {ECO:0000256\|RuleBase:RU362034} | | 1.58061441 | 0.47725508 | 0.373439153 | | 54 | 54 | 57.3 | 57.3 | 95.44 |
| A0A0H3NAY0 | 28.2996 | 28.5279 | 28.263 | 28.576 | 28.7732 | 28.8588 | Aminotran_3 | Succinylornithine transaminase {ECO:0000256\|HAMAP-Rule:MF_01173} | | 1.466946774 | 0.47805357 | 0.372465769 | | 15 | 15 | 46.8 | 46.8 | 43.83 |
| A0A0H3NXF6 | 26.5156 | 26.8116 | 26.9494 | 27.1227 | 26.6079 | 24.8413 | Pro_CA | | | 0.333473839 | 0.4781387 | -0.568232854 | | 10 | 10 | 46.7 | 46.7 | 26.64 |

### All proteins

| Uniprot accession number | LFQ intensity WT_1 | LFQ intensity WT_2 | LFQ intensity WT_3 | LFQ intensity OrfSwap_1 | LFQ intensity OrfSwap_2 | LFQ intensity OrfSwap_3 | Pfam name | Uniprot full protein name | Student's T-test Significant OrfSwap_WT | -Log Student's T-test p-value OrfSwap_WT | Student's T-test q-value OrfSwap_WT | Student's T-test Difference OrfSwap_WT | Cluster | Peptides | Unique peptides | Sequence coverage [%] | Unique sequence coverage [%] | Mol. weight [kDa] |
|---|---|---|---|---|---|---|---|---|---|---|---|---|---|---|---|---|---|---|
| A0A0H3NKP0 | 30.7368 | 30.4581 | 30.3505 | 30.4232 | 30.1105 | 29.7947 | IF3_C;IF3_N | Translation initiation factor IF-3 {ECO:0000256\|HAMAP-Rule:MF_00080, ECO:0000256\|RuleBase:RU000646} | | 0.879172969 | 0.47878924 | -0.405639648 | | 14 | 14 | 83.3 | 83.3 | 20.59 |
| A0A0H3NCM7 | 29.1419 | 29.1573 | 29.1182 | 28.8633 | 28.8958 | 28.5357 | G6PD_C;G6PD_N | Glucose-6-phosphate 1-dehydrogenase {ECO:0000256\|HAMAP-Rule:MF_00966} | | 1.498445572 | 0.47902703 | -0.374195099 | | 20 | 20 | 45 | 45 | 55.92 |
| A0A0H3NA48 | 28.9188 | 28.6772 | 28.9615 | 29.1981 | 29.2504 | 29.2042 | ADK_lid | Adenylate kinase {ECO:0000256\|HAMAP-Rule:MF_00235, ECO:0000256\|RuleBase:RU003331, ECO:0000256\|SAAS:SAAS00914410} | | 1.811771577 | 0.47917303 | 0.365025838 | | 11 | 11 | 57.9 | 57.9 | 23.49 |
| A0A0H3NEM2 | 26.9168 | 27.0124 | 26.8777 | 26.7007 | 26.0895 | 26.8129 | DEAD;Helicase_C | ATP-dependent RNA helicase SrmB {ECO:0000256\|HAMAP-Rule:MF_00967} | | 0.813611214 | 0.4841674 | -0.401301702 | | 9 | 9 | 23.9 | 23.9 | 50.06 |

#### All proteins

| Uniprot accession number | LFQ intensity WT_1 | LFQ intensity WT_2 | LFQ intensity WT_3 | LFQ intensity OrfSwap _1 | LFQ intensity OrfSwap _2 | LFQ intensity OrfSwap _3 | Pfam name | Uniprot full protein name | Student's T-test Significant OrfSwap_WT | -Log Student's T-test p-value OrfSwap_WT | Student's T-test q-value OrfSwap_WT | Student's T-test Difference OrfSwap_WT | Cluster | Peptides | Unique peptides | Sequence coverage [%] | Unique sequence coverage [%] | Mol. weight [kDa] |
|---|---|---|---|---|---|---|---|---|---|---|---|---|---|---|---|---|---|---|
| A0A0H3NC32 | 26.7622 | 26.3219 | 26.5256 | 26.9365 | 26.5613 | 27.3493 | FlaE;Flg_bb_rod;Flg_bbr_C | Flagellar hook protein FlgE {ECO:0000256\|RuleBase:RU362116} | | 0.724093275 | 0.48475938 | 0.412491481 | | 5 | 5 | 13.2 | 13.2 | 42.21 |
| A0A0H3NI81 | 27.4886 | 27.4128 | 24.1576 | 27.5535 | 24.8457 | 24.2022 | EPSP_synthase | UDP-N-acetylglucosamine 1-carboxyvinyltransferase {ECO:0000256\|HAMAP-Rule:MF_00111} | | 0.21130918 | 0.48528319 | -0.819207509 | | 5 | 5 | 14.3 | 14.3 | 44.74 |
| A0A0H3NHL9 | 28.8369 | 29.0635 | 28.8586 | 29.2688 | 29.2904 | 29.2563 | EIID-AGA | | | 2.073532307 | 0.48540444 | 0.352140427 | | 10 | 10 | 26.9 | 26.9 | 31.26 |
| A0A0H3NLH5 | 30.8464 | 30.7825 | 30.7447 | 30.4732 | 30.4427 | 30.4504 | Response_reg;Trans_reg_C | | | 3.381623656 | 0.48545934 | -0.335751851 | | 14 | 14 | 52.9 | 52.9 | 27.29 |
| A0A0H3NIK3 | 25.9703 | 24.5246 | 24.7043 | 24.2189 | 24.6897 | 24.8261 | Pentapeptide_4;SopA;SopA_C | | | 0.424694013 | 0.48558758 | -0.488164266 | | 3 | 3 | 5.1 | 5.1 | 86.78 |
| A0A0H3NBC6 | 27.8071 | 27.4559 | 25.2089 | 27.4999 | 27.5051 | 27.2405 | AAA;AAA_2;Clp_N;ClpB_D2-small | | | 0.29233487 | 0.48564912 | 0.591192245 | | 9 | 9 | 13.5 | 13.5 | 84.04 |
| A0A0H3NI01 | 30.3428 | 30.2839 | 30.2656 | 29.7011 | 30.048 | 30.0423 | OEP | | | 1.455954598 | 0.48620935 | -0.366942724 | | 24 | 24 | 42.8 | 42.8 | 53.69 |
| A0A0H3NIL5 | 32.223 | 32.2285 | 32.3724 | 32.0673 | 31.934 | 31.7566 | Ribosomal_L1 | 50S ribosomal protein L1 {ECO:0000256\|HAMAP-Rule:MF_01318} | | 1.591638159 | 0.49097593 | -0.355350494 | | 13 | 13 | 51.7 | 51.7 | 24.73 |
| A0A0H3NKP5 | 30.2653 | 30.3437 | 30.4405 | 30.5861 | 30.6708 | 30.8225 | Bac_DNA_binding | Integration host factor subunit alpha {ECO:0000256\|RuleBase:RU004485, ECO:0000256\|SAAS:SAAS00084680} | | 1.794298329 | 0.4938961 | 0.343341192 | | 8 | 8 | 65 | 65 | 11.76 |

### All proteins

| Uniprot accession number | LFQ intensity WT_1 | LFQ intensity WT_2 | LFQ intensity WT_3 | LFQ intensity OrfSwap_1 | LFQ intensity OrfSwap_2 | LFQ intensity OrfSwap_3 | Pfam name | Uniprot full protein name | Student's T-test Significant OrfSwap_WT | -Log Student's T-test p-value OrfSwap_WT | Student's T-test q-value OrfSwap_WT | Student's T-test Difference OrfSwap_WT | Cluster | Peptides | Unique peptides | Sequence coverage [%] | Unique sequence coverage [%] | Mol. weight [kDa] |
|---|---|---|---|---|---|---|---|---|---|---|---|---|---|---|---|---|---|---|
| A0A0H3ND82 | 27.5946 | 28.3741 | 28.3324 | 28.4748 | 28.382 | 28.6587 | tRNA_bind;tRNA-synt_1g | Methionine--tRNA ligase {ECO:0000256\|HAMAP-Rule:MF_00098} | | 0.693548965 | 0.49404793 | 0.404878616 | | 12 | 12 | 19.5 | 19.5 | 76.27 |
| A0A0H3NJU7 | 30.5939 | 30.6695 | 30.3958 | 30.0503 | 30.2192 | 30.3289 | ketoacyl-synt;Ketoacyl-synt_C | | | 1.430337728 | 0.4940564 | -0.353614171 | | 13 | 13 | 27.7 | 27.7 | 42.37 |
| A0A0H3N8W7 | 27.2227 | 27.1875 | 27.0141 | 25.3452 | 27.2824 | 27.2308 | LptE | LPS-assembly lipoprotein LptE {ECO:0000256\|HAMAP-Rule:MF_01186, ECO:0000256\|SAAS:SAAS00963593} | | 0.336428871 | 0.49437391 | -0.522002538 | | 4 | 4 | 33.2 | 33.2 | 21.46 |
| A0A0H3NAW8 | 30.34 | 30.5536 | 30.6259 | 30.0615 | 30.1438 | 30.2711 | PK;PK_C | Pyruvate kinase {ECO:0000256\|RuleBase:RU000504} | | 1.524965299 | 0.49530886 | -0.347717285 | | 19 | 19 | 38.3 | 38.3 | 48.65 |
| A0A0H3NDK2 | 29.9244 | 29.8875 | 29.7436 | 29.3936 | 29.4827 | 29.6542 | ATP-cone;Ribonuc_red_lgC;Ribonuc_red_lgN | Ribonucleoside-diphosphate reductase {ECO:0000256\|RuleBase:RU003410} | | 1.651952654 | 0.49562151 | -0.341632843 | | 36 | 36 | 43.9 | 43.9 | 85.74 |

### All proteins

| Uniprot accession number | LFQ intensity WT_1 | LFQ intensity WT_2 | LFQ intensity WT_3 | LFQ intensity OrfSwap_1 | LFQ intensity OrfSwap_2 | LFQ intensity OrfSwap_3 | Pfam name | Uniprot full protein name | Student's T-test Significant OrfSwap_WT | -Log Student's T-test p-value OrfSwap_WT | Student's T-test q-value OrfSwap_WT | Student's T-test Difference OrfSwap_WT | Cluster | Peptides | Unique peptides | Sequence coverage [%] | Unique sequence coverage [%] | Mol. weight [kDa] |
|---|---|---|---|---|---|---|---|---|---|---|---|---|---|---|---|---|---|---|
| A0A0H3N9I3 | 25.7438 | 24.9588 | 25.4718 | 26.0708 | 25.4967 | 25.8139 | Radical_SAM;TRAM;UPF0004 | tRNA-2-methylthio-N(6)-dimethylallyladenosine synthase {ECO:0000256\|HAMAP-Rule:MF_01864, ECO:0000256\|SAAS:SAAS00623688} | | 0.639830464 | 0.49615517 | 0.402385712 | | 5 | 5 | 9.1 | 9.1 | 53.72 |
| A0A0H3NHH4 | 24.9756 | 25.7707 | 25.2975 | 25.633 | 24.939 | 24.0607 | Guanylate_kin | Guanylate kinase {ECO:0000256\|HAMAP-Rule:MF_00328, ECO:0000256\|SAAS:SAAS00152752} | | 0.388578932 | 0.49626609 | -0.470362981 | | 2 | 2 | 10.6 | 10.6 | 23.5 |
| A0A0H3NHB9 | 28.8888 | 29.121 | 28.9497 | 29.1602 | 29.528 | 29.3066 | GST_C;GST_N_3 | | | 1.269703087 | 0.49856237 | 0.345087687 | | 7 | 7 | 32.2 | 32.2 | 22.57 |
| A0A0H3NF14 | 29.8539 | 29.8579 | 30.1055 | 29.668 | 29.7041 | 29.4131 | Aminotran_3 | | | 1.302700706 | 0.49885593 | -0.34402593 | | 22 | 22 | 53.2 | 53.2 | 45.64 |
| A0A0H3NK71 | 29.7034 | 29.5928 | 29.6802 | 29.4385 | 29.2722 | 29.2897 | Pyr_redox_2;Pyr_redox_dim | Soluble pyridine nucleotide transhydrogenase {ECO:0000256\|HAMAP-Rule:MF_00247} | | 2.185164322 | 0.49991507 | -0.325329463 | | 13 | 13 | 39.7 | 39.7 | 51.61 |

### All proteins

| Uniprot accession number | LFQ intensity WT_1 | LFQ intensity WT_2 | LFQ intensity WT_3 | LFQ intensity OrfSwap_1 | LFQ intensity OrfSwap_2 | LFQ intensity OrfSwap_3 | Pfam name | Uniprot full protein name | Student's T-test Significant OrfSwap_WT | -Log Student's T-test p-value OrfSwap_WT | Student's T-test q-value OrfSwap_WT | Student's T-test Difference OrfSwap_WT | Cluster | Peptides | Unique peptides | Sequence coverage [%] | Unique sequence coverage [%] | Mol. weight [kDa] |
|---|---|---|---|---|---|---|---|---|---|---|---|---|---|---|---|---|---|---|
| A0A0H3NIU7 | 30.8404 | 30.5089 | 30.9749 | 31.1953 | 31.0182 | 31.1684 | ACAS_N;AMP-binding;AMP-binding_C | Acetyl-coenzyme A synthetase {ECO:0000256\|HAMAP-Rule:MF_01123} | | 1.112291305 | 0.5000766 | 0.352527618 | | 40 | 40 | 67.6 | 67.6 | 72.15 |
| A0A0H3NFE6 | 24.4798 | 26.362 | 26.8404 | 27.1801 | 27.3664 | 25.0183 | DnaJ;DnaJ_C | Curved DNA-binding protein {ECO:0000256\|HAMAP-Rule:MF_01154, ECO:0000256\|SAAS:SAAS00724453} | | 0.236799793 | 0.50027292 | 0.627510071 | | 7 | 7 | 25.5 | 25.5 | 34.69 |
| A0A0H3NHN2 | 32.3472 | 32.2009 | 32.2341 | 31.9392 | 31.8343 | 32.0303 | PGK | Phosphoglycerate kinase {ECO:0000256\|HAMAP-Rule:MF_00145, ECO:0000256\|RuleBase:RU000532, ECO:0000256\|SAAS:SAAS00041088} | | 1.977662924 | 0.50031092 | -0.326096217 | | 26 | 26 | 60.5 | 60.5 | 41.13 |

### All proteins

| Uniprot accession number | LFQ intensity WT_1 | LFQ intensity WT_2 | LFQ intensity WT_3 | LFQ intensity OrfSwap_1 | LFQ intensity OrfSwap_2 | LFQ intensity OrfSwap_3 | Pfam name | Uniprot full protein name | Student's T-test Significant OrfSwap_WT | -Log Student's T-test p-value OrfSwap_WT | Student's T-test q-value OrfSwap_WT | Student's T-test Difference OrfSwap_WT | Cluster | Peptides | Unique peptides | Sequence coverage [%] | Unique sequence coverage [%] | Mol. weight [kDa] |
|---|---|---|---|---|---|---|---|---|---|---|---|---|---|---|---|---|---|---|
| A0A0H3NH94 | 25.5215 | 25.908 | 25.4876 | 25.0049 | 25.6243 | 25.1524 | Glyco_tran_28_C;Glyco_transf_28 | UDP-N-acetylglucosamine--N-acetylmuramyl-(pentapeptide) pyrophosphoryl-undecaprenol N-acetylglucosamine transferase {ECO:0000256\|HAMAP-Rule:MF_00033, ECO:0000256\|SAAS:SAAS00765687} | | 0.755090836 | 0.50084615 | -0.37852033 | | 5 | 5 | 15.2 | 15.2 | 37.86 |
| A0A0H3N7V0 | 29.7046 | 29.6098 | 29.5099 | 29.2895 | 29.3198 | 29.247 | Lipoprotein_9 | Lipoprotein {ECO:0000256\|PIRNR:PIRNR002854} | | 2.236567098 | 0.50136421 | -0.322648366 | | 13 | 13 | 50.9 | 50.9 | 29.44 |
| A0A0H3NJK9 | 25.2637 | 25.2742 | 25.3518 | 25.0646 | 24.9434 | 24.9072 | ABC_membrane;ABC_tran | | | 2.382747169 | 0.50143897 | -0.324887594 | | 2 | 2 | 3.1 | 3.1 | 62.99 |
| A0A0H3NHF3 | 26.811 | 27.0606 | 24.3591 | 26.5786 | 26.477 | 26.8848 | Pribosyltran | Orotate phosphoribosyltransferase {ECO:0000256\|HAMAP-Rule:MF_01208, ECO:0000256\|SAAS:SAAS00615437} | | 0.260860235 | 0.50242194 | 0.569905599 | | 5 | 5 | 18.3 | 18.3 | 23.56 |

### All proteins

| Uniprot accession number | LFQ intensity WT_1 | LFQ intensity WT_2 | LFQ intensity WT_3 | LFQ intensity OrfSwap_1 | LFQ intensity OrfSwap_2 | LFQ intensity OrfSwap_3 | Pfam name | Uniprot full protein name | Student's T-test Significant OrfSwap_WT | -Log Student's T-test p-value OrfSwap_WT | Student's T-test q-value OrfSwap_WT | Student's T-test Difference OrfSwap_WT | Cluster | Peptides | Unique peptides | Sequence coverage [%] | Unique sequence coverage [%] | Mol. weight [kDa] |
|---|---|---|---|---|---|---|---|---|---|---|---|---|---|---|---|---|---|---|
| A0A0H3NG32 | 27.5397 | 27.4414 | 27.0481 | 27.1592 | 26.5062 | 27.2131 | DUF1190 | UPF0441 protein YgiB {ECO:0000256\|HAMAP-Rule:MF_01188} | | 0.634937523 | 0.50485535 | -0.383595785 | | 4 | 4 | 21.5 | 21.5 | 23.39 |
| A0A0H3NAK6 | 30.1143 | 29.93 | 29.9289 | 29.5371 | 29.8086 | 29.6342 | Glutaredoxin2_C;GST_N_3 | | | 1.521298959 | 0.50520502 | -0.33108902 | | 16 | 16 | 67.9 | 67.9 | 24.46 |
| A0A0H3NGS5 | 32.2547 | 32.3731 | 32.4545 | 31.8226 | 32.1112 | 32.1445 | Ribosomal_L4 | 50S ribosomal protein L4 {ECO:0000256\|HAMAP-Rule:MF_01328} | | 1.33247664 | 0.50687265 | -0.334659576 | | 19 | 19 | 67.2 | 67.2 | 22.09 |
| A0A0H3NGB2 | 27.9044 | 27.7668 | 27.4369 | 28.078 | 28.1148 | 27.9387 | Lum_binding | | | 1.078296488 | 0.51273333 | 0.341130575 | | 8 | 8 | 34.7 | 34.7 | 23.39 |
| A0A0H3NDW4 | 29.5599 | 29.957 | 29.6336 | 29.4803 | 29.3861 | 29.2826 | PK;PK_C | Pyruvate kinase {ECO:0000256\|RuleBase:RU000504} | | 1.165708618 | 0.51300207 | -0.333822886 | | 19 | 19 | 39.8 | 39.8 | 51.39 |
| A0A0H3NP62 | 24.7261 | 25.3577 | 25.5512 | 25.2465 | 24.9926 | 24.1397 | MTS | 50S ribosomal protein L3 glutamine methyltransferase {ECO:0000256\|HAMAP-Rule:MF_02125} | | 0.428993307 | 0.51406237 | -0.418738683 | | 3 | 3 | 11.3 | 11.3 | 35.06 |
| A0A0H3NCZ1 | 32.9464 | 32.7999 | 32.9681 | 32.638 | 32.6522 | 32.4726 | Ribosomal_S2 | 30S ribosomal protein S2 {ECO:0000256\|HAMAP-Rule:MF_00291} | | 1.812872625 | 0.51406639 | -0.31722641 | | 28 | 28 | 68.5 | 68.5 | 26.76 |
| A0A0H3N9E2 | 26.775 | 24.6099 | 25.8556 | 25.0285 | 26.1676 | 24.4591 | CSD | | | 0.262239345 | 0.51445455 | -0.528481166 | | 4 | 4 | 52.9 | 52.9 | 7.583 |
| A0A0H3NHA5 | 27.6724 | 25.8367 | 27.8664 | 26.4877 | 28.0865 | 24.9712 | OmpW | | | 0.213859905 | 0.51963711 | -0.610064189 | | 2 | 2 | 9.9 | 9.9 | 22.96 |
| A0A0H3NBY2 | 31.505 | 31.6605 | 31.7031 | 31.3392 | 31.2286 | 31.3735 | SBP_bac_5 | | | 1.847548955 | 0.5201893 | -0.309071859 | | 28 | 28 | 61 | 61 | 61.29 |

#### All proteins

| Uniprot accession number | LFQ intensity WT_1 | LFQ intensity WT_2 | LFQ intensity WT_3 | LFQ intensity OrfSwap _1 | LFQ intensity OrfSwap _2 | LFQ intensity OrfSwap _3 | Pfam name | Uniprot full protein name | Student's T-test Significant OrfSwap_WT | -Log Student's T-test p-value OrfSwap_WT | Student's T-test q-value OrfSwap_WT | Student's T-test Difference OrfSwap_WT | Cluster | Peptides | Unique peptides | Sequence coverage [%] | Unique sequence coverage [%] | Mol. weight [kDa] |
|---|---|---|---|---|---|---|---|---|---|---|---|---|---|---|---|---|---|---|
| A0A0H3N9S4 | 28.584 | 28.4189 | 28.5453 | 28.3268 | 28.2938 | 27.9853 | Lyase_aromatic | Histidine ammonia-lyase {ECO:0000256\|HAMAP-Rule:MF_00229, ECO:0000256\|RuleBase:RU004479, ECO:0000256\|SAAS:SAAS00831227} | | 1.233345131 | 0.52678615 | -0.314107259 | | 13 | 13 | 31.4 | 31.4 | 53.84 |
| A0A0H3NU72 | 27.8506 | 27.8758 | 28.0001 | 27.5558 | 28.6611 | 28.6658 | ATP-synt_DE;ATP-synt_DE_N | ATP synthase epsilon chain {ECO:0000256\|HAMAP-Rule:MF_00530} | | 0.445105649 | 0.52720816 | 0.38538297 | | 5 | 5 | 23.7 | 23.7 | 15.06 |
| A0A0H3NNW3 | 27.5724 | 27.3873 | 28.0198 | 27.4257 | 27.3992 | 27.1278 | peroxidase | Catalase-peroxidase {ECO:0000256\|HAMAP-Rule:MF_01961, ECO:0000256\|RuleBase:RU003451} | | 0.746426783 | 0.52757377 | -0.34226799 | | 10 | 10 | 15.4 | 15.4 | 79.66 |
| A0A0H3NAC8 | 27.456 | 27.4866 | 28.1056 | 27.3126 | 27.3337 | 27.3719 | ASL_C;Lyase_1 | Adenylosuccinate lyase {ECO:0000256\|RuleBase:RU361172} | | 0.74191419 | 0.52795072 | -0.343343099 | | 11 | 11 | 21.5 | 21.5 | 51.55 |

#### All proteins

| Uniprot accession number | LFQ intensity WT_1 | LFQ intensity WT_2 | LFQ intensity WT_3 | LFQ intensity OrfSwap_1 | LFQ intensity OrfSwap_2 | LFQ intensity OrfSwap_3 | Pfam name | Uniprot full protein name | Student's T-test Significant OrfSwap_WT | -Log Student's T-test p-value OrfSwap_WT | Student's T-test q-value OrfSwap_WT | Student's T-test Difference OrfSwap_WT | Cluster | Peptides | Unique peptides | Sequence coverage [%] | Unique sequence coverage [%] | Mol. weight [kDa] |
|---|---|---|---|---|---|---|---|---|---|---|---|---|---|---|---|---|---|---|
| A0A0H3NCF5 | 28.711 | 29.0179 | 28.9886 | 28.5588 | 28.8845 | 28.2583 | NAD_synthase | NH(3)-dependent NAD(+) synthetase {ECO:0000256\|HAMAP-Rule:MF_00193, ECO:0000256\|RuleBase:RU003812, ECO:0000256\|SAAS:SAAS00694129} | | 0.757477907 | 0.5282863 | -0.338647207 | | 11 | 11 | 42.5 | 42.5 | 30.48 |
| A0A0H3NEC9 | 26.9601 | 26.4211 | 24.2388 | 26.1755 | 26.6444 | 26.3396 | DUF4115 | Cytoskeleton protein RodZ {ECO:0000256\|HAMAP-Rule:MF_02017, ECO:0000256\|SAAS:SAAS00892895} | | 0.23981901 | 0.53076673 | 0.513157527 | | 7 | 7 | 24 | 24 | 35.66 |
| A0A0H3NPA9 | 30.0985 | 29.3624 | 29.3355 | 29.374 | 29.026 | 29.3323 | PTS-HPr | | | 0.57887334 | 0.53165041 | -0.35467275 | | 7 | 7 | 88.2 | 88.2 | 9.119 |
| A0A0H3NMB1 | 33.1149 | 33.1179 | 33.2371 | 32.7776 | 32.8151 | 33.0014 | Ldh_1_C;Ldh_1_N | Malate dehydrogenase {ECO:0000256\|HAMAP-Rule:MF_01516, ECO:0000256\|RuleBase:RU004066, ECO:0000256\|SAAS:SAAS00064143} | | 1.660888416 | 0.536384 | -0.291900635 | | 24 | 24 | 62.5 | 62.5 | 32.48 |
| A0A0H3NBW6 | 31.1311 | 31.2938 | 30.8495 | 30.4522 | 31.1029 | 30.707 | Mn_catalase | | | 0.666576691 | 0.53679352 | -0.337415695 | | 13 | 13 | 44.2 | 44.2 | 31.85 |

### All proteins

| Uniprot accession number | LFQ intensity WT_1 | LFQ intensity WT_2 | LFQ intensity WT_3 | LFQ intensity OrfSwap_1 | LFQ intensity OrfSwap_2 | LFQ intensity OrfSwap_3 | Pfam name | Uniprot full protein name | Student's T-test Significant OrfSwap_WT | -Log Student's T-test p-value OrfSwap_WT | Student's T-test q-value OrfSwap_WT | Student's T-test Difference OrfSwap_WT | Cluster | Peptides | Unique peptides | Sequence coverage [%] | Unique sequence coverage [%] | Mol. weight [kDa] |
|---|---|---|---|---|---|---|---|---|---|---|---|---|---|---|---|---|---|---|
| A0A0H3NGS0 | 32.9575 | 32.9095 | 33.0406 | 32.6261 | 32.7388 | 32.6896 | KH_2;Ribosomal_S3_C | 30S ribosomal protein S3 {ECO:0000256\|HAMAP-Rule:MF_01309} | | 2.31655548 | 0.53695391 | -0.28438441 | | 29 | 29 | 88.4 | 88.4 | 25.98 |
| A0A0H3NEA5 | 28.8196 | 28.3237 | 28.3886 | 28.0334 | 28.0773 | 28.4426 | HGTP_anticodon | Histidine--tRNA ligase {ECO:0000256\|HAMAP-Rule:MF_00127} | | 0.738273466 | 0.53734137 | -0.326145172 | | 12 | 12 | 40.3 | 40.3 | 46.93 |
| A0A0H3NDQ5 | 29.0579 | 28.9635 | 29.5036 | 28.8431 | 28.8415 | 28.884 | Oxidored_q6 | NADH-quinone oxidoreductase subunit B {ECO:0000256\|HAMAP-Rule:MF_01356} | | 0.888888744 | 0.5373629 | -0.318779627 | | 8 | 8 | 35 | 35 | 25.09 |
| A0A0H3N8I8 | 27.6029 | 27.3198 | 27.8847 | 28.4576 | 27.7218 | 27.6933 | 5_nucleotid_C;Metallophos | | | 0.522721108 | 0.53793939 | 0.355143865 | | 10 | 10 | 18.2 | 18.2 | 60.44 |
| A0A0H3NQ01 | 25.951 | 25.2177 | 24.8878 | 25.6359 | 25.8792 | 25.6225 | HsdM_N;N6_Mtase | | | 0.481796992 | 0.53798793 | 0.360378901 | | 6 | 6 | 12.9 | 12.9 | 59.33 |
| A0A0H3NFA1 | 31.4317 | 31.7693 | 31.3679 | 31.2713 | 31.1562 | 31.2509 | DUF883 | | | 1.077606385 | 0.54051866 | -0.296854655 | | 10 | 10 | 38.8 | 38.8 | 11.64 |
| A0A0H3NGZ8 | 26.4929 | 26.669 | 26.3717 | 26.8321 | 26.3081 | 25.1976 | ATP-synt_ab;ATP-synt_ab_N | | | 0.336135643 | 0.5407505 | -0.398608526 | | 6 | 6 | 16.7 | 16.7 | 47.61 |
| A0A0H3N9H9 | 31.5758 | 31.5565 | 31.5995 | 31.9014 | 31.7326 | 31.9384 | SBP_bac_3 | | | 1.912422033 | 0.54140157 | 0.280209223 | | 27 | 27 | 69.5 | 69.5 | 34.09 |
| A0A0H3NHH6 | 30.5029 | 30.2896 | 30.2625 | 29.9757 | 30.1842 | 30.0267 | Trehalase | Periplasmic trehalase {ECO:0000256\|HAMAP-Rule:MF_01060} | | 1.370864631 | 0.54225641 | -0.289454142 | | 17 | 17 | 27 | 27 | 63.56 |

### All proteins

| Uniprot accession number | LFQ intensity WT_1 | LFQ intensity WT_2 | LFQ intensity WT_3 | LFQ intensity OrfSwap_1 | LFQ intensity OrfSwap_2 | LFQ intensity OrfSwap_3 | Pfam name | Uniprot full protein name | Student's T-test Significant OrfSwap_WT | -Log Student's T-test p-value OrfSwap_WT | Student's T-test q-value OrfSwap_WT | Student's T-test Difference OrfSwap_WT | Cluster | Peptides | Unique peptides | Sequence coverage [%] | Unique sequence coverage [%] | Mol. weight [kDa] |
|---|---|---|---|---|---|---|---|---|---|---|---|---|---|---|---|---|---|---|
| A0A0H3NC19 | 29.1757 | 29.1602 | 28.6556 | 28.7773 | 28.5277 | 28.7466 | MMR_HSR1; YchF-GTPase_C | Ribosome-binding ATPase YchF {ECO:0000256\|HAMAP-Rule:MF_00944} | | 0.767041775 | 0.5427747 | -0.313322703 | | 13 | 13 | 43.8 | 43.8 | 39.63 |
| A0A0H3NEL4 | 30.5566 | 30.6101 | 30.7343 | 31.368 | 30.8898 | 30.621 | Gly_radical | Autonomous glycyl radical cofactor {ECO:0000256\|HAMAP-Rule:MF_00806, ECO:0000256\|SAAS:SAAS00058385} | | 0.656409804 | 0.54281275 | 0.325878143 | | 13 | 12 | 91.3 | 91.3 | 14.34 |
| A0A0H3N9L3 | 30.0436 | 29.8009 | 30.2795 | 30.5439 | 30.414 | 30.1097 | M20_dimer; Peptidase_M20 | | | 0.766869259 | 0.54316832 | 0.314517975 | | 14 | 14 | 34.4 | 34.4 | 52.44 |
| A0A0H3NDF6 | 26.4353 | 26.4834 | 26.6064 | 26.8169 | 26.9139 | 26.6602 | ALAD | Delta-aminolevulinic acid dehydratase {ECO:0000256\|RuleBase:RU000515} | | 1.489057431 | 0.54331746 | 0.28864034 | | 5 | 5 | 20.7 | 20.7 | 35.55 |
| A0A0H3P010 | 24.3162 | 26.1456 | 26.0773 | 25.381 | 25.3267 | 24.5079 | Transposase_31 | | | 0.266278172 | 0.54352286 | -0.441139857 | | 6 | 6 | 21.4 | 21.4 | 35.32 |
| A0A0H3NKR1 | 31.9238 | 31.8276 | 32.0099 | 32.0296 | 32.2467 | 32.3398 | Ribosomal_L19 | 50S ribosomal protein L19 {ECO:0000256\|HAMAP-Rule:MF_00402, ECO:0000256\|RuleBase:RU000559} | | 1.262609684 | 0.55183529 | 0.284978867 | | 19 | 19 | 81.7 | 81.7 | 13.13 |

### All proteins

| Uniprot accession number | LFQ intensity WT_1 | LFQ intensity WT_2 | LFQ intensity WT_3 | LFQ intensity OrfSwap_1 | LFQ intensity OrfSwap_2 | LFQ intensity OrfSwap_3 | Pfam name | Uniprot full protein name | Student's T-test Significant OrfSwap_WT | -Log Student's T-test p-value OrfSwap_WT | Student's T-test q-value OrfSwap_WT | Student's T-test Difference OrfSwap_WT | Cluster | Peptides | Unique peptides | Sequence coverage [%] | Unique sequence coverage [%] | Mol. weight [kDa] |
|---|---|---|---|---|---|---|---|---|---|---|---|---|---|---|---|---|---|---|
| A0A0H3NSC0 | 25.554 | 25.602 | 24.7296 | 24.8755 | 26.1064 | 26.0453 | GTP1_OBG; MMR_HSR1 | GTPase Obg {ECO:0000256\|HAMAP-Rule:MF_01454} | | 0.317725995 | 0.55614873 | 0.380566915 | | 2 | 2 | 5.9 | 5.9 | 43.1 |
| A0A0H3NDN2 | 25.857 | 24.5737 | 24.775 | 25.2991 | 24.87 | 26.2233 | SpoVR | | | 0.282461667 | 0.55985156 | 0.395547231 | | 4 | 4 | 9.8 | 9.8 | 60.54 |
| A0A0H3NFQ6 | 32.7406 | 32.9229 | 32.7596 | 32.4746 | 32.4026 | 32.7062 | Iso_dh | Isocitrate dehydrogenase [NADP] {ECO:0000256\|RuleBase:RU004446} | | 1.214100873 | 0.55985965 | -0.279916128 | | 31 | 31 | 55.5 | 55.5 | 45.79 |
| A0A0H3NM77 | 29.4067 | 28.9552 | 28.884 | 29.5388 | 28.2591 | 28.3598 | KH_5;NusA_N | Transcription termination/antitermination protein NusA {ECO:0000256\|HAMAP-Rule:MF_00945} | | 0.338976444 | 0.56128405 | -0.362711589 | | 14 | 14 | 25.4 | 25.4 | 55.43 |
| A0A0H3NFZ2 | 26.6071 | 24.4634 | 26.0245 | 24.6678 | 25.5745 | 25.5697 | NiFeSe_Hases | | | 0.23799449 | 0.56284272 | -0.427640915 | | 4 | 4 | 10.9 | 10.9 | 62.44 |
| A0A0H3NVR3 | 32.5416 | 32.5808 | 32.7243 | 33.0888 | 32.896 | 32.6991 | FumaraseC_C;Lyase_1 | Aspartate ammonia-lyase {ECO:0000256\|RuleBase:RU362017} | | 1.044887865 | 0.56381467 | 0.279062907 | | 35 | 35 | 68 | 68 | 52.29 |
| A0A0H3NDA5 | 25.6134 | 28.1059 | 27.8834 | 27.7725 | 27.6366 | 27.5322 | DUF1100 | Esterase FrsA {ECO:0000256\|HAMAP-Rule:MF_01063} | | 0.217159072 | 0.56490522 | 0.446171443 | | 9 | 9 | 22.9 | 22.9 | 47.16 |
| A0A0H3N8A6 | 28.3676 | 28.0619 | 28.5528 | 27.7546 | 28.3712 | 27.9399 | YajC | | | 0.587856626 | 0.566 | -0.305545171 | | 5 | 5 | 30 | 30 | 11.84 |

### All proteins

| Uniprot accession number | LFQ intensity WT_1 | LFQ intensity WT_2 | LFQ intensity WT_3 | LFQ intensity OrfSwap_1 | LFQ intensity OrfSwap_2 | LFQ intensity OrfSwap_3 | Pfam name | Uniprot full protein name | Student's T-test Significant OrfSwap_WT | -Log Student's T-test p-value OrfSwap_WT | Student's T-test q-value OrfSwap_WT | Student's T-test Difference OrfSwap_WT | Cluster | Peptides | Unique peptides | Sequence coverage [%] | Unique sequence coverage [%] | Mol. weight [kDa] |
|---|---|---|---|---|---|---|---|---|---|---|---|---|---|---|---|---|---|---|
| A0A0H3N8Z5 | 27.1196 | 27.2223 | 26.7915 | 27.2385 | 27.5518 | 27.2076 | LptD;OstA | LPS-assembly protein LptD {ECO:0000256\|HAMAP-Rule:MF_01411, ECO:0000256\|SAAS:SAAS00073013} | | 0.780626539 | 0.56675915 | 0.288120906 | | 13 | 13 | 22.8 | 22.8 | 89.82 |
| A0A0H3NI82 | 35.528 | 35.516 | 35.6908 | 35.3091 | 35.3545 | 35.2927 | FGGY_C;FGGY_N | Glycerol kinase {ECO:0000256\|HAMAP-Rule:MF_00186} | | 1.923259758 | 0.56826154 | -0.259525299 | | 53 | 53 | 73.9 | 73.9 | 56.05 |
| A0A0H3NH87 | 24.8703 | 25.5028 | 25.4567 | 26.0649 | 25.4577 | 25.2642 | Mur_ligase;Mur_ligase_C;Mur_ligase_M | UDP-N-acetylmuramoyl-L-alanyl-D-glutamate--2,6-diaminopimelate ligase {ECO:0000256\|HAMAP-Rule:MF_00208} | | 0.432505876 | 0.57013793 | 0.318966548 | | 5 | 5 | 11.9 | 11.9 | 53.29 |
| A0A0H3NU77 | 30.9222 | 30.8733 | 30.7368 | 30.4293 | 30.6938 | 30.6106 | ATP-synt_B | ATP synthase subunit b {ECO:0000256\|HAMAP-Rule:MF_01398} | | 1.302385071 | 0.57080998 | -0.26620992 | | 22 | 22 | 68.6 | 68.6 | 17.37 |
| A0A0H3ND92 | 27.1442 | 27.6721 | 28.0206 | 27.5502 | 28.363 | 27.8993 | OpuAC | | | 0.396300866 | 0.57147992 | 0.325215658 | | 7 | 7 | 19.3 | 19.3 | 32.76 |
| A0A0H3P198 | 26.4916 | 26.1384 | 24.0566 | 25.4077 | 25.5325 | 24.4399 | ProQ | RNA chaperone ProQ {ECO:0000256\|SAAS:SAAS00876659} | | 0.201132221 | 0.57234652 | -0.435484568 | | 1 | 1 | 11 | 11 | 22.46 |

### All proteins

| Uniprot accession number | LFQ intensity WT_1 | LFQ intensity WT_2 | LFQ intensity WT_3 | LFQ intensity OrfSwap_1 | LFQ intensity OrfSwap_2 | LFQ intensity OrfSwap_3 | Pfam name | Uniprot full protein name | Student's T-test Significant OrfSwap_WT | -Log Student's T-test p-value OrfSwap_WT | Student's T-test q-value OrfSwap_WT | Student's T-test Difference OrfSwap_WT | Cluster | Peptides | Unique peptides | Sequence coverage [%] | Unique sequence coverage [%] | Mol. weight [kDa] |
|---|---|---|---|---|---|---|---|---|---|---|---|---|---|---|---|---|---|---|
| A0A0H3NMW9 | 27.1909 | 27.388 | 27.5513 | 27.8391 | 27.3619 | 27.7713 | PTS_EIIA_2; PTS_EIIC;PTS_IIB | | | 0.704375139 | 0.57246038 | 0.280719757 | | 12 | 12 | 20.4 | 20.4 | 68.16 |
| A0A0H3NDR3 | 31.1496 | 30.9038 | 31.2273 | 31.0919 | 30.7121 | 30.6453 | DAO;Fer2_BFD | Glycerol-3-phosphate dehydrogenase {ECO:0000256\|RuleBase:RU361217} | | 0.749511437 | 0.57254135 | -0.277175268 | | 34 | 34 | 56.6 | 56.6 | 59.03 |
| A0A0H3NBT7 | 28.4347 | 28.4969 | 28.3853 | 27.7023 | 28.4786 | 28.2562 | DUF3313 | | | 0.55795539 | 0.57297543 | -0.293275833 | | 2 | 2 | 12.6 | 12.6 | 24.44 |
| A0A0H3NBY3 | 31.5978 | 31.6725 | 31.5155 | 31.8277 | 31.6402 | 32.1414 | Usp | | | 0.831416628 | 0.5735 | 0.274446487 | | 15 | 15 | 75.7 | 75.7 | 15.71 |
| A0A0H3NKZ8 | 28.8591 | 29.0826 | 28.8912 | 28.7163 | 28.8932 | 28.392 | Aldedh | | | 0.7867754 | 0.57417078 | -0.277066549 | | 19 | 19 | 50.2 | 50.2 | 51.85 |
| A0A0H3NGQ1 | 31.7492 | 32.3431 | 32.1879 | 31.7474 | 31.8536 | 31.8433 | Ribosomal_L27A | 50S ribosomal protein L15 {ECO:0000256\|HAMAP-Rule:MF_01341, ECO:0000256\|RuleBase:RU003889, ECO:0000256\|SAAS:SAAS00402269} | | 0.70191181 | 0.57428893 | -0.278601329 | | 15 | 15 | 64.6 | 64.6 | 14.97 |
| A0A0H3NIN9 | 31.0967 | 31.1331 | 30.8894 | 30.9247 | 30.8934 | 30.4626 | ICL | | | 0.769217797 | 0.57472 | -0.279493968 | | 23 | 23 | 56.7 | 56.7 | 47.56 |
| A0A0H3NHN3 | 27.0276 | 27.4854 | 26.8591 | 27.2759 | 27.4344 | 27.5226 | HTH_IclR;IclR | | | 0.646311487 | 0.5748365 | 0.286909739 | | 8 | 8 | 35.4 | 35.4 | 29.86 |
| A0A0H3N9D6 | 32.9912 | 33.2807 | 33.1001 | 33.3228 | 33.5274 | 33.3193 | Ferritin | DNA protection during starvation protein {ECO:0000256\|HAMAP-Rule:MF_01441, ECO:0000256\|SAAS:SAAS00760410} | | 1.147986372 | 0.57542748 | 0.265858968 | | 23 | 23 | 86.8 | 86.8 | 18.72 |

### All proteins

| Uniprot accession number | LFQ intensity WT_1 | LFQ intensity WT_2 | LFQ intensity WT_3 | LFQ intensity OrfSwap_1 | LFQ intensity OrfSwap_2 | LFQ intensity OrfSwap_3 | Pfam name | Uniprot full protein name | Student's T-test Significant OrfSwap_WT | -Log Student's T-test p-value OrfSwap_WT | Student's T-test q-value OrfSwap_WT | Student's T-test Difference OrfSwap_WT | Cluster | Peptides | Unique peptides | Sequence coverage [%] | Unique sequence coverage [%] | Mol. weight [kDa] |
|---|---|---|---|---|---|---|---|---|---|---|---|---|---|---|---|---|---|---|
| A0A0H3NCT0 | 31.7248 | 31.7063 | 31.8592 | 32.0821 | 31.9384 | 32.0166 | Pyr_redox_2;Pyr_redox_dim | Dihydrolipoyl dehydrogenase {ECO:0000256\|RuleBase:RU003692} | | 1.760172539 | 0.5756791 | 0.248902003 | | 39 | 39 | 66.5 | 66.5 | 50.64 |
| A0A0H3NHR4 | 29.7073 | 29.493 | 29.622 | 30.085 | 29.878 | 29.6601 | ATP-synt | ATP synthase gamma chain {ECO:0000256\|HAMAP-Rule:MF_00815} | | 0.905261926 | 0.5759028 | 0.266909281 | | 22 | 22 | 51.9 | 51.9 | 31.56 |
| A0A0H3NMK2 | 25.1106 | 25.3407 | 25.0968 | 25.6204 | 25.9161 | 24.9288 | HSP33 | 33 kDa chaperonin {ECO:0000256\|HAMAP-Rule:MF_00117} | | 0.431665324 | 0.57698127 | 0.305733363 | | 2 | 2 | 7.8 | 7.8 | 32.64 |
| A0A0H3ND65 | 27.3952 | 27.6751 | 27.2579 | 27.5962 | 27.6572 | 27.8734 | ADH_zinc_N | | | 0.829263774 | 0.5787635 | 0.266199748 | | 7 | 7 | 27.2 | 27.2 | 37.76 |
| A0A0H3NGS7 | 32.6162 | 32.5331 | 32.4843 | 32.7171 | 32.7274 | 32.936 | Ribosomal_S7 | 30S ribosomal protein S7 {ECO:0000256\|HAMAP-Rule:MF_00480} | | 1.428995178 | 0.57951763 | 0.248958588 | | 22 | 22 | 84.6 | 84.6 | 17.59 |
| A0A0H3ND87 | 27.1123 | 27.3553 | 27.1265 | 27.4494 | 27.3487 | 27.5527 | DUF1338 | | | 1.205675675 | 0.58016296 | 0.252232869 | | 12 | 12 | 25.5 | 25.5 | 50.96 |
| A0A0H3NJA8 | 29.015 | 29.0712 | 28.9864 | 29.4008 | 29.273 | 29.1413 | Anticodon_1;tRNA-synt_1;Val_tRNA-synt_C | Valine--tRNA ligase {ECO:0000256\|HAMAP-Rule:MF_02004} | | 1.455261121 | 0.58034011 | 0.24745369 | | 23 | 23 | 23.6 | 23.6 | 108.2 |

### All proteins

| Uniprot accession number | LFQ intensity WT_1 | LFQ intensity WT_2 | LFQ intensity WT_3 | LFQ intensity OrfSwap_1 | LFQ intensity OrfSwap_2 | LFQ intensity OrfSwap_3 | Pfam name | Uniprot full protein name | Student's T-test Significant OrfSwap_WT | -Log Student's T-test p-value OrfSwap_WT | Student's T-test q-value OrfSwap_WT | Student's T-test Difference OrfSwap_WT | Cluster | Peptides | Unique peptides | Sequence coverage [%] | Unique sequence coverage [%] | Mol. weight [kDa] |
|---|---|---|---|---|---|---|---|---|---|---|---|---|---|---|---|---|---|---|
| A0A0H3NG01 | 29.0741 | 29.6171 | 29.3683 | 29.3991 | 29.5345 | 29.9807 | NDK | Nucleoside diphosphate kinase {ECO:0000256\|HAMAP-Rule:MF_00451, ECO:0000256\|RuleBase:RU004013} | | 0.533022163 | 0.5805948 | 0.284938176 | | 8 | 8 | 69.2 | 69.2 | 15.52 |
| A0A0H3NDT2 | 23.6174 | 27.2447 | 24.8378 | 26.1151 | 26.4519 | 24.6264 | SecIII_SopE_N;SopE_GEF | Guanine nucleotide exchange factor {ECO:0000256\|PIRNR:PIRNR034781} | | 0.154588908 | 0.58314128 | 0.497846603 | | 5 | 5 | 27.5 | 27.5 | 26.45 |
| A0A0H3NIW5 | 27.6574 | 27.7617 | 27.8132 | 28.061 | 27.91 | 27.983 | 5_nucleotid_C;Metallophos | | | 1.72012696 | 0.58316912 | 0.240551631 | | 11 | 11 | 23.8 | 23.8 | 70.52 |
| A0A0H3NIG9 | 31.3677 | 31.3122 | 31.4686 | 31.7435 | 31.6829 | 31.4663 | HSP90 | Chaperone protein HtpG {ECO:0000256\|HAMAP-Rule:MF_00505, ECO:0000256\|SAAS:SAAS00869959} | | 1.216742575 | 0.58407366 | 0.248072942 | | 45 | 45 | 60.7 | 60.7 | 71.49 |
| A0A0H3NRJ2 | 27.5266 | 27.7892 | 27.945 | 28.0615 | 27.9683 | 27.9954 | AMP_N;Peptidase_M24 | | | 0.952698732 | 0.58491513 | 0.254805883 | | 8 | 8 | 22.1 | 22.1 | 42.48 |
| A0A0H3NFR8 | 30.8858 | 31.0907 | 30.8772 | 31.3021 | 31.2489 | 31.0428 | tRNA_anti-codon;tRNA-synt_2 | Lysine--tRNA ligase {ECO:0000256\|HAMAP-Rule:MF_00252} | | 1.099939552 | 0.58625547 | 0.24672699 | | 28 | 28 | 48.1 | 48.1 | 57.58 |
| A0A0H3NGZ6 | 27.9901 | 28.38 | 27.9452 | 28.3078 | 27.7687 | 27.3636 | T3SS_TC | | | 0.403558158 | 0.5871298 | -0.291743596 | | 9 | 9 | 35.6 | 35.6 | 37.11 |
| A0A0H3NL96 | 29.0318 | 28.7623 | 29.1962 | 29.1741 | 29.0202 | 29.6124 | HAMP;MCPsignal;TarH | | | 0.553170932 | 0.58774359 | 0.272109985 | | 18 | 15 | 37.1 | 31.1 | 60.17 |

#### All proteins

| Uniprot accession number | LFQ intensity WT_1 | LFQ intensity WT_2 | LFQ intensity WT_3 | LFQ intensity OrfSwap_1 | LFQ intensity OrfSwap_2 | LFQ intensity OrfSwap_3 | Pfam name | Uniprot full protein name | Student's T-test Significant OrfSwap_WT | -Log Student's T-test p-value OrfSwap_WT | Student's T-test q-value OrfSwap_WT | Student's T-test Difference OrfSwap_WT | Cluster | Peptides | Unique peptides | Sequence coverage [%] | Unique sequence coverage [%] | Mol. weight [kDa] |
|---|---|---|---|---|---|---|---|---|---|---|---|---|---|---|---|---|---|---|
| A0A0H3NDE7 | 25.1327 | 24.8555 | 24.1497 | 25.591 | 25.2109 | 24.3189 | NA37 | Nucleoid-associated protein YejK {ECO:0000256\|HAMAP-Rule:MF_00730} | | 0.275656243 | 0.588051 | 0.327667236 | | 3 | 3 | 11.3 | 11.3 | 37.88 |
| A0A0H3NJK0 | 30.9697 | 31.0388 | 30.9866 | 30.8072 | 30.8622 | 30.6163 | ATP-synt_ab;Rho_N;Rho_RNA_bind | Transcription termination factor Rho {ECO:0000256\|HAMAP-Rule:MF_01884} | | 1.422340992 | 0.59157818 | -0.236412684 | | 33 | 33 | 56.6 | 56.6 | 46.99 |
| A0A0H3NIQ6 | 24.1953 | 26.3124 | 24.6534 | 25.5135 | 25.5384 | 25.1851 | PS_Dcarbxylase | Phosphatidylserine decarboxylase proenzyme {ECO:0000256\|HAMAP-Rule:MF_00662} | | 0.21314918 | 0.59629764 | 0.358632406 | | 4 | 4 | 17.4 | 17.4 | 35.89 |
| A0A0H3NBR4 | 30.7913 | 30.6707 | 30.851 | 30.8613 | 30.2572 | 30.4307 | Redoxin | Thiol peroxidase {ECO:0000256\|HAMAP-Rule:MF_00269} | | 0.609946416 | 0.60173913 | -0.254587809 | | 15 | 15 | 62.5 | 62.5 | 18.03 |
| A0A0H3NXR8 | 26.9344 | 27.0252 | 27.2318 | 26.8906 | 26.9523 | 26.6223 | | | | 0.836888317 | 0.60287884 | -0.242052714 | | 5 | 5 | 36 | 36 | 12.43 |
| A0A0H3NPH5 | 29.3795 | 29.7998 | 29.4952 | 29.2266 | 29.2519 | 29.4651 | EFP;EFP_N;Elong-fact-P_C | Elongation factor P {ECO:0000256\|HAMAP-Rule:MF_00141} | | 0.766026664 | 0.60407942 | -0.243666967 | | 7 | 7 | 31.9 | 31.9 | 20.62 |

### All proteins

| Uniprot accession number | LFQ intensity WT_1 | LFQ intensity WT_2 | LFQ intensity WT_3 | LFQ intensity OrfSwap_1 | LFQ intensity OrfSwap_2 | LFQ intensity OrfSwap_3 | Pfam name | Uniprot full protein name | Student's T-test Significant OrfSwap_WT | -Log Student's T-test p-value OrfSwap_WT | Student's T-test q-value OrfSwap_WT | Student's T-test Difference OrfSwap_WT | Cluster | Peptides | Unique peptides | Sequence coverage [%] | Unique sequence coverage [%] | Mol. weight [kDa] |
|---|---|---|---|---|---|---|---|---|---|---|---|---|---|---|---|---|---|---|
| A0A0H3NIZ3 | 34.663 | 34.5744 | 34.5816 | 34.8101 | 34.792 | 34.8586 | Cpn60_TCP1 | 60 kDa chaperonin {ECO:0000256\|HAMAP-Rule:MF_00600, ECO:0000256\|RuleBase:RU000419} | | 2.453731468 | 0.60450355 | 0.213873545 | | 58 | 58 | 75.7 | 75.7 | 57.29 |
| A0A0H3NMT2 | 23.8971 | 24.8103 | 25.1312 | 25.1082 | 24.9494 | 24.6484 | CBP_GIL | | | 0.29820662 | 0.60471048 | 0.289128621 | | 3 | 3 | 7.1 | 7.1 | 59.31 |
| A0A0H3NNY9 | 31.3262 | 31.418 | 31.3242 | 31.0835 | 31.2624 | 31.0404 | Ribosomal_L11;Ribosomal_L11_N | 50S ribosomal protein L11 {ECO:0000256\|HAMAP-Rule:MF_00736, ECO:0000256\|SAAS:SAAS00731162} | | 1.418063934 | 0.60486486 | -0.227379481 | | 10 | 10 | 45.1 | 45.1 | 14.88 |
| A0A0H3NVV1 | 29.5556 | 29.6818 | 29.4921 | 28.8869 | 29.557 | 29.5305 | Band_7 | Protein HflC {ECO:0000256\|PIRNR:PIRNR005651} | | 0.484072684 | 0.60490459 | -0.25164032 | | 19 | 19 | 57.5 | 57.5 | 37.54 |
| A0A0H3NJ70 | 31.4326 | 31.327 | 31.3125 | 31.1074 | 31.1468 | 31.1745 | His_Phos_1 | 2,3-bisphosphoglycerate-dependent phosphoglycerate mutase {ECO:0000256\|HAMAP-Rule:MF_01039, ECO:0000256\|RuleBase:RU004512} | | 2.137263471 | 0.60521062 | -0.214448293 | | 19 | 19 | 58.8 | 58.8 | 28.49 |
| A0A0H3NI93 | 29.0441 | 28.8768 | 29.032 | 28.797 | 28.5832 | 28.8849 | DJ-1_PfpI | Glyoxalase {ECO:0000256\|PIRNR:PIRNR006320} | | 1.030087029 | 0.60578648 | -0.229297002 | | 7 | 7 | 54.8 | 54.8 | 22.9 |

### All proteins

| Uniprot accession number | LFQ intensity WT_1 | LFQ intensity WT_2 | LFQ intensity WT_3 | LFQ intensity OrfSwap_1 | LFQ intensity OrfSwap_2 | LFQ intensity OrfSwap_3 | Pfam name | Uniprot full protein name | Student's T-test Significant OrfSwap_WT | -Log Student's T-test p-value OrfSwap_WT | Student's T-test q-value OrfSwap_WT | Student's T-test Difference OrfSwap_WT | Cluster | Peptides | Unique peptides | Sequence coverage [%] | Unique sequence coverage [%] | Mol. weight [kDa] |
|---|---|---|---|---|---|---|---|---|---|---|---|---|---|---|---|---|---|---|
| A0A0H3NPA2 | 28.425 | 28.1159 | 27.9749 | 27.6609 | 28.267 | 27.8121 | OEP | | | 0.50128766 | 0.60596403 | -0.258629481 | | 13 | 13 | 31.7 | 31.7 | 50.51 |
| A0A0H3NG02 | 31.3815 | 31.5676 | 31.3694 | 31.4765 | 31.9065 | 31.6476 | | | | 0.779050531 | 0.60625312 | 0.237372716 | | 20 | 20 | 70.4 | 70.4 | 31.44 |
| A0A0H3NFY2 | 29.4877 | 29.6819 | 29.2331 | 29.5327 | 29.2887 | 28.7943 | Polysacc_deac_1 | | | 0.445469469 | 0.60668336 | -0.262335459 | | 12 | 12 | 30.9 | 30.9 | 35.19 |
| A0A0H3NE38 | 25.7181 | 24.4853 | 26.4807 | 25.5499 | 25.0255 | 25.1072 | FliG_C;FliG_M;FliG_N | Flagellar motor switch protein FliG {ECO:0000256\|PIRNR:PIRNR003161} | | 0.214849283 | 0.60733571 | -0.333819071 | | 3 | 3 | 11.2 | 11.2 | 36.85 |
| A0A0H3NFZ7 | 26.7393 | 26.8081 | 26.582 | 26.5556 | 25.9766 | 26.827 | HAMP;MCPsignal | | | 0.421856468 | 0.60740035 | -0.256755193 | | 8 | 6 | 15.7 | 12.8 | 58.9 |
| A0A0H3NB27 | 27.6356 | 26.064 | 25.554 | 27.0981 | 28.2601 | 25.2074 | Sod_Cu | Superoxide dismutase [Cu-Zn] {ECO:0000256\|RuleBase:RU000393} | | 0.149806786 | 0.60777061 | 0.437334061 | | 3 | 3 | 9.8 | 9.8 | 17.74 |
| A0A0H3NM79 | 27.2074 | 27.4179 | 27.3312 | 27.4855 | 27.6324 | 27.5165 | PGM_PMM_I;PGM_PMM_II;PGM_PMM_III;PGM_PMM_IV | Phosphoglucosamine mutase {ECO:0000256\|HAMAP-Rule:MF_01554, ECO:0000256\|RuleBase:RU004327, ECO:0000256\|SAAS:SAAS00358419} | | 1.392200495 | 0.60782765 | 0.225961685 | | 10 | 10 | 24 | 24 | 47.44 |

### All proteins

| Uniprot accession number | LFQ intensity WT_1 | LFQ intensity WT_2 | LFQ intensity WT_3 | LFQ intensity OrfSwap_1 | LFQ intensity OrfSwap_2 | LFQ intensity OrfSwap_3 | Pfam name | Uniprot full protein name | Student's T-test Significant OrfSwap_WT | -Log Student's T-test p-value OrfSwap_WT | Student's T-test q-value OrfSwap_WT | Student's T-test Difference OrfSwap_WT | Cluster | Peptides | Unique peptides | Sequence coverage [%] | Unique sequence coverage [%] | Mol. weight [kDa] |
|---|---|---|---|---|---|---|---|---|---|---|---|---|---|---|---|---|---|---|
| A0A0H3NAQ6 | 24.955 | 24.151 | 25.4144 | 25.0691 | 25.1724 | 25.1117 | ROK | N-acetyl-D-glucosamine kinase {ECO:0000256\|HAMAP-Rule:MF_01271, ECO:0000256\|SAAS:SAAS00693724} | | 0.305182033 | 0.60806338 | 0.277616501 | | 3 | 3 | 8.6 | 8.6 | 33.06 |
| A0A0H3NH43 | 25.1597 | 23.6869 | 24.9329 | 25.3214 | 25.036 | 24.3518 | SRP54;SRP54_N | Signal recognition particle receptor FtsY {ECO:0000256\|HAMAP-Rule:MF_00920} | | 0.223834297 | 0.60810508 | 0.30988884 | | 4 | 4 | 10.8 | 10.8 | 53.73 |
| A0A0H3NBP3 | 30.2556 | 30.7005 | 30.4565 | 30.5781 | 30.733 | 30.7954 | ADH_N;ADH_zinc_N | | | 0.736767419 | 0.60876626 | 0.231296539 | | 12 | 12 | 46.9 | 46.9 | 32.55 |
| A0A0H3NIM4 | 27.9466 | 27.7004 | 27.857 | 28.1076 | 28.0106 | 28.0322 | Fe-S_biosyn;NifU | Fe/S biogenesis protein NfuA {ECO:0000256\|HAMAP-Rule:MF_01637, ECO:0000256\|SAAS:SAAS01079107} | | 1.299190936 | 0.60889354 | 0.21546936 | | 5 | 5 | 24.1 | 24.1 | 20.94 |
| A0A0H3NAG7 | 31.3214 | 31.4182 | 31.4122 | 31.2623 | 31.2311 | 31.0015 | Aldedh;Pro_dh;Pro_dh-DNA_bdg | Bifunctional protein PutA {ECO:0000256\|PIRNR:PIRNR000197} | | 1.1700411 | 0.60898246 | -0.218968074 | | 50 | 50 | 35.2 | 35.2 | 144.1 |

#### All proteins

| Uniprot accession number | LFQ intensity WT_1 | LFQ intensity WT_2 | LFQ intensity WT_3 | LFQ intensity OrfSwap_1 | LFQ intensity OrfSwap_2 | LFQ intensity OrfSwap_3 | Pfam name | Uniprot full protein name | Student's T-test Significant OrfSwap_WT | -Log Student's T-test p-value OrfSwap_WT | Student's T-test q-value OrfSwap_WT | Student's T-test Difference OrfSwap_WT | Cluster | Peptides | Unique peptides | Sequence coverage [%] | Unique sequence coverage [%] | Mol. weight [kDa] |
|---|---|---|---|---|---|---|---|---|---|---|---|---|---|---|---|---|---|---|
| A0A0H3NK26 | 26.8715 | 27.1166 | 26.9824 | 26.6775 | 26.8496 | 26.794 | PHP | Probable phosphatase YcdX {ECO:0000256\|HAMAP-Rule:MF_01561} | | 1.1680235 | 0.60976307 | -0.216481527 | | 5 | 5 | 21.6 | 21.6 | 26.91 |
| A0A0H3NH08 | 31.7541 | 31.7201 | 31.648 | 31.4871 | 31.5424 | 31.4699 | Ribosomal_S20p | 30S ribosomal protein S20 {ECO:0000256\|HAMAP-Rule:MF_00500} | | 2.256444283 | 0.60995804 | -0.207614899 | | 10 | 10 | 50.6 | 50.6 | 9.655 |
| A0A0H3NH64 | 28.3733 | 28.761 | 28.5483 | 28.4667 | 28.5268 | 27.9688 | SopD | | | 0.501192432 | 0.61050435 | -0.240128835 | | 11 | 11 | 34.7 | 34.7 | 36.14 |
| A0A0H3NMF0 | 32.1846 | 32.4434 | 32.3553 | 32.569 | 32.5512 | 32.4976 | Ribosomal_L6 | 50S ribosomal protein L6 {ECO:0000256\|HAMAP-Rule:MF_01365, ECO:0000256\|RuleBase:RU003870} | | 1.257500359 | 0.6128125 | 0.211518606 | | 22 | 22 | 71.8 | 71.8 | 18.86 |
| A0A0H3N8Z8 | 27.5933 | 27.3868 | 27.5497 | 27.3456 | 27.1202 | 27.4146 | PTS_EIIA_1;PTS_EIIB;PTS_EIIC | | | 0.929505312 | 0.61387175 | -0.216497421 | | 9 | 9 | 13.1 | 13.1 | 68.49 |
| A0A0H3NP98 | 27.1931 | 26.9754 | 26.9828 | 26.7744 | 26.8676 | 26.8881 | ABC_tran | UvrABC system protein A {ECO:0000256\|HAMAP-Rule:MF_00205, ECO:0000256\|SAAS:SAAS00385002} | | 1.224093393 | 0.62136332 | -0.207073212 | | 12 | 12 | 16.7 | 16.7 | 103.9 |
| A0A0H3NLC3 | 28.9052 | 29.247 | 29.3013 | 29.5524 | 29.2804 | 29.2781 | OsmC | | | 0.643270485 | 0.62535406 | 0.21916453 | | 9 | 9 | 39.9 | 39.9 | 15.1 |
| A0A0H3NH80 | 27.1851 | 26.8836 | 26.8656 | 27.0735 | 26.8463 | 26.3094 | LacI;Periplа_BP_1 | | | 0.399224768 | 0.6263821 | -0.235059102 | | 7 | 7 | 23.4 | 23.4 | 38.04 |

### All proteins

| Uniprot accession number | LFQ intensity WT_1 | LFQ intensity WT_2 | LFQ intensity WT_3 | LFQ intensity OrfSwap_1 | LFQ intensity OrfSwap_2 | LFQ intensity OrfSwap_3 | Pfam name | Uniprot full protein name | Student's T-test Significant OrfSwap_WT | -Log Student's T-test p-value OrfSwap_WT | Student's T-test q-value OrfSwap_WT | Student's T-test Difference OrfSwap_WT | Cluster | Peptides | Unique peptides | Sequence coverage [%] | Unique sequence coverage [%] | Mol. weight [kDa] |
|---|---|---|---|---|---|---|---|---|---|---|---|---|---|---|---|---|---|---|
| A0A0H3NAW3 | 24.9837 | 24.9306 | 24.6271 | 24.9142 | 24.4986 | 24.46 | DUF444 | UPF0229 protein YeaH {ECO:0000256 \|HAMAP-Rule:MF_01232} | | 0.537394813 | 0.62746207 | -0.222870509 | | 5 | 5 | 9.6 | 9.6 | 49.52 |
| A0A0H3NHY9 | 28.9714 | 29.2601 | 29.3024 | 29.0749 | 28.9642 | 28.8708 | Peptidase_M24 | Xaa-Pro dipeptidase {ECO:0000256 \|HAMAP-Rule:MF_01279} | | 0.804300909 | 0.63142955 | -0.207989375 | | 13 | 13 | 35.4 | 35.4 | 50.17 |
| A0A0H3NHR1 | 30.0083 | 29.5662 | 29.3757 | 29.5014 | 29.3167 | 29.4707 | OSCP | ATP synthase subunit delta {ECO:0000256 \|HAMAP-Rule:MF_01416} | | 0.490236033 | 0.63514237 | -0.220423381 | | 8 | 8 | 49.2 | 49.2 | 19.41 |
| A0A0H3NID6 | 31.8456 | 31.8299 | 31.7992 | 32.1328 | 31.9612 | 31.9565 | Ribosomal_S13 | 30S ribosomal protein S13 {ECO:0000256 \|HAMAP-Rule:MF_01315} | | 1.491823001 | 0.64347945 | 0.191942215 | | 17 | 17 | 80.5 | 80.5 | 13.16 |
| A0A0H3NMJ6 | 36.057 | 35.4642 | 36.0784 | 35.8975 | 35.444 | 35.5871 | Flagellin_C;Flagellin_D3;Flagellin_N | Flagellin {ECO:0000256 \|RuleBase:RU362073} | | 0.390217786 | 0.64523077 | -0.22366333 | | 36 | 14 | 65.5 | 34.7 | 51.61 |
| A0A0H3NAU2 | 31.7684 | 31.8009 | 31.8695 | 31.959 | 32.114 | 31.9359 | 2-oxoacid_dh;Biotin_lipoyl;E3_binding | Dihydrolipoyllysine-residue succinyltransferase component of 2-oxoglutarate dehydrogenase complex {ECO:0000256 \|RuleBase:RU361138} | | 1.398437155 | 0.64602385 | 0.190045039 | | 36 | 36 | 47.8 | 47.8 | 43.86 |

### All proteins

| Uniprot accession number | LFQ intensity WT_1 | LFQ intensity WT_2 | LFQ intensity WT_3 | LFQ intensity OrfSwap_1 | LFQ intensity OrfSwap_2 | LFQ intensity OrfSwap_3 | Pfam name | Uniprot full protein name | Student's T-test Significant OrfSwap_WT | -Log Student's T-test p-value OrfSwap_WT | Student's T-test q-value OrfSwap_WT | Student's T-test Difference OrfSwap_WT | Cluster | Peptides | Unique peptides | Sequence coverage [%] | Unique sequence coverage [%] | Mol. weight [kDa] |
|---|---|---|---|---|---|---|---|---|---|---|---|---|---|---|---|---|---|---|
| A0A0H3NXL4 | 25.9112 | 24.4299 | 25.8222 | 25.6723 | 25.643 | 25.6378 | CbiA | | | 0.213002151 | 0.64685034 | 0.263261795 | | 4 | 4 | 23.3 | 23.3 | 22.4 |
| A0A0H3NFT0 | 30.865 | 31.0839 | 31.069 | 30.8385 | 30.7671 | 30.8361 | GCV_T;GCV_T_C | Aminomethyltransferase {ECO:0000256\|HAMAP-Rule:MF_00259} | | 1.21350255 | 0.64691468 | -0.192082087 | | 14 | 14 | 49.5 | 49.5 | 40.22 |
| A0A0H3N850 | 26.5864 | 24.6526 | 25.3622 | 25.4652 | 24.8385 | 25.4571 | ACAS_N;AMP-binding;AMP-binding_C | | | 0.176698292 | 0.65339219 | -0.280147552 | | 5 | 5 | 7.5 | 7.5 | 69.33 |
| A0A0H3NQV4 | 29.6878 | 29.4187 | 29.4693 | 29.902 | 29.7273 | 29.5381 | DHHA1;tRNA_SAD;tRNA-synt_2c | Alanine--tRNA ligase {ECO:0000256\|HAMAP-Rule:MF_00036} | | 0.669214047 | 0.65409137 | 0.197139104 | | 28 | 28 | 33.4 | 33.4 | 95.9 |
| A0A0H3NKH8 | 31.3835 | 31.1895 | 31.302 | 31.0654 | 31.1015 | 31.1564 | AAA_PrkA;PrkA | | | 1.380698136 | 0.65478378 | -0.183882395 | | 42 | 42 | 59.9 | 59.9 | 68.51 |
| A0A0H3ND26 | 30.894 | 30.9402 | 30.7791 | 30.9656 | 30.8212 | 31.4513 | DegT_DnrJ_EryC1 | | | 0.457289714 | 0.65501695 | 0.208275477 | | 23 | 23 | 47.6 | 47.6 | 48.1 |
| A0A0H3NGW9 | 25.8847 | 26.1553 | 26.0948 | 26.4216 | 26.0261 | 24.9346 | DHQ_synthase | 3-dehydroquinate synthase {ECO:0000256\|HAMAP-Rule:MF_00110, ECO:0000256\|SAAS:SAAS00210916} | | 0.215644134 | 0.65651939 | -0.250836054 | | 4 | 4 | 11.9 | 11.9 | 38.71 |
| A0A0H3NUE5 | 27.4964 | 27.6295 | 27.5956 | 27.6368 | 27.7445 | 27.9008 | HemY_N;TPR_2 | | | 1.014651549 | 0.65775462 | 0.186817169 | | 12 | 12 | 35.4 | 35.4 | 43.26 |
| A0A0H3NKE9 | 29.7723 | 29.1953 | 29.7741 | 29.0322 | 29.7157 | 29.3346 | Rhodanese | Sulfurtransferase {ECO:0000256\|RuleBase:RU000507} | | 0.327302519 | 0.65779125 | -0.219727198 | | 11 | 11 | 36.8 | 36.8 | 30.83 |

### All proteins

| Uniprot accession number | LFQ intensity WT_1 | LFQ intensity WT_2 | LFQ intensity WT_3 | LFQ intensity OrfSwap_1 | LFQ intensity OrfSwap_2 | LFQ intensity OrfSwap_3 | Pfam name | Uniprot full protein name | Student's T-test Significant OrfSwap_WT | -Log Student's T-test p-value OrfSwap_WT | Student's T-test q-value OrfSwap_WT | Student's T-test Difference OrfSwap_WT | Cluster | Peptides | Unique peptides | Sequence coverage [%] | Unique sequence coverage [%] | Mol. weight [kDa] |
|---|---|---|---|---|---|---|---|---|---|---|---|---|---|---|---|---|---|---|
| A0A0H3NMA3 | 31.4215 | 32.0059 | 31.7707 | 30.9946 | 31.5329 | 31.9862 | Ribosomal_S9 | 30S ribosomal protein S9 {ECO:0000256\|HAMAP-Rule:MF_00532, ECO:0000256\|RuleBase:RU003816, ECO:0000256\|SAAS:SAAS00015375} | | 0.27488336 | 0.65830872 | -0.228157679 | | 12 | 12 | 67.7 | 67.7 | 14.83 |
| A0A0H3NCE6 | 27.8135 | 27.8221 | 27.9559 | 28.3247 | 27.4157 | 27.1641 | CheB_methylest;Response_reg | Protein-glutamate methylesterase/protein-glutamine glutaminase {ECO:0000256\|HAMAP-Rule:MF_00099} | | 0.256070595 | 0.66309548 | -0.229002635 | | 6 | 6 | 25.5 | 25.5 | 37.55 |
| A0A0H3NT12 | 33.1477 | 33.2741 | 33.2291 | 33.2873 | 32.8736 | 32.918 | PEPCK_ATP | Phosphoenolpyruvate carboxykinase (ATP) {ECO:0000256\|HAMAP-Rule:MF_00453, ECO:0000256\|SAAS:SAAS01086626} | | 0.630049874 | 0.66338 | -0.190594991 | | 49 | 49 | 70.1 | 70.1 | 59.65 |

### All proteins

| Uniprot accession number | LFQ intensity WT_1 | LFQ intensity WT_2 | LFQ intensity WT_3 | LFQ intensity OrfSwap_1 | LFQ intensity OrfSwap_2 | LFQ intensity OrfSwap_3 | Pfam name | Uniprot full protein name | Student's T-test Significant OrfSwap_WT | -Log Student's T-test p-value OrfSwap_WT | Student's T-test q-value OrfSwap_WT | Student's T-test Difference OrfSwap_WT | Cluster | Peptides | Unique peptides | Sequence coverage [%] | Unique sequence coverage [%] | Mol. weight [kDa] |
|---|---|---|---|---|---|---|---|---|---|---|---|---|---|---|---|---|---|---|
| A0A0H3NSB3 | 30.3342 | 30.2318 | 30.6851 | 30.2392 | 30.2882 | 30.1482 | AAA;FtsH_ext;Peptidase_M41 | ATP-dependent zinc metalloprotease FtsH {ECO:0000256\|HAMAP-Rule:MF_01458} | | 0.599201317 | 0.66416694 | -0.191806793 | | 30 | 30 | 45.3 | 45.3 | 70.83 |
| A0A0H3NE09 | 30.0616 | 30.0235 | 30.0716 | 30.5312 | 30.2258 | 29.9839 | PTS_EIIA_1 | | | 0.540605078 | 0.66444147 | 0.194708506 | | 9 | 9 | 50.3 | 50.3 | 18.25 |
| A0A0H3NCR0 | 25.812 | 24.7445 | 25.4424 | 24.493 | 26.4388 | 25.8941 | Mur_ligase_C;Mur_ligase_M | UDP-N-acetylmuramoylalanine--D-glutamate ligase {ECO:0000256\|HAMAP-Rule:MF_00639, ECO:0000256\|RuleBase:RU003664, ECO:0000256\|SAAS:SAAS00382177} | | 0.156747377 | 0.66575042 | 0.275692622 | | 4 | 4 | 17.8 | 17.8 | 47 |
| A0A0H3NIN1 | 33.2423 | 33.0483 | 32.9929 | 32.922 | 32.8553 | 32.9709 | Bac_DNA_binding | | | 1.012552447 | 0.66957475 | -0.178457896 | | 9 | 9 | 75.6 | 75.6 | 9.521 |
| A0A0G2PMS1 | 35.6836 | 35.6187 | 35.8652 | 35.5113 | 35.4796 | 35.6415 | GTP_EFTU;GTP_EFTU_D2;GTP_EFTU_D3 | Elongation factor Tu {ECO:0000256\|HAMAP-Rule:MF_00118, ECO:0000256\|RuleBase:RU004061} | | 0.938616189 | 0.67115755 | -0.178351084 | | 45 | 45 | 80.2 | 80.2 | 43.28 |
| A0A0H3NIW2 | 25.5001 | 25.7123 | 24.929 | 24.3925 | 25.0075 | 26.0107 | Osmo_CC;Sugar_tr | | | 0.175453048 | 0.67978182 | -0.243555705 | | 3 | 3 | 6.2 | 6.2 | 54.79 |
| A0A0H3NDY7 | 27.8312 | 27.9364 | 27.7336 | 27.7303 | 27.6951 | 27.5595 | Nitroreductase | | | 1.031765201 | 0.68075497 | -0.17209816 | | 10 | 10 | 48.4 | 48.4 | 23.96 |

#### All proteins

| Uniprot accession number | LFQ intensity WT_1 | LFQ intensity WT_2 | LFQ intensity WT_3 | LFQ intensity OrfSwap_1 | LFQ intensity OrfSwap_2 | LFQ intensity OrfSwap_3 | Pfam name | Uniprot full protein name | Student's T-test Significant OrfSwap_WT | -Log Student's T-test p-value OrfSwap_WT | Student's T-test q-value OrfSwap_WT | Student's T-test Difference OrfSwap_WT | Cluster | Peptides | Unique peptides | Sequence coverage [%] | Unique sequence coverage [%] | Mol. weight [kDa] |
|---|---|---|---|---|---|---|---|---|---|---|---|---|---|---|---|---|---|---|
| A0A0H3N8P5 | 29.6874 | 29.377 | 29.6026 | 29.4171 | 29.3119 | 29.4139 | Anti-adapt_IraP | Anti-adapter protein IraP {ECO:0000256\|HAMAP-Rule:MF_01198} | | 0.818425927 | 0.68143894 | -0.174692154 | | 7 | 7 | 64.8 | 64.8 | 10.11 |
| A0A0H3N957 | 32.248 | 31.8994 | 32.0812 | 32.1103 | 31.7632 | 31.8089 | Transketolase_N | Pyruvate dehydrogenase E1 component {ECO:0000256\|PIRNR:PIRNR000156} | | 0.542285232 | 0.68210873 | -0.182070414 | | 49 | 49 | 51.7 | 51.7 | 99.58 |
| A0A0H3NGY2 | 27.862 | 27.2515 | 27.2904 | 27.8006 | 27.4336 | 27.7528 | S1;Tex_N;Tex_YqgF | | | 0.35370157 | 0.68296053 | 0.1943868 | | 14 | 14 | 22.5 | 22.5 | 84.86 |
| A0A0H3N7Q0 | 24.9092 | 25.7289 | 25.6599 | 25.4915 | 25.6499 | 25.7567 | PolyA_pol;PolyA_pol_arg_C;PolyA_pol_RNAbd | Poly(A) polymerase I {ECO:0000256\|HAMAP-Rule:MF_00957} | | 0.296604237 | 0.68302951 | 0.200063705 | | 5 | 5 | 11.7 | 11.7 | 54.68 |
| A0A0H3NB52 | 27.5142 | 27.2197 | 27.3167 | 27.3972 | 27.4039 | 27.7926 | PMI_typeI | Mannose-6-phosphate isomerase {ECO:0000256\|RuleBase:RU000611} | | 0.505066336 | 0.68395417 | 0.181035995 | | 6 | 6 | 23.3 | 23.3 | 42.59 |
| A0A0H3NCR8 | 28.2348 | 28.1812 | 28.0884 | 27.9447 | 27.9716 | 28.0878 | SecA_DEAD;SecA_PP_bind;SecA_SW;SEC-C | Protein translocase subunit SecA {ECO:0000256\|HAMAP-Rule:MF_01382, ECO:0000256\|RuleBase:RU003874} | | 1.276574475 | 0.68415107 | -0.166775386 | | 12 | 12 | 14.2 | 14.2 | 101.8 |
| A0A0H3NB41 | 27.7927 | 27.8182 | 27.8837 | 27.5644 | 27.8351 | 27.5849 | LysM;YkuD | | | 0.868213359 | 0.68483007 | -0.17004776 | | 6 | 6 | 26.1 | 26.1 | 36.13 |

### All proteins

| Uniprot accession number | LFQ intensity WT_1 | LFQ intensity WT_2 | LFQ intensity WT_3 | LFQ intensity OrfSwap_1 | LFQ intensity OrfSwap_2 | LFQ intensity OrfSwap_3 | Pfam name | Uniprot full protein name | Student's T-test Significant OrfSwap_WT | -Log Student's T-test p-value OrfSwap_WT | Student's T-test q-value OrfSwap_WT | Student's T-test Difference OrfSwap_WT | Cluster | Peptides | Unique peptides | Sequence coverage [%] | Unique sequence coverage [%] | Mol. weight [kDa] |
|---|---|---|---|---|---|---|---|---|---|---|---|---|---|---|---|---|---|---|
| A0A0H3NDB8 | 25.427 | 24.9666 | 25.9314 | 26.2615 | 24.6453 | 26.155 | GDP_Man_Dehyd | dTDP-glucose 4,6-dehydratase {ECO:0000256\|RuleBase:RU004473} | | 0.155305165 | 0.68500813 | 0.245586395 | | 5 | 5 | 17.2 | 17.2 | 40.72 |
| A0A0H3NFL6 | 30.0605 | 30.0221 | 30.2136 | 30.4663 | 30.3311 | 30.0276 | PEP-utilisers_N;PEP-utilizers;PEP-utilizers_C | Phosphoenolpyruvate-protein phosphotransferase {ECO:0000256\|PIRNR:PIRNR000732} | | 0.548023341 | 0.68612378 | 0.176262538 | | 25 | 25 | 37.9 | 37.9 | 63.37 |
| A0A0H3NF91 | 29.6975 | 29.6275 | 29.2936 | 29.404 | 29.3088 | 29.3908 | Sod_Cu | Superoxide dismutase [Cu-Zn] {ECO:0000256\|RuleBase:RU000393} | | 0.599585123 | 0.6862589 | -0.171646118 | | 6 | 6 | 42.4 | 42.4 | 18.37 |
| A0A0H3NUS7 | 25.5428 | 25.4644 | 25.4676 | 24.5659 | 25.0132 | 26.2131 | FdhE | Protein FdhE {ECO:0000256\|HAMAP-Rule:MF_00611} | | 0.175184949 | 0.68666451 | -0.227526347 | | 5 | 5 | 20.4 | 20.4 | 34.71 |
| A0A0H3NMF3 | 31.2288 | 30.9813 | 31.5676 | 31.6021 | 31.455 | 31.274 | Ribosomal_L14 | 50S ribosomal protein L14 {ECO:0000256\|HAMAP-Rule:MF_01367, ECO:0000256\|RuleBase:RU003950} | | 0.401334513 | 0.68683197 | 0.184476852 | | 11 | 11 | 73.2 | 73.2 | 13.57 |
| A0A0H3NNA2 | 32.7971 | 32.6072 | 32.6347 | 32.9843 | 32.8402 | 32.7173 | ATP-synt_ab;ATP-synt_ab_C;ATP-synt_ab_N | ATP synthase subunit alpha {ECO:0000256\|HAMAP-Rule:MF_01346} | | 0.795820948 | 0.68757792 | 0.167588552 | | 42 | 42 | 62.8 | 62.8 | 55.11 |

### All proteins

| Uniprot accession number | LFQ intensity WT_1 | LFQ intensity WT_2 | LFQ intensity WT_3 | LFQ intensity OrfSwap_1 | LFQ intensity OrfSwap_2 | LFQ intensity OrfSwap_3 | Pfam name | Uniprot full protein name | Student's T-test Significant OrfSwap_WT | -Log Student's T-test p-value OrfSwap_WT | Student's T-test q-value OrfSwap_WT | Student's T-test Difference OrfSwap_WT | Cluster | Peptides | Unique peptides | Sequence coverage [%] | Unique sequence coverage [%] | Mol. weight [kDa] |
|---|---|---|---|---|---|---|---|---|---|---|---|---|---|---|---|---|---|---|
| A0A0H3NWH8 | 30.1809 | 30.15 | 29.7652 | 29.7805 | 29.8105 | 29.9905 | ABC_tran;ABC_tran_Xtn | | | 0.503723601 | 0.69296618 | -0.171532313 | | 23 | 23 | 36.4 | 36.4 | 62.31 |
| A0A0H3NIQ0 | 30.8963 | 30.7656 | 30.6958 | 31.0815 | 31.0059 | 30.7693 | FAD_binding_2;Succ_DH_flav_C | Fumarate reductase flavoprotein subunit {ECO:0000256\|RuleBase:RU362050} | | 0.682030638 | 0.69300645 | 0.166360219 | | 32 | 32 | 51 | 51 | 65.49 |
| A0A0H3NAM4 | 26.18 | 25.9064 | 25.7908 | 25.4069 | 25.9307 | 25.9919 | AstB | N-succinylarginine dihydrolase {ECO:0000256\|HAMAP-Rule:MF_01172, ECO:0000256\|SAAS:SAAS00376480} | | 0.346243877 | 0.6937706 | -0.182580312 | | 4 | 4 | 11.4 | 11.4 | 49.17 |
| A0A0H3NRH2 | 28.2558 | 28.5306 | 28.1311 | 28.7197 | 28.3494 | 28.3665 | GCV_T | tRNA-modifying protein YgfZ {ECO:0000256\|HAMAP-Rule:MF_01175, ECO:0000256\|SAAS:SAAS00035597} | | 0.438825619 | 0.6961672 | 0.172705332 | | 11 | 11 | 43.3 | 43.3 | 35.97 |
| E1WFA1 | 31.7553 | 31.472 | 31.7973 | 31.7751 | 31.3681 | 31.3686 | Response_reg;Trans_reg_C | Virulence transcriptional regulatory protein PhoP | | 0.430615038 | 0.69937721 | -0.170899709 | | 20 | 20 | 69.6 | 69.6 | 25.63 |

### All proteins

| Uniprot accession number | LFQ intensity WT_1 | LFQ intensity WT_2 | LFQ intensity WT_3 | LFQ intensity OrfSwap_1 | LFQ intensity OrfSwap_2 | LFQ intensity OrfSwap_3 | Pfam name | Uniprot full protein name | Student's T-test Significant OrfSwap_WT | -Log Student's T-test p-value OrfSwap_WT | Student's T-test q-value OrfSwap_WT | Student's T-test Difference OrfSwap_WT | Cluster | Peptides | Unique peptides | Sequence coverage [%] | Unique sequence coverage [%] | Mol. weight [kDa] |
|---|---|---|---|---|---|---|---|---|---|---|---|---|---|---|---|---|---|---|
| A0A0H3N8B3 | 28.4956 | 28.6516 | 28.6056 | 28.5006 | 28.4149 | 28.3771 | HGTP_anticodon;tRNA_edit;tRNA-synt_2b | Proline--tRNA ligase {ECO:0000256\|HAMAP-Rule:MF_01569} | | 1.222382801 | 0.70105128 | -0.153394699 | | 13 | 13 | 22.9 | 22.9 | 63.54 |
| A0A0H3NKR2 | 27.4334 | 27.6848 | 27.5769 | 27.4536 | 27.4295 | 27.3465 | Fer2_3;Fer4_8 | Succinate dehydrogenase iron-sulfur subunit {ECO:0000256\|RuleBase:RU361237} | | 0.908732597 | 0.7035584 | -0.155172348 | | 5 | 5 | 27.5 | 27.5 | 27.18 |
| A0A0H3N8I2 | 27.4644 | 27.5261 | 27.7075 | 27.2261 | 27.5788 | 27.4225 | Crl | Sigma factor-binding protein Crl {ECO:0000256\|HAMAP-Rule:MF_01178, ECO:0000256\|SAAS:SAAS00371097} | | 0.554124208 | 0.71254226 | -0.156885783 | | 3 | 3 | 12.8 | 12.8 | 15.8 |
| A0A0H3NPJ4 | 24.7256 | 26.3759 | 26.0988 | 26.1305 | 24.985 | 25.4115 | TsaE | | | 0.135768459 | 0.71368051 | -0.224454244 | | 1 | 1 | 9.2 | 9.2 | 16.9 |
| A0A0H3NUR8 | 25.3015 | 25.322 | 25.1028 | 25.2719 | 25.005 | 24.9853 | DeoRC;HTH_DeoR | | | 0.597209175 | 0.7137707 | -0.154690425 | | 2 | 2 | 8.2 | 8.2 | 29.05 |
| A0A0H3NM65 | 28.2654 | 27.907 | 27.9416 | 28.1116 | 28.1019 | 28.3706 | BON | | | 0.471168475 | 0.7172337 | 0.156735738 | | 8 | 8 | 41.4 | 41.4 | 20.1 |
| A0A0H3NDV8 | 30.1074 | 29.4718 | 30.1325 | 30.0997 | 30.0205 | 30.089 | Pro_isomerase | Peptidyl-prolyl cis-trans isomerase {ECO:0000256\|RuleBase:RU363019} | | 0.311297138 | 0.7175873 | 0.165873845 | | 13 | 13 | 64.6 | 64.6 | 18.14 |

### All proteins

| Uniprot accession number | LFQ intensity WT_1 | LFQ intensity WT_2 | LFQ intensity WT_3 | LFQ intensity OrfSwap_1 | LFQ intensity OrfSwap_2 | LFQ intensity OrfSwap_3 | Pfam name | Uniprot full protein name | Student's T-test Significant OrfSwap_WT | -Log Student's T-test p-value OrfSwap_WT | Student's T-test q-value OrfSwap_WT | Student's T-test Difference OrfSwap_WT | Cluster | Peptides | Unique peptides | Sequence coverage [%] | Unique sequence coverage [%] | Mol. weight [kDa] |
|---|---|---|---|---|---|---|---|---|---|---|---|---|---|---|---|---|---|---|
| A0A0H3NIL3 | 31.1277 | 31.1657 | 31.208 | 31.2283 | 31.3529 | 31.3442 | 6PGD;NAD_binding_2 | 6-phosphogluconate dehydrogenase, decarboxylating {ECO:0000256\|PIRNR:PIRNR000109, ECO:0000256\|RuleBase:RU000485} | | 1.419081102 | 0.7210523 | 0.14139239 | | 25 | 25 | 49.1 | 49.1 | 51.4 |
| A0A0H3NEG7 | 29.1816 | 28.9334 | 28.924 | 28.987 | 28.7405 | 28.8633 | Lactonase | 6-phosphogluconolactonase {ECO:0000256\|HAMAP-Rule:MF_01605, ECO:0000256\|SAAS:SAAS00634850} | | 0.607054614 | 0.72117722 | -0.149438222 | | 13 | 13 | 44.1 | 44.1 | 36.37 |
| A0A0H3NFW3 | 30.3049 | 29.6174 | 29.7639 | 30.3407 | 30.1374 | 29.7203 | Asparaginase | | | 0.242796561 | 0.72210726 | 0.17074585 | | 14 | 14 | 46 | 46 | 36.93 |
| A0A0H3NH96 | 28.6807 | 28.2548 | 28.3644 | 28.6496 | 28.5317 | 28.5737 | DUF3412;DUF4478;Lysine_decarbox | | | 0.501115774 | 0.72269826 | 0.151693344 | | 16 | 16 | 34.4 | 34.4 | 50.75 |
| A0A0H3NIH8 | 24.9376 | 29.302 | 28.8674 | 28.5185 | 28.6753 | 24.7775 | ZapB | Cell division protein ZapB {ECO:0000256\|HAMAP-Rule:MF_01196, ECO:0000256\|SAAS:SAAS00371340} | | 0.070293144 | 0.72627673 | -0.378538767 | | 5 | 5 | 53.2 | 53.2 | 9.312 |

### All proteins

| Uniprot accession number | LFQ intensity WT_1 | LFQ intensity WT_2 | LFQ intensity WT_3 | LFQ intensity OrfSwap_1 | LFQ intensity OrfSwap_2 | LFQ intensity OrfSwap_3 | Pfam name | Uniprot full protein name | Student's T-test Significant OrfSwap_WT | -Log Student's T-test p-value OrfSwap_WT | Student's T-test q-value OrfSwap_WT | Student's T-test Difference OrfSwap_WT | Cluster | Peptides | Unique peptides | Sequence coverage [%] | Unique sequence coverage [%] | Mol. weight [kDa] |
|---|---|---|---|---|---|---|---|---|---|---|---|---|---|---|---|---|---|---|
| A0A0H3NJF2 | 32.6891 | 32.7789 | 32.7474 | 32.7789 | 32.6449 | 32.3455 | ATP-synt_ab;ATP-synt_ab_N | ATP synthase subunit beta {ECO:0000256\|HAMAP-Rule:MF_01347} | | 0.496159185 | 0.72702992 | -0.148696899 | | 29 | 29 | 64.8 | 64.8 | 50.28 |
| A0A0H3NG11 | 27.4829 | 27.5878 | 27.4594 | 27.5976 | 27.1803 | 27.3119 | DUF3663;Peptidase_M17 | Peptidase B {ECO:0000256\|HAMAP-Rule:MF_00504} | | 0.495074958 | 0.72897959 | -0.146792094 | | 16 | 16 | 39.1 | 39.1 | 46.36 |
| A0A0H3N950 | 32.8587 | 32.7333 | 32.7567 | 32.7537 | 32.6209 | 32.5584 | ATP-grasp_2;Ligase_CoA | Succinate--CoA ligase [ADP-forming] subunit beta {ECO:0000256\|HAMAP-Rule:MF_00558} | | 0.935297938 | 0.72943574 | -0.138577779 | | 30 | 30 | 58.5 | 58.5 | 41.5 |
| A0A0H3NGT9 | 25.8862 | 25.1505 | 23.8099 | 24.9301 | 25.2006 | 25.3344 | MarR | | | 0.121496711 | 0.73622535 | 0.206135432 | | 5 | 5 | 36.4 | 36.4 | 20.54 |
| A0A0H3NSF7 | 25.2855 | 24.8395 | 25.0804 | 24.9332 | 26.9825 | 24.0085 | ROK | N-acetylmannosamine kinase {ECO:0000256\|HAMAP-Rule:MF_01234, ECO:0000256\|SAAS:SAAS00693515} | | 0.096539069 | 0.7367375 | 0.239597956 | | 3 | 3 | 14.4 | 14.4 | 30.08 |
| A0A0H3N9Z8 | 35.0094 | 34.5099 | 34.8783 | 34.1396 | 34.9475 | 34.8223 | OmpA;OmpA_membrane | | | 0.216633105 | 0.73785959 | -0.162743886 | | 32 | 32 | 70.6 | 70.6 | 37.52 |

### All proteins

| Uniprot accession number | LFQ intensity WT_1 | LFQ intensity WT_2 | LFQ intensity WT_3 | LFQ intensity OrfSwap_1 | LFQ intensity OrfSwap_2 | LFQ intensity OrfSwap_3 | Pfam name | Uniprot full protein name | Student's T-test Significant OrfSwap_WT | -Log Student's T-test p-value OrfSwap_WT | Student's T-test q-value OrfSwap_WT | Student's T-test Difference OrfSwap_WT | Cluster | Peptides | Unique peptides | Sequence coverage [%] | Unique sequence coverage [%] | Mol. weight [kDa] |
|---|---|---|---|---|---|---|---|---|---|---|---|---|---|---|---|---|---|---|
| A0A0H3NI90 | 30.9131 | 31.2561 | 31.1022 | 31.0392 | 30.6337 | 31.1531 | Ribosomal_L31 | 50S ribosomal protein L31 {ECO:0000256\|HAMAP-Rule:MF_00501, ECO:0000256\|SAAS:SAAS00804274} | | 0.327967517 | 0.73820218 | -0.148462931 | | 12 | 12 | 97.1 | 97.1 | 7.719 |
| A0A0H3NJJ2 | 24.6282 | 25.3363 | 25.9028 | 25.5026 | 25.401 | 24.3957 | DUF2813 | | | 0.1368971 | 0.73880997 | -0.189306259 | | 4 | 4 | 10 | 10 | 63.55 |
| E1WAB5 | 25.7931 | 25.9226 | 24.3596 | 24.8154 | 25.0902 | 25.6147 | OrgA_MxiK | Oxygen-regulated invasion protein OrgA | | 0.122162689 | 0.75232298 | -0.184972763 | | 2 | 2 | 13.6 | 13.6 | 22.83 |
| A0A0H3NRK4 | 31.7116 | 30.5951 | 31 | 31.2965 | 31.28 | 31.1892 | F_bP_aldolase | | | 0.177050961 | 0.76315659 | 0.15298907 | | 23 | 23 | 63.8 | 63.8 | 39.16 |
| A0A0H3NVE4 | 27.6475 | 27.3926 | 27.9993 | 27.736 | 27.5138 | 27.3772 | Acyltransferase | Glycerol-3-phosphate acyltransferase {ECO:0000256\|HAMAP-Rule:MF_00393} | | 0.268942093 | 0.76583901 | -0.137517293 | | 16 | 16 | 18.1 | 18.1 | 91.13 |
| A0A0H3NA74 | 27.6686 | 28.0115 | 28.1157 | 28.0441 | 27.9735 | 28.1673 | DUF480 | UPF0502 protein YceH {ECO:0000256\|HAMAP-Rule:MF_01584} | | 0.370689021 | 0.7678949 | 0.129697164 | | 11 | 11 | 52.6 | 52.6 | 24.12 |
| A0A0H3NGC5 | 30.4171 | 30.6236 | 30.7155 | 30.3865 | 30.2469 | 30.7354 | Ribosomal_S15 | 30S ribosomal protein S15 {ECO:0000256\|HAMAP-Rule:MF_01343, ECO:0000256\|RuleBase:RU003920} | | 0.31003796 | 0.77398148 | -0.129110336 | | 7 | 7 | 39.3 | 39.3 | 10.2 |

### All proteins

| Uniprot accession number | LFQ intensity WT_1 | LFQ intensity WT_2 | LFQ intensity WT_3 | LFQ intensity OrfSwap_1 | LFQ intensity OrfSwap_2 | LFQ intensity OrfSwap_3 | Pfam name | Uniprot full protein name | Student's T-test Significant OrfSwap_WT | -Log Student's T-test p-value OrfSwap_WT | Student's T-test q-value OrfSwap_WT | Student's T-test Difference OrfSwap_WT | Cluster | Peptides | Unique peptides | Sequence coverage [%] | Unique sequence coverage [%] | Mol. weight [kDa] |
|---|---|---|---|---|---|---|---|---|---|---|---|---|---|---|---|---|---|---|
| A0A0H3NJ13 | 29.5046 | 29.0538 | 29.2659 | 29.5182 | 29.4651 | 29.221 | FUR | Ferric uptake regulation protein {ECO:0000256\|RuleBase:RU364037} | | 0.327285549 | 0.77635077 | 0.126676559 | | 8 | 8 | 70.7 | 70.7 | 17.01 |
| A0A0H3NJI1 | 25.3056 | 26.2536 | 26.3227 | 24.7332 | 26.2273 | 26.3945 | His_Phos_1 | Probable phosphoglycerate mutase GpmB {ECO:0000256\|HAMAP-Rule:MF_01040, ECO:0000256\|SAAS:SAAS00727155} | | 0.101505191 | 0.77664417 | -0.17563947 | | 6 | 6 | 35.3 | 35.3 | 23.87 |
| A0A0H3NC78 | 26.2427 | 26.2841 | 26.3444 | 26.4331 | 26.2671 | 25.7799 | ABC_tran;oligo_HPY | | | 0.262278875 | 0.77693395 | -0.130371094 | | 7 | 7 | 30.1 | 30.1 | 36.86 |
| E1W874 | 27.6217 | 27.745 | 27.2549 | 27.4092 | 27.4181 | 27.4178 | Aminotran_3 | Glutamate-1-semialdehyde 2,1-aminomutase | | 0.354623941 | 0.77715871 | -0.125501633 | | 9 | 9 | 29.3 | 29.3 | 45.33 |
| A0A0H3NF87 | 30.7587 | 31.1697 | 30.9017 | 31.4275 | 31.1556 | 30.6525 | SseC;T3SSipB | | | 0.20289883 | 0.77780704 | 0.135164897 | | 33 | 33 | 51.8 | 51.8 | 62.45 |
| A0A0H3NHF0 | 31.2882 | 31.2935 | 31.0688 | 31.3536 | 31.4017 | 31.2393 | Ribosomal_L28 | 50S ribosomal protein L28 {ECO:0000256\|HAMAP-Rule:MF_00373} | | 0.578688075 | 0.78200612 | 0.114691416 | | 9 | 9 | 66.7 | 66.7 | 9.05 |

### All proteins

| Uniprot accession number | LFQ intensity WT_1 | LFQ intensity WT_2 | LFQ intensity WT_3 | LFQ intensity OrfSwap_1 | LFQ intensity OrfSwap_2 | LFQ intensity OrfSwap_3 | Pfam name | Uniprot full protein name | Student's T-test Significant OrfSwap_WT | -Log Student's T-test p-value OrfSwap_WT | Student's T-test q-value OrfSwap_WT | Student's T-test Difference OrfSwap_WT | Cluster | Peptides | Unique peptides | Sequence coverage [%] | Unique sequence coverage [%] | Mol. weight [kDa] |
|---|---|---|---|---|---|---|---|---|---|---|---|---|---|---|---|---|---|---|
| A0A0H3NHJ8 | 28.0159 | 28.1238 | 28.2178 | 27.8722 | 28.4179 | 27.6824 | MinE | Cell division topological specificity factor {ECO:0000256\|HAMAP-Rule:MF_00262, ECO:0000256\|SAAS:SAAS01078064} | | 0.21918627 | 0.78461679 | -0.128313065 | | 8 | 8 | 46.6 | 46.6 | 10.18 |
| A0A0H3NIM7 | 27.9155 | 27.7853 | 27.8403 | 27.7718 | 27.6247 | 27.8268 | GDP_Man_Dehyd | | | 0.675908775 | 0.79256193 | -0.105896632 | | 14 | 14 | 36.5 | 36.5 | 41.02 |
| A0A0H3NKE0 | 31.2103 | 30.4085 | 30.6182 | 30.4585 | 31.2148 | 30.9555 | | | | 0.148783303 | 0.79286275 | 0.130602519 | | 9 | 9 | 57.1 | 57.1 | 17.17 |
| A0A0H3NCB9 | 24.4846 | 25.371 | 25.4874 | 25.6452 | 24.6426 | 24.6201 | Lip_A_acyltrans | Lipid A biosynthesis myristoyltransferase {ECO:0000256\|HAMAP-Rule:MF_01944} | | 0.113570684 | 0.79288956 | -0.144990921 | | 2 | 2 | 6.8 | 6.8 | 37.29 |
| A0A0H3NI43 | 24.8434 | 26.0174 | 25.0533 | 25.2427 | 24.6917 | 25.5505 | DUF1090 | | | 0.118378458 | 0.79358788 | -0.143090566 | | 3 | 3 | 11.5 | 11.5 | 14.07 |
| A0A0H3NKZ4 | 28.2968 | 28.7859 | 28.6823 | 28.4955 | 28.8632 | 28.7634 | Glutaredoxin | Glutaredoxin {ECO:0000256\|PIRNR:PIRNR005894} | | 0.255794524 | 0.79477626 | 0.119003932 | | 10 | 10 | 73.9 | 73.9 | 12.91 |
| A0A0H3N9B4 | 29.1002 | 29.0981 | 28.8932 | 29.097 | 29.1023 | 29.2138 | PDZ | Periplasmic serine endoprotease DegP-like {ECO:0000256\|RuleBase:RU364067} | | 0.612765135 | 0.79479211 | 0.107159297 | | 12 | 12 | 31.4 | 31.4 | 49.32 |

### All proteins

| Uniprot accession number | LFQ intensity WT_1 | LFQ intensity WT_2 | LFQ intensity WT_3 | LFQ intensity OrfSwap_1 | LFQ intensity OrfSwap_2 | LFQ intensity OrfSwap_3 | Pfam name | Uniprot full protein name | Student's T-test Significant OrfSwap_WT | -Log Student's T-test p-value OrfSwap_WT | Student's T-test q-value OrfSwap_WT | Student's T-test Difference OrfSwap_WT | Cluster | Peptides | Unique peptides | Sequence coverage [%] | Unique sequence coverage [%] | Mol. weight [kDa] |
|---|---|---|---|---|---|---|---|---|---|---|---|---|---|---|---|---|---|---|
| A0A0H3NKX3 | 29.0261 | 29.681 | 29.6052 | 29.3263 | 29.6919 | 29.6565 | Pyrophosphatase | Inorganic pyrophosphatase {ECO:0000256\|HAMAP-Rule:MF_00209} | | 0.195465312 | 0.79490964 | 0.120784124 | | 10 | 10 | 53.4 | 53.4 | 19.68 |
| A0A0H3NC72 | 27.9959 | 23.6475 | 28.1385 | 27.9395 | 24.8787 | 27.8041 | YCII | | | 0.054247315 | 0.79518293 | 0.280146917 | | 3 | 3 | 20.4 | 20.4 | 10.59 |
| A0A0H3NK66 | 26.3105 | 26.0941 | 26.2316 | 26.5853 | 26.4823 | 25.9307 | M20_dimer;Peptidase_M20 | Acetylornithine deacetylase {ECO:0000256\|HAMAP-Rule:MF_01108} | | 0.22123186 | 0.79564134 | 0.120737076 | | 7 | 7 | 20.1 | 20.1 | 42.2 |
| A0A0H3NFF1 | 28.6373 | 28.7959 | 28.6128 | 28.8046 | 28.5228 | 28.3955 | His_Phos_2 | | | 0.33179739 | 0.80280301 | -0.107749939 | | 10 | 10 | 25.9 | 25.9 | 45.56 |
| A0A0H3N9J8 | 27.6118 | 27.8787 | 27.6537 | 27.538 | 27.6619 | 27.6385 | SeqA;SeqA_N | Negative modulator of initiation of replication {ECO:0000256\|HAMAP-Rule:MF_00908, ECO:0000256\|PIRNR:PIRNR019401} | | 0.486657561 | 0.80695495 | -0.101923625 | | 5 | 5 | 37.8 | 37.8 | 20.15 |
| A0A0H3NPH9 | 25.4904 | 26.4306 | 25.4927 | 25.8945 | 25.3495 | 25.804 | Lipocalin_2 | Outer membrane lipoprotein Blc {ECO:0000256\|PIRNR:PIRNR036893} | | 0.125514685 | 0.81467066 | -0.121884028 | | 5 | 5 | 32.2 | 32.2 | 19.96 |
| A0A0H3NGR7 | 26.8848 | 27.1741 | 27.1643 | 27.4745 | 27.1531 | 26.9167 | malic;Malic_M | NAD-dependent malic enzyme {ECO:0000256\|HAMAP-Rule:MF_01619} | | 0.222932967 | 0.81541829 | 0.107024511 | | 8 | 8 | 17.7 | 17.7 | 62.9 |

### All proteins

| Uniprot accession number | LFQ intensity WT_1 | LFQ intensity WT_2 | LFQ intensity WT_3 | LFQ intensity OrfSwap_1 | LFQ intensity OrfSwap_2 | LFQ intensity OrfSwap_3 | Pfam name | Uniprot full protein name | Student's T-test Significant OrfSwap_WT | -Log Student's T-test p-value OrfSwap_WT | Student's T-test q-value OrfSwap_WT | Student's T-test Difference OrfSwap_WT | Cluster | Peptides | Unique peptides | Sequence coverage [%] | Unique sequence coverage [%] | Mol. weight [kDa] |
|---|---|---|---|---|---|---|---|---|---|---|---|---|---|---|---|---|---|---|
| A0A0H3NI02 | 24.7628 | 25.6682 | 25.1949 | 26.0384 | 24.9461 | 25.0191 | TatC | Sec-independent protein translocase protein TatC {ECO:0000256\|HAMAP-Rule:MF_00902} | | 0.103213069 | 0.8200597 | 0.12590917 | | 1 | 1 | 6.9 | 6.9 | 29.07 |
| A0A0H3NWG0 | 25.6536 | 25.5301 | 25.0936 | 25.6902 | 25.6733 | 25.2367 | GTP_EFTU;GTP_EFTU_D2;RF3_C | Peptide chain release factor 3 {ECO:0000256\|HAMAP-Rule:MF_00072, ECO:0000256\|SAAS:SAAS00669814} | | 0.181726302 | 0.82014948 | 0.107659022 | | 6 | 6 | 12.1 | 12.1 | 59.56 |
| A0A0H3NCT4 | 25.3182 | 24.2247 | 25.4631 | 24.7172 | 24.96 | 25.7178 | Aldo_ket_red | | | 0.093805846 | 0.82069151 | 0.129671733 | | 4 | 4 | 15.8 | 15.8 | 33.68 |
| A0A0H3NE04 | 29.4943 | 28.9639 | 29.4387 | 29.6657 | 28.6472 | 29.2378 | Ferritin | Ferritin {ECO:0000256\|RuleBase:RU361145} | | 0.124229602 | 0.82250595 | -0.115422567 | | 7 | 7 | 52.1 | 52.1 | 19.28 |
| A0A0H3NDZ5 | 29.2937 | 30.0952 | 29.8941 | 30.1096 | 29.7105 | 29.7899 | CheW;CheY-binding;HATPase_c;H-kinase_dim;Hpt | | | 0.150649776 | 0.82252006 | 0.109036128 | | 20 | 20 | 32.3 | 32.3 | 73.01 |
| A0A0H3NC48 | 31.1389 | 31.2929 | 31.3953 | 31.0987 | 31.0648 | 31.3869 | Aconitase;Aconitase_C | Aconitate hydratase {ECO:0000256\|RuleBase:RU361275} | | 0.295562149 | 0.82904579 | -0.092209498 | | 45 | 45 | 52 | 52 | 97.5 |

### All proteins

| Uniprot accession number | LFQ intensity WT_1 | LFQ intensity WT_2 | LFQ intensity WT_3 | LFQ intensity OrfSwap _1 | LFQ intensity OrfSwap _2 | LFQ intensity OrfSwap _3 | Pfam name | Uniprot full protein name | Student's T-test Significant OrfSwap_WT | -Log Student's T-test p-value OrfSwap_WT | Student's T-test q-value OrfSwap_WT | Student's T-test Difference OrfSwap_WT | Cluster | Peptides | Unique peptides | Sequence coverage [%] | Unique sequence coverage [%] | Mol. weight [kDa] |
|---|---|---|---|---|---|---|---|---|---|---|---|---|---|---|---|---|---|---|
| A0A0H3NMI8 | 25.231 | 25.1363 | 24.1994 | 25.5213 | 23.614 | 25.0233 | PALP | D-cysteine desulfhydrase {ECO:0000256\|HAMAP-Rule:MF_01045} | | 0.072318752 | 0.83013609 | -0.136027018 | | 2 | 2 | 10.1 | 10.1 | 34.91 |
| A0A0H3NFY6 | 28.2672 | 28.157 | 28.3607 | 28.4837 | 28.4142 | 28.1577 | GATase;GMP_synt_C;NAD_synthase | GMP synthase [glutamine-hydrolyzing] {ECO:0000256\|HAMAP-Rule:MF_00344, ECO:0000256\|SAAS:SAAS00723629} | | 0.321112113 | 0.83097935 | 0.090261459 | | 17 | 17 | 39.8 | 39.8 | 58.7 |
| A0A0H3NR00 | 27.2873 | 27.3061 | 27.1302 | 27.3014 | 27.1089 | 27.0428 | SicP-binding;Y_phosphatase;YopE | | | 0.399258642 | 0.8310637 | -0.090145747 | | 11 | 11 | 25.2 | 25.2 | 60.05 |
| A0A0H3NGP6 | 32.885 | 33.1556 | 32.9369 | 32.7522 | 32.9393 | 33.0103 | Ribosomal_S4;S4 | 30S ribosomal protein S4 {ECO:0000256\|HAMAP-Rule:MF_01306} | | 0.33526983 | 0.83152522 | -0.091917674 | | 26 | 26 | 75.2 | 75.2 | 23.49 |
| A0A0H3NXK9 | 28.8979 | 28.7937 | 28.9837 | 28.6501 | 28.7369 | 29.035 | Colicin;Colicin_Ia | | | 0.261056419 | 0.84904118 | -0.084404627 | | 21 | 21 | 39.1 | 39.1 | 70 |
| A0A0H3N925 | 27.9281 | 28.0177 | 27.838 | 28.1132 | 28.1132 | 27.8091 | NIF3 | | | 0.299162709 | 0.84977909 | 0.083895365 | | 5 | 5 | 27.5 | 27.5 | 26.95 |
| A0A0H3NAI9 | 27.7879 | 27.6471 | 27.9586 | 27.6249 | 27.7785 | 27.745 | MdoG | Glucans biosynthesis protein G {ECO:0000256\|HAMAP-Rule:MF_01069} | | 0.332144742 | 0.850279 | -0.081732432 | | 9 | 9 | 23.1 | 23.1 | 57.85 |

### All proteins

| Uniprot accession number | LFQ intensity WT_1 | LFQ intensity WT_2 | LFQ intensity WT_3 | LFQ intensity OrfSwap_1 | LFQ intensity OrfSwap_2 | LFQ intensity OrfSwap_3 | Pfam name | Uniprot full protein name | Student's T-test Significant OrfSwap_WT | -Log Student's T-test p-value OrfSwap_WT | Student's T-test q-value OrfSwap_WT | Student's T-test Difference OrfSwap_WT | Cluster | Peptides | Unique peptides | Sequence coverage [%] | Unique sequence coverage [%] | Mol. weight [kDa] |
|---|---|---|---|---|---|---|---|---|---|---|---|---|---|---|---|---|---|---|
| A0A0H3NNZ4 | 32.5274 | 32.5683 | 32.6166 | 32.5623 | 32.5126 | 32.4083 | RNA_pol_Rpb1_1;RNA_pol_Rpb1_2;RNA_pol_Rpb1_3;RNA_pol_Rpb1_4;RNA_pol_Rpb1_5 | DNA-directed RNA polymerase subunit beta' {ECO:0000256\|HAMAP-Rule:MF_01322} | | 0.662792516 | 0.85195906 | -0.076356252 | | 87 | 87 | 58.2 | 58.2 | 155.2 |
| A0A0H3NBD6 | 28.6175 | 27.8323 | 28.0412 | 28.1904 | 28.0645 | 27.9627 | FumaraseC_C;Lyase_1 | Fumarate hydratase class II {ECO:0000256\|HAMAP-Rule:MF_00743} | | 0.138230731 | 0.85197654 | -0.091200511 | | 11 | 11 | 33.4 | 33.4 | 50.3 |
| A0A0H3NG44 | 27.6513 | 27.6017 | 25.8338 | 27.3244 | 27.0887 | 27.0245 | CTP_transf_like;PfkB | Bifunctional protein HldE {ECO:0000256\|HAMAP-Rule:MF_01603} | | 0.067511675 | 0.85226354 | 0.116973241 | | 12 | 12 | 32.5 | 32.5 | 51.12 |
| A0A0H3NQ15 | 26.7613 | 26.9604 | 26.5038 | 27.0055 | 26.9099 | 26.5666 | SIS | | | 0.172471718 | 0.85254307 | 0.085518519 | | 8 | 8 | 29.5 | 29.5 | 38.39 |
| A0A0H3NGD9 | 30.6328 | 30.3312 | 30.537 | 30.5358 | 30.694 | 30.507 | GST_C;GST_N_2 | | | 0.300739964 | 0.85465306 | 0.078660329 | | 13 | 13 | 71.6 | 71.6 | 22.44 |
| A0A0H3NJ64 | 31.4018 | 31.592 | 31.3452 | 31.3698 | 31.6839 | 31.5196 | Aldedh | | | 0.265627558 | 0.85645415 | 0.078111013 | | 27 | 27 | 58.1 | 58.1 | 50.04 |
| A0A0H3N9D1 | 28.9847 | 28.8986 | 28.893 | 28.5966 | 28.89 | 29.054 | Bac_surface_Ag;POTRA | Outer membrane protein assembly factor BamA {ECO:0000256\|HAMAP-Rule:MF_01430} | | 0.223822382 | 0.85708721 | -0.078536352 | | 18 | 18 | 23.4 | 23.4 | 89.53 |

### All proteins

| Uniprot accession number | LFQ intensity WT_1 | LFQ intensity WT_2 | LFQ intensity WT_3 | LFQ intensity OrfSwap_1 | LFQ intensity OrfSwap_2 | LFQ intensity OrfSwap_3 | Pfam name | Uniprot full protein name | Student's T-test Significant OrfSwap_WT | -Log Student's T-test p-value OrfSwap_WT | Student's T-test q-value OrfSwap_WT | Student's T-test Difference OrfSwap_WT | Cluster | Peptides | Unique peptides | Sequence coverage [%] | Unique sequence coverage [%] | Mol. weight [kDa] |
|---|---|---|---|---|---|---|---|---|---|---|---|---|---|---|---|---|---|---|
| A0A0H3NVT1 | 26.1092 | 27.4496 | 26.9672 | 26.769 | 27.057 | 26.9887 | Fumarate_red_C | Fumarate reductase subunit C {ECO:0000256\|HAMAP-Rule:MF_00708, ECO:0000256\|SAAS:SAAS00819766} | | 0.08493812 | 0.85764877 | 0.096227646 | | 3 | 3 | 13.7 | 13.7 | 15 |
| A0A0H3NNN9 | 28.3017 | 28.1649 | 28.253 | 28.2896 | 28.0924 | 28.1182 | Molybdop_Fe4S4;Molybdopterin;Molydop_binding | | | 0.423175477 | 0.85798261 | -0.073103587 | | 16 | 14 | 24.7 | 21.7 | 112.4 |
| A0A0H3NKD1 | 30.0437 | 30.1677 | 29.8045 | 29.7866 | 29.9858 | 30.0149 | PGI | Glucose-6-phosphate isomerase {ECO:0000256\|HAMAP-Rule:MF_00473} | | 0.232887006 | 0.85895224 | -0.076215744 | | 29 | 29 | 45.5 | 45.5 | 61.43 |
| A0A0H3NBJ3 | 27.9658 | 27.9339 | 27.8897 | 27.6249 | 28.0259 | 28.3823 | adh_short | | | 0.136424994 | 0.86028324 | 0.081265767 | | 7 | 7 | 36.7 | 36.7 | 27.04 |
| A0A0H3NHR0 | 26.0826 | 25.5369 | 24.482 | 25.6584 | 25.4681 | 25.2498 | GSH-S_ATP;GSH-S_N | Glutathione synthetase {ECO:0000256\|HAMAP-Rule:MF_00162} | | 0.065890896 | 0.87419913 | 0.091590881 | | 3 | 3 | 18.1 | 18.1 | 35.43 |
| A0A0H3NLY2 | 25.085 | 24.2289 | 25.4975 | 24.3364 | 26.2286 | 24.5572 | DUF2884 | | | 0.050503737 | 0.875683 | 0.103595734 | | 3 | 3 | 10.9 | 10.9 | 26.27 |
| A0A0H3NJR8 | 29.0665 | 29.3923 | 29.102 | 28.8419 | 28.9345 | 29.5661 | 3HCDH;3HCDH_N;ECH_1 | Fatty acid oxidation complex subunit alpha {ECO:0000256\|HAMAP-Rule:MF_01621} | | 0.104919112 | 0.87918508 | -0.072758993 | | 25 | 25 | 31.8 | 31.8 | 79.59 |

### All proteins

| Uniprot accession number | LFQ intensity WT_1 | LFQ intensity WT_2 | LFQ intensity WT_3 | LFQ intensity OrfSwap_1 | LFQ intensity OrfSwap_2 | LFQ intensity OrfSwap_3 | Pfam name | Uniprot full protein name | Student's T-test Significant OrfSwap_WT | -Log Student's T-test p-value OrfSwap_WT | Student's T-test q-value OrfSwap_WT | Student's T-test Difference OrfSwap_WT | Cluster | Peptides | Unique peptides | Sequence coverage [%] | Unique sequence coverage [%] | Mol. weight [kDa] |
|---|---|---|---|---|---|---|---|---|---|---|---|---|---|---|---|---|---|---|
| A0A0H3NLC1 | 27.5923 | 27.7822 | 27.3795 | 27.3655 | 27.7771 | 27.4058 | Alpha-amylase;CBM_48 | | | 0.14527862 | 0.87974713 | -0.068511963 | | 9 | 9 | 17.2 | 17.2 | 78.54 |
| A0A0H3NBX8 | 27.1427 | 26.9271 | 27.3643 | 27.1187 | 27.3561 | 27.1607 | 2-Hacid_dh;2-Hacid_dh_C | | | 0.174457243 | 0.88054101 | 0.067153931 | | 10 | 10 | 26.7 | 26.7 | 36.42 |
| A0A0H3NDR9 | 26.441 | 25.5529 | 25.8128 | 26.235 | 25.1041 | 26.2292 | DUF3340;PDZ;Peptidase_S41 | | | 0.059982081 | 0.892149 | -0.079474767 | | 7 | 7 | 10.1 | 10.1 | 76.78 |
| A0A0H3NCA7 | 25.7666 | 25.4199 | 25.2405 | 25.6048 | 24.8501 | 25.7619 | PCRF;RF-1 | Peptide chain release factor 1 {ECO:0000256\|HAMAP-Rule:MF_00093} | | 0.076761558 | 0.89386857 | -0.070027669 | | 4 | 4 | 12.8 | 12.8 | 40.46 |
| A0A0H3NJN6 | 30.0116 | 30.0994 | 30.0947 | 30.1215 | 30.2155 | 30.0388 | Molybdop_Fe4S4;Molybdopterin;Molydop_binding;NADH-G_4Fe-4S_3 | NADH-quinone oxidoreductase {ECO:0000256\|RuleBase:RU003525} | | 0.412248768 | 0.89389986 | 0.05670166 | | 33 | 33 | 35.5 | 35.5 | 100 |
| A0A0H3NG03 | 27.5379 | 27.5237 | 25.1284 | 27.451 | 27.4554 | 25.5985 | | | | 0.035105118 | 0.8939886 | 0.104980469 | | 4 | 4 | 23 | 23 | 24.29 |
| A0A0H3NK42 | 25.1984 | 25.6309 | 25.3778 | 25.8243 | 25.174 | 25.0075 | Glycos_transf_2 | Glucans biosynthesis glucosyltransferase H {ECO:0000256\|HAMAP-Rule:MF_01072, ECO:0000256\|SAAS:SAAS00342634} | | 0.085237448 | 0.89445364 | -0.067089081 | | 5 | 5 | 7.9 | 7.9 | 97.1 |

#### All proteins

| Uniprot accession number | LFQ intensity WT_1 | LFQ intensity WT_2 | LFQ intensity WT_3 | LFQ intensity OrfSwap_1 | LFQ intensity OrfSwap_2 | LFQ intensity OrfSwap_3 | Pfam name | Uniprot full protein name | Student's T-test Significant OrfSwap_WT | -Log Student's T-test p-value OrfSwap_WT | Student's T-test q-value OrfSwap_WT | Student's T-test Difference OrfSwap_WT | Cluster | Peptides | Unique peptides | Sequence coverage [%] | Unique sequence coverage [%] | Mol. weight [kDa] |
|---|---|---|---|---|---|---|---|---|---|---|---|---|---|---|---|---|---|---|
| A0A0H3NKT7 | 29.3446 | 29.4069 | 29.3024 | 29.3966 | 29.3489 | 29.4683 | Adenylsucc_synt | Adenylosuccinate synthetase {ECO:0000256\|HAMAP-Rule:MF_00011, ECO:0000256\|RuleBase:RU000520} | | 0.50581225 | 0.8963357 | 0.053297679 | | 21 | 21 | 41.7 | 41.7 | 47.38 |
| A0A0H3NGE9 | 26.8392 | 25.1662 | 26.5241 | 26.6922 | 25.0183 | 27.0893 | MlaC | | | 0.037441282 | 0.89670822 | 0.090117137 | | 7 | 7 | 25.6 | 25.6 | 24 |
| A0A0H3NRQ8 | 25.1015 | 24.5978 | 25.2667 | 24.6516 | 25.4046 | 25.1043 | Methyltrans_RNA | Ribosomal RNA small subunit methyltransferase E {ECO:0000256\|PIRNR:PIRNR015601} | | 0.076765476 | 0.89712908 | 0.064848582 | | 3 | 3 | 13.9 | 13.9 | 28.9 |
| A0A0H3NGR5 | 31.8371 | 31.8153 | 31.7048 | 31.9002 | 31.8543 | 31.7547 | KOW;ribosomal_L24 | 50S ribosomal protein L24 {ECO:0000256\|HAMAP-Rule:MF_01326} | | 0.355113279 | 0.90143706 | 0.050661723 | | 15 | 15 | 89.4 | 89.4 | 11.32 |
| A0A0H3NC49 | 30.9394 | 30.937 | 30.9185 | 31.1227 | 30.9401 | 30.873 | CbiA | Site-determining protein {ECO:0000256\|PIRNR:PIRNR003092} | | 0.248277897 | 0.9115662 | 0.046971003 | | 21 | 21 | 65.9 | 65.9 | 29.5 |
| A0A0H3NV41 | 28.0315 | 27.9024 | 28.0008 | 28.021 | 27.9074 | 27.8664 | HTH_1;LysR_substrate | | | 0.315432109 | 0.91177433 | -0.046614965 | | 11 | 11 | 39 | 39 | 34.18 |
| A0A0H3NIS1 | 28.4696 | 28.2862 | 28.2596 | 27.9548 | 28.5005 | 28.4044 | SBP_bac_8 | Maltodextrin-binding protein {ECO:0000256\|RuleBase:RU365005} | | 0.103384552 | 0.91306215 | -0.051902771 | | 8 | 8 | 24.6 | 24.6 | 43.48 |

### All proteins

| Uniprot accession number | LFQ intensity WT_1 | LFQ intensity WT_2 | LFQ intensity WT_3 | LFQ intensity OrfSwap_1 | LFQ intensity OrfSwap_2 | LFQ intensity OrfSwap_3 | Pfam name | Uniprot full protein name | Student's T-test Significant OrfSwap_WT | -Log Student's T-test p-value OrfSwap_WT | Student's T-test q-value OrfSwap_WT | Student's T-test Difference OrfSwap_WT | Cluster | Peptides | Unique peptides | Sequence coverage [%] | Unique sequence coverage [%] | Mol. weight [kDa] |
|---|---|---|---|---|---|---|---|---|---|---|---|---|---|---|---|---|---|---|
| A0A0H3N830 | 27.8879 | 27.8953 | 28.1037 | 27.8983 | 27.9139 | 28.216 | Pro_CA | Carbonic anhydrase {ECO:0000256\|RuleBase:RU003956} | | 0.138941334 | 0.9151236 | 0.047068914 | | 6 | 6 | 32.7 | 32.7 | 24.82 |
| A0A0H3N9N1 | 30.7581 | 30.8535 | 30.8015 | 31.0001 | 30.9769 | 30.5794 | Fer2_3 | Succinate dehydrogenase iron-sulfur subunit {ECO:0000256\|RuleBase:RU361237} | | 0.125632674 | 0.91610127 | 0.04779307 | | 18 | 18 | 58.2 | 58.2 | 26.86 |
| A0A0H3NEA9 | 29.8783 | 29.7625 | 29.7351 | 30.0292 | 29.7822 | 29.4227 | AhpC-TSA | | | 0.092979908 | 0.91786555 | -0.047293981 | | 6 | 6 | 51.3 | 51.3 | 17.61 |
| A0A0H3NIH0 | 30.0808 | 30.3977 | 30.4504 | 30.2103 | 30.3032 | 30.5544 | DeoC | 3-hydroxy-5-phosphonooxypentane-2,4-dione thiolase {ECO:0000256\|HAMAP-Rule:MF_02052} | | 0.108199036 | 0.91897896 | 0.046288808 | | 13 | 13 | 49.5 | 49.5 | 31.74 |
| A0A0H3NKB2 | 28.0416 | 28.2948 | 28.0722 | 28.2157 | 28.0592 | 28.2636 | ArsC | Arsenate reductase {ECO:0000256\|RuleBase:RU362029} | | 0.161057218 | 0.91946294 | 0.043252945 | | 6 | 6 | 46.2 | 46.2 | 13.38 |
| A0A0H3NH20 | 27.9009 | 27.7403 | 28.1742 | 27.766 | 27.8999 | 28.0279 | Semialdhyde_dh;Semialdhyde_dhC | Aspartate-semialdehyde dehydrogenase {ECO:0000256\|HAMAP-Rule:MF_02121} | | 0.098485909 | 0.92875104 | -0.040529887 | | 9 | 9 | 24.7 | 24.7 | 40.14 |
| E1WGG9 | 24.6036 | 26.5196 | 26.0535 | 26.0078 | 26.312 | 24.6647 | Trehalose_PPase | Trehalose-phosphate phosphatase | | 0.028046112 | 0.92893855 | -0.064082464 | | 3 | 3 | 12 | 12 | 29.26 |
| A0A0H3NS87 | 28.8824 | 28.9119 | 28.7824 | 28.7343 | 29.0023 | 28.7233 | DJ-1_PfpI | | | 0.145721281 | 0.92955989 | -0.038924535 | | 9 | 9 | 65.7 | 65.7 | 18.87 |

### All proteins

| Uniprot accession number | LFQ intensity WT_1 | LFQ intensity WT_2 | LFQ intensity WT_3 | LFQ intensity OrfSwap_1 | LFQ intensity OrfSwap_2 | LFQ intensity OrfSwap_3 | Pfam name | Uniprot full protein name | Student's T-test Significant OrfSwap_WT | -Log Student's T-test p-value OrfSwap_WT | Student's T-test q-value OrfSwap_WT | Student's T-test Difference OrfSwap_WT | Cluster | Peptides | Unique peptides | Sequence coverage [%] | Unique sequence coverage [%] | Mol. weight [kDa] |
|---|---|---|---|---|---|---|---|---|---|---|---|---|---|---|---|---|---|---|
| A0A0H3NBF5 | 28.1649 | 28.2406 | 27.8088 | 27.9381 | 28.3813 | 28.0222 | Alpha-amylase;DUF3459 | Malto-oligosyltrehalose trehalohydrolase {ECO:0000256\|PIRNR:PIRNR006337} | | 0.078570049 | 0.92980195 | 0.042411168 | | 11 | 11 | 28.8 | 28.8 | 65.77 |
| A0A0H3NIG0 | 24.7694 | 25.0644 | 25.0578 | 25.2876 | 25.2856 | 24.4497 | Aminotran_1_2 | Aminotransferase {ECO:0000256\|RuleBase:RU000481} | | 0.050880454 | 0.93010526 | 0.04374822 | | 2 | 2 | 8.3 | 8.3 | 43.53 |
| A0A0H3NDU3 | 28.5976 | 29.0039 | 28.8376 | 28.7793 | 28.8491 | 28.9259 | CheW;Response_reg | | | 0.110880409 | 0.93023024 | 0.038371404 | | 15 | 15 | 47.4 | 47.4 | 37.01 |
| A0A0H3NFG7 | 29.8748 | 29.7291 | 29.6573 | 29.8473 | 29.7129 | 29.8127 | CTP_synth_N;GATase | CTP synthase {ECO:0000256\|HAMAP-Rule:MF_01227} | | 0.188171207 | 0.93044444 | 0.037218094 | | 19 | 19 | 34.5 | 34.5 | 60.12 |
| A0A0H3NLW5 | 31.7516 | 31.4594 | 31.5361 | 31.7572 | 31.5418 | 31.5594 | Transket_pyr;Transketolase_C;Transketolase_N | | | 0.121556543 | 0.9304675 | 0.037112554 | | 27 | 27 | 44.1 | 44.1 | 70.5 |
| A0A0H3NH09 | 34.3384 | 34.4589 | 34.3704 | 34.4355 | 34.5339 | 34.3033 | DAO;DAO_C | Glycerol-3-phosphate dehydrogenase {ECO:0000256\|RuleBase:RU361217} | | 0.174648368 | 0.93076414 | 0.034979502 | | 62 | 62 | 80.5 | 80.5 | 56.92 |
| A0A0H3NA75 | 32.8563 | 32.7429 | 32.8796 | 32.8902 | 32.8778 | 32.6012 | S1 | 30S ribosomal protein S1 {ECO:0000256\|PIRNR:PIRNR002111} | | 0.129777257 | 0.93090055 | -0.036514282 | | 49 | 49 | 55.1 | 55.1 | 61.17 |

### All proteins

| Uniprot accession number | LFQ intensity WT_1 | LFQ intensity WT_2 | LFQ intensity WT_3 | LFQ intensity OrfSwap_1 | LFQ intensity OrfSwap_2 | LFQ intensity OrfSwap_3 | Pfam name | Uniprot full protein name | Student's T-test Significant OrfSwap_WT | -Log Student's T-test p-value OrfSwap_WT | Student's T-test q-value OrfSwap_WT | Student's T-test Difference OrfSwap_WT | Cluster | Peptides | Unique peptides | Sequence coverage [%] | Unique sequence coverage [%] | Mol. weight [kDa] |
|---|---|---|---|---|---|---|---|---|---|---|---|---|---|---|---|---|---|---|
| A0A0H3NI80 | 31.207 | 31.1933 | 30.9261 | 31.0016 | 30.8566 | 31.3602 | TIM | Triosephosphate isomerase {ECO:0000256\|HAMAP-Rule:MF_00147, ECO:0000256\|RuleBase:RU363013, ECO:0000256\|SAAS:SAAS00728730} | | 0.071839007 | 0.9349146 | -0.03596433 | | 16 | 16 | 48.6 | 48.6 | 26.92 |
| A0A0H3NV38 | 27.57 | 27.4016 | 27.4316 | 27.6815 | 27.4669 | 27.1606 | PEPcase | Phosphoenolpyruvate carboxylase {ECO:0000256\|HAMAP-Rule:MF_00595, ECO:0000256\|SAAS:SAAS01082397} | | 0.068633911 | 0.94301099 | -0.031396866 | | 15 | 15 | 19.1 | 19.1 | 99 |
| A0A0H3NPF9 | 30.1282 | 30.1027 | 30.189 | 30.1294 | 30.0864 | 30.2912 | malic;Malic_M;PTA_PTB | | | 0.161941077 | 0.94353783 | 0.029040655 | | 26 | 26 | 42.4 | 42.4 | 82.32 |
| A0A0H3NTT7 | 28.6936 | 28.6422 | 28.6313 | 28.8908 | 28.6226 | 28.3622 | RNA_pol_Rpb6 | DNA-directed RNA polymerase subunit omega {ECO:0000256\|HAMAP-Rule:MF_00366, ECO:0000256\|SAAS:SAAS00387783} | | 0.069297921 | 0.94420302 | -0.030489604 | | 9 | 9 | 96.7 | 96.7 | 10.24 |

### All proteins

| Uniprot accession number | LFQ intensity WT_1 | LFQ intensity WT_2 | LFQ intensity WT_3 | LFQ intensity OrfSwap_1 | LFQ intensity OrfSwap_2 | LFQ intensity OrfSwap_3 | Pfam name | Uniprot full protein name | Student's T-test Significant OrfSwap_WT | -Log Student's T-test p-value OrfSwap_WT | Student's T-test q-value OrfSwap_WT | Student's T-test Difference OrfSwap_WT | Cluster | Peptides | Unique peptides | Sequence coverage [%] | Unique sequence coverage [%] | Mol. weight [kDa] |
|---|---|---|---|---|---|---|---|---|---|---|---|---|---|---|---|---|---|---|
| A0A0H3NGI3 | 31.9199 | 31.9568 | 31.8859 | 31.959 | 32.0144 | 31.8657 | Ribosomal_L13 | 50S ribosomal protein L13 {ECO:0000256\|HAMAP-Rule:MF_01366, ECO:0000256\|RuleBase:RU003878, ECO:0000256\|SAAS:SAAS00725370} | | 0.205395644 | 0.94864022 | 0.025496165 | | 20 | 20 | 94.4 | 94.4 | 16.02 |
| A0A0H3NLX1 | 23.3897 | 26.8697 | 27.1245 | 25.7958 | 24.3591 | 27.4118 | Orn_Arg_deC_N | Biosynthetic arginine decarboxylase {ECO:0000256\|HAMAP-Rule:MF_01417} | | 0.01349516 | 0.94979178 | 0.06092453 | | 9 | 9 | 23.4 | 23.4 | 71.75 |
| A0A0H3NIT6 | 26.9691 | 27.8297 | 27.6972 | 27.7164 | 27.5795 | 27.2904 | DcrB | | | 0.034414479 | 0.9506776 | 0.030085882 | | 5 | 5 | 38.4 | 38.4 | 19.69 |
| A0A0H3NAN4 | 29.9406 | 30.0976 | 30.2774 | 30.0794 | 29.9917 | 30.1688 | | | | 0.081045335 | 0.95096862 | -0.025223414 | | 8 | 8 | 41.8 | 41.8 | 25.55 |
| A0A0H3NB31 | 30.5324 | 30.4358 | 30.2229 | 30.0719 | 30.6111 | 30.4388 | Rick_17kDa_Anti | | | 0.042916221 | 0.9583619 | -0.023087819 | | 8 | 8 | 54.8 | 54.8 | 15.55 |
| A0A0H3NLZ8 | 30.623 | 30.7836 | 30.8916 | 30.7895 | 30.7271 | 30.7167 | CZB;MCPsignal | | | 0.095063578 | 0.95864305 | -0.021624247 | | 21 | 21 | 64.8 | 64.8 | 39.38 |
| A0A0H3ND48 | 27.5304 | 27.7333 | 27.3659 | 27.4576 | 28.0096 | 27.2334 | Alpha-amylase | | | 0.03140738 | 0.95869565 | 0.023662567 | | 19 | 19 | 28 | 28 | 95.03 |
| A0A0H3NK81 | 30.4741 | 31.3061 | 31.2217 | 30.851 | 30.8443 | 31.2377 | Ribosomal_L12;Ribosomal_L12_N | 50S ribosomal protein L7/L12 {ECO:0000256\|HAMAP-Rule:MF_00368} | | 0.02616262 | 0.96001628 | -0.022996267 | | 5 | 5 | 45.5 | 45.5 | 12.3 |

### All proteins

| Uniprot accession number | LFQ intensity WT_1 | LFQ intensity WT_2 | LFQ intensity WT_3 | LFQ intensity OrfSwap_1 | LFQ intensity OrfSwap_2 | LFQ intensity OrfSwap_3 | Pfam name | Uniprot full protein name | Student's T-test Significant OrfSwap_WT | -Log Student's T-test p-value OrfSwap_WT | Student's T-test q-value OrfSwap_WT | Student's T-test Difference OrfSwap_WT | Cluster | Peptides | Unique peptides | Sequence coverage [%] | Unique sequence coverage [%] | Mol. weight [kDa] |
|---|---|---|---|---|---|---|---|---|---|---|---|---|---|---|---|---|---|---|
| A0A0H3NEH2 | 28.3551 | 28.4703 | 28.3774 | 28.4243 | 28.3747 | 28.3503 | Urocanase;Urocanase_C;Urocanase_N | Urocanate hydratase {ECO:0000256\|HAMAP-Rule:MF_00577, ECO:0000256\|SAAS:SAAS00693702} | | 0.161889337 | 0.96086721 | -0.017843246 | | 16 | 16 | 32.4 | 32.4 | 61.47 |
| A0A0H3NEZ1 | 30.3937 | 30.5558 | 30.1628 | 30.4128 | 30.3823 | 30.2623 | Peripla_BP_4 | | | 0.051239251 | 0.96181867 | -0.018326441 | | 18 | 18 | 50.3 | 50.3 | 35.81 |
| A0A0H3NLV7 | 31.6748 | 31.8085 | 31.5429 | 31.739 | 31.7062 | 31.62 | Aldedh;Fe-ADH | Aldehyde-alcohol dehydrogenase {ECO:0000256\|PIRNR:PIRNR000111} | | 0.052987237 | 0.9749027 | 0.01300176 | | 39 | 39 | 50.4 | 50.4 | 96.21 |
| A0A0H3NC79 | 26.727 | 26.8634 | 26.3854 | 26.2517 | 27.9943 | 25.6725 | ProQ;ProQ_C | RNA chaperone ProQ {ECO:0000256\|HAMAP-Rule:MF_00749, ECO:0000256\|SAAS:SAAS00038850} | | 0.008807848 | 0.97640486 | -0.01906395 | | 7 | 7 | 33.3 | 33.3 | 25.44 |
| A0A0H3NAL3 | 27.8227 | 27.6938 | 27.7912 | 27.7743 | 27.7806 | 27.7808 | Nitroreductase | | | 0.08529314 | 0.98078167 | 0.009348551 | | 10 | 10 | 60.7 | 60.7 | 20.13 |
| A0A0H3N8W6 | 27.1476 | 27.8716 | 27.8089 | 27.2387 | 27.4312 | 28.1517 | CPSase_sm_chain;GATase | Carbamoyl-phosphate synthase small chain {ECO:0000256\|HAMAP-Rule:MF_01209} | | 0.001998113 | 0.99495187 | -0.002213796 | | 8 | 8 | 26.4 | 26.4 | 41.65 |

### All proteins

| Uniprot accession number | LFQ intensity WT_1 | LFQ intensity WT_2 | LFQ intensity WT_3 | LFQ intensity OrfSwap_1 | LFQ intensity OrfSwap_2 | LFQ intensity OrfSwap_3 | Pfam name | Uniprot full protein name | Student's T-test Significant OrfSwap_WT | -Log Student's T-test p-value OrfSwap_WT | Student's T-test q-value OrfSwap_WT | Student's T-test Difference OrfSwap_WT | Cluster | Peptides | Unique peptides | Sequence coverage [%] | Unique sequence coverage [%] | Mol. weight [kDa] |
|---|---|---|---|---|---|---|---|---|---|---|---|---|---|---|---|---|---|---|
| A0A0H3NK19 | 29.2068 | 29.5378 | 29.4342 | 29.3601 | 29.4605 | 29.37 | CDH | CDP-diacylglycerol pyrophosphatase {ECO:0000256\|HAMAP-Rule:MF_00319} | | 0.012617325 | 0.99554839 | 0.003928502 | | 14 | 14 | 61.8 | 61.8 | 28.37 |
| A0A0H3NAD2 | 28.9885 | 28.8328 | 29.1335 | 28.7704 | 29.1711 | 29.0219 | DUF3458;DUF3458_C;Peptidase_M1 | | | 0.006510881 | 0.99562466 | 0.002889633 | | 17 | 17 | 25.1 | 25.1 | 98.71 |
| E1WGG8 | 28.1307 | 28.0075 | 27.8724 | 27.8389 | 28.0881 | 28.0898 | Glyco_transf_20 | Trehalose-6-phosphate synthase {ECO:0000250\|UniProtKB:P31677} | | 0.006012866 | 0.9958929 | 0.0020504 | | 17 | 17 | 31.7 | 31.7 | 53.58 |
| A0A0H3NPR4 | 27.4724 | 27.7203 | 27.6866 | 27.5393 | 27.6593 | 27.6714 | PmbA_TldD | | | 0.011579982 | 0.99627383 | -0.003099442 | | 8 | 8 | 23.3 | 23.3 | 48.41 |
| A0A0H3NH69 | 26.91 | 26.9363 | 27.0439 | 26.9108 | 26.9189 | 27.0629 | OB_RNB;RNB;S1 | Exoribonuclease 2 {ECO:0000256\|HAMAP-Rule:MF_01036} | | 0.004176291 | 0.99637917 | 0.000818888 | | 10 | 10 | 17.5 | 17.5 | 72.44 |
| A0A0H3NFB5 | 27.9107 | 27.2997 | 27.9453 | 27.4302 | 27.9603 | 27.7788 | MGS | Methylglyoxal synthase {ECO:0000256\|HAMAP-Rule:MF_00549, ECO:0000256\|SAAS:SAAS01090967} | | 0.005650947 | 0.99688829 | 0.004500071 | | 4 | 4 | 26.3 | 26.3 | 16.99 |
